# A Unified De Novo Approach for Predicting the Structures of Ordered and Disordered Proteins

**DOI:** 10.1101/2020.01.30.925636

**Authors:** John J. Ferrie, E. James Petersson

## Abstract

As recognition of the abundance and relevance of intrinsically disordered proteins (IDPs) continues to grow, demand increases for methods that can rapidly predict the conformational ensembles populated by these proteins. To date, IDP simulations have largely been dominated by molecular dynamics (MD) simulations, which require significant compute times and/or complex hardware. Recent developments in MD have afforded methods capable of simulating both ordered and disordered proteins, yet to date accurate fold prediction from sequence has been dominated by Monte-Carlo (MC) based methods such as Rosetta. To overcome the limitations of current approaches in IDP simulation using Rosetta while maintaining its utility for modeling folded domains, we developed PyRosetta-based algorithms that allow for the accurate *de novo* prediction of proteins across all degrees of foldedness along with structural ensembles of disordered proteins. Our simulations have an accuracy comparable to state-of-the-art MD with vastly reduced computational demands.

## Introduction

Intrinsically disordered proteins (IDPs) and intrinsically disordered regions (IDRs) of proteins have garnered increasing attention due to growing recognition of their roles in cellular health and disease ^1,2^. These highly dynamic systems play crucial roles in signaling pathways, the function of the nuclear pore, lipid transport, membraneless organelles, and several pathologies ^1–3^. These functions are often related to structural transitions, where protein-protein interactions or ligand binding events facilitate conversion from a broad conformational ensemble to a much smaller number of states ^2^. The ability to model both the conformationally diverse states as well as the induced ordered structures of IDPs is essential for mechanistic understanding of their activity and for potential therapeutic development for IDP-related proteinopathies.

As interest in IDPs and IDRs has continued to grow, experimental and theoretical techniques originally developed for structured proteins have been adapted to accommodate these dynamic systems. Although NMR, fluorescence and scattering-based methods have all been effectively utilized to characterize unstructured systems, the production of representative structural ensembles for IDPs using computational methods has proven far more challenging ^4–14^. Many approaches have constructed ensembles by using experimental data to filter sets of potential structures generated from random-coil or Protein Data Bank (PDB) fragment libraries ^6,9,15^. Molecular dynamics (MD) force fields and sampling methods, such as replica-exchange, have been developed and optimized to predict IDP ensembles ^16–22^. Monte Carlo (MC) methods, such as those employed in CAMPARI, have demonstrated efficacy in modeling structural ensembles with and without data restraints, but protocols developed to date cannot generally accommodate both folded proteins and IDPs ^4,11,23,24^.

Our goal was to develop an architecture that effectively handles both ordered and disordered proteins and can generate accurate structural representations of both, *de novo*. Furthermore, we wanted to develop algorithms that do not suffer the hardware and computational time burdens associated with MD simulations. With these ideas in mind, we have turned our attention towards the Rosetta Modeling Suite. This platform has demonstrated extensive success in predicting structures for ordered systems through the use of fragment sampling in combination with a simulated annealing MC approach with limited gradient-based minimization ^25^. Additionally, Rosetta has been utilized to identify IDRs and to model IDRs and IDPs with the application of experimental constraints ^4,11,24,26–28^.

Here, we focus on adapting two algorithms: AbInitio, which can predict folded protein structure from sequence, and FloppyTail, which can produce structural ensembles of IDRs ^24,26,29–31^. We tested these methods’ ability to produce an accurate structural ensemble of an IDP, α-synuclein (αS), for which there exists a battery of experimental data. This allowed us to identify issues with Rosetta’s knowledge-based terms and fragment sampling approach, which we modified to improve the energy term accuracy and the fragment selection process. Lastly, we demonstrate that the incorporation of these improvements, along with additional changes to sampling schemes, affords two improved algorithms, AbInitioVO and FastFloppyTail, which in concert allow for the accurate prediction of structural representations of proteins across all degrees of foldedness.

## Results and Discussion

When considering a unified architecture, we noted that Rosetta-based algorithms such as AbInitio, designed for folded structures, and FloppyTail, designed for disordered structures, vary dramatically in their approaches ^24,29,31^. AbInitio utilizes simulated annealing, coarse-grained representations and knowledge-based scoring terms to sample nine-residue (9-mer) and three-residue (3-mer) fragments culled from the PDB ^29–31^. With each simulation stage, additional score terms are added to the MC score function and the number of fragment insertions tested between score events are reduced. Lastly, the simulated structures are processed via the Relax algorithm, which performs a series of gradient based minimizations ^32^. By comparison, FloppyTail also utilizes knowledge-based scoring and coarse-grained representations to perform initial 3-mer sampling, but follows this with additional sampling under the “score12” all-atom score function ^24^. In this later stage, fragment insertion steps are coupled to gradient-based minimization and sidechain rotamer sampling.

To determine the effective capacity of each method, structural predictions of αS were preformed using the AbInitio and FloppyTail methods, and were compared to 1) Förster resonance energy transfer (FRET) data from four different probe pairs, as well as inter-residue distances determined by other groups using 2) FRET, 3) electron transfer experiments, and 4) paramagnetic relaxation enhancement (PRE) measurements, to assess global ensemble accuracy ^4,5,8,11,33–35^. Furthermore, the accuracy of residue level information for αS ensembles was analyzed through comparison to 5) chemical shift (δ) data, 6) residual dipolar couplings (RDCs) and 7) NMR J-couplings ^8,36^. Detailed descriptions of the methods and equations used to compute comparisons of the structural ensembles to each piece of experimental data can be found in the Supporting Information (SI).

Of the two methods, FloppyTail significantly outperformed AbInitio in generating accurate disordered ensembles of αS, as might be expected (referred to as *FlopppyTail_score12* and *AbInitio*, respectively, in Tables S2-S4 and all Figures). The FloppyTail ensemble demonstrated an impressive degree of agreement with global descriptors of the protein’s overall topology, including radius of gyration (R_g_), FRET efficiency (E_FRET_), distance and PRE data (Figs. S4, S6, S8 and S25) However, upon comparison to residue-level information, we noted that the predicted C and Cβ chemical shifts (Fig. S54) as well as ^2^J_NCα_ J-couplings (Fig. S91) deviated from the experimental data (Tables S3-S4). These deviations are attributable to clear overpopulations of helical structure observed when plotting per residue helical propensity across all members of the resultant ensemble (Fig. S2). Indeed, overpopulation of helical architectures was not isolated to the FloppyTail simulation and was even more pronounced in the AbInitio simulated structures (Fig. S2). In addition, the AbInitio algorithm presented severe discrepancies in the overall R_g_ and other global parameters compared to experimental values (Figs S4, S6, S8). Overcompaction and overpopulation of helices are common problems in both MD and MC-based methods which have been optimized for ordered proteins ^16,18,20,21,37^.

### Employment of a Generalized Simulation Format

To overcome these barriers, a new method was generated in PyRosetta allowing for a comparison of 17 different combinations of score functions, sampling schemes and atomic representations ^38^. These are described in detail in the SI. We observed that over-compaction was most severe when using the centroid coarse-grained representation (*CenStd_PP* and *CenStd_PPFI* simulations, Table S2, Figs. S3, S5, S7 and S13-S14), but was alleviated via either subsequent all-atom sampling (all *SimAnn* simulations Table S2, Figs. S3, S5, S7 and S18-S21) or sampling with the specific centroid score function utilized by the FloppyTail algorithm (all *CenStd_Ext* simulations Table S2, Figs. S3, S5, S7 and S15-S16) ^24^. Therefore, we were able to determine that the “rg” score term, which is excluded from the score terms used in the centroid phase of the FloppyTail algorithm, was the likely source of overcompation.

Unlike overcompaction, overpopulation of helical structures appeared to occur independently of the atomic representation or score function and was solely an artifact of sampling. Indeed, overpopulation occurred in all cases where fragments were sampled (all *PPFI* and *PPFISC* simulations, Figs. S1-S2) except when the protein was reduced to a self-avoiding polymer representation (*VDW_PPFI* and *VDW_PPFISC* simulations, Figs. S1-S2). Therefore, we hypothesized that fragment selection, as opposed to the fragment sampling process *per se*, was the likely source of error.

Lastly, we observed that sidechain sampling had a relatively minimal impact in the absence of fragment sampling. Inspecting the all-atom simulations and simulated annealing simulations (*Beta*, *SimAnn* and *VDW* simulations, Table S2-S4) one sees that the impact of sidechain sampling is relatively small compared to the impact of sampling fragments (which includes backbone movement). Since sidechain packing in Rosetta accounts for a significant portion of the simulation time, we reasoned that simulation speed might be enhanced by reducing the frequency of sidechain sampling events.

### Increased Accuracy through Improved Fragment Selection

Fragment libraries in Rosetta are typically generated from 3-state secondary structure predictions based on protein primary sequences, and can be generated automatically using the Robetta server or manually using the FragmentPicker application ^39,40^. As predictions from most servers (Robetta, PsiPred, RaptorX) contained a significant helical probability corresponding to the amphipathic region of αS that forms α-helices when bound to membranes, all fragment selection strategies yielded structures with significant helical content (*FloppyTail_score12* and *FloppyTail_Quota* simulations, Fig. S2) ^41–44^. Recent studies by Bax and coworkers demonstrated that loops from PDB structures populated a similar Ramanchandran backbone dihedral space to that of IDPs ^45^. Therefore, we hypothesized that a fragment library composed exclusively of loops might allow us to use fragment insertion without overpopulating helical structures. Indeed, replacement of the initial fragment library with an all-loop library in the FloppyTail algorithm resulted in an ensemble which was devoid of extended secondary structure (*FloppyTail_Loops* simulation, Fig. S2). We were encouraged by the efficacy of this strategy, but recognized that fragment library construction required prior knowledge of a protein sequence’s degree of order. Therefore, as surrogate for this knowledge, we posited that disordered probability predictions from primary sequence could be used to control the probability of introducing loops as follows ^43,44^:

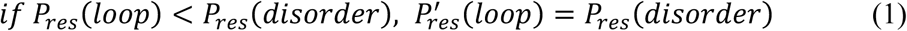

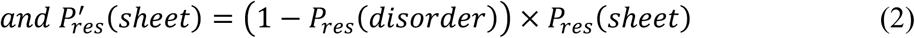

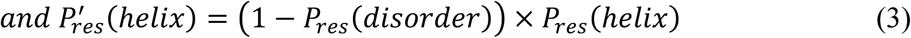

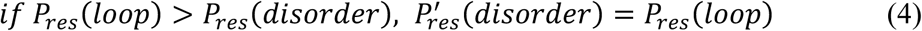

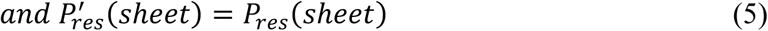

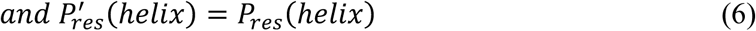

Here, P_res_(disorder) represents the predicted probability of a given residue to be disordered, while P_res_(loop), P_res_(helix), and P_res_(sheet) represent the probability of the residue being a loop, helix or sheet, respectively. Additionally, primed variables, P’_res_ (loop), P’_res_(helix), and P’_res_(sheet), represent the reweighted predictions. Finally, for residues with a loop probability greater than the predicted disorder probability, reweighed disorder probability will be set equal to the initial loop probability. Although many servers incorrectly characterized αS as being probabilistically ordered in at least a single stretch of residues, the RaptorX sever provided a correct prediction that the protein was entirely disordered ^43,44^. Therefore, we initially favored RaptorX for disordered probability predictions for αS with PsiPred predictions of secondary structure within the ordered sections, termed *best-reweighting*, although we also explored other combinations when looking at a larger collection of proteins (below). Gratifyingly, the resultant fragment library constructed using these reweighted loop predictions as inputs resulted in a highly accurate FloppyTail output ensemble (*FloppyTail*, Tables S2-S4) comprised of very few helices.

### Re-defining Knowledge Based Terms

Although improved fragment selection sufficiently reduced overpopulation of helices in the output from the FloppyTail method, the overcompaction identified in the AbInitio simulation required additional attention due to its reliance on knowledge-based terms. The final stages of the AbInitio simulation are reliant on the “rg” score term, whose energetic penalty is derived from the R_g_ of the structure ^31^. Although this facilitates the compaction necessary for the adoption of appropriate structures for folded proteins, this produces severely overcompacted IDPs and IDRs. Therefore, we tested replacement of this term with a score term comparing the R_g_ of the current simulated protein structure to that expected from polymer-scaling physics. Employing the following equations from Schuler and co-workers, we can predict an expected R_g_ from the hydrophobic or charge content of a protein with a given sequence:

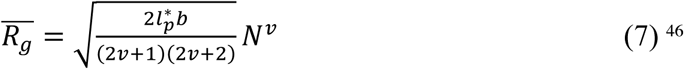

Here, 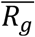 is the predicted mean R_g_ value determined from the persistence length (*l*_*p*_* = 0.53 nm) and the average distance between two Cα atoms (*b* = 0.38 nm), which have been previously determined, along with the number of residues in the sequence, *N*, and the scaling factor, *v*. Determination of the value used for the scaling factor is based again on the predicted disordered probability as detailed in the SI. In short, regions predicted to be ordered are assigned a scaling factor *ν* = 0.33, while the scaling factor for disordered regions is determined via Eqs. S13-S16 depending on whether net charge or hydrophobicity serves as the dominating characteristic. To incorporate the predicted 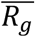 within an energy term, we elected to craft a potential from a general version of a self-avoiding walk (SAW) probability distribution, previously employed by Zheng, Best, Schuler and coworkers, that accommodates changes in the scaling exponent ^47^.

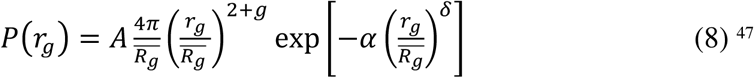

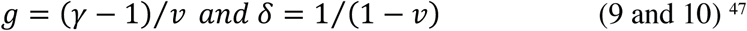

Above, *r*_*g*_ represents the Rg of the structure being assessed by the scoring function and 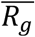 again represents the predicted mean R_g_. The constants *g* and *δ* in Eq. 8 are defined in the subsequent equations Eq. 9 and 10. The constant *γ* (1.1615) has been previously estimated for proteins, while the constants *A* and *α* are determined for a given 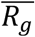 and *v* pair based on the normalization conditions detailed in the SI (Eqs. SX and SX). This function provides a broad distribution of radii of gyration while at the same time preventing structures from becoming overly compact. The per segment score term is defined as:

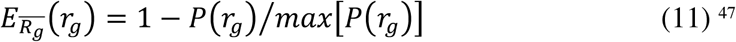

**Figure 1.**
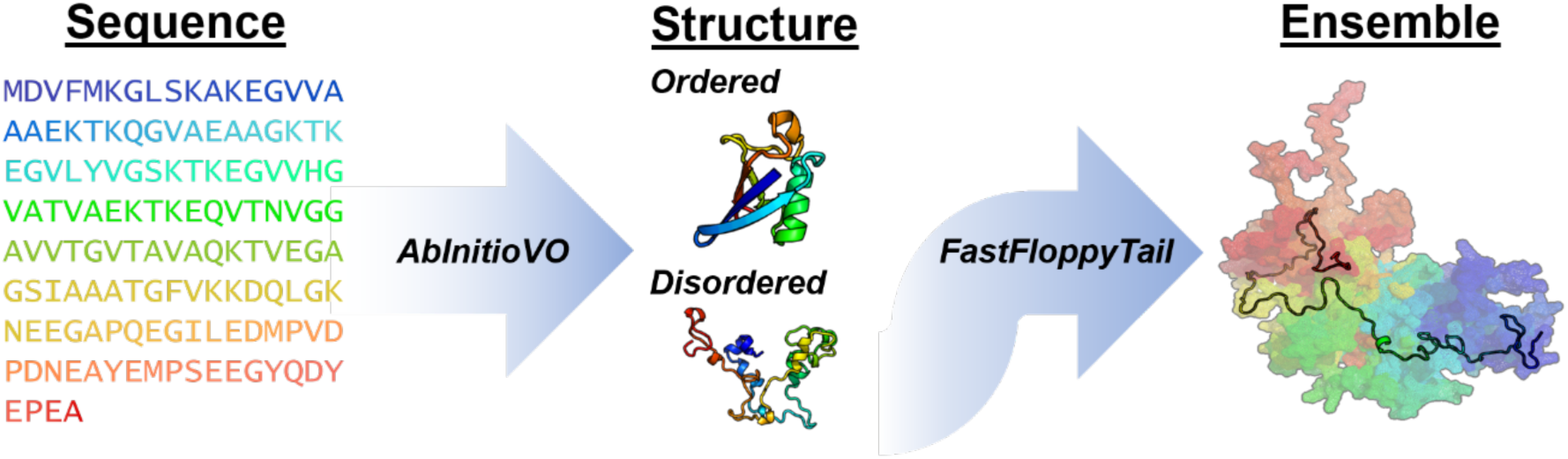
Schematic depicting how improved versions of AbInitio (AbInitioVO) and FloppyTail (FastFloppyTail) would allow for accurate structural prediction of disordered and ordered proteins from sequence.

Through this form, the energy value scales from zero to one for any 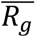, allowing the weight of this score term within our overall score function to determine the depth of the potential. Subsequently, we identified the optimal weighting value as that which maximized the impact of the score term on the average radius of gyration while minimizing the restriction on the conformational diversity (Fig. S104). Combination of this novel scoring approach with the previously described fragment selection strategy results in the new simulation termed AbInitioVO, or AbInitio Variable Order.

### Acceleration and Improved Accuracy via Adjustment to Sampling

As previously stated, results from the generalized simulation suggested that a reduction of sidechain sampling could afford an accelerated simulation with comparable accuracy. Therefore, we removed sidechain sampling from one of the interior loops of the FloppyTail protocol, reducing the sampling frequency 15-fold. This resulted in a ~10-fold reduction in compute time and produced an ensemble (*FloppyTail_Rot* simulation, Tables S2-S4) that was nearly identical to the *FloppyTail* ensemble across all experimental comparisons. Additionally, unlike a traditional MC approach, at several stages of the FloppyTail simulation, the structure is returned to the lowest energy structure encountered in the search up until that point, restricting sampling to states near minima ^24^. Serendipitously, we discovered that returning to the lowest energy conformer more frequently further improved agreement with the experimental data, which in combination with our improved fragment selection scheme resulted in the algorithm now called FastFloppyTail. Finally, we employed the Relax algorithm, commonly used after the AbInitio algorithm, to the outputs from FastFloppyTail simulations in an algorithm termed FastFloppyTail-Relax, and again were met with an additional improvement in the overall agreement with experimental data (*FloppyTail_Relax*, *FloppyTail_Rot_Relax* and *FastFloppyTail_Relax* simulations, Tables S2-S4) ^29,30,48^.

**Figure 2.**
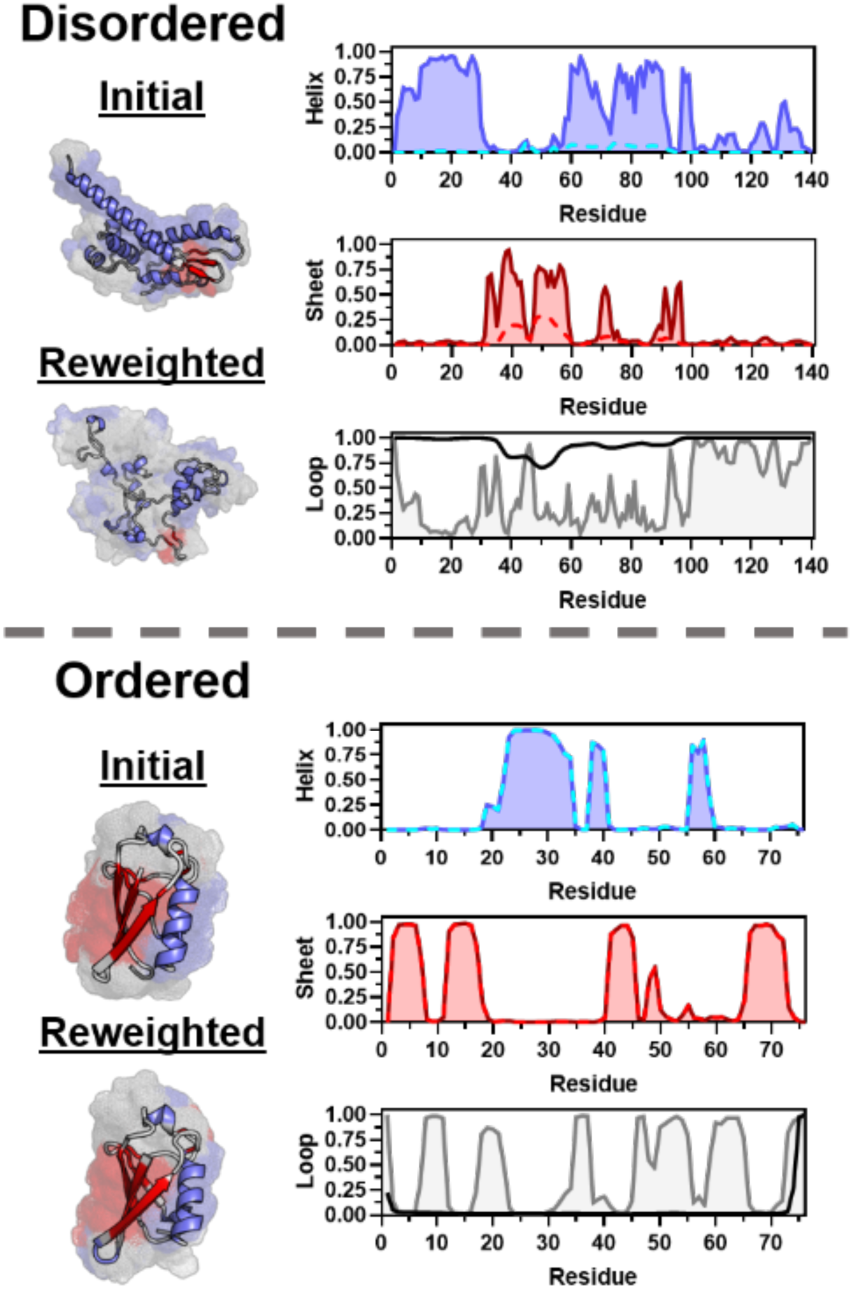
Impact of reweighting secondary structure probabilities during fragment selection on ordered and disordered protein structural predictions. For αS (Disordered) and ubiquitin (Ordered), structural outputs from simulations (left) utilizing initial secondary structure predictions from PsiPred (Initial) and reweighted predictions according to disorder predictions from RaptorX (Reweighted) ^43,44^. Structural images show a single structure as a cartoon with ten representative structures shown in surface rendering to illustrate helical (blue), sheet (red) and loop (grey) population. Plots (right) depict the initial secondary structure predictions from PsiPred (solid blue, red and grey) along with reweighted predictions (dashed cyan and red) based on the RaptorX disordered probability (solid black) ^43,44^.

### Accurate De Novo Prediction of Disordered and Ordered Proteins

To confirm the generalizability of the AbInitioVO and FastFloppyTail protocols, a set of 25 proteins containing a variety of different secondary structural elements and spanning varying levels of order/disorder was selected. For each AbInitioVO simulation, disordered probability predictions were performed based on the input sequence using RaptorX and fragment libraries were assembled using a best fragment selection protocol based on a loop-reweighted secondary structure prediction from PsiPred, using Eqs. 1–6. The resultant structural predictions from AbIntioVO were compared to structures generated using the standard AbInitio protocol (Table S5). For ordered and partially-ordered proteins, prediction accuracy was confirmed via comparison to x-ray crystal structures and NMR structures, while disordered proteins were inspected for agreement with the experimentally determined R_g_ of each protein. As exemplified by ubiquitin, histone H1 and αS in Figure 3, AbInitioVO can assign the correct secondary structural elements and level of disorder across many scenarios and performs comparably to AbInitio in folded regions. Additionally, we find that the resultant average R_g_ values for disordered proteins from AbInitioVO are more consistent with the experimental R_g_ values than those from AbInitio, as shown for αS. These findings are generalizable to the full set of results from all 25 proteins (Table S5, Figs S105-S107) and support the notion that AbInitioVO is able to correctly predict the folded portions of ordered and partially-ordered proteins, while identifying disordered regions with the correct secondary structure and overall size.

**Figure 3.**
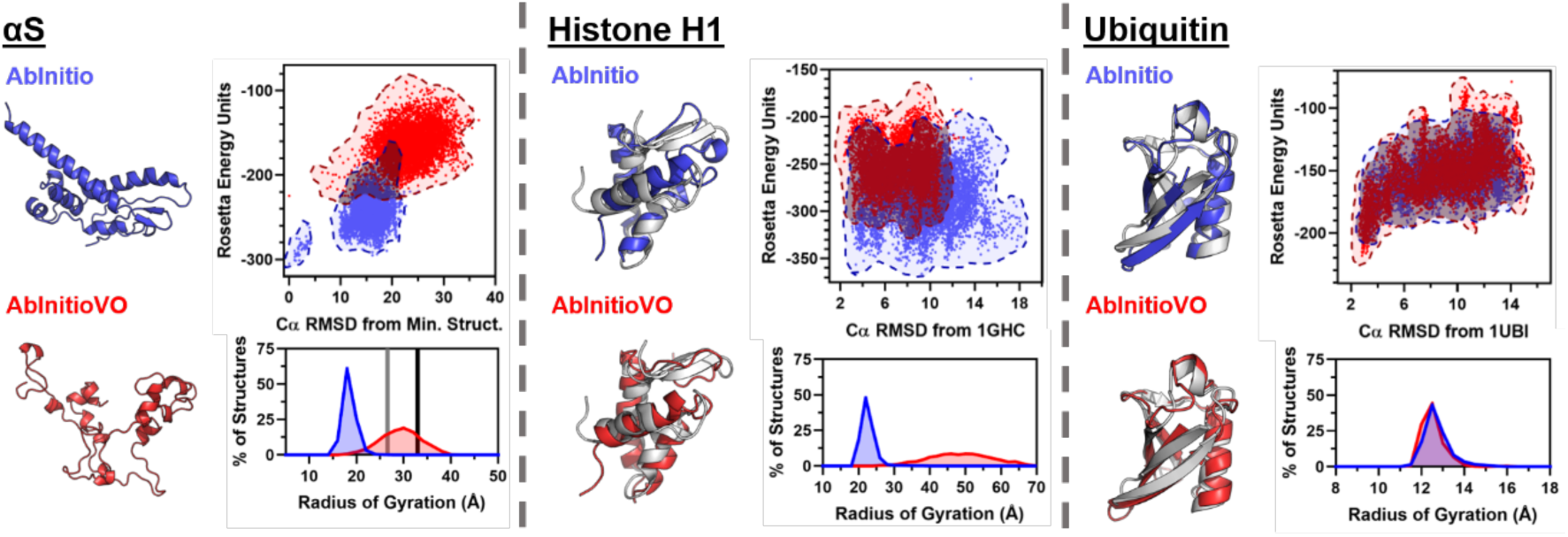
Comparison of secondary structure and disordered region predictions from AbInitio and AbInitioVO. Example disordered (αS), partially-disordered (Histone H1) and ordered (Ubiquitin) protein predictions. Each panel contains (left) single structural predictions from AbInitio (top left) and AbInitioVO (bottom left) with the lowest energy structure (αS) or lowest RMSD structure as compared to the PDB structure (1GHC for histone H1 and 1UBI for ubiquitin) along with folding funnels (top right) showing energy vs RMSD for all output structures and a histogram of output structure R_g_ values (bottom right). The experimental R_g_ values for αS from NMR (grey) and X-ray scattering (black) experiments are shown for comparison ^50,51^.

**Figure 4.**
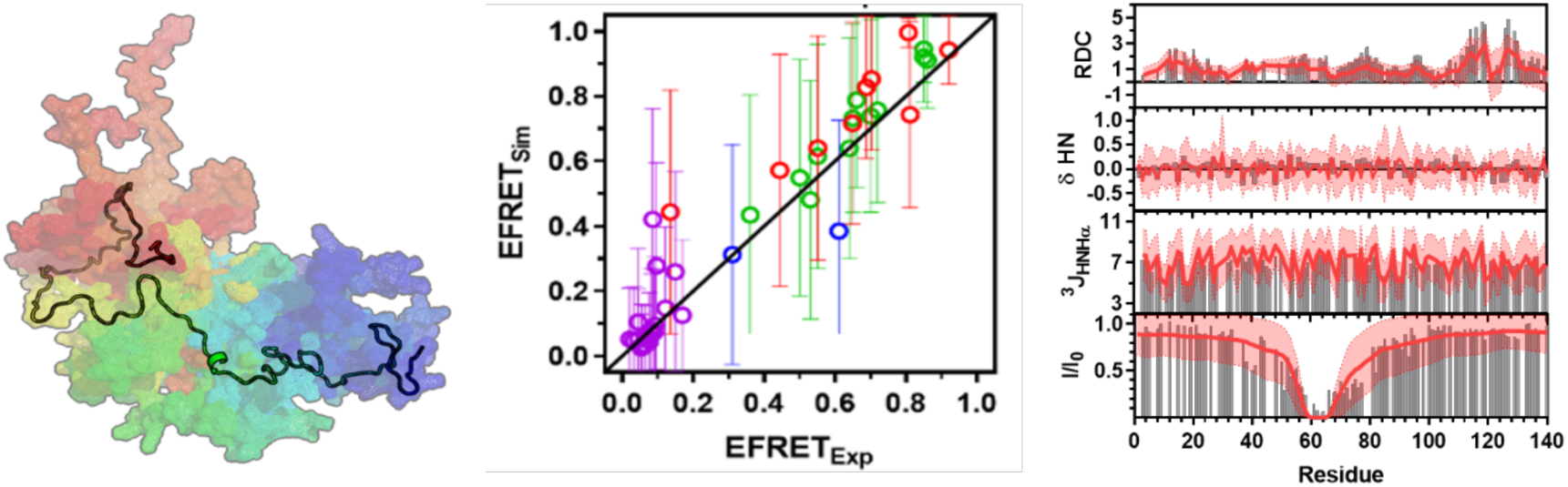
Comparison of simulated FastFloppyTail αS ensemble to global and residue-level experimental data. (Left) The single lowest energy structure (cartoon) overlaid on the ten lowest energy structures (surface) of αS from FastFloppyTail simulation. (Middle) Comparison of E_FRET_ values computed from output structures (EFRET_Sim_) with experimental data (EFRET_Exp_) to assess global accuracy of the ensemble with error bars showing the standard deviation. (Right) Comparison of (from top) residual dipolar couplings (RDC, Hz), amide proton secondary chemical shifts (δ HN, ppm), J-couplings (^3^J_HNHα_, Hz) and PRE values (I/I_0_, unitless) computed from the output structures (red line) with experimental data (grey bars) to illustrate residue level accuracy. Lines represent the average value computed on a per residue basis across a set of 1000 structures, while shaded regions show the computed standard deviation.

Following correct identification by AbInitioVO, IDRs can subsequently be simulated using the FastFloppyTail algorithm to generate accurate structural ensembles. Although AbInitioVO can accurately predict global structural features, accurate ensemble generation requires subsequent sampling as disordered regions are still populated with multiple short helical fragments which can be observed in the structural output for αS from AbInitioVO (Fig. 3). The effectiveness of FastFloppyTail was tested against the 6 IDPs in the 25-protein set, which were compared to available experimental data (Table S6). Figure 5 shows the per residue helical fraction determined from the simulated structures compared to the helical abundance computed from chemical shifts.

**Figure 5.**
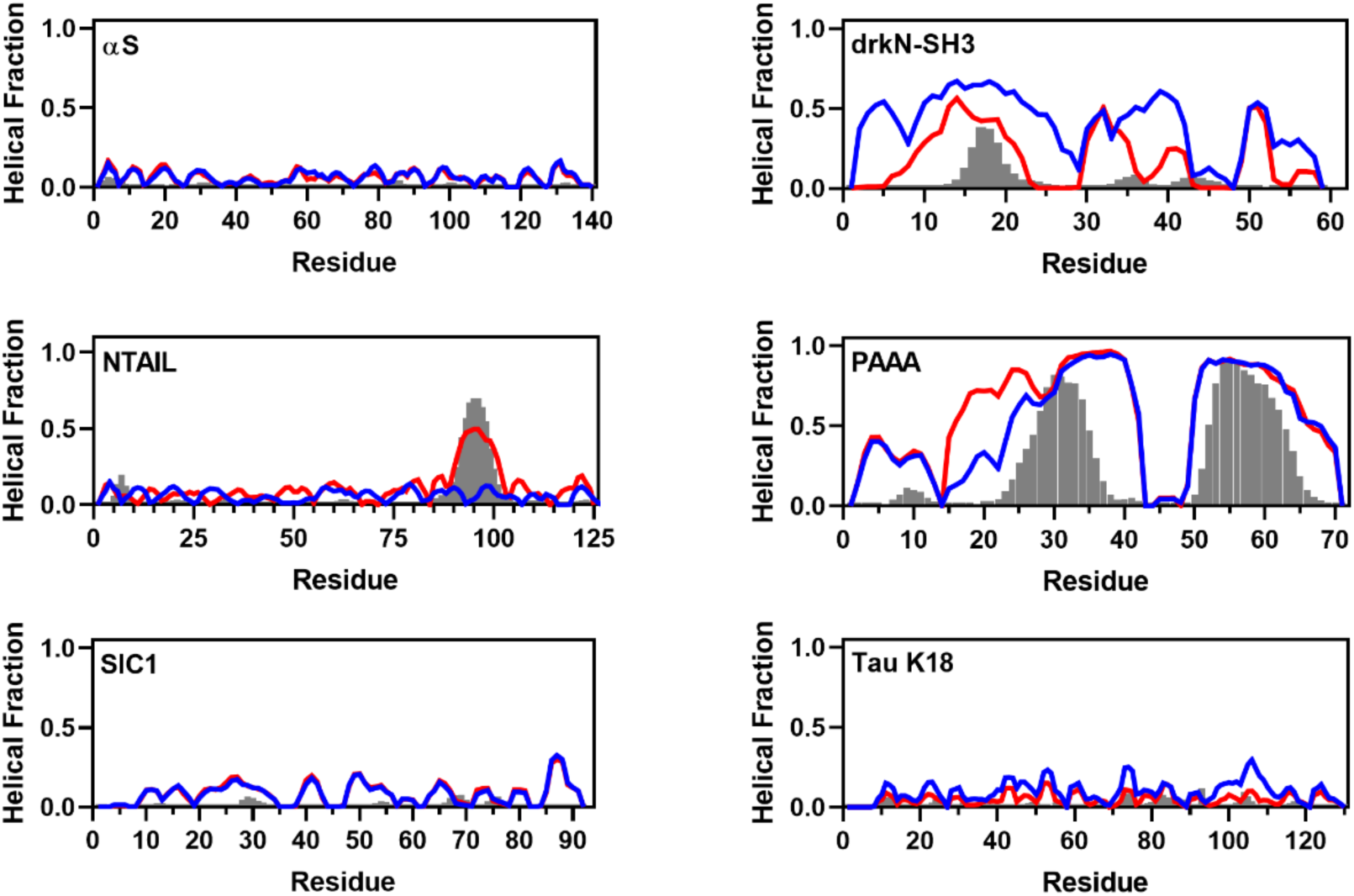
Comparison of per residue helical fraction between FastFloppyTail simulated ensembles and experimentally derived data. Plots of the helical fraction for α-synuclein, drkN-SH3, NTAIL, PAAA, SIC1 and tau K18 computed by DSSP from simulated ensembles as decribed in SI. Simulations were performed with fragments derived from *best-reweighting* (blue) or *self-reweighting* (red) schemes and are compared to helical fractions computed from chemical shift data using D2D (grey bars) as described in SI ^52^.

Unsurprisingly, we find that the ensembles from FastFloppyTail are highly sensitive to the secondary structure predictions and disordered probability predictions, encouraging us to test additional methods of reweighting loops probabilities. The first was identical to the approach used for the AbInitioVO simulations, using secondary structure predictions from PsiPred and disordered predictions from RaptorX, termed the *best-reweighting* scheme. The second method, here termed *self-reweighting*, used disordered predictions from PsiPred to weight secondary structure predictions from PsiPred and disordered predictions from RaptorX to weight secondary structure predictions from RaptorX, which are each used to select fragments for 50% of the final fragment library. By selecting fragments using self-reweighted predictions, as shown in Figure 5, our resultant ensembles show better overall agreement in predicting locations with transient helices as well as reasonable matches with quantitative helical propensities. Comparison of the ensembles to the available experimental data (Table S6, Figs. S165-S220) reveals a striking degree of agreement, comparable to state-of-the-art MD simulations ^18^. We find that the average R_g_ values from self-reweighted FastFloppyTail-Relax simulations were within 5 Å of the experimentally determined values across all tested proteins, highlighting agreement with global structural features of the simulated proteins (Table 1). Additionally, comparisons to NMR RDC measurements (Table 1), which capture local conformational sampling, demonstrate a high degree of agreement, with all Q-values falling below 0.5. To further assess residue-level accuracy, the overall RMSD compared to NMR chemical shift data from this approach is 0.72, which is comparable to 0.63 from previous MD methods for the same set of proteins ^18^. These, as well as comparisons to NMR J-coupling values and PRE and FRET experiments (Table S6), demonstrate that the resultant ensembles from the self-reweighted FastFloppyTail-Relax approach accurately capture both residue-level features, such as backbone torsion angle propensities, as well as more global structural features.

**Table 1.**
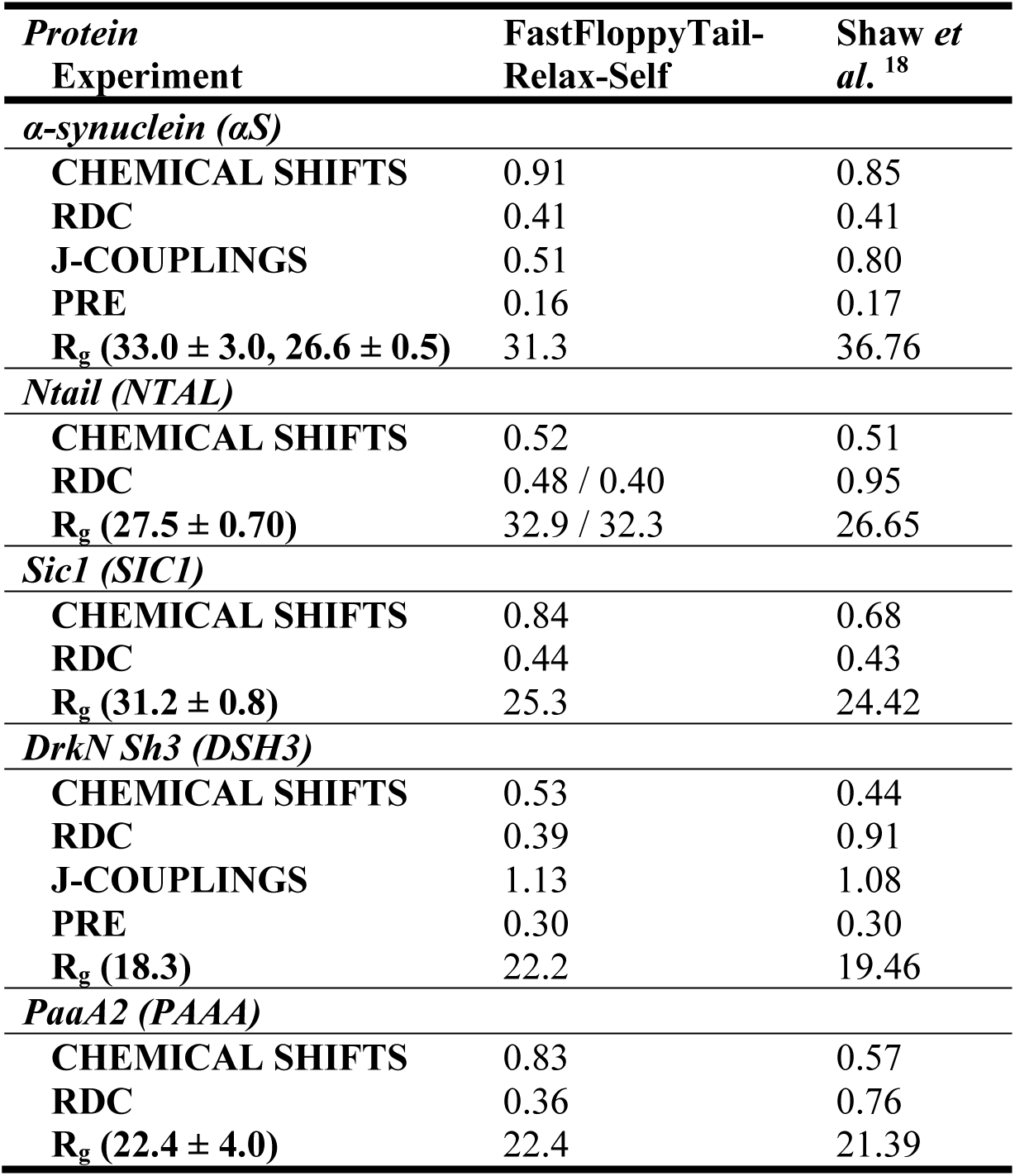
Comparison of simulated data with experimental values for FastFloppyTail-Relax outputs using self-reweighting with MD results from Shaw *et al*. ^18^. Chemical shift (Hz) and J-coupling (ppm) values are reported as RMSD while RDC values are reported as Q-values (unitless) and R_g_ values are reported in angstroms. All values for comparisons were averages computed across all 1000 outputs.

Given the nature of these algorithms, optimal predictions require accurate predictions of both secondary structure and disordered propensity. However, we find that the built-in redundancy of our reweighting scheme improves our ability to capture disordered proteins. For example, although tau K18 is not predicted by RaptorX to be disordered, it is predicted to be comprised exclusively of loops, allowing for accurate structural prediction. This indicates that our reweighting strategy only requires that either the disordered probability or the loop probability be correct to accurately predict disordered proteins.

Lastly, we find that in the particularly interesting case of the drkN SH3 protein, which NMR experiments show to exist in a 1:1 ratio of ordered and disordered states, our combined method accurately captures the full range of states sampled by a protein from a single set of predictions ^49^. Unlike previously published approaches that focus exclusively on ordered or disordered protein prediction, application of AbInitioVO results in the beta-sheet rich SH3 domain fold (Fig. S160) for the supplied sequence. The ensemble generated by application of FastFloppyTail for this sequence matches well with NMR data acquired for the disordered population of drkN (Table S6, Figs S190-S197), including observation of a helical stretch in ensemble structures that lack the SH3 fold (Fig. 5). Successful prediction of protein ensembles with these types of transient features, which are distinct from proteins with IDRs that contain consistent ordered and disordered domains, underscores the efficacy of the AbInitioVO/FastFloppyTail-Relax approach to capture many aspects of protein structure across a wide array of folds a degree of order.

## Conclusions

Our analysis shows that the refined AbInitioVO and FastFloppyTail algorithms are capable of accurately predicting the structure of proteins across a wide array of folds and degrees of order. Both methods rely on the use of secondary structure and disordered probability predictions to facilitate tailored fragment sampling approaches. This reduces the necessary sampling time, and necessary hardware, and improves the efficacy of predicting both ordered and disordered proteins. We have demonstrated that the AbInitio method within the Rosetta Modeling Suite, which has long been recognized for its ability to accurately predict the structure of ordered proteins given their primary sequence, can be amended to allow for the accurate prediction of disordered (IDP) and partially ordered systems (both IDRs and transient systems) without any noticeable reduction in efficacy for folded proteins. Furthermore, following initial structural prediction in AbInitioVO, conformational ensembles of disordered systems can be expanded using fragment-based approaches like FastFloppyTail-Relax with comparable accuracy to state-of-the-art MD methods, but with dramatically shortened compute times. The remaining limitations of these approaches are highly tethered to limitations in the accuracy of the initial secondary structure and disorder predictions from servers, indicating that further improvements in RaptorX, PsiPred, or similar servers can improve the accuracy of AbInitioVO/ FastFloppyTail-Relax. Similar to CAMPARI, we are able to demonstrate that both ordered and disordered domain sampling is fully accommodated. However, here this is accomplished *de novo* without the need for starting structures. Ultimately, we hypothesize that these algorithms can be useful in a variety of contexts. By rapidly providing accurate ensembles, our algorithms allow disordered ensembles to be further refined through the application of experimental data without burdensome computing times. Experimental data could include NMR, crosslinking, or FRET measurements supplying domain-level or local information. The AbInitioVO/FastFloppyTail-Relax outputs can also be used as starting points for MD trajectories serving as surrogates or supplements to enhances sampling schemes. Moreover, since our MC simulations enable an efficient sampling of conformational space, subsequent MD trajectories can be used for studying aspects such as mechanisms of structural transitions or interactions with solvent molecules and membranes. In summary, the algorithms presented herein demonstrate, for the first time, that simulations that capture the structures and conformational distributions of both ordered and disordered proteins can be accomplished with limited computational requirements.

## Methods

All simulations were performed using PyRosetta, where versions of AbInitio and FloppyTail were drafted in PyRosetta based on original code ^38^. For the Generalized Simulation, AbInitioVO, FastFloppyTail and all derivatives, the nomenclature and specifics of each simulation are detailed in the SI. In short, for the generalized simulation, sampling consisted of six sampling stages where the temperature used for assessing the Metropolis criteria is decreased at each stage. Sampling consists of φ/ψ backbone torsion angle sampling, fragment insertion and/or sidechain rotamer sampling (indicated by PP, FI and SC suffixes in the generalized simulation names, respectively). Simulations were also tested under different score function and atomic representations utilizing the standard centroid scoring procedure in Rosetta, the standard full-atom score function in Rosetta, or the scoring procedure utilized by AbInitio (CenStd, Beta, and SimAnn prefixes respectively) ^29–31,48^. Additionally, polymer-like self-avoiding walk simulations were also performed where the score function was exclusively comprised of a repulsive van der Waals energy term (VDW prefix) ^48^. The AbInitio and FloppyTail simulations follow the previously described algorithm formats with the changes indicated in the main text and SI. Comparison of all simulated structures and structural ensembles are described in detail for each measurement in the SI.

## Data Availability

All simulation and analysis scripts as well as demonstration simulations are available on Github: https://github.com/jferrie3/AbInitioVO-and-FastFloppyTail.

## Acknowledgments

We thank David Eliezer for sharing the chemical shift, RDC and PRE data for αS, Christopher Dobson and Michele Vendruscolo for sharing PRE data for αS and Markus Zweckstetter for sharing RDC data. We also thank Abhinav Nath for advice and for reading of the manuscript and Wenwei Zheng for sharing his code related to computing polymer-scaling equations. We are grateful to David Baker and his laboratory for guidance of JJF during his summer internship. This research was supported by the National Institutes of Health (NIH R01-NS081033 and R01-NS103873 to EJP). JJF thanks the National Science Foundation (NSF DGE-1321851) and the Parkinson’s Disease Foundation (PF-RVSA-SFW-1754) for fellowship support.

## Author Contributions

JJF performed all computational simulations and comparisons to experimental data. Both JJF and EJP contributed to writing the manuscript.

## Competing Interests

The authors declare that they have no conflict of interest.

## Materials & Correspondence

All simulation and analysis scripts are available upon request from EJP or JJF.

## Notes

https://github.com/jferrie3/AbInitioVO-and-FastFloppyTail.

