## Supplementary Information for "A Unified De Novo Approach for Predicting the Structures of Ordered and Disordered Proteins"

J. J. Ferrie* and E. J. Petersson*

*Department of Chemistry, University of Pennsylvania, 231 South 34th Street, Philadelphia, Pennsylvania 19104-6323, USA*

### Table of Contents

General Information ....S2

Computational Resources ....S2

Generalized Simulation Format ....S2

Construction of Fragment Libraries ....S3

Calculation of Data from Ensembles....S4

FRET Data....S4

Distance Data....S4

PRE Data ....S4

Chemical Shift Data....S5

Residual Dipolar Couplings ....S5

J-Couplings....S5

Percent Helicity....S6

Comparison of αS Ensembles from Generalized Simulation to Experimental Data ....S7

Summary of Generalized Simulations of αS....S7

Comparison of αS Per Residue Percent Helicity ....S11

Comparison with αS Radius of Gyration Data....S13

Comparison with αS FRET Data....S15

Comparison with αS Distance Data ....S17

Comparison with αS PRE Data ....S19

Comparison with αS Chemical Shift Data ....S48

Comparison with αS Residual Dipolar Coupling Data ....S77

Comparison with αS J-Coupling Data....S85

Construction of the New Rg Score Term.... S114

Implementation of the New Rg Score Term in PyRosetta .... S115

Determination of the Optimal Weight for the New Rg Score Term .................................................. S116
Comparison of AbInitio and AbInitioVO Simulations to Experimental Data ......................................... S117
Simulated Proteins and Abbreviations ............................................................................................. S117
Summary of Simulated Proteins ....................................................................................................... S118
Comparison of Radii of Gyration...................................................................................................... S119
Comparison of Folding Funnels ....................................................................................................... S122
Comparisons of Folded Domain Structures of Ordered and Partially-Ordered Proteins.................. S134
Structural Comparisons of Disordered Domains of Partially-Ordered and Disordered Proteins....... S140
Comparison of FastFloppyTail and FastFloppyTail-Relax Simulations to Experimental Data ................ S145
Summary of Simulated Proteins ....................................................................................................... S145
Comparisons of αS Data................................................................................................................... S147
Comparisons of DSH3 Data.............................................................................................................. S161
Comparisons of NTAL Data .............................................................................................................. S168
Comparisons of PAAA Data............................................................................................................... S173
Comparisons of SIC1 Data................................................................................................................ S178
Comparisons of TAUK Data............................................................................................................... S183
References:........................................................................................................................................ S188

### General Information

**Computational Resources:** All simulations were performed on the University of Pennsylvania School of Arts and Sciences General Purpose Cluster (GPC). Each of the four GPC compute nodes contains 24 cores at 2.6 Ghz, 256 GB RAM, 1 GbE Networking, 56 Gb Infiniband and 3.6 TB OS/Scratch storage. All times reported herein were from 140-residue αS. On average, Full-atom Generalized Simulations required ~3 hours per structure while simulations using exclusively Centroid coarse-graining required ~30 minutes per structure. Simulations using both coarse-grained and all-atom molecular representations (SimAnn) took ~1.5 hours per structure and the previously reported FloppyTail simulation method took ~30 minutes per structure whereas FastFloppyTail simulation method took ~3 minutes per structure. The AbInitio and AbInitioVO algorithms both required ~45 minutes per structure and Relaxes took ~3 minutes a structure.

**Generalized Simulation Format:** We chose to sample the effects of full-atom and centroid AtomTypes on the simulation, in an effort to assess the possibility of using the coarse-grained method to speed up the simulation during initial sampling steps as is frequently done in Rosetta. Additionally, we assessed the utility of different score terms and focused on six different score weights (VDW: exclusively repulsive van der Waals term "fa_rep", CenStd: "cen_std" score term weights, CenStdExt: "cen_std" score term weights adding "rama", "cenpack", "hbond_lr_bb" and "hbond_sr_bb" all at weights of 1.0, Beta: "beta_nov15" score term weights, SimAnn: simulated annealing where additional score terms were included as sampling progressed utilizing "score0", "score1", "score2", "score3" and "beta_nov15", CenNath: "vdw", "rama", "pair", "env", "hbond_lr_bb" and "hbond_sr_bb" all at weights of 1.0, and CenOpt: identical to the SimAnn centroid score functions with “rg” at a weight of 0 and “hbond_lr_bb” and “hbond_sr_bb” at weights of 1.0).[1] Finally, we utilized three different types of Movers within each of the different AtomType/ScoreFunction combinations (PP: φ/ψ torsion angle changes using Small/Shear Movers, FI: fragment insertion, and SC: side-chain rotamer optimization using the PackRotamersMover). The simulation nomenclature is in the form AtomType_Scoring_Sampling. For example, FA_Beta_PPFI utilizes the full-atom AtomType, the "beta_nov15" scoring term weights for the ScoreFunction and application of both phi/psi sampling and fragment insertion. To assess the impact of each of these parameters on the resultant ensemble, a base script was devised which could be easily altered to analyze variable effects while keeping the number of Movers applied to the protein backbone over the course of the simulation constant. In brief, the method consisted primarily of RandomMovers, which select and apply a single move from a detailed set of movers comprised of SmallMovers and ShearMovers (apply φ/ψ torsion angle changes), as well as MinMovers (perform gradient-descent minimization to locate local minima). Fragment insertion in the form of the ClassicFragmentMover was added directly to RandomMovers for both 9-mer and 3-mer fragments. Side-chain sampling was not added directly to the RandomMover, but was applied after a move from the RandomMover. The number of moves applied by the RandomMover prior to application of the Metropolis-Hastings acceptance criteria were decreased over the course of the simulation. The temperature (specified kT) value was also decreased. Lastly, the structure was set to the lowest energy structure found after each sampling stage and at the end of the simulation. For each simulation the resultant ensembles consisted of ~1000 structures.

**Construction of Fragment Libraries:** The Robetta server was used to generate the initial fragment library for testing all variants of the “Generalized Simulation” along with “FloppyTail” and “FloppyTail_ref2015”.[2] Production of the custom fragment libraries used were prepared using the FragmentPicker application in Rosetta [3]. All fragment libraries contained 200 fragments. The "FloppyTail_Quota" library was prepared using secondary structure probability predictions from the primary sequence of α-synuclein (αS) using the Jufo, PsiPred and RaptorX servers along with the quota protocol of the FragmentPicker application.[4-6] The probability of selecting fragments based on Jufo, PsiPred and RaptorX predictions were set to 0.33, 0.33 and 0.34 respectively with priorities for SecondarySimilarity of 250, 300 and 350 with weights of 1.0 for all and priorities ofor RamaScore of 150 with a weight of 2.0 for all predictions in the input weight file. The input weight file also contained ProfileScoreL1 with priority and weight of 200 and 2.0 respectively based on a sequence profile from PSI-BLAST. The "FloppyTail_Loop" library was prepared by exchanging the secondary structure predictions with a single manually crafted prediction input that contains 1.0 loop probabilities at every residue of the sequence. All “FloppyTail” simulations not previously addressed, including all FastFloppyTail simulations in the Generalized Simulation section, utilized a quota-style fragment library using the same inputs as detailed above, where the disordered probability prediction was used to re-weight loop contributions as described in the main text based on a disordered probability prediction from RaptorX.

AbInitio and AbInitioVO simulations were generated using the FragmentPicker following the best protocol, using only PsiPred secondary structure predictions as inputs. AbInitioVO PsiPred predictions were loop reweighted using disordered probability predictions from RaptorX, termed *best-reweighting*, as described in the main text. FastFloppyTail simulations of αS and other disordered proteins were performed using libraries generated using the quota protocol of the FragmentPicker application and the *best-reweighting* approach, identical to that employed for AbInitioVO simulations. This contained SecondarySimilary (priority: 350, weight 1.0), RamaScore (priority: 150, weight: 2.0) and ProfileScoreL1 (priority: 200, weight: 2.0) as inputs. Libraries were also generated using *self-reweighting*, where secondary structure predictions from PsiPred and RaptorX were reweighted based on disordered probability predictions from PsiPred and RaptorX respectively. Additionally, all residues were set to probabilities of being 100% loop unless, following reweighting, the residue helix or loop probability was > 0.75 and at least 3 or 5 consecutive residues were identified as sheets or helices respectively. The input file values were adjusted accordingly (PsiPred:: SecondarySimilary: priority: 300 weight: 1.0, RamaScore: priority: 150 weight: 2.0, RaptorX:: SecondarySimilary: priority: 350 weight: 1.0, RamaScore: priority: 150 weight: 2.0).

**Calculation of Data from Ensembles**

**FRET Data:** FRET efficiencies ($E_{FRET}$s) for each residue pair were computed for each individual structure and averaged across all structures in a given ensemble to determine the average $E_{FRET}$ of the ensemble. This was performed through application of the Förster Equation:

$$E_{FRET} = \frac{1}{1+(r/R_0)^6} \qquad \text{(Eqn. S1)}^{7}$$

which is dependent on the interfluorophore distance, *r*, and the Förster distance ($R_0$). $R_0$ values were taken from the references listed with figures referring to FRET data. For expedience, we approximated *r* as the distance between the Cα atoms of the labeled residues.

**Distance Data**: Distances for each residue pair were determined from the distances between the Cα atoms of each residue for each structure in each ensemble and were averaged over all members of the ensemble.

**PRE Data:** Paramagnetic relaxation enhancement (PRE) values ($\Gamma_2$) were computed and converted to the experimentally observed value ($I_{ox}/I_{red}$) from each structure in the ensemble using the formulas:

$$\Gamma_2 = \left[\frac{K}{r^6}\left(4\tau_c + \frac{3\tau_c}{1+\omega^2{\tau_c}^2}\right)\right] \qquad \text{(Eqn. S2)}$$

$$\frac{I_{ox}}{I_{red}} = \frac{R2_{red}\, exp(-\Gamma_2 t)}{R2_{red}+\Gamma_2} \qquad \text{(Eqn. S3)}$$

previously detailed by Sung *et al.* [8] Here, *K* is a constant that describes the spin properties for the nitroxide radical (1.23 x $10^{-32}$ $cm^6$ $s^{-2}$), $\tau_c$ is the correlation time for the electron-nuclear interaction vector (4 ns) and $\omega/2\pi$ is the Larmour frequency of an amide proton (computed in all cases for a 700 MHz field), $R2_{red}$ is the transverse relaxation rate in the diamagnetic state (set to 4 $s^{-1}$) and *t* is the total INEPT evolution time of the HSQC experiment, which was 10 ms for comparisons to data from Dedmon *et al.* [9] (as previously done by Piana *et al.* [10]) and 4 ms for comparisons to data from Sung *et al.* [8].

**Chemical Shift Data:** The amide proton (HN), amide nitrogen (N), carbonyl carbon (C), α-carbon (Cα), α-carbon proton (HA), and β-carbon (Cβ) chemical shifts were all computed using the SPARTA+ package developed by Shen and Bax.[11] The chemical shift values were computed for each structure and averages were computed uniformly across a given ensemble. Secondary chemical shifts were calculated by subtracting random coil chemical shifts for each residue as assigned by SPARTA+ [11].

**Residual Dipolar Couplings:** All residual dipolar couplings (RDCs) were computed using PALES [12]. For each structure in a given ensemble, the RDC for a given residue was computed using the "*-bestFit*" flag along suppling input data for comparison. Calculations were performed on 15 residue segments where the residue of interest occupied the central position, as previously done by Piana *et al.* [10]. For the first and last seven residues in a given protein, the N- and C-terminal 15 residue segments were used. The averages were computed for the 1000 lowest energy structures in each ensemble.

**J-Couplings:** The NMR *J*-couplings were calculated using fitted Karplus equations previously utilized by Shen and Bax from backbone φ/ψ dihedral angles [13]. For each structure, φ and ψ were computed in DSSP [14]. As with other values, the *J*-coupling was computed for each residue in each structure and averaged across all members of the ensemble. The Karplus equations used to compute each *J*-coupling are detailed below:

$$^3J_{HN-HA} = 7.97\cos^2(\varphi - 60°) - 1.26\cos(\varphi - 60°) + 0.63 \quad \text{(Eqn. S4)}^{15}$$

$$^1J_{C\alpha-H\alpha} = A_{RC} + 1.4\sin(\psi + 138°) - 4.1\cos(2[\psi + 138°]) + 1.7\cos(2[\varphi + 60°]) \quad \text{(Eqn. S5)}^{16}$$

$$^1J_{C\alpha-N} = 9.5098 - 0.9799\cos(\psi) + 1.7040\cos^2(\psi) \quad \text{(Eqn. S6)}^{17}$$

$$^2J_{C\alpha-N} = C - 1.5176\cos(\psi) - 0.2047\cos^2(\psi) \quad \text{(Eqn. S7)}^{13}$$

$$^3J_{C'-C'} = 0.46 - 0.95\cos(\psi) + 1.78\cos^2(\psi) \quad \text{(Eqn. S7)}^{18}$$

For both the $^1J_{C\alpha\text{-}H\alpha}$ and the $^2J_{C\alpha\text{-}N}$ couplings, the value of the constants $A_{RC}$ and $C$ are amino acid dependent. For Val, Thr, Ile, and Ser $C$ = 7.65 while for all other amino acids, $C$ = 8.15 [13]. The constant $A_{RC}$ values are listed in Table S1 below and originate from the random coil $^1J_{C\alpha\text{-}H\alpha}$ values [16].

**Table S1.**

| Residue | $A_{RC}$ | Residue | $A_{RC}$ | Residue | $A_{RC}$ | Residue | $A_{RC}$ |
|---|---|---|---|---|---|---|---|
| Ala | 143.7 | Glu | 141.9 | Met | 142.2 | Trp | 143.0 |
| Arg | 141.5 | His | 143.8 | Phe | 142.9 | Tyr | 143.0 |
| Asn | 141.5 | Ile | 141.3 | Pro | 148.4 | Val | 141.3 |
| Asp | 142.5 | Leu | 141.1 | Ser | 142.1 | Other | 140.3 |
| Gln | 141.1 | Lys | 141.5 | Thr | 141.4 | | |

**Percent Helicity:** For each structure, secondary structures were assigned using DSSP [14]. Residues assigned as a $3_{10}$ helix, α-helix or π-helix, corresponding to G, H and I assignments in DSSP respectively, were counted as helical.

### Comparison of αS Ensembles from Generalized Simulation to Experimental Data

### Summary of Generalized Simulations of αS

**Table S2:** Comparison of Simulated αS Ensembles to Global Experimental Data.

| Name | AtomType | ScoreFunction | Sampling PP | FI | SC | *Rg* (Å) | $E_{FRET}$ RMSD | $I_O/I_R$ RMSD | Dist. RMSD |
|---|---|---|---|---|---|---|---|---|---|
| | | ***Generalized Simulation*** | | | | | | | |
| VDW_PP | FA | fa_rep | X | | | 43.7 + 8.0 | 0.18 | 0.23 | 17.5 |
| VDW_PPFI | FA | fa_rep | X | X | | 39.7 + 6.8 | 0.15 | 0.21 | 13.17 |
| VDW_PPFISC | FA | fa_rep | X | X | X | 40.0 + 7.0 | 0.15 | 0.21 | 14.0 |
| CenStd_PP | CEN | cen_std | X | | | 19.6 + 2.5 | 0.28 | 0.20 | 11.9 |
| CenStd_PPFI | CEN | cen_std | X | X | | 19.2 + 2.5 | 0.28 | 0.22 | 12.5 |
| CenNath | CEN | CenNath | X | | | 33.4 + 7.9 | 0.11 | 0.18 | 8.09 |
| CenStd_Ext_PP | CEN | CenStdExt | X | | | 32.7 + 8.7 | 0.11 | 0.17 | 7.02 |
| CenStd_Ext_PPFI | CEN | CenStdExt | X | X | | 25.6 + 5.6 | 0.17 | 0.16 | 6.69 |
| Beta_PP | FA | ref2015 | X | | | 30.6 + 5.5 | 0.13 | 0.17 | 8.23 |
| Beta_PPFI | FA | ref2015 | X | X | | 36.4 + 6.9 | 0.12 | 0.19 | 11.7 |
| Beta_PPFISC | FA | ref2015 | X | X | X | 40.5 + 6.7 | 0.18 | 0.20 | 15.2 |
| Beta_PPSC | FA | ref2015 | X | | X | 30.3 + 5.3 | 0.13 | 0.17 | 7.74 |
| SimAnn_PP | CEN/FA | score0-3/ref2015 | X | | | 33.9 + 7.0 | 0.10 | 0.18 | 8.54 |
| SimAnn_PPFI | CEN/FA | score0-3/ref2015 | X | X | | 33.4 + 6.8 | 0.11 | 0.18 | 9.02 |
| SimAnn_PPFISC | CEN/FA | score0-3/ref2015 | X | X | X | 25.5 + 4.0 | 0.18 | 0.17 | 8.14 |
| SimAnn_PPSC | CEN/FA | score0-3/ref2015 | X | | X | 31.7 + 6.3 | 0.12 | 0.17 | 7.38 |
| | | ***FloppyTail*** | | | | | | | |
| FloppyTail_score12 | CEN/FA | CenStdExt/score12 | X | X | X | 32.8 + 6.0 | 0.12 | 0.18 | 8.36 |
| FloppyTail_ref2015 | CEN/FA | CenStdExt/ref2015 | X | X | X | 33.9 + 6.4 | 0.12 | 0.18 | 9.39 |
| FloppyTail_Quota | CEN/FA | CenStdExt /ref2015 | X | X | X | 34.2 + 6.1 | 0.12 | 0.18 | 9.02 |
| FloppyTail_Loops | CEN/FA | CenStdExt /ref2015 | X | X | X | 35.3 + 6.7 | 0.12 | 0.18 | 9.75 |
| FloppyTail_NoFrags | CEN/FA | CenStdExt /ref2015 | X | X | X | 34.8 + 6.3 | 0.12 | 0.18 | 9.00 |
| FloppyTail-Relax | CEN/FA | CenStdExt /ref2015 | X | X | X | 27.7 + 8.1 | 0.15 | 0.15 | 7.03 |
| FloppyTail | CEN/FA | CenStdExt /ref2015 | X | X | X | 34.0 + 7.7 | 0.11 | 0.18 | 7.94 |
| FastFloppyTail-Relax | CEN/FA | CenStdExt /ref2015 | X | X | X | 31.0 + 9.0 | 0.11 | 0.17 | 5.37 |
| FastFloppyTail | CEN/FA | CenStdExt /ref2015 | X | X | X | 38.0 + 9.4 | 0.12 | 0.20 | 10.1 |
| FloppyTail_Rot | CEN/FA | CenStdExt /ref2015 | X | X | X | 37.2 + 8.6 | 0.12 | 0.19 | 10.4 |
| FloppyTail_Rot-Relax | CEN/FA | CenStdExt /ref2015 | X | X | X | 29.2 + 8.7 | 0.13 | 0.15 | 6.25 |
| | | ***AbInitio*** | | | | | | | |
| AbInitio | CEN/FA | score0-3/ref2015 | X | X | X | 17.3 + 1.0 | 0.32 | 0.31 | 12.2 |
| AbInitioVO | CEN/FA | score0-3/ref2015 | X | X | X | 22.7 + 2.6 | 0.24 | 0.22 | 12.9 |
| | | ***Robustelli et. al.*** | | | | | | | |
| a99SB-*disp* | | Molecular Dynamics | | | | 36.73 | - | 0.17 | - |

*Atom type abbreviations: CEN = centroid, FA = full-atom.

** $R_g$ values can be compared to experimentally determined values of 33.0 ± 3.0 Å and 26.6 ± 0.5 Å from SAXS [19] and NMR [20] data, respectively. $E_{FRET}$ RMSD values were computed from data from Ferrie *et. al.*, Ferrie *et. al*. and Nath *et. al*. [1,21,22]. Distance RMSD values were computed from Grupi *et. al*. and Lee *et al*. [23,24] PRE RMSD values were computed from data from Sung *et. al.* and Dedmon *et. al.* [8,9].

**Table S3:** Comparison of Simulated αS Ensembles to Chemical Shift and RDC Data.

| **Simulation Name** | **N** | **H** | **C** | **Cα** | **Cβ** | **All CS** | **RDC M.Z.** |
|---|---|---|---|---|---|---|---|
| VDW_PP | 1.19 | 0.88 | 0.84 | 0.76 | 1.05 | 0.87 | 0.55 |
| VDW_PPFI | 1.52 | 0.17 | 0.55 | 0.65 | 1.06 | 0.91 | 0.55 |
| VDW_PPFISC | 1.83 | 0.19 | 0.49 | 0.50 | 1.08 | 1.01 | 0.52 |
| CenStd_PP | 3.42 | 0.25 | 0.70 | 0.65 | 1.28 | 1.69 | 0.55 |
| CenStd_PPFI | 3.66 | 0.20 | 0.66 | 1.07 | 1.31 | 1.83 | 0.44 |
| CenNath | 2.10 | 0.22 | 0.40 | 0.50 | 1.37 | 1.16 | 0.51 |
| CenStd_Ext_PP | 2.07 | 0.22 | 0.40 | 0.49 | 1.35 | 1.15 | 0.51 |
| CenStd_Ext_PPFI | 3.60 | 0.28 | 1.28 | 1.52 | 1.42 | 1.95 | 0.36 |
| Beta_PP | 0.95 | 1.06 | 0.57 | 2.67 | 0.14 | 1.38 | 0.54 |
| Beta_PPFI | 3.37 | 0.31 | 1.18 | 1.58 | 1.18 | 1.83 | 0.39 |
| Beta_PPFISC | 2.93 | 0.26 | 1.68 | 2.60 | 1.23 | 1.99 | 0.32 |
| Beta_PPSC | 1.59 | 0.13 | 1.00 | 0.41 | 1.11 | 0.99 | 0.57 |
| SimAnn_PP | 2.74 | 0.14 | 0.65 | 0.49 | 1.06 | 1.37 | 0.52 |
| SimAnn_PPFI | 3.63 | 0.31 | 0.83 | 1.10 | 1.14 | 1.81 | 0.43 |
| SimAnn_PPFISC | 3.10 | 0.27 | 1.20 | 1.81 | 1.08 | 1.77 | 0.37 |
| SimAnn_PPSC | 2.17 | 0.12 | 0.64 | 0.43 | 1.03 | 1.13 | 0.51 |
| FloppyTail_score12 | 2.03 | 0.18 | 0.39 | 0.58 | 1.02 | 1.07 | 0.48 |
| FloppyTail_ref2015 | 1.93 | 0.17 | 0.39 | 0.58 | 1.02 | 1.02 | 0.48 |
| FloppyTail_Quota | 1.75 | 0.18 | 0.38 | 0.46 | 1.03 | 0.95 | 0.49 |
| FloppyTail_Loops | 1.52 | 0.13 | 0.40 | 0.39 | 1.02 | 0.86 | 0.55 |
| FloppyTail_NoFrags | 1.52 | 0.13 | 0.41 | 0.38 | 1.01 | 0.86 | 0.55 |
| FloppyTail-Relax | 1.79 | 0.13 | 0.38 | 0.34 | 0.98 | 0.94 | 0.44 |
| FloppyTail | 1.69 | 0.13 | 0.36 | 0.35 | 0.98 | 0.90 | 0.44 |
| FastFloppyTail-Relax | 1.62 | 0.11 | 0.47 | 0.36 | 1.02 | 0.90 | 0.42 |
| FastFloppyTail | 1.55 | 0.12 | 0.42 | 0.33 | 1.01 | 0.86 | 0.41 |
| FloppyTail_Rot | 1.66 | 0.13 | 0.36 | 0.33 | 0.98 | 0.89 | 0.44 |
| FloppyTail_Rot-Relax | 1.77 | 0.12 | 0.38 | 0.34 | 0.97 | 0.93 | 0.44 |
| AbInitio | 2.68 | 0.25 | 1.37 | 2.08 | 1.23 | 1.73 | 0.38 |
| AbInitioVO | 2.24 | 0.24 | 0.61 | 0.72 | 1.03 | 1.19 | 0.47 |
| a99SB-*disp* | 1.46 | 0.14 | 0.31 | 0.51 | 1.04 | 0.85 | 0.41 |

*All chemical shift data are reported as RMSD values from computed from data from Sung *et al.* [8]. RDC values are computed as Q-values, as described in Zweckstetter *et. al.* [12], based on data from Bertoncini *et al*. (RDC M.Z.) [25].

**Table S4:** Comparison of Simulated αS Ensembles to *J*-Coupling Data.

| Simulation Name | $^3J_{HNH\alpha}$ | $^1J_{C\alpha H\alpha}$ | $^1J_{NC\alpha}$ | $^2J_{NC\alpha}$ | $^3J_{C'C'}$ | All *J* |
|---|---|---|---|---|---|---|
| ***Generalized Simulation*** | | | | | | |
| VDW_PP | 2.38 | 1.31 | 0.63 | 0.42 | 0.69 | 1.30 |
| VDW_PPFI | 2.63 | 1.10 | 0.72 | 0.74 | 0.57 | 1.38 |
| VDW_PPFISC | 2.39 | 1.51 | 0.94 | 0.93 | 0.52 | 1.42 |
| CenStd_PP | 1.41 | 1.37 | 0.66 | 0.46 | 0.61 | 0.99 |
| CenStd_PPFI | 1.21 | 1.68 | 0.98 | 1.25 | 0.14 | 1.17 |
| CenNath | 1.67 | 1.46 | 0.25 | 0.45 | 0.41 | 1.03 |
| CenStd_Ext_PP | 1.61 | 1.43 | 0.25 | 0.45 | 0.39 | 1.00 |
| CenStd_Ext_PPFI | 1.85 | 3.15 | 0.89 | 1.55 | 0.42 | 1.83 |
| Beta_PP | 0.89 | 1.39 | 0.43 | 0.40 | 0.73 | 0.85 |
| Beta_PPFI | 1.22 | 2.81 | 0.91 | 1.29 | 0.38 | 1.55 |
| Beta_PPFISC | 2.41 | 4.31 | 1.15 | 1.48 | 0.40 | 2.37 |
| Beta_PPSC | 0.82 | 1.28 | 0.38 | 0.40 | 0.74 | 0.79 |
| SimAnn_PP | 0.67 | 1.68 | 0.55 | 0.45 | 0.58 | 0.91 |
| SimAnn_PPFI | 0.84 | 2.08 | 0.91 | 1.37 | 0.24 | 1.25 |
| SimAnn_PPFISC | 1.50 | 3.21 | 0.99 | 1.39 | 0.34 | 1.77 |
| SimAnn_PPSC | 0.43 | 1.39 | 0.60 | 0.49 | 0.50 | 0.76 |
| ***FloppyTail*** | | | | | | |
| FloppyTail_score12 | 0.48 | 1.64 | 0.54 | 0.70 | 0.16 | 0.88 |
| FloppyTail_ref2015 | 0.52 | 1.68 | 0.52 | 0.79 | 0.17 | 0.88 |
| FloppyTail_Quota | 0.62 | 1.55 | 0.49 | 0.74 | 0.19 | 0.85 |
| FloppyTail_Loops | 0.51 | 1.10 | 0.35 | 0.43 | 0.10 | 0.60 |
| FloppyTail_NoFrags | 0.53 | 1.11 | 0.35 | 0.43 | 0.10 | 0.60 |
| FloppyTail-Relax | 0.59 | 1.25 | 0.41 | 0.54 | 0.16 | 0.69 |
| FloppyTail | 0.64 | 1.45 | 0.40 | 0.53 | 0.16 | 0.77 |
| FastFloppyTail-Relax | 0.65 | 1.14 | 0.36 | 0.39 | 0.26 | 0.64 |
| FastFloppyTail | 0.56 | 1.22 | 0.35 | 0.39 | 0.24 | 0.65 |
| FloppyTail_Rot | 0.61 | 1.41 | 0.40 | 0.51 | 0.16 | 0.75 |
| FloppyTail_Rot-Relax | 0.60 | 1.23 | 0.40 | 0.52 | 0.17 | 0.68 |
| ***AbInitio*** | | | | | | |
| AbInitio | 2.15 | 3.49 | 0.99 | 1.32 | 0.37 | 1.98 |
| AbInitioVO | 1.16 | 1.82 | 0.71 | 1.12 | 0.25 | 1.13 |
| ***Robustelli et. al*** | | | | | | |
| a99SB-*disp* | 1.11 | - | - | - | 0.18 | - |

*All data are reported as RMSD values computed from data from Mantsyzov *et al.* and Lee *et. al.* [13,18].

**Comparison of αS Per Residue Percent Helicity**

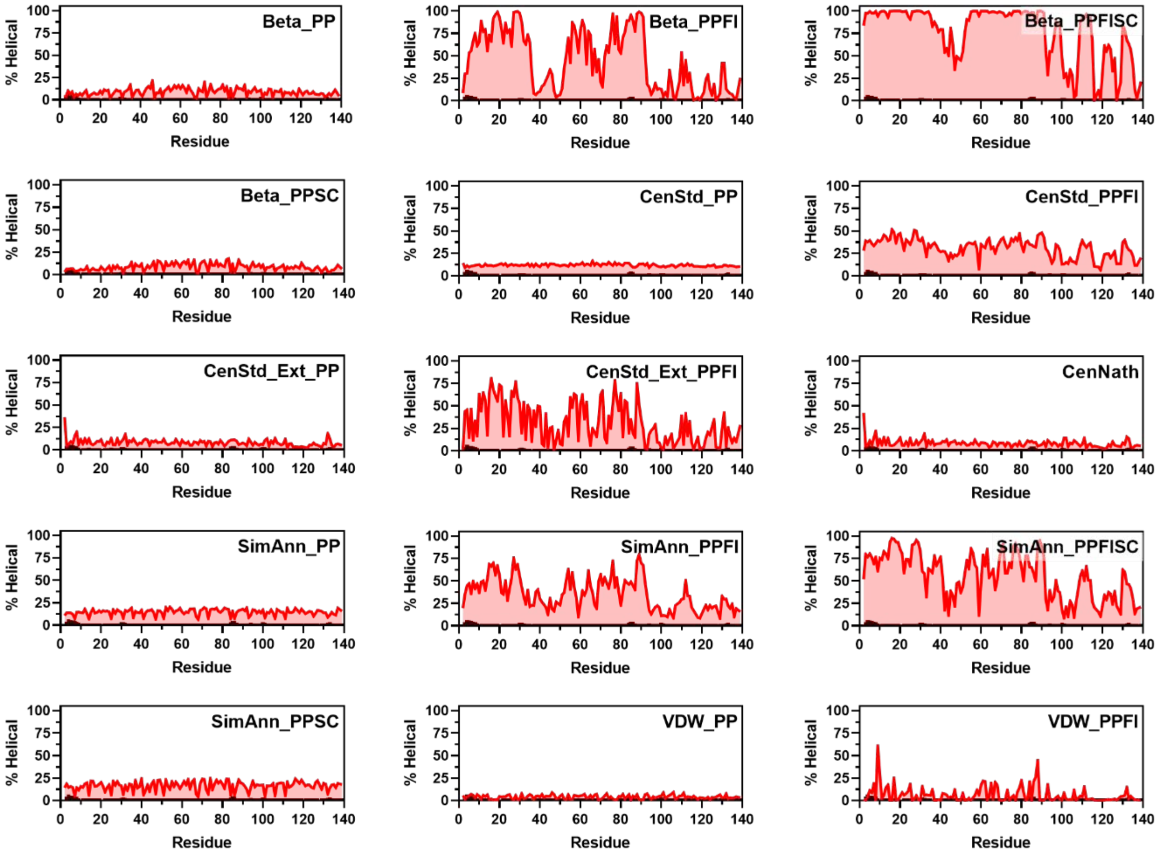


**Figure S1:** Plots of Percent Helicity. Plots of the percentage of structures containing helices on a per residue basis from Beta_PP (Row 1 Left), Beta_PPFI (Row 1 Middle), Beta_PPFISC (Row 1 Right), Beta_PPSC (Row 2 Left), CenStd_PP (Row 2 Middle), CenStd_PPFI (Row 2 Right), CenStd_Ext_PP (Row 3 Left), CenStd_Ext_PPFI (Row 3 Middle), CenNath (Row 3 Right), SimAnn_PP (Row 4 Left), SimAnn_PPFI (Row 4 Middle), SimAnn_PPFISC (Row 4 Right), SimAnn_PPSC (Row 5 Left), VDW_PP (Row 5 Middle) and VDW_PPFI (Row 5 Right) ensembles compared to percentages computed from Sung *et al*. chemical shift data using the D2D method from Camilloni *et al*. [8,26].

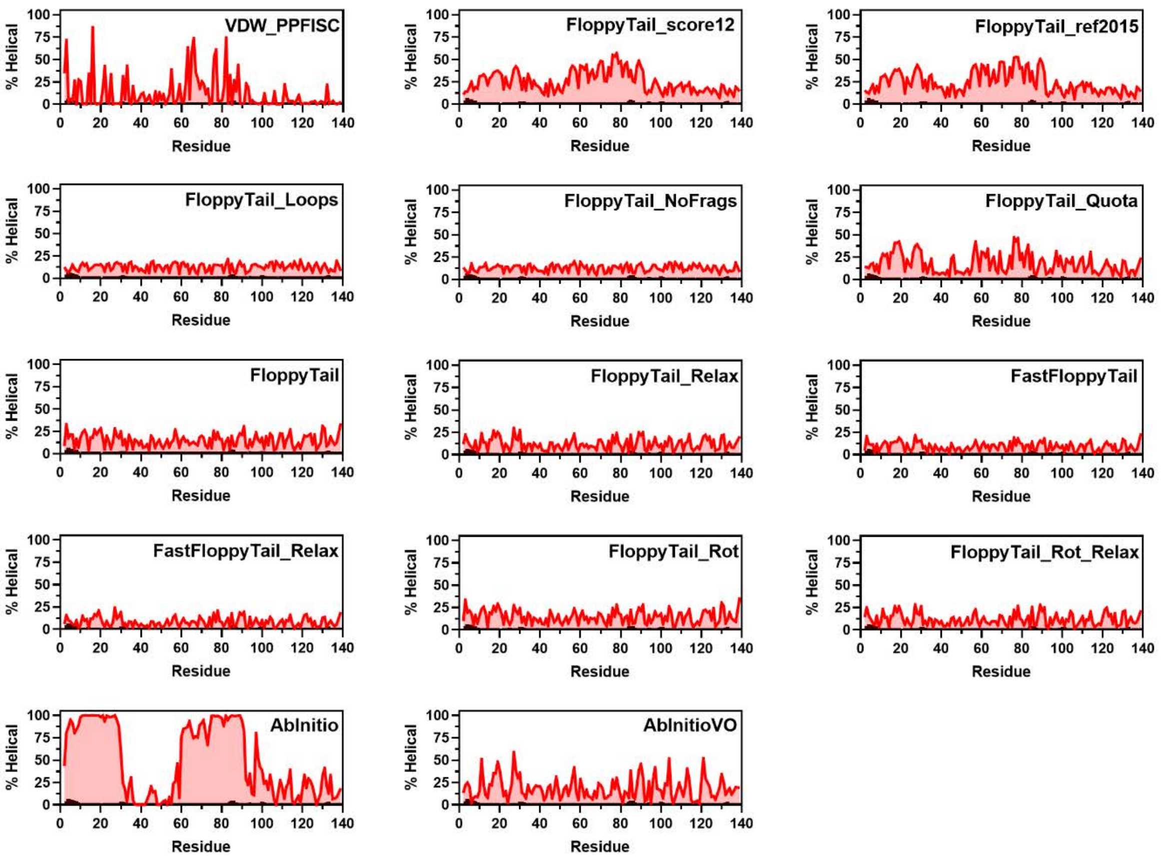


**Figure S2:** Plots of Percent Helicity. Plots of the percentage of structures containing helices on a per residue basis from VDW_PPFISC (Row 1 Left), FloppyTail_score12 (Row 1 Middle), FloppyTail_ref2015 (Row 1 Right), FloppyTail_Quota (Row 2 Left), FloppyTail_NoFrags (Row 2 Middle), FloppyTail_Loops (Row 2 Right), FloppyTail (Row 3 Left), FloppyTail_Relax (Row 3 Middle), FastFloppyTail (Row 3 Right), FastFloppyTail_Relax (Row 4 Left), FloppyTail_Rot (Row 4 Middle), FloppyTail_Rot_Relax (Row 4 Right), AbInitio (Row 5 Left) and AbInitioVO (Row 5 Middle) ensembles compared to percentages computed from Sung *et al*. chemical shift data using the D2D method from Camilloni *et al*. [8,26].

### Comparison with αS Radius of Gyration Data

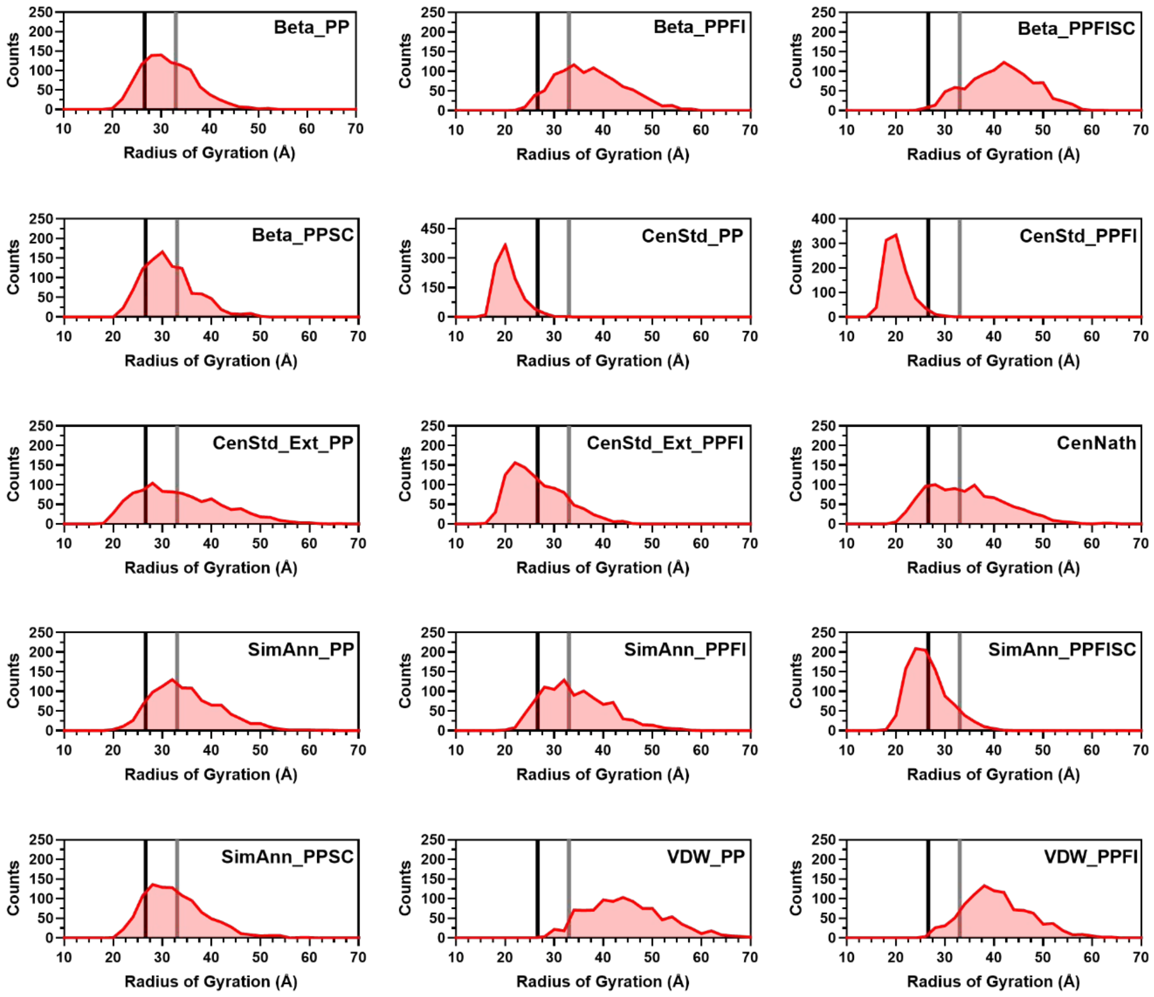


**Figure S3:** Histograms of Radii of Gyration. Histograms of the radius of gyration from Beta_PP (Row 1 Left), Beta_PPFI (Row 1 Middle), Beta_PPFISC (Row 1 Right), Beta_PPSC (Row 2 Left), CenStd_PP (Row 2 Middle), CenStd_PPFI (Row 2 Right), CenStd_Ext_PP (Row 3 Left), CenStd_Ext_PPFI (Row 3 Middle), CenNath (Row 3 Right), SimAnn_PP (Row 4 Left), SimAnn_PPFI (Row 4 Middle), SimAnn_PPFISC (Row 4 Right), SimAnn_PPSC (Row 5 Left), VDW_PP (Row 5 Middle) and VDW_PPFI (Row 5 Right) ensembles compared to experimental values from SAXS [19] (grey) and NMR [20] (black).

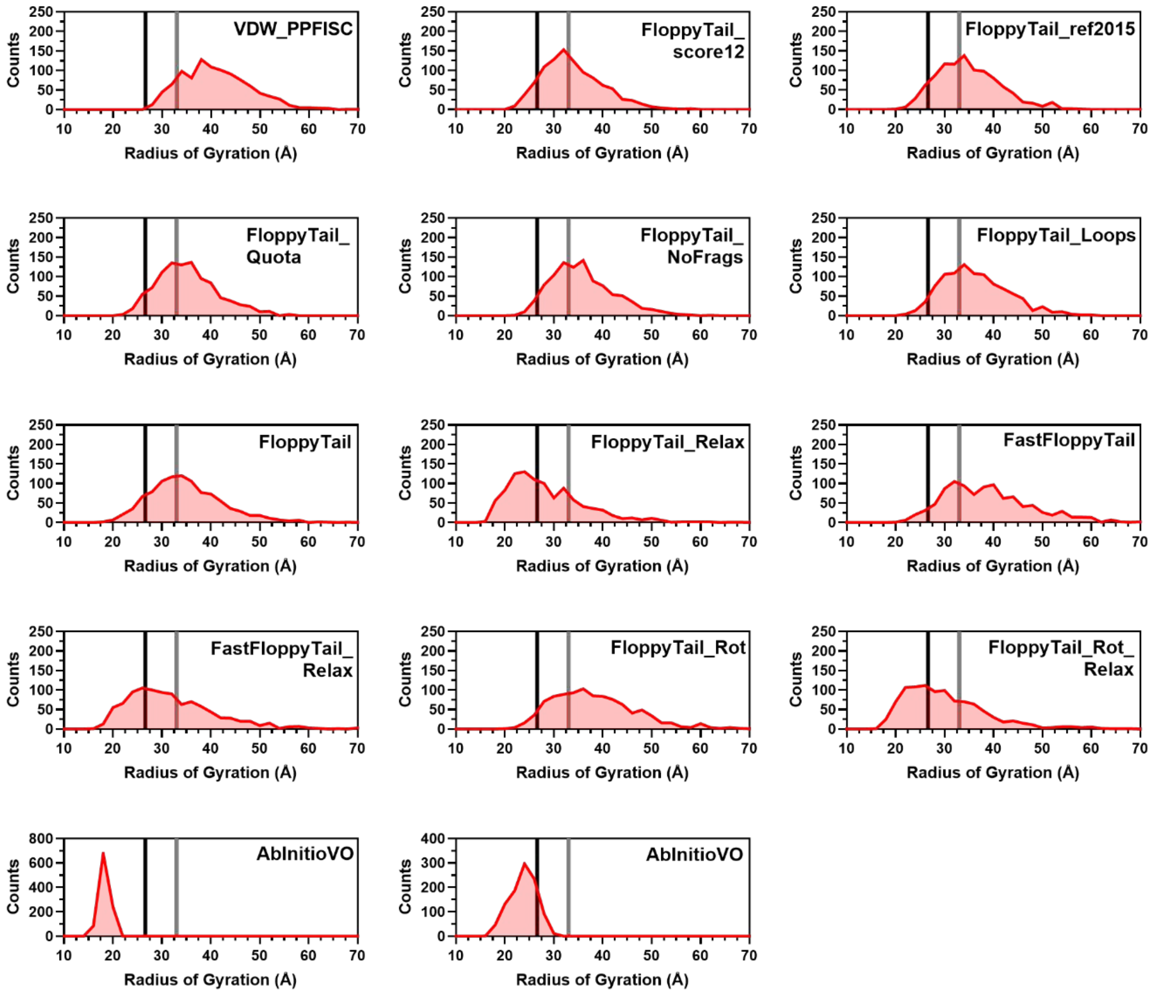


**Figure S4:** Histograms of Radii of Gyration. Histograms of the radius of gyration from VDW_PPFISC (Row 1 Left), FloppyTail_score12 (Row 1 Middle), FloppyTail_ref2015 (Row 1 Right), FloppyTail_Quota (Row 2 Left), FloppyTail_NoFrags (Row 2 Middle), FloppyTail_Loops (Row 2 Right), FloppyTail (Row 3 Left), FloppyTail_Relax (Row 3 Middle), FastFloppyTail (Row 3 Right), FastFloppyTail_Relax (Row 4 Left), FloppyTail_Rot (Row 4 Middle), FloppyTail_Rot_Relax (Row 4 Right), AbInitio (Row 5 Left) and AbInitioVO (Row 5 Middle) ensembles compared to experimental values from SAXS [19] (grey) and NMR [20] (black).

### Comparison with αS FRET Data

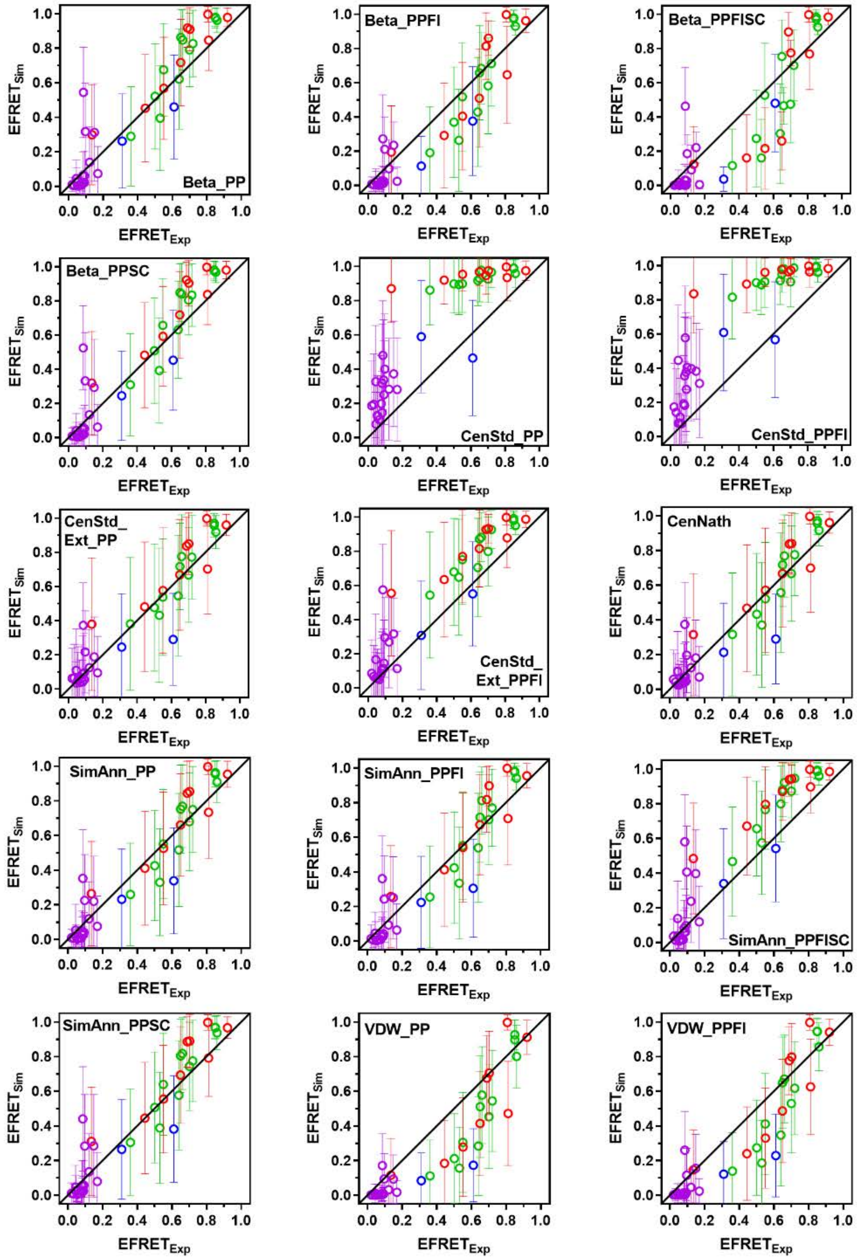


**Figure S5:** Comparison of Simulated $E_{FRET}$ with Experimental $E_{FRET}$. Simulated $E_{FRET}$s for Beta_PP (Row 1 Left), Beta_PPFI (Row 1 Center), Beta_PPFISC (Row 1 Right), Beta_PPSC (Row 2 Left), CenStd_PP (Row 2 Center), CenStd_PPFI (Row 2 Right), CenStd_Ext_PP (Row 3 Left), CenStd_Ext_PPFI (Row 3 Center), CenNath (Row 3 Right), SimAnn_PP (Row 4 Left), SimAnn_PPFI (Row 4 Center), SimAnn_PPFISC (Row 4 Right), SimAnn_PPSC (Row 5 Left) , VDW_PP (Row 5 Middle) , VDW_PPFI (Row 5 Right) with data from Ferrie *et al*. Cnf-Trp [21] (Purple) and Fam-Raz [21] (Red) Pairs, Ferrie *et al*. Mcm-Acd pair [22] (Blue), and Nath *et al*. [1] (Green).

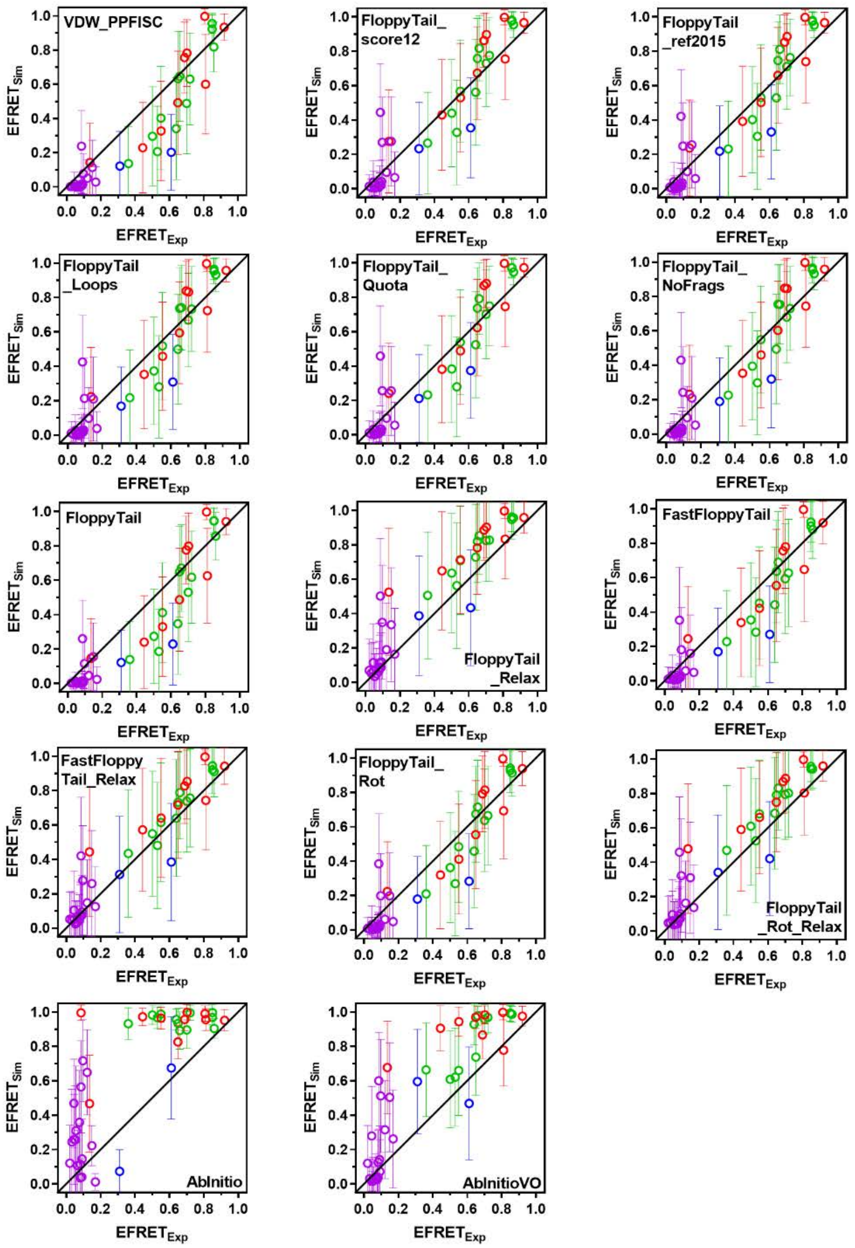


**Figure S6:** Comparison of Simulated $E_{FRET}$ with Experimental $E_{FRET}$. Simulated $E_{FRET}$s for VDW_PPFISC (Row 1 Left), FloppyTail_score12 (Row 1 Center), FloppyTail_ref2015 (Row 1 Right), FloppyTail_Loops (Row 2 Left), FloppyTail_Quota (Row 2 Center), FloppyTail_NoFrags (Row 2 Right), FloppyTail (Row 3 Left), FloppyTail_Relax (Row 3 Center), FastFloppyTail (Row 3 Right), FastFloppyTail_Relax (Row 4 Left), FloppyTail_Rot (Row 4 Center), FloppyTail_Rot_Relax (Row 4 Right), AbInitio (Row 5 Left) and AbInitioVO (Row 5 Middle) with data from Ferrie *et al*. Cnf-Trp [21] (Purple) and Fam-Raz [21] (Red) Pairs, Ferrie *et al*. Mcm-Acd pair[22] (Blue), and Nath *et al*. [1] (Green).

### Comparison with αS Distance Data

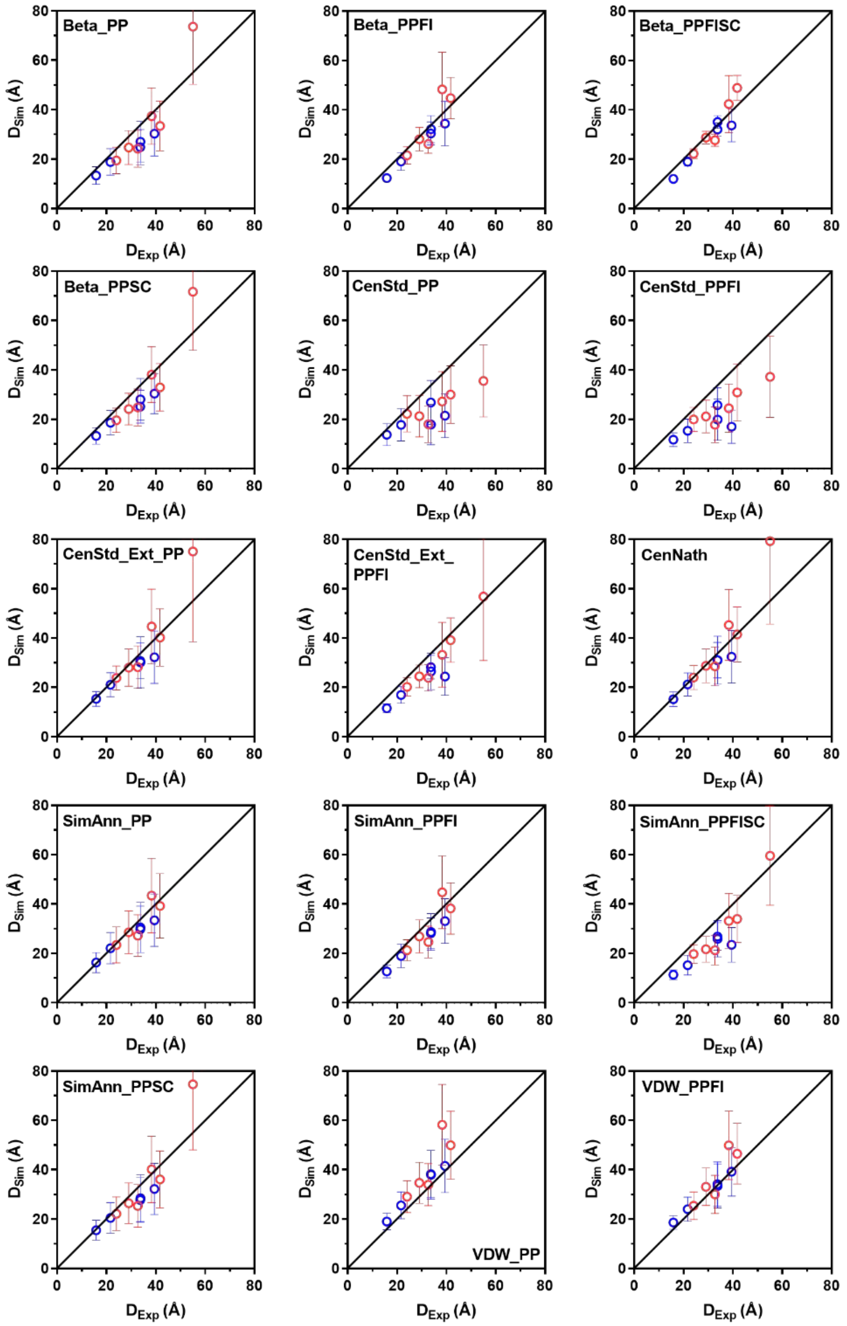


**Figure S7:** Comparison of Simulated Distance with Experimental Distances. Simulated distances for Beta_PP (Row 1 Left), Beta_PPFI (Row 1 Center), Beta_PPFISC (Row 1 Right), Beta_PPSC (Row 2 Left), CenStd_PP (Row 2 Center), CenStd_PPFI (Row 2 Right), CenStd_Ext_PP (Row 3 Left), CenStd_Ext_PPFI (Row 3 Center), CenNath (Row 3 Right), SimAnn_PP (Row 4 Left), SimAnn_PPFI (Row 4 Center), SimAnn_PPFISC (Row 4 Right), SimAnn_PPSC (Row 5 Left) , VDW_PP (Row 5 Middle) , VDW_PPFI (Row 5 Right) with data from Lee *et al.* [24] (Red) and Grupi *et al.* [23] (Blue).

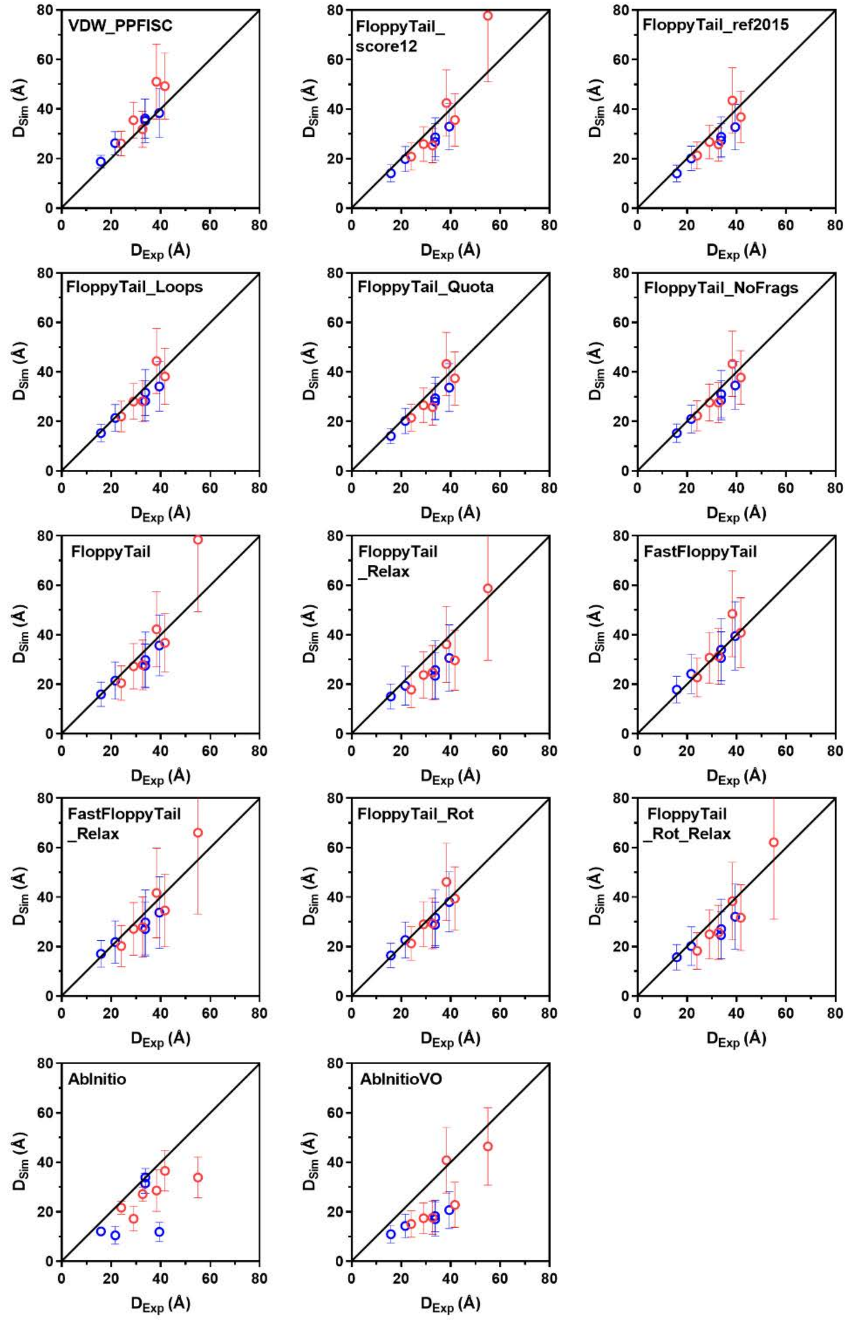


**Figure S8:** Comparison of Simulated Distance with Experimental Distances. Simulated distances for VDW_PPFISC (Row 1 Left), FloppyTail_score12 (Row 1 Center), FloppyTail_ref2015 (Row 1 Right), FloppyTail_Loops (Row 2 Left), FloppyTail_Quota (Row 2 Center), FloppyTail_NoFrags (Row 2 Right), FloppyTail (Row 3 Left), FloppyTail_Relax (Row 3 Center), FastFloppyTail (Row 3 Right), FastFloppyTail_Relax (Row 4 Left), FloppyTail_Rot (Row 4 Center), FloppyTail_Rot_Relax (Row 4 Right), AbInitio (Row 5 Left) and AbInitioVO (Row 5 Middle) with data from Lee *et al*. [24] (Red) and Grupi *et al*. [23] (Blue).

**Comparison with αS PRE Data**

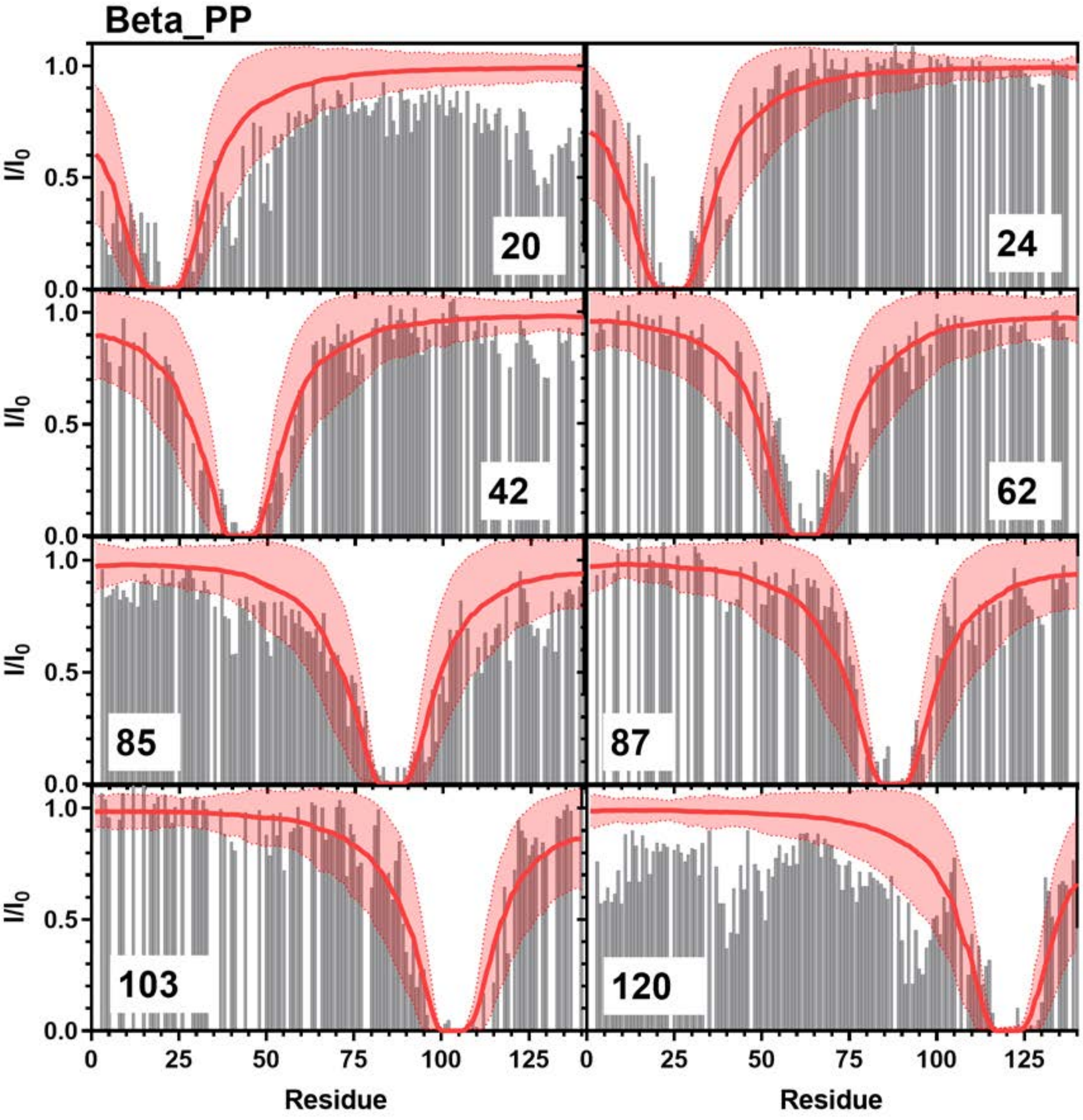


**Figure S9:** Comparison of Simulated PRE Values and Experimental PRE Values: Simulated PRE values from Beta_PP (red line) overlayed on top of experimental data (grey bars) from positions 20 (Top Left), 24 (Top Right), 42 (Upper Middle Left), 62 (Upper Middle Right), 85 (Lower Middle Left), 87 (Lower Middle Right), 103 (Bottom Left), 120 (Bottom Right). Experimental data for positions 20, 85, and 120 are from Sung *et al.* [8] and data for positions 24, 42, 62, 87, and 103 are from Dedmon *et al.* [9].

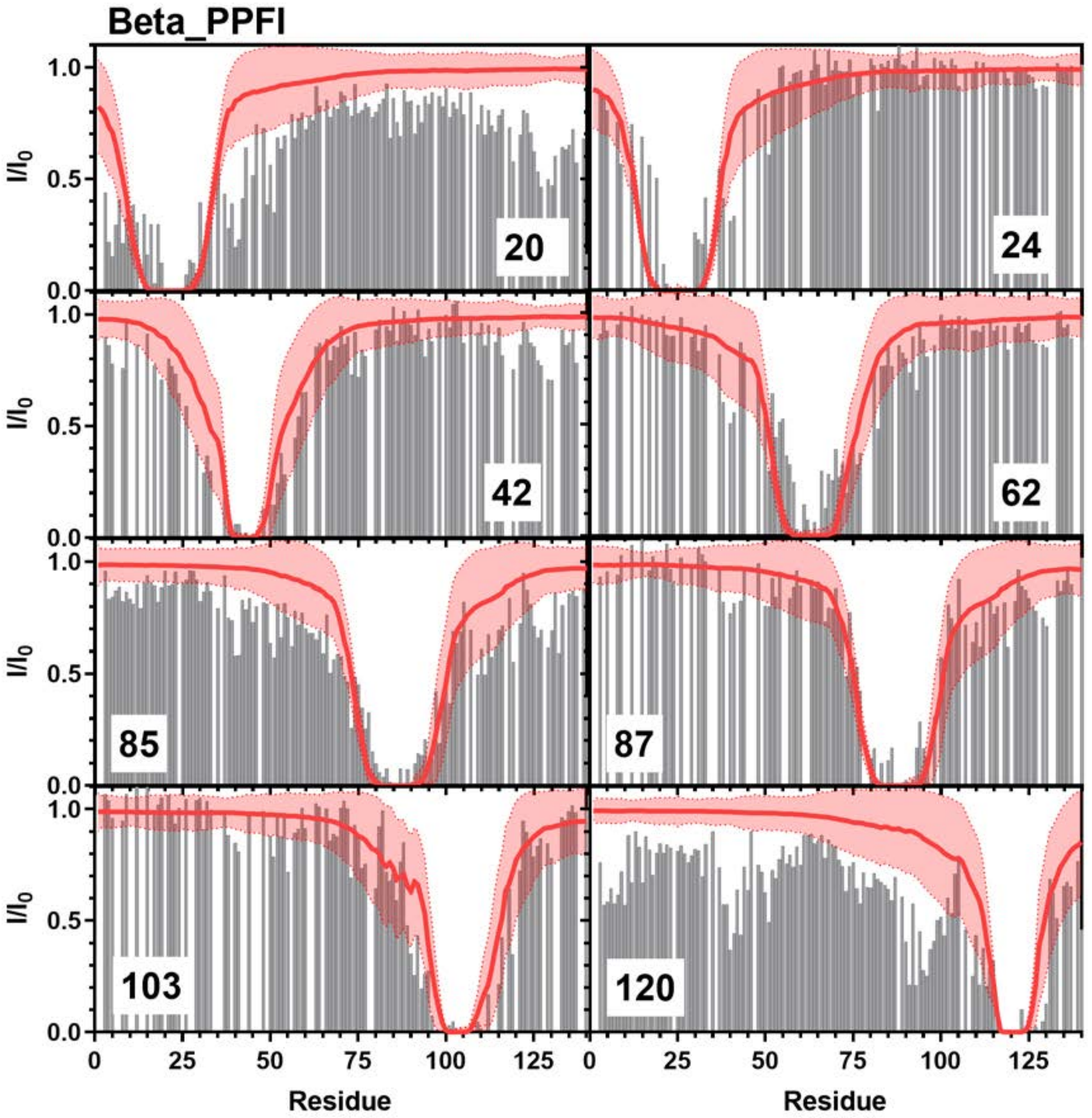


**Figure S10:** Comparison of Simulated PRE Values and Experimental PRE Values: Simulated PRE values from Beta_PPFI (red line) overlayed on top of experimental data (grey bars) from positions 20 (Top Left), 24 (Top Right), 42 (Upper Middle Left), 62 (Upper Middle Right), 85 (Lower Middle Left), 87 (Lower Middle Right), 103 (Bottom Left), 120 (Bottom Right). Experimental data for positions 20, 85, and 120 are from Sung *et al.* [8] and data for positions 24, 42, 62, 87, and 103 are from Dedmon *et al.* [9].

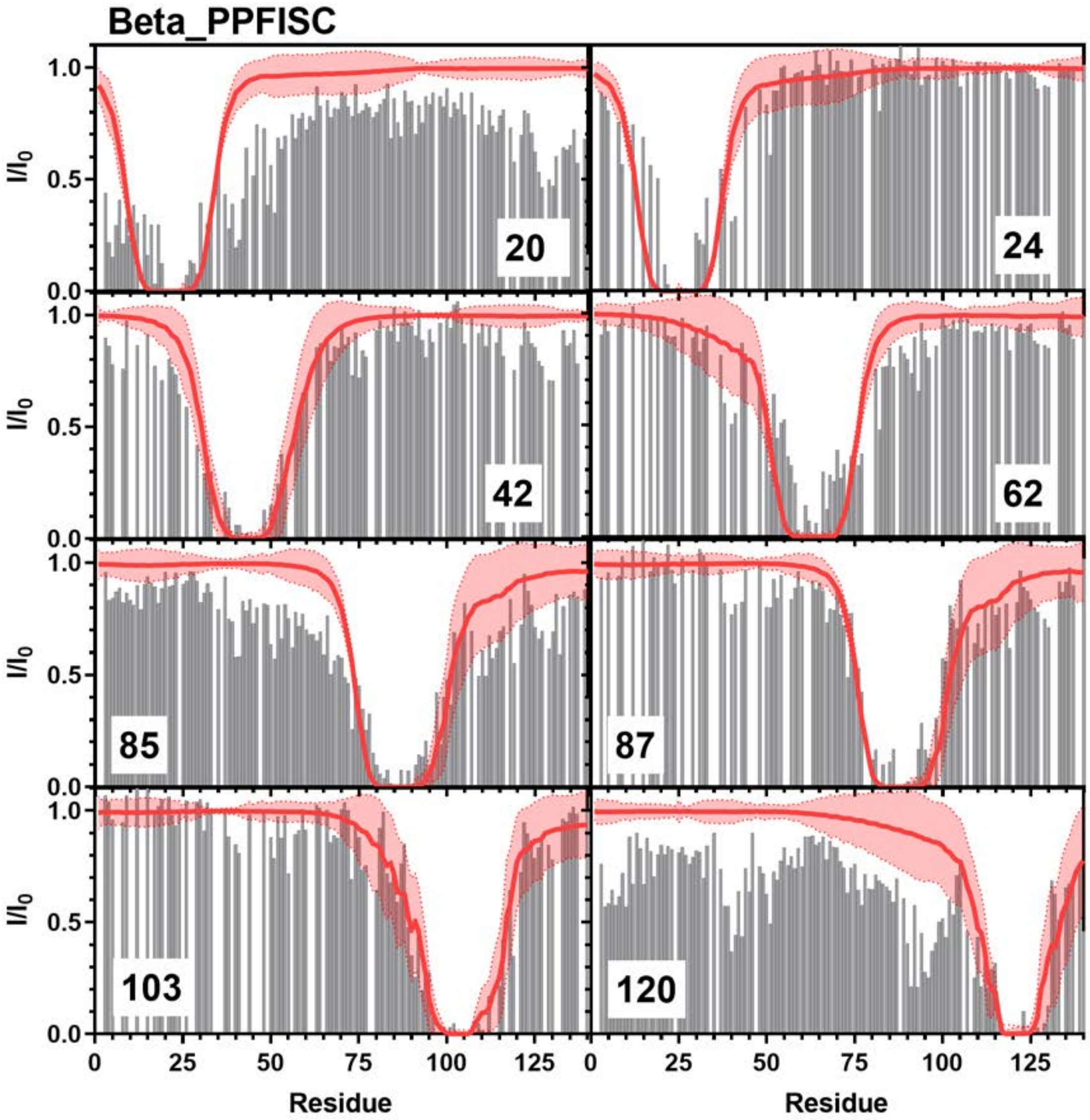


**Figure S11:** Comparison of Simulated PRE Values and Experimental PRE Values: Simulated PRE values from Beta_PPFISC (red line) overlayed on top of experimental data (grey bars) from positions 20 (Top Left), 24 (Top Right), 42 (Upper Middle Left), 62 (Upper Middle Right), 85 (Lower Middle Left), 87 (Lower Middle Right), 103 (Bottom Left), 120 (Bottom Right). Experimental data for positions 20, 85, and 120 are from Sung *et al.* [8] and data for positions 24, 42, 62, 87, and 103 are from Dedmon *et al.* [9].

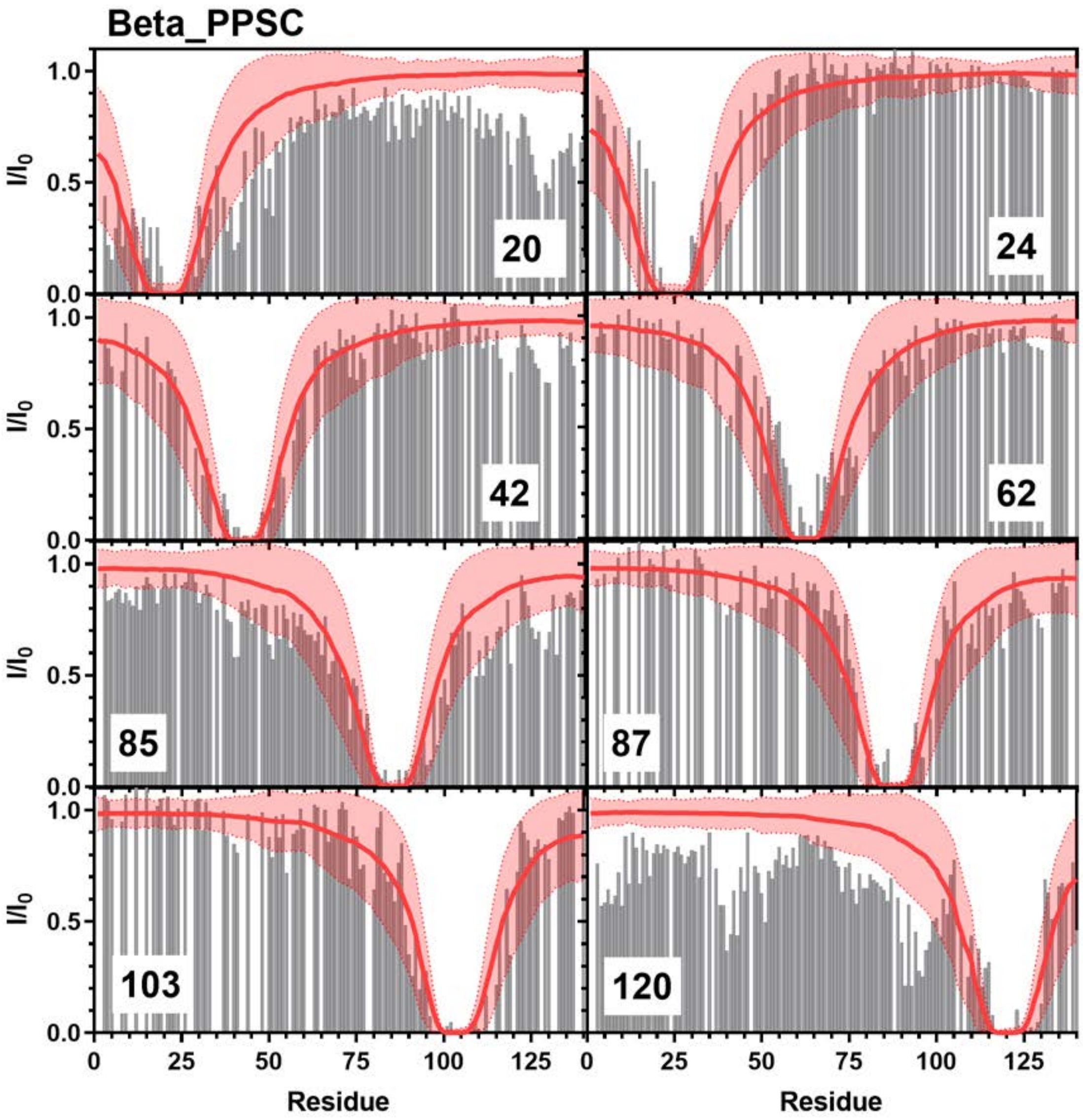


**Figure S12:** Comparison of Simulated PRE Values and Experimental PRE Values: Simulated PRE values from Beta_PPSC (red line) overlayed on top of experimental data (grey bars) from positions 20 (Top Left), 24 (Top Right), 42 (Upper Middle Left), 62 (Upper Middle Right), 85 (Lower Middle Left), 87 (Lower Middle Right), 103 (Bottom Left), 120 (Bottom Right). Experimental data for positions 20, 85, and 120 are from Sung *et al.* [8] and data for positions 24, 42, 62, 87, and 103 are from Dedmon *et al.* [9].

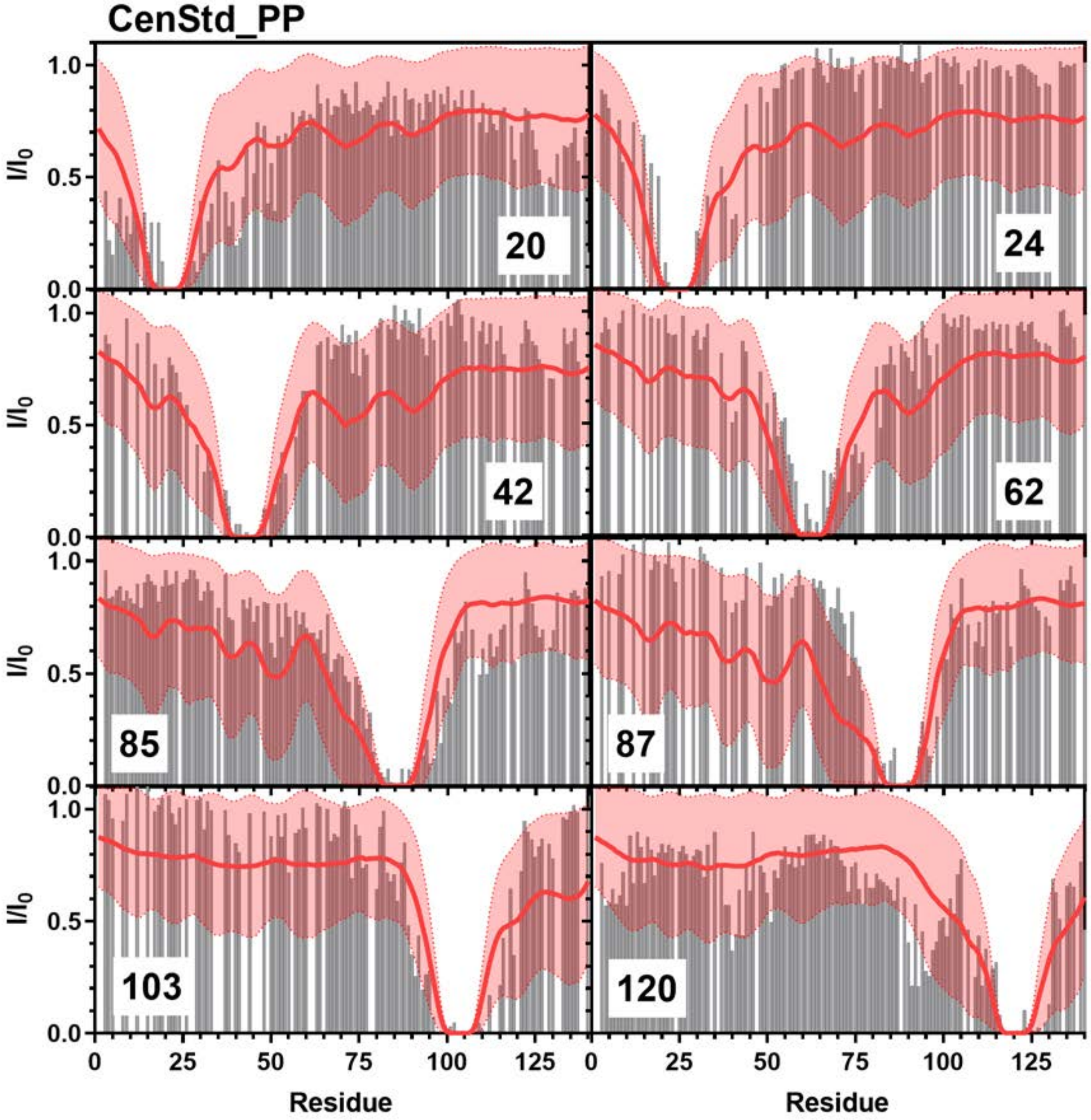


**Figure S13:** Comparison of Simulated PRE Values and Experimental PRE Values: Simulated PRE values from CenStd_PP (red line) overlayed on top of experimental data (grey bars) from positions 20 (Top Left), 24 (Top Right), 42 (Upper Middle Left), 62 (Upper Middle Right), 85 (Lower Middle Left), 87 (Lower Middle Right), 103 (Bottom Left), 120 (Bottom Right). Experimental data for positions 20, 85, and 120 are from Sung *et al.* [8] and data for positions 24, 42, 62, 87, and 103 are from Dedmon *et al.* [9].

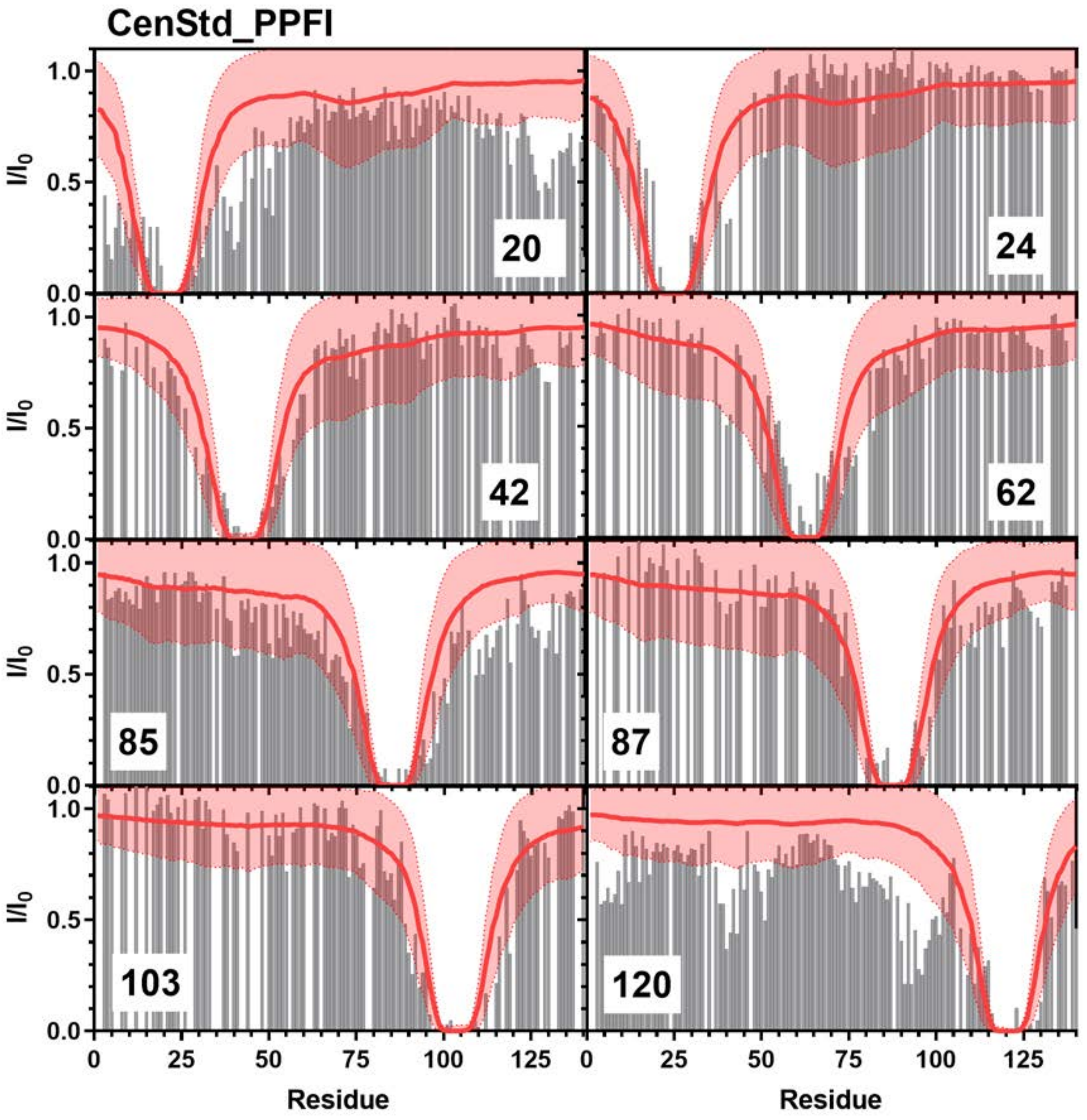


**Figure S14:** Comparison of Simulated PRE Values and Experimental PRE Values: Simulated PRE values from CenStd_PPFI (red line) overlayed on top of experimental data (grey bars) from positions 20 (Top Left), 24 (Top Right), 42 (Upper Middle Left), 62 (Upper Middle Right), 85 (Lower Middle Left), 87 (Lower Middle Right), 103 (Bottom Left), 120 (Bottom Right). Experimental data for positions 20, 85, and 120 are from Sung *et al*. [8] and data for positions 24, 42, 62, 87, and 103 are from Dedmon *et al*. [9].

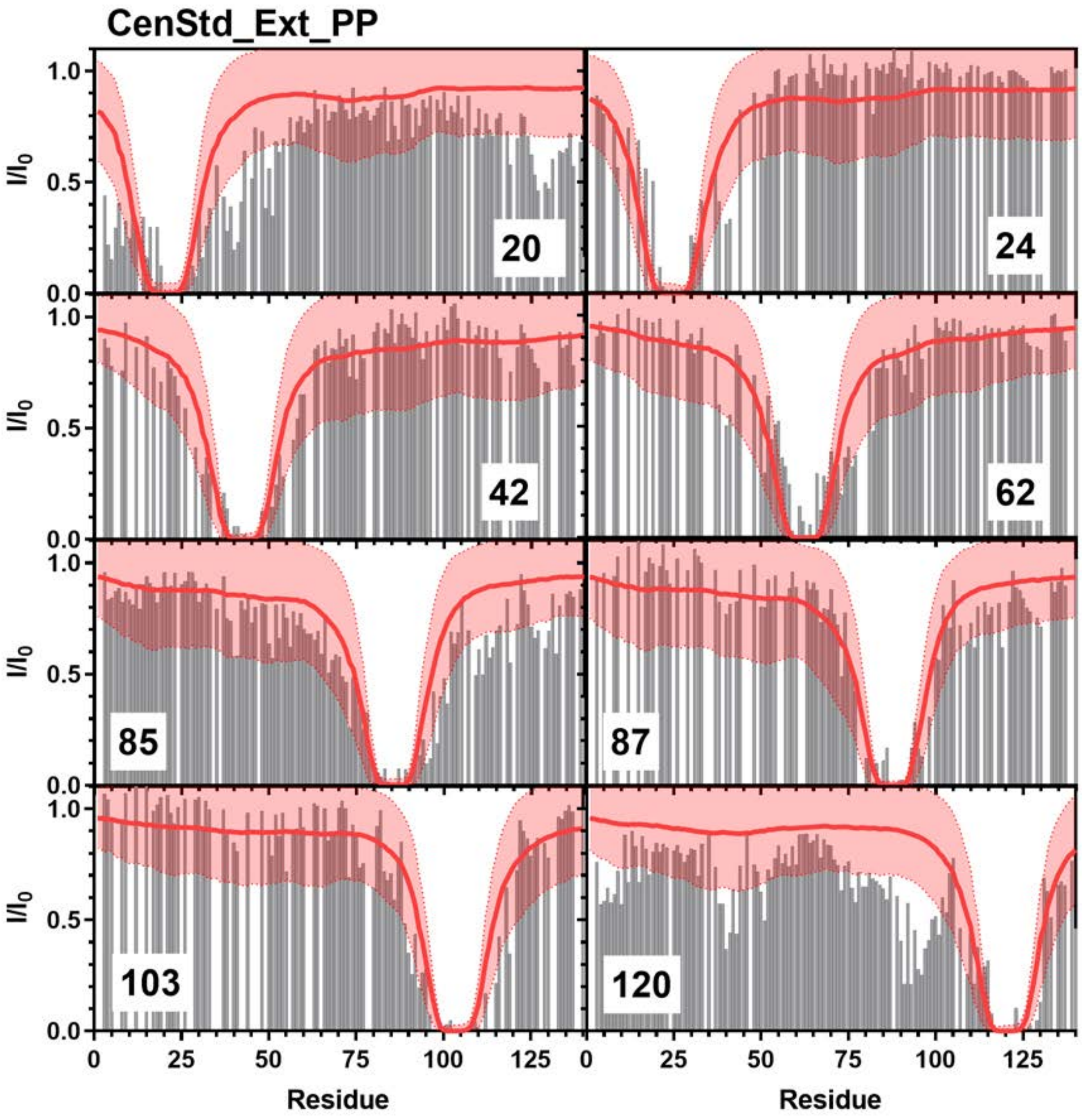


**Figure S15:** Comparison of Simulated PRE Values and Experimental PRE Values: Simulated PRE values from CenStd_Ext_PP (red line) overlayed on top of experimental data (grey bars) from positions 20 (Top Left), 24 (Top Right), 42 (Upper Middle Left), 62 (Upper Middle Right), 85 (Lower Middle Left), 87 (Lower Middle Right), 103 (Bottom Left), 120 (Bottom Right). Experimental data for positions 20, 85, and 120 are from Sung *et al.* [8] and data for positions 24, 42, 62, 87, and 103 are from Dedmon *et al.* [9].

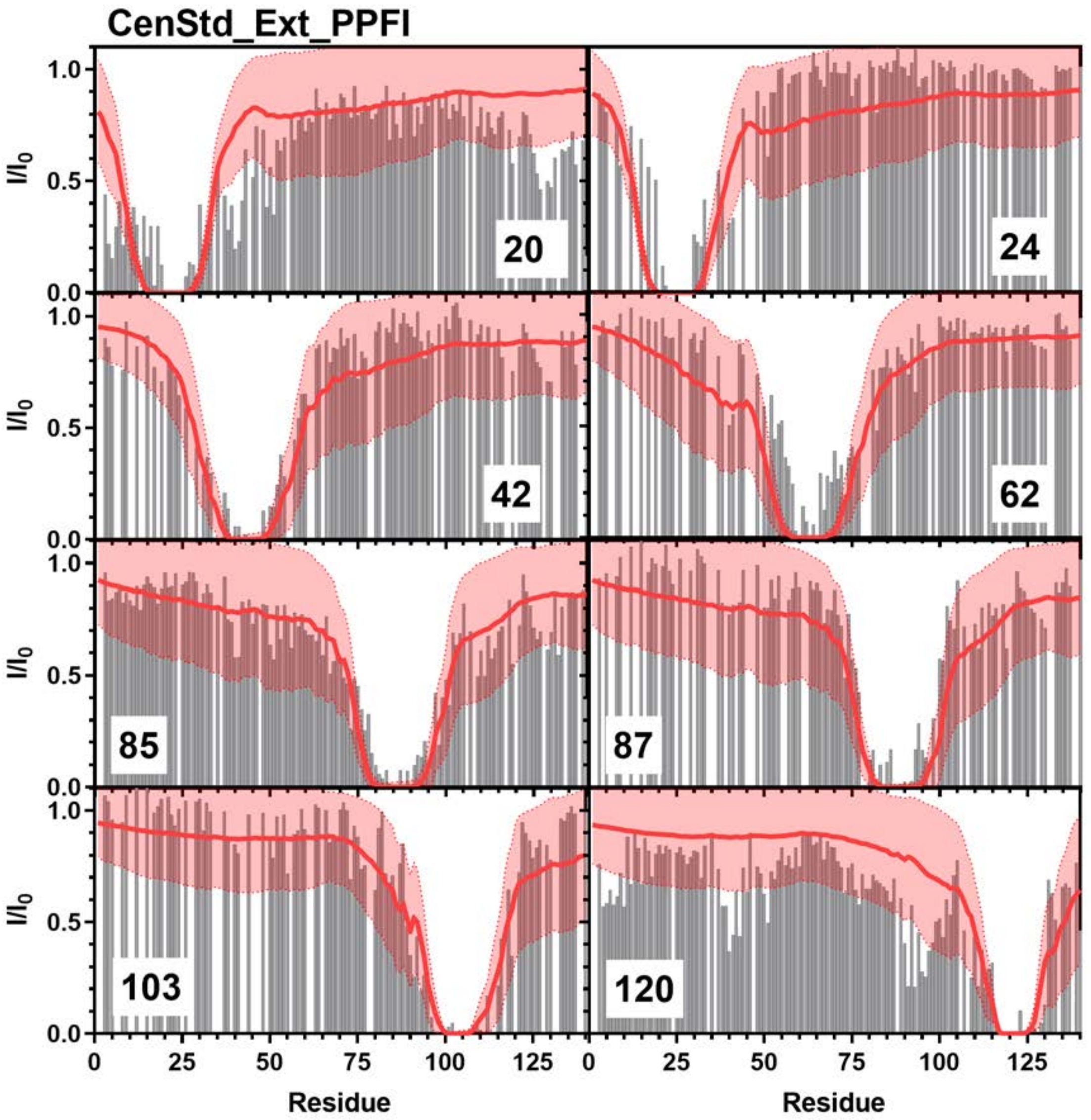


**Figure S16:** Comparison of Simulated PRE Values and Experimental PRE Values: Simulated PRE values from CenStd_Ext_PPFI (red line) overlayed on top of experimental data (grey bars) from positions 20 (Top Left), 24 (Top Right), 42 (Upper Middle Left), 62 (Upper Middle Right), 85 (Lower Middle Left), 87 (Lower Middle Right), 103 (Bottom Left), 120 (Bottom Right). Experimental data for positions 20, 85, and 120 are from Sung *et al*. [8] and data for positions 24, 42, 62, 87, and 103 are from Dedmon *et al*. [9].

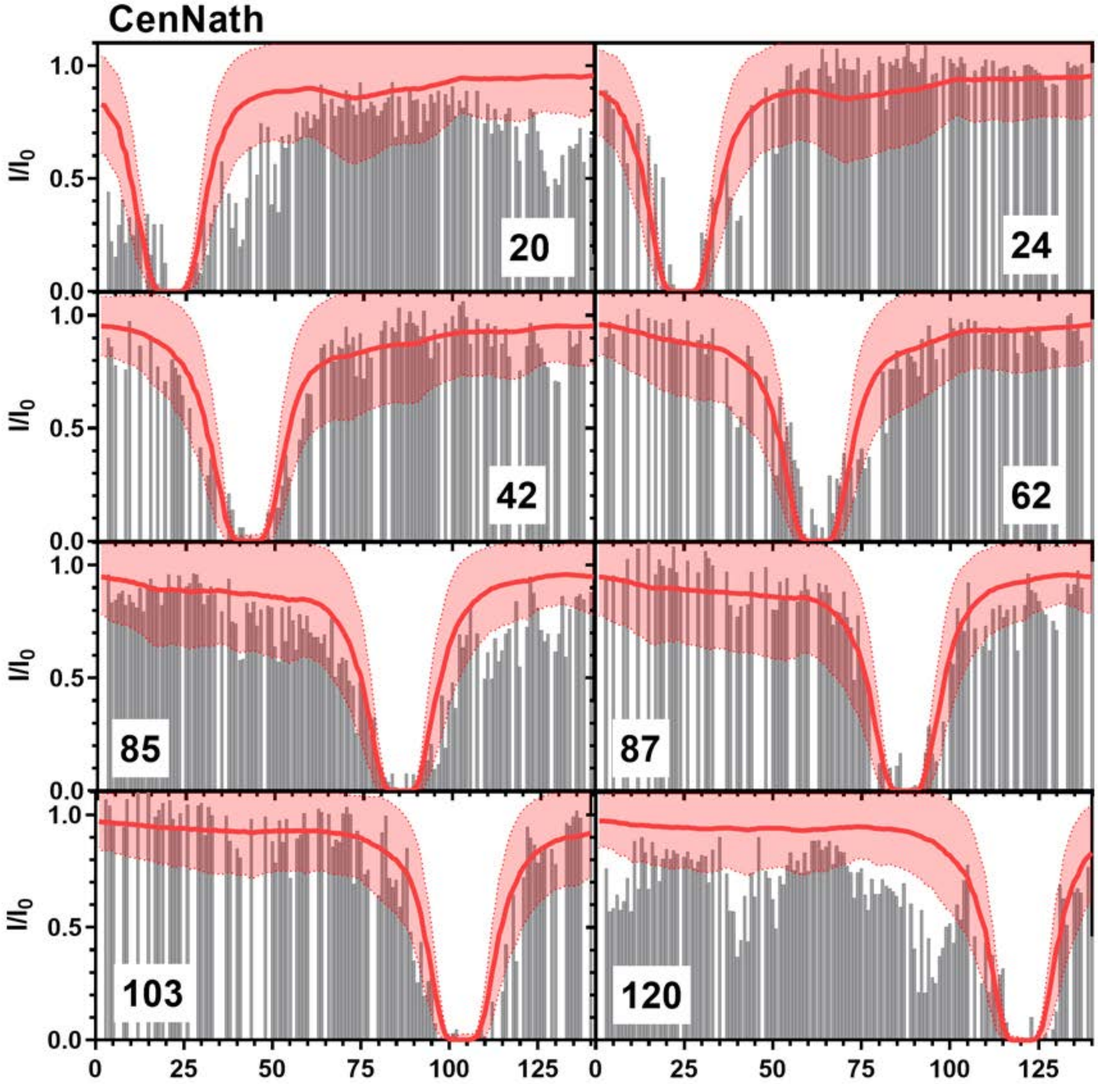


**Figure S17:** Comparison of Simulated PRE Values and Experimental PRE Values: Simulated PRE values from CenNath (red line) overlayed on top of experimental data (grey bars) from positions 20 (Top Left), 24 (Top Right), 42 (Upper Middle Left), 62 (Upper Middle Right), 85 (Lower Middle Left), 87 (Lower Middle Right), 103 (Bottom Left), 120 (Bottom Right). Experimental data for positions 20, 85, and 120 are from Sung *et al*. [8] and data for positions 24, 42, 62, 87, and 103 are from Dedmon *et al*. [9].

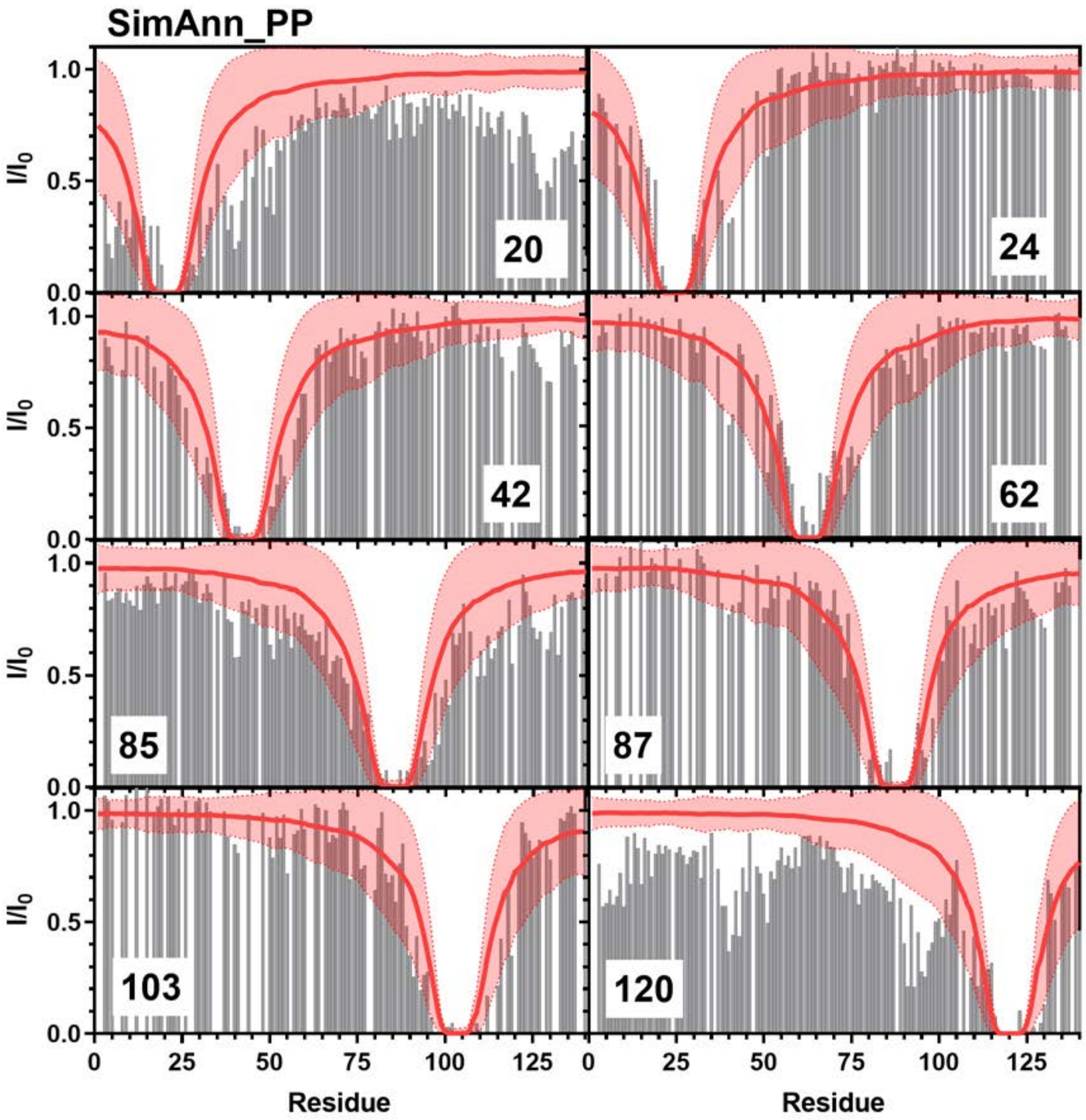


**Figure S18:** Comparison of Simulated PRE Values and Experimental PRE Values: Simulated PRE values from SimAnn_PP (red line) overlayed on top of experimental data (grey bars) from positions 20 (Top Left), 24 (Top Right), 42 (Upper Middle Left), 62 (Upper Middle Right), 85 (Lower Middle Left), 87 (Lower Middle Right), 103 (Bottom Left), 120 (Bottom Right). Experimental data for positions 20, 85, and 120 are from Sung *et al*. [8] and data for positions 24, 42, 62, 87, and 103 are from Dedmon *et al*. [9].

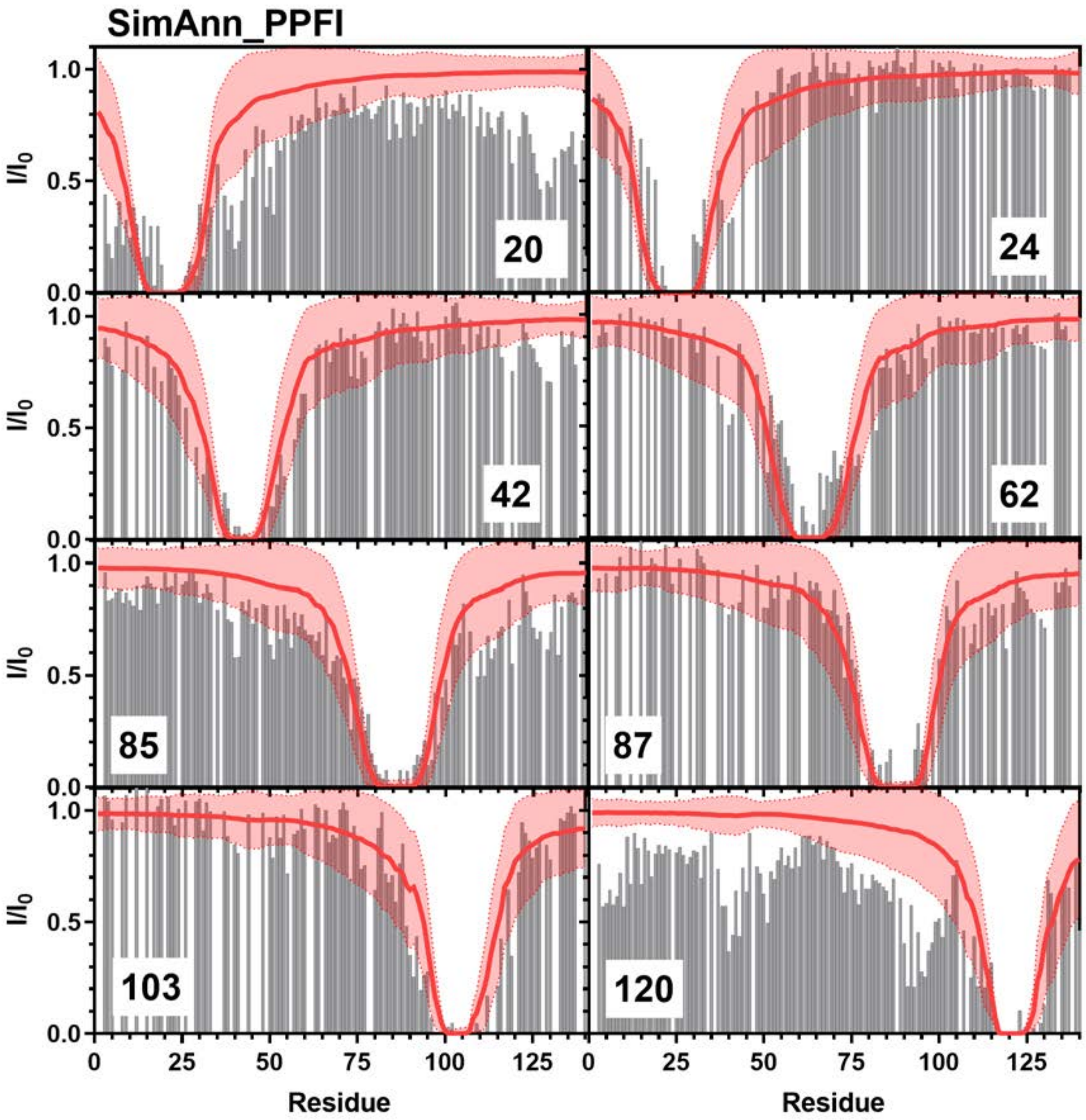


**Figure S19:** Comparison of Simulated PRE Values and Experimental PRE Values: Simulated PRE values from SimAnn_PPFI (red line) overlayed on top of experimental data (grey bars) from positions 20 (Top Left), 24 (Top Right), 42 (Upper Middle Left), 62 (Upper Middle Right), 85 (Lower Middle Left), 87 (Lower Middle Right), 103 (Bottom Left), 120 (Bottom Right). Experimental data for positions 20, 85, and 120 are from Sung *et al*. [8] and data for positions 24, 42, 62, 87, and 103 are from Dedmon *et al*. [9].

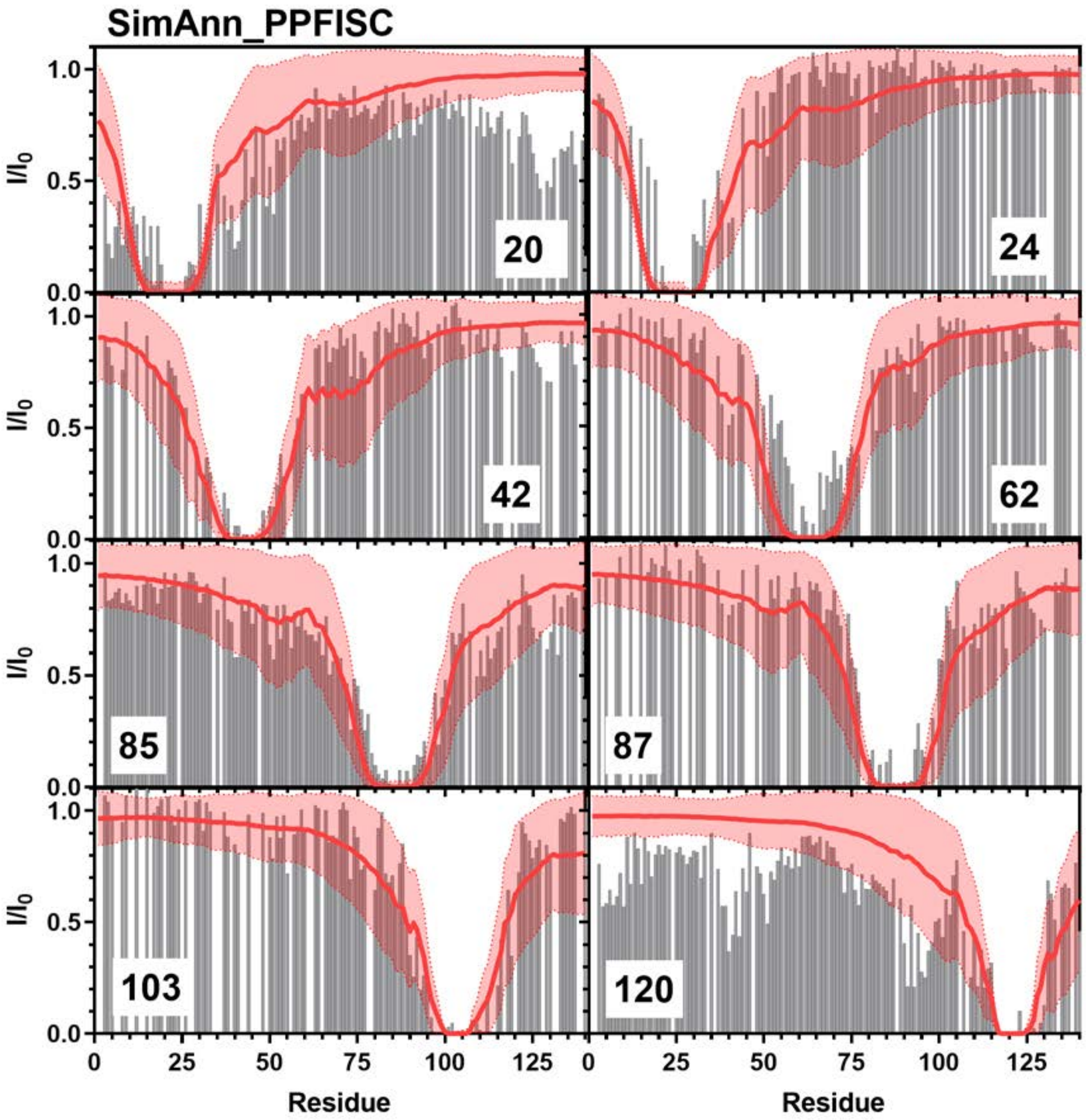


**Figure S20:** Comparison of Simulated PRE Values and Experimental PRE Values: Simulated PRE values from SimAnn_PPFISC (red line) overlayed on top of experimental data (grey bars) from positions 20 (Top Left), 24 (Top Right), 42 (Upper Middle Left), 62 (Upper Middle Right), 85 (Lower Middle Left), 87 (Lower Middle Right), 103 (Bottom Left), 120 (Bottom Right). Experimental data for positions 20, 85, and 120 are from Sung *et al*. [8] and data for positions 24, 42, 62, 87, and 103 are from Dedmon *et al*. [9].

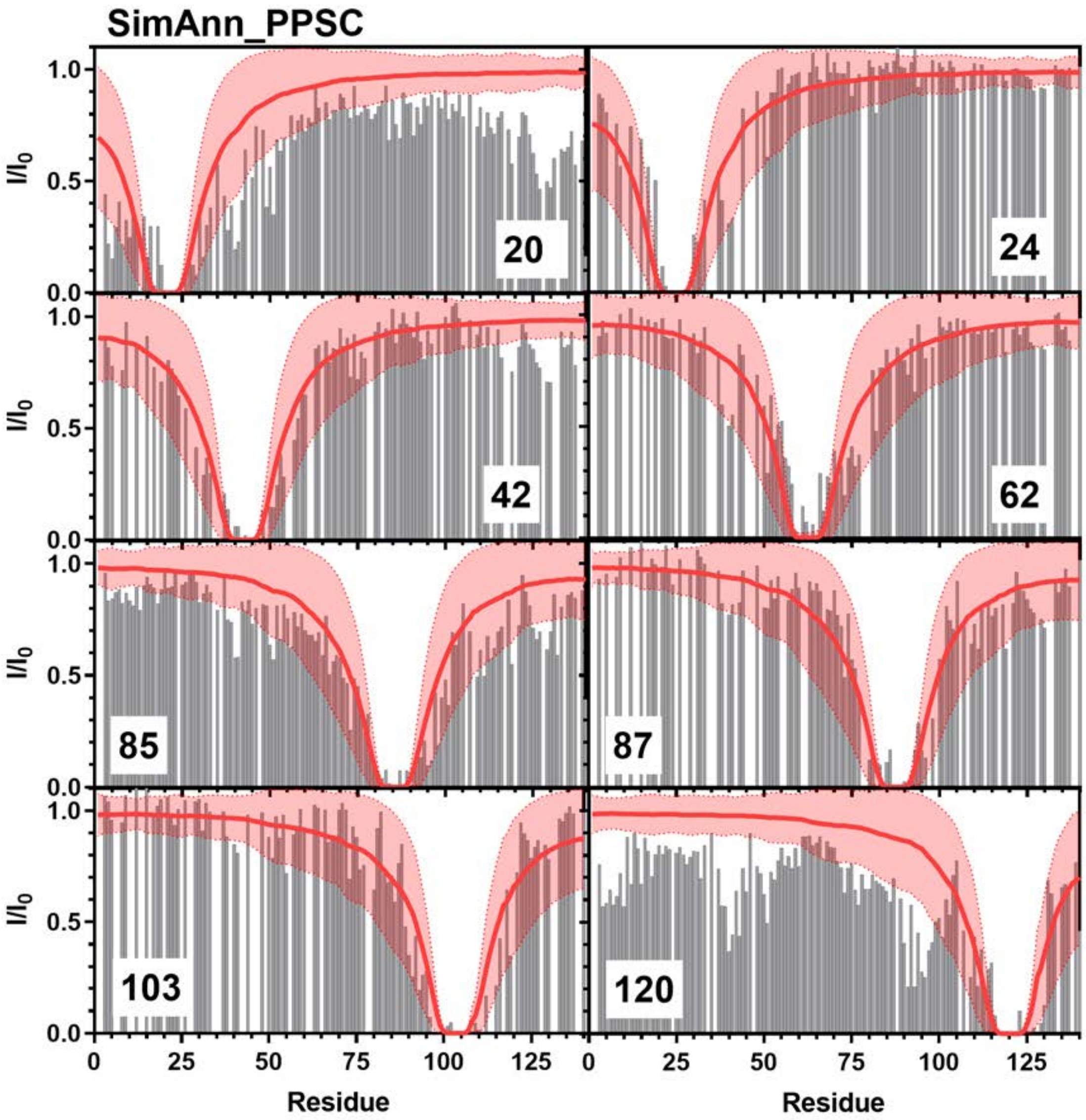


**Figure S21:** Comparison of Simulated PRE Values and Experimental PRE Values: Simulated PRE values from SimAnn_PPSC (red line) overlayed on top of experimental data (grey bars) from positions 20 (Top Left), 24 (Top Right), 42 (Upper Middle Left), 62 (Upper Middle Right), 85 (Lower Middle Left), 87 (Lower Middle Right), 103 (Bottom Left), 120 (Bottom Right). Experimental data for positions 20, 85, and 120 are from Sung *et al.* [8] and data for positions 24, 42, 62, 87, and 103 are from Dedmon *et al.* [9].

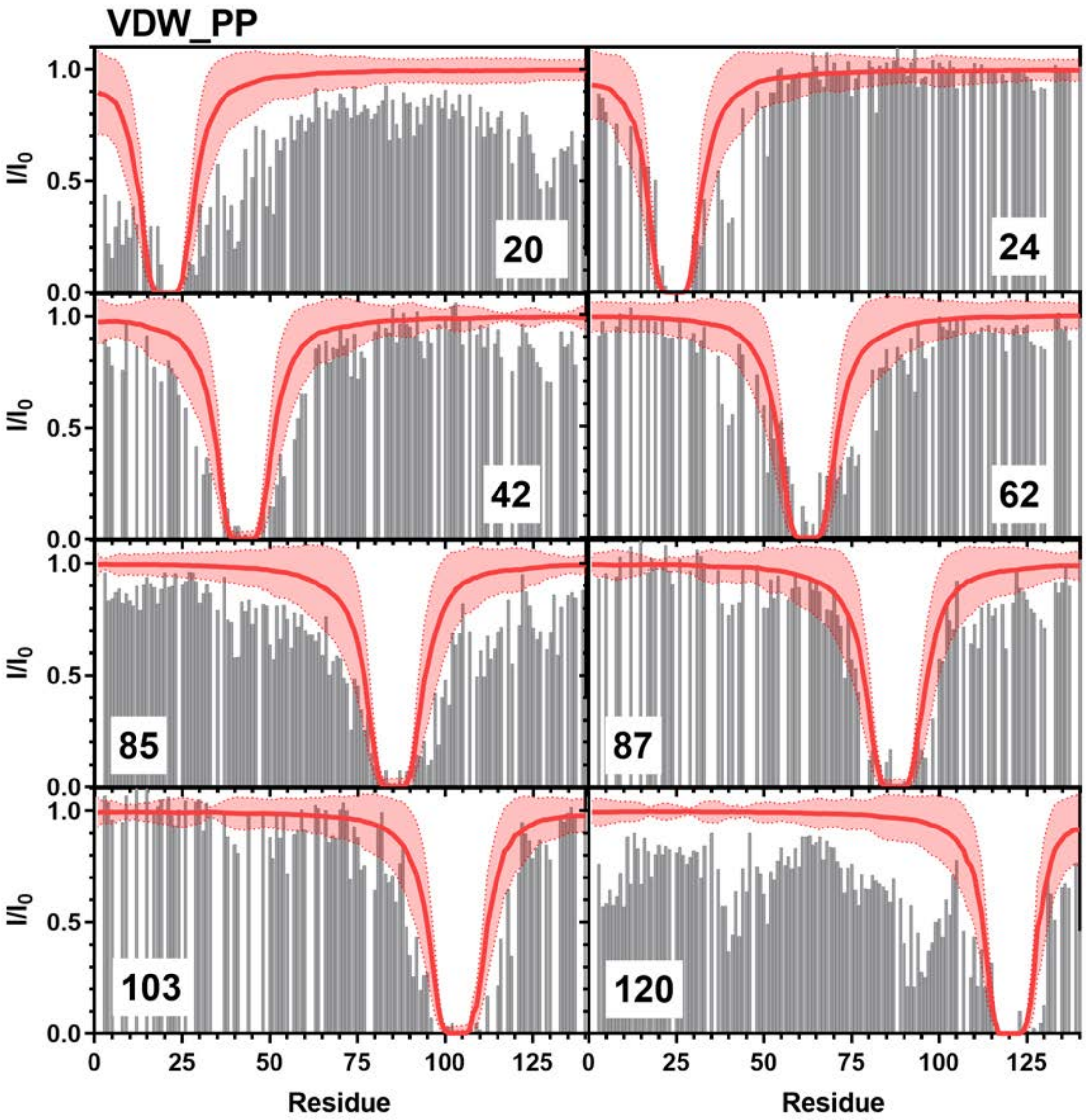


**Figure S22:** Comparison of Simulated PRE Values and Experimental PRE Values: Simulated PRE values from VDW_PP (red line) overlayed on top of experimental data (grey bars) from positions 20 (Top Left), 24 (Top Right), 42 (Upper Middle Left), 62 (Upper Middle Right), 85 (Lower Middle Left), 87 (Lower Middle Right), 103 (Bottom Left), 120 (Bottom Right). Experimental data for positions 20, 85, and 120 are from Sung *et al.* [8] and data for positions 24, 42, 62, 87, and 103 are from Dedmon *et al.* [9].

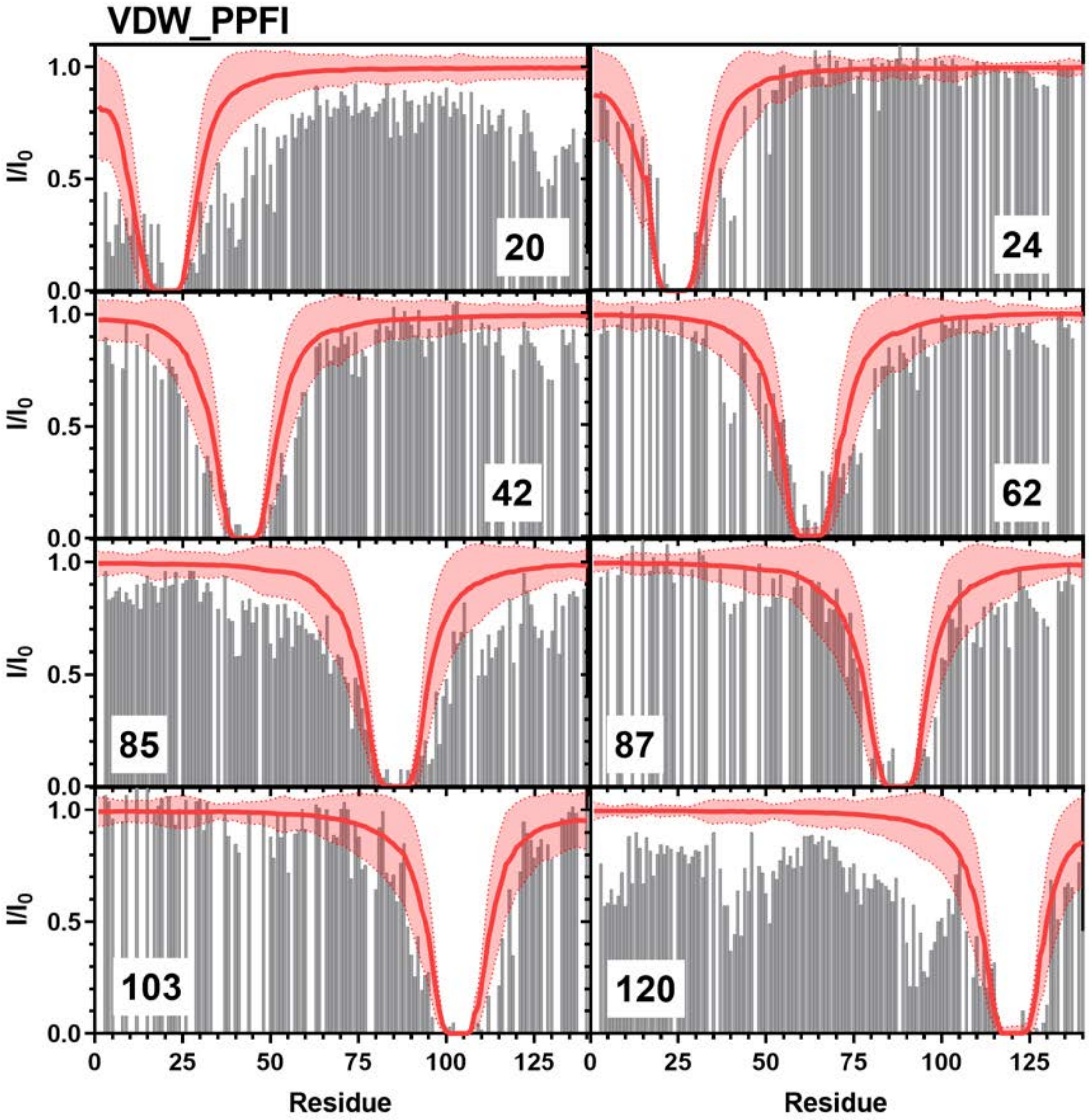


**Figure S23:** Comparison of Simulated PRE Values and Experimental PRE Values: Simulated PRE values from VDW_PPFI (red line) overlayed on top of experimental data (grey bars) from positions 20 (Top Left), 24 (Top Right), 42 (Upper Middle Left), 62 (Upper Middle Right), 85 (Lower Middle Left), 87 (Lower Middle Right), 103 (Bottom Left), 120 (Bottom Right). Experimental data for positions 20, 85, and 120 are from Sung *et al*. [8] and data for positions 24, 42, 62, 87, and 103 are from Dedmon *et al*. [9].

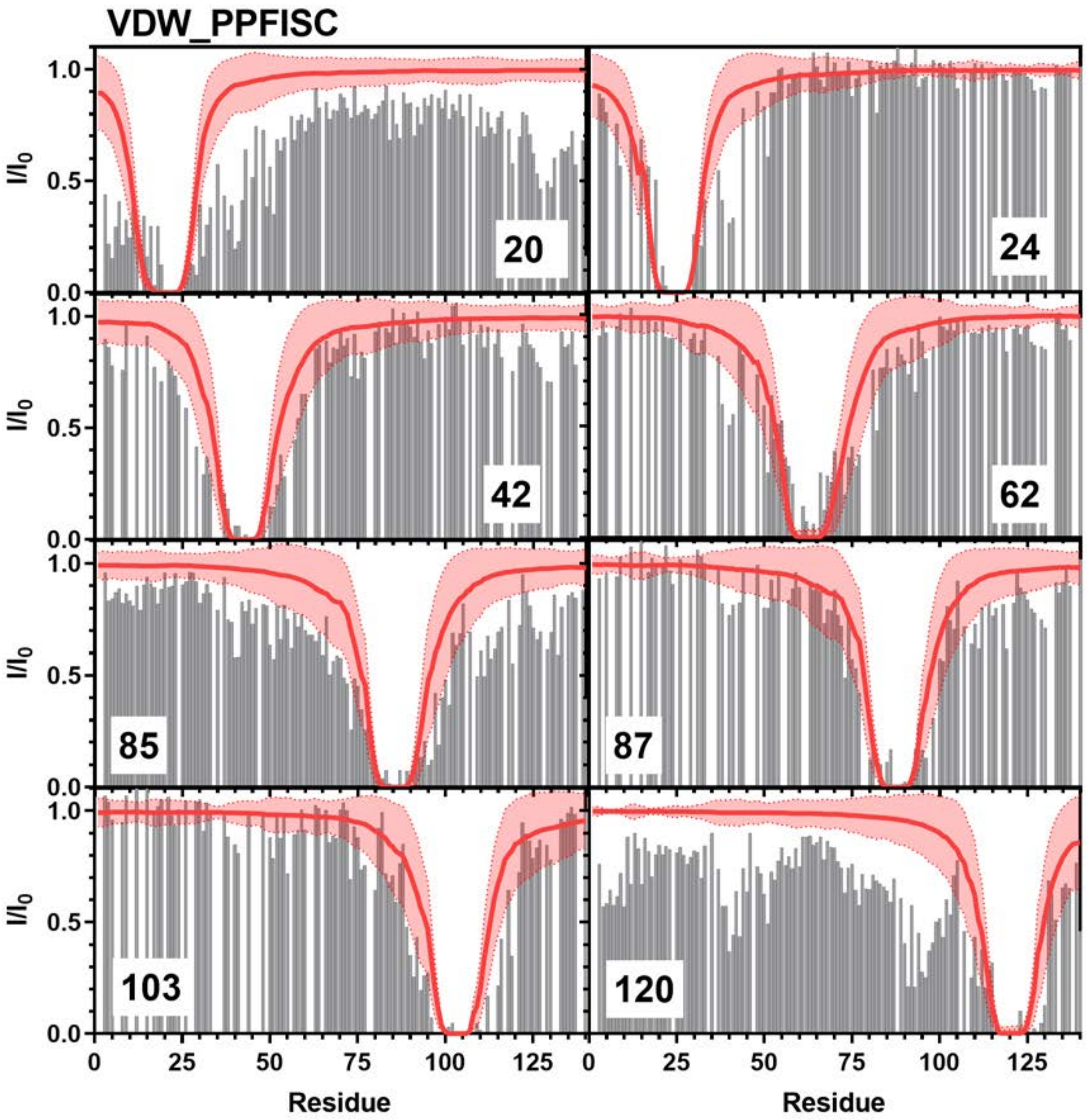


**Figure S24:** Comparison of Simulated PRE Values and Experimental PRE Values: Simulated PRE values from VDW_PPFISC (red line) overlayed on top of experimental data (grey bars) from positions 20 (Top Left), 24 (Top Right), 42 (Upper Middle Left), 62 (Upper Middle Right), 85 (Lower Middle Left), 87 (Lower Middle Right), 103 (Bottom Left), 120 (Bottom Right). Experimental data for positions 20, 85, and 120 are from Sung *et al*. [8] and data for positions 24, 42, 62, 87, and 103 are from Dedmon *et al*. [9].

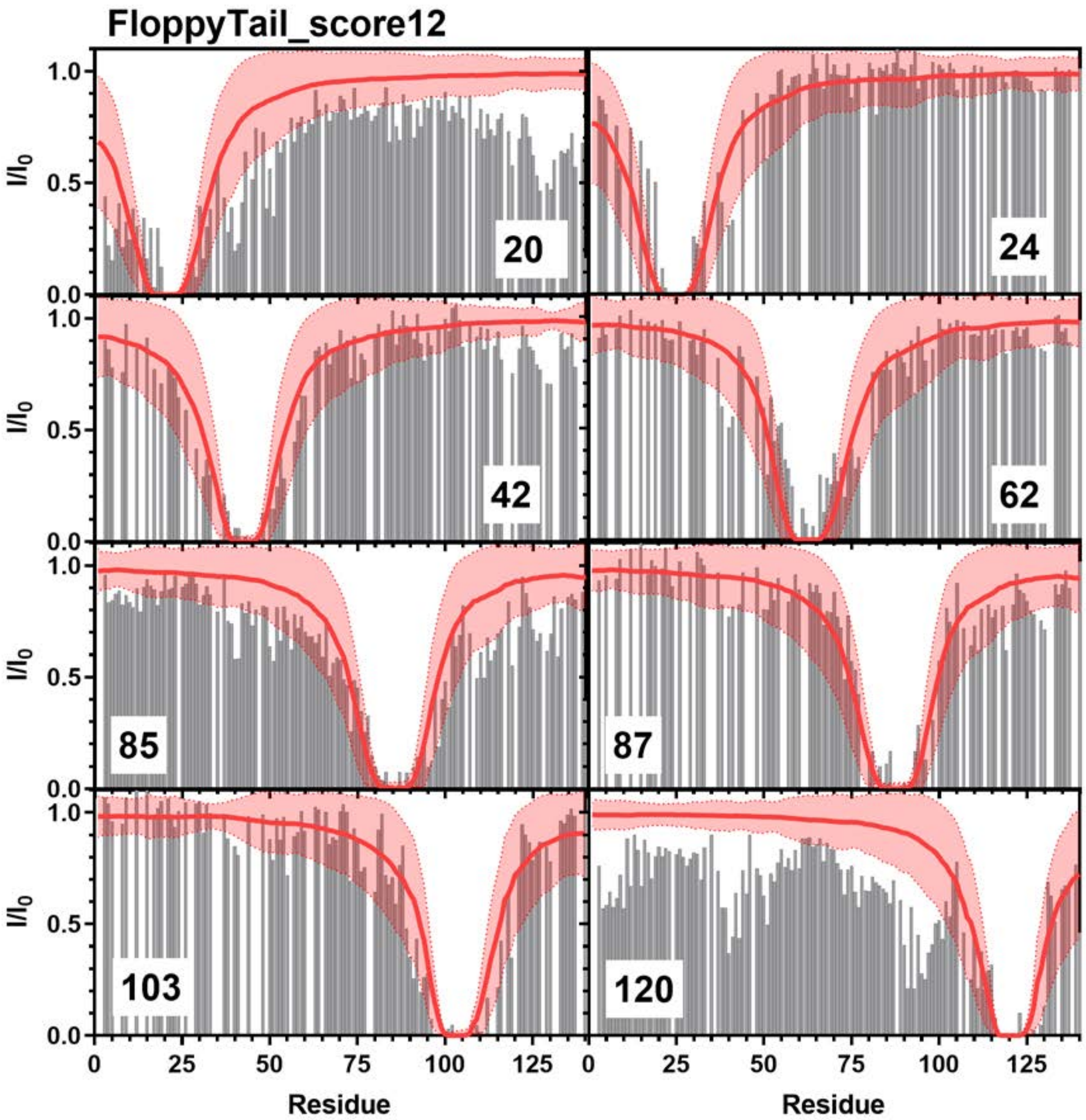


**Figure S25:** Comparison of Simulated PRE Values and Experimental PRE Values: Simulated PRE values from FloppyTail_score12 (red line) overlayed on top of experimental data (grey bars) from positions 20 (Top Left), 24 (Top Right), 42 (Upper Middle Left), 62 (Upper Middle Right), 85 (Lower Middle Left), 87 (Lower Middle Right), 103 (Bottom Left), 120 (Bottom Right). Experimental data for positions 20, 85, and 120 are from Sung *et al*. [8] and data for positions 24, 42, 62, 87, and 103 are from Dedmon *et al*. [9].

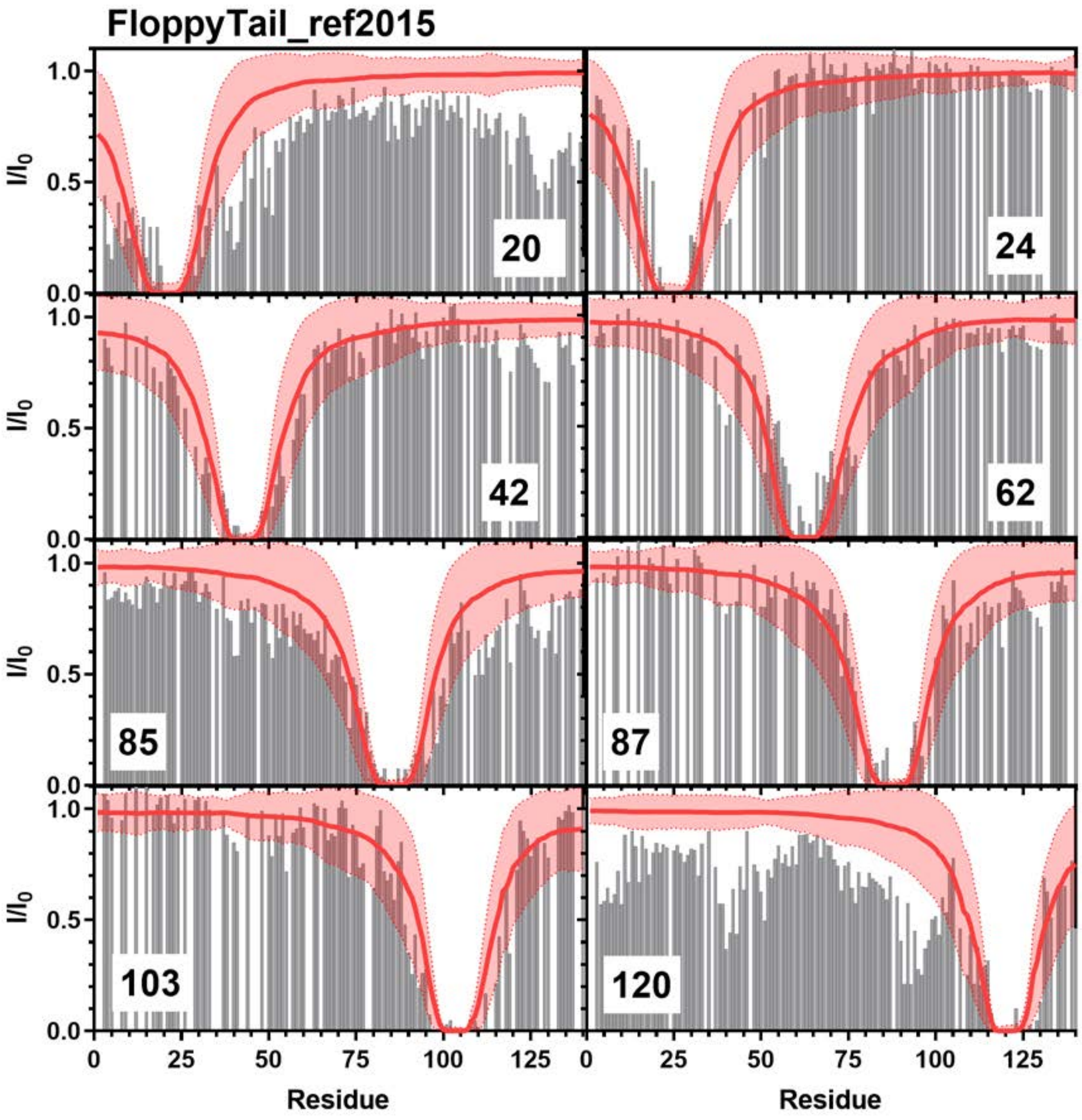


**Figure S26:** Comparison of Simulated PRE Values and Experimental PRE Values: Simulated PRE values from FloppyTail_ref2015 (red line) overlayed on top of experimental data (grey bars) from positions 20 (Top Left), 24 (Top Right), 42 (Upper Middle Left), 62 (Upper Middle Right), 85 (Lower Middle Left), 87 (Lower Middle Right), 103 (Bottom Left), 120 (Bottom Right). Experimental data for positions 20, 85, and 120 are from Sung *et al.* [8] and data for positions 24, 42, 62, 87, and 103 are from Dedmon *et al.* [9].

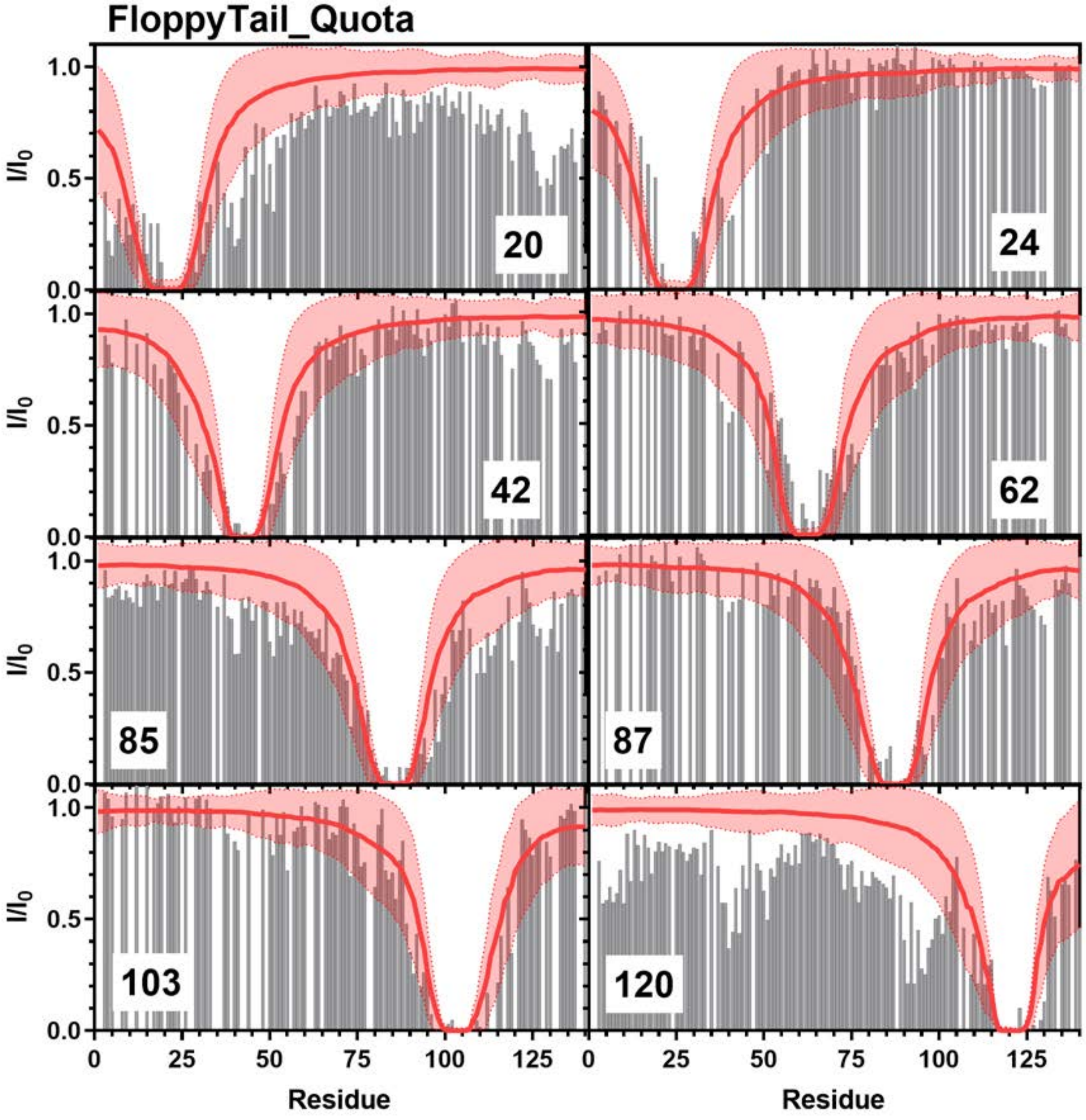


**Figure S27:** Comparison of Simulated PRE Values and Experimental PRE Values: Simulated PRE values from FloppyTail_Quota (red line) overlayed on top of experimental data (grey bars) from positions 20 (Top Left), 24 (Top Right), 42 (Upper Middle Left), 62 (Upper Middle Right), 85 (Lower Middle Left), 87 (Lower Middle Right), 103 (Bottom Left), 120 (Bottom Right). Experimental data for positions 20, 85, and 120 are from Sung *et al.* [8] and data for positions 24, 42, 62, 87, and 103 are from Dedmon *et al.* [9].

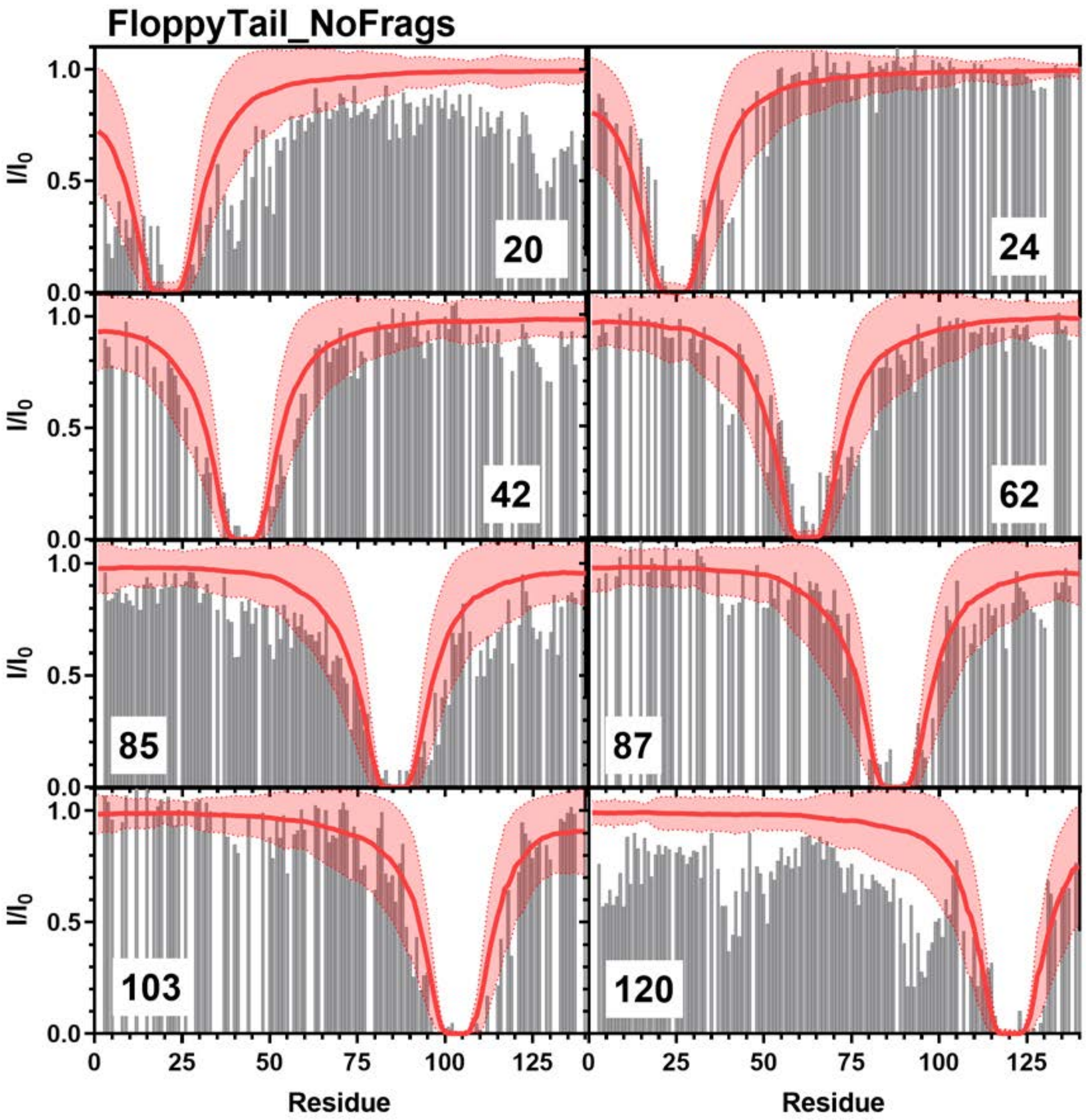


**Figure S28:** Comparison of Simulated PRE Values and Experimental PRE Values: Simulated PRE values from FloppyTail_NoFrags (red line) overlayed on top of experimental data (grey bars) from positions 20 (Top Left), 24 (Top Right), 42 (Upper Middle Left), 62 (Upper Middle Right), 85 (Lower Middle Left), 87 (Lower Middle Right), 103 (Bottom Left), 120 (Bottom Right). Experimental data for positions 20, 85, and 120 are from Sung *et al.* [8] and data for positions 24, 42, 62, 87, and 103 are from Dedmon *et al.* [9].

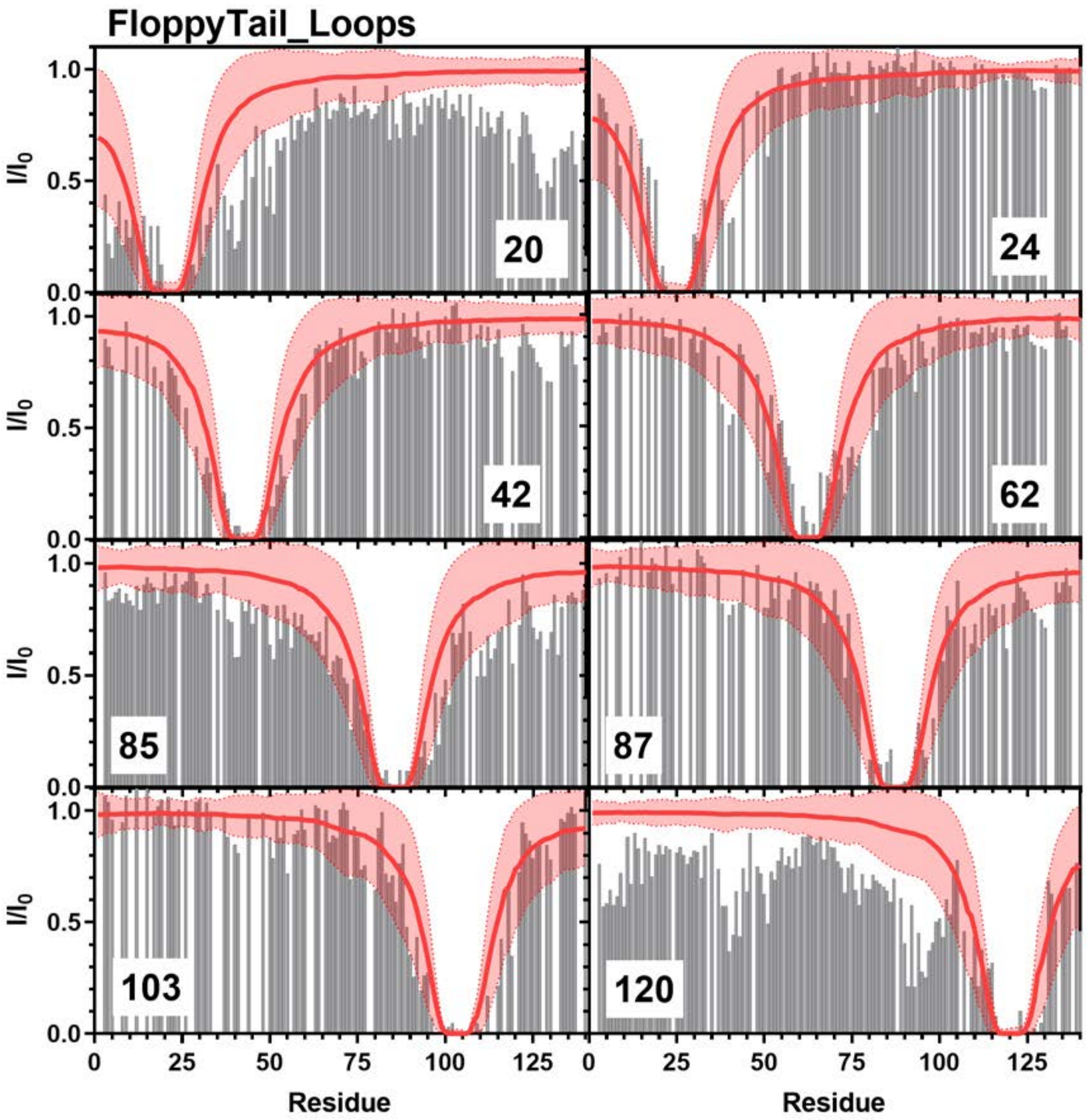


**Figure S29:** Comparison of Simulated PRE Values and Experimental PRE Values: Simulated PRE values from FloppyTail_Loops (red line) overlayed on top of experimental data (grey bars) from positions 20 (Top Left), 24 (Top Right), 42 (Upper Middle Left), 62 (Upper Middle Right), 85 (Lower Middle Left), 87 (Lower Middle Right), 103 (Bottom Left), 120 (Bottom Right). Experimental data for positions 20, 85, and 120 are from Sung *et al*. [8] and data for positions 24, 42, 62, 87, and 103 are from Dedmon *et al*. [9],

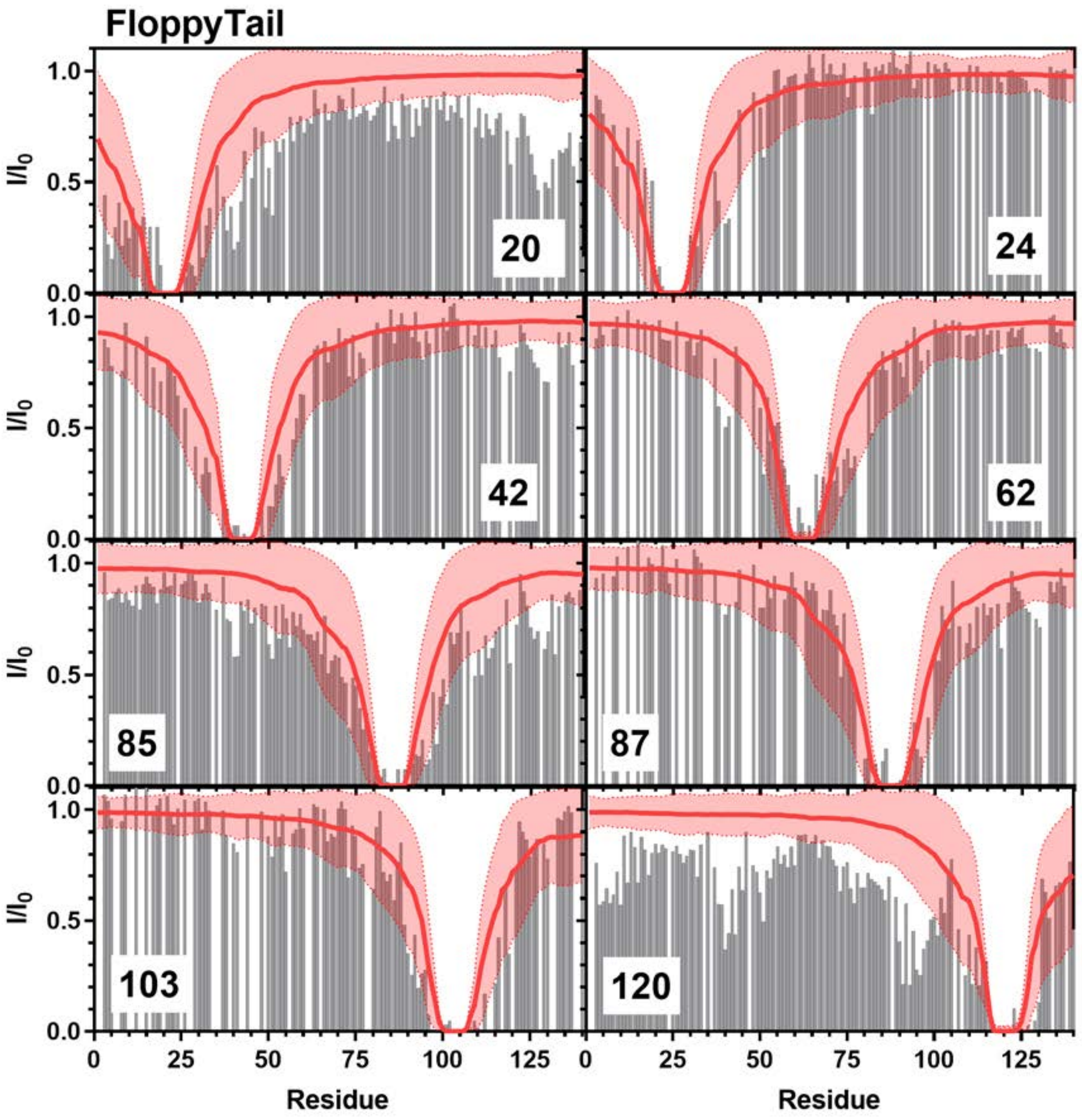


**Figure S30:** Comparison of Simulated PRE Values and Experimental PRE Values: Simulated PRE values from FloppyTail (red line) overlayed on top of experimental data (grey bars) from positions 20 (Top Left), 24 (Top Right), 42 (Upper Middle Left), 62 (Upper Middle Right), 85 (Lower Middle Left), 87 (Lower Middle Right), 103 (Bottom Left), 120 (Bottom Right). Experimental data for positions 20, 85, and 120 are from Sung *et al.* [8] and data for positions 24, 42, 62, 87, and 103 are from Dedmon *et al.* [9].

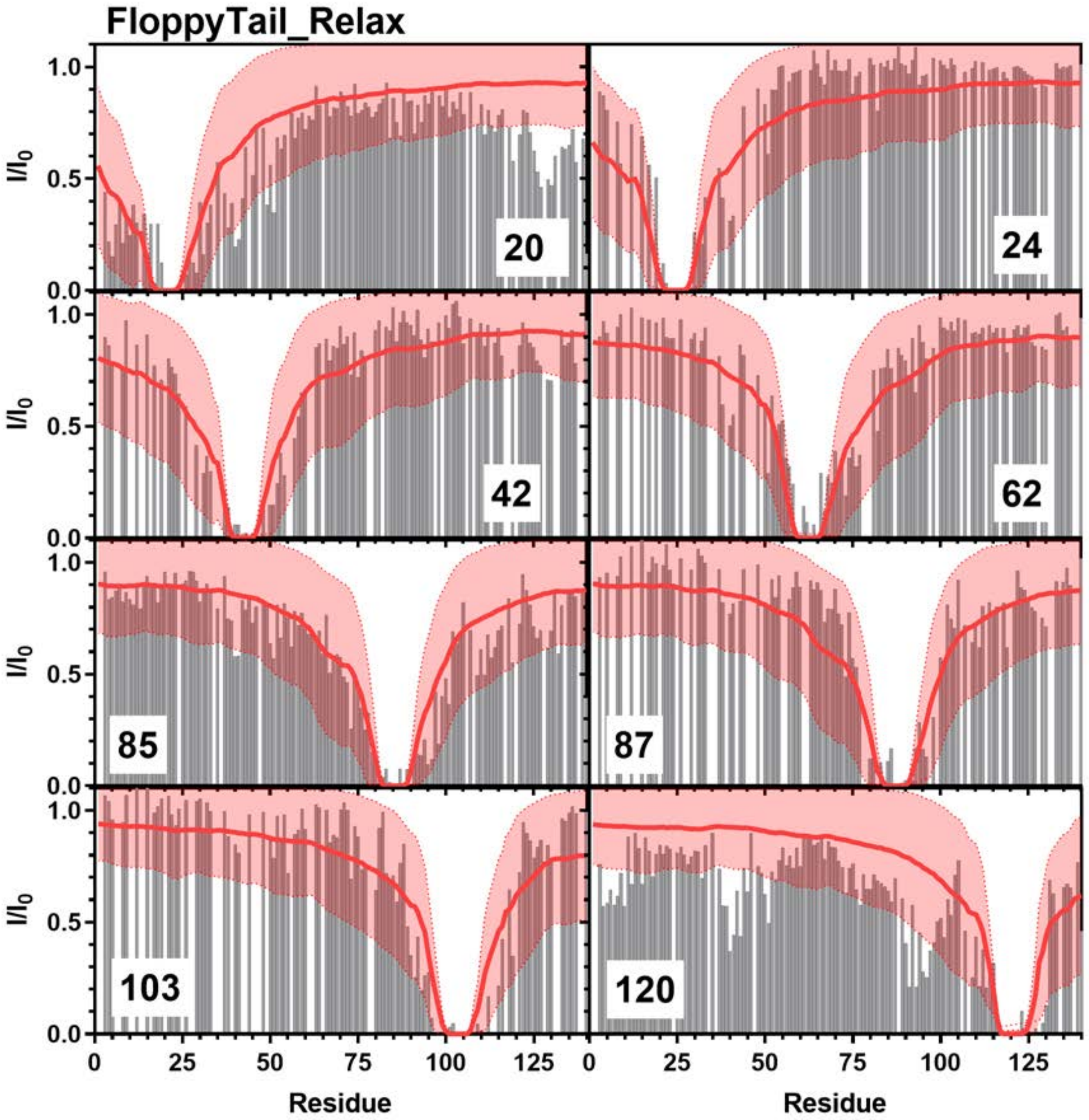


**Figure S31:** Comparison of Simulated PRE Values and Experimental PRE Values: Simulated PRE values from FloppyTail_Relax (red line) overlayed on top of experimental data (grey bars) from positions 20 (Top Left), 24 (Top Right), 42 (Upper Middle Left), 62 (Upper Middle Right), 85 (Lower Middle Left), 87 (Lower Middle Right), 103 (Bottom Left), 120 (Bottom Right). Experimental data for positions 20, 85, and 120 are from Sung *et al*. [8] and data for positions 24, 42, 62, 87, and 103 are from Dedmon *et al*. [9].

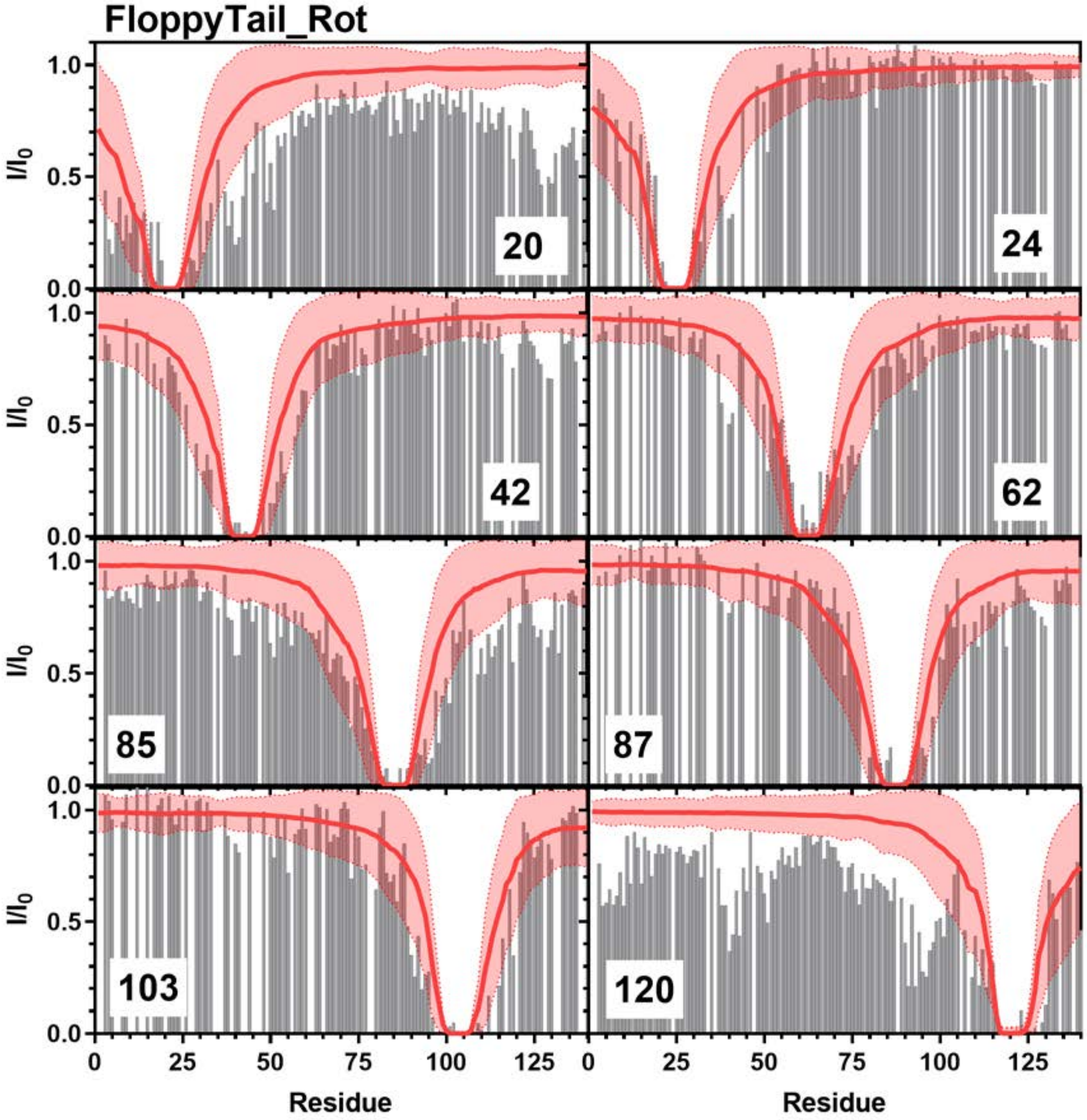


**Figure S32:** Comparison of Simulated PRE Values and Experimental PRE Values: Simulated PRE values from FloppyTail_Rot (red line) overlayed on top of experimental data (grey bars) from positions 20 (Top Left), 24 (Top Right), 42 (Upper Middle Left), 62 (Upper Middle Right), 85 (Lower Middle Left), 87 (Lower Middle Right), 103 (Bottom Left), 120 (Bottom Right). Experimental data for positions 20, 85, and 120 are from Sung *et al*. [8] and data for positions 24, 42, 62, 87, and 103 are from Dedmon *et al*. [9].

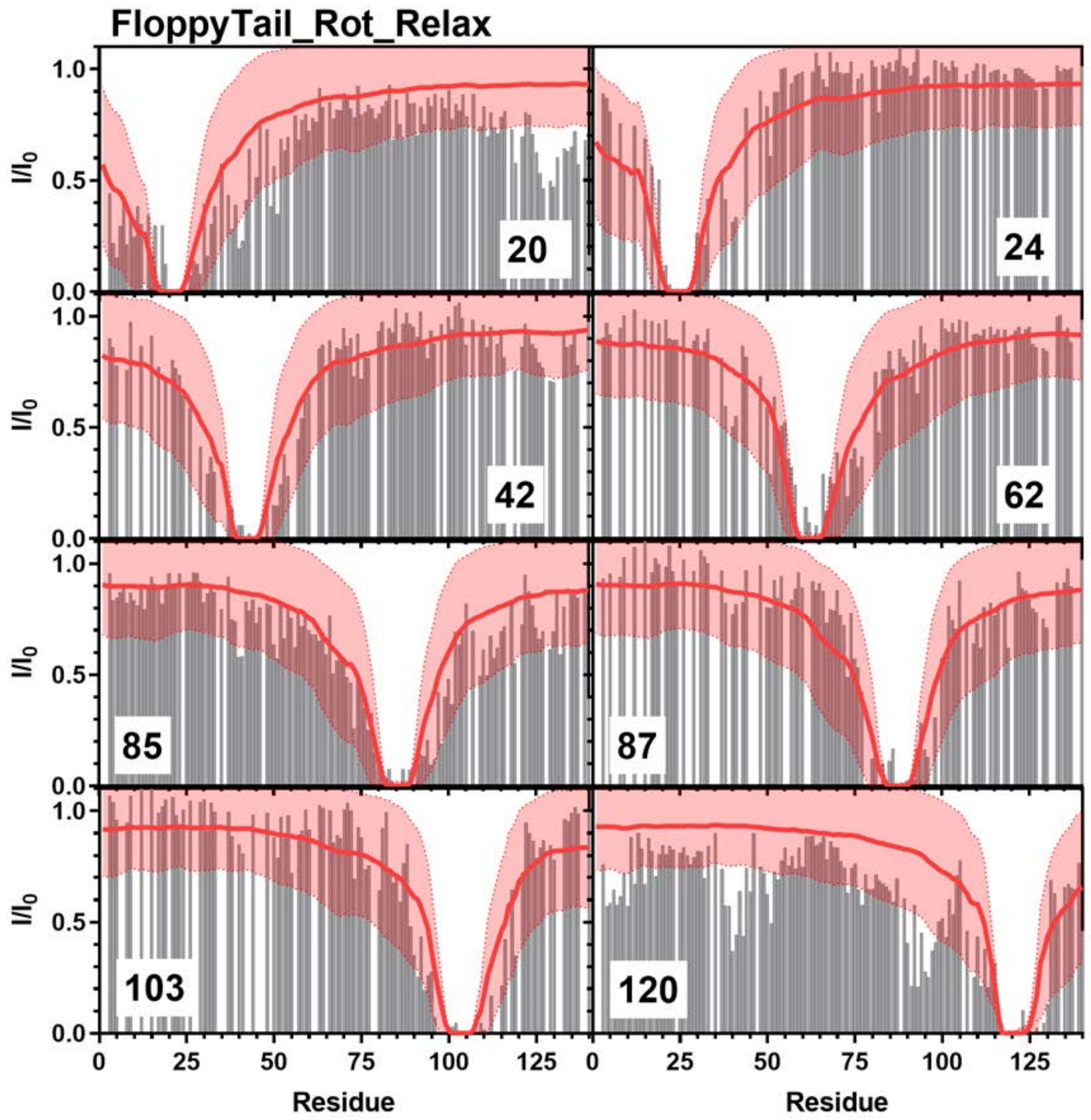


**Figure S33:** Comparison of Simulated PRE Values and Experimental PRE Values: Simulated PRE values from FloppyTail_Rot_Relax (red line) overlayed on top of experimental data (grey bars) from positions 20 (Top Left), 24 (Top Right), 42 (Upper Middle Left), 62 (Upper Middle Right), 85 (Lower Middle Left), 87 (Lower Middle Right), 103 (Bottom Left), 120 (Bottom Right). Experimental data for positions 20, 85, and 120 are from Sung *et al*. [8] and data for positions 24, 42, 62, 87, and 103 are from Dedmon *et al*. [9].

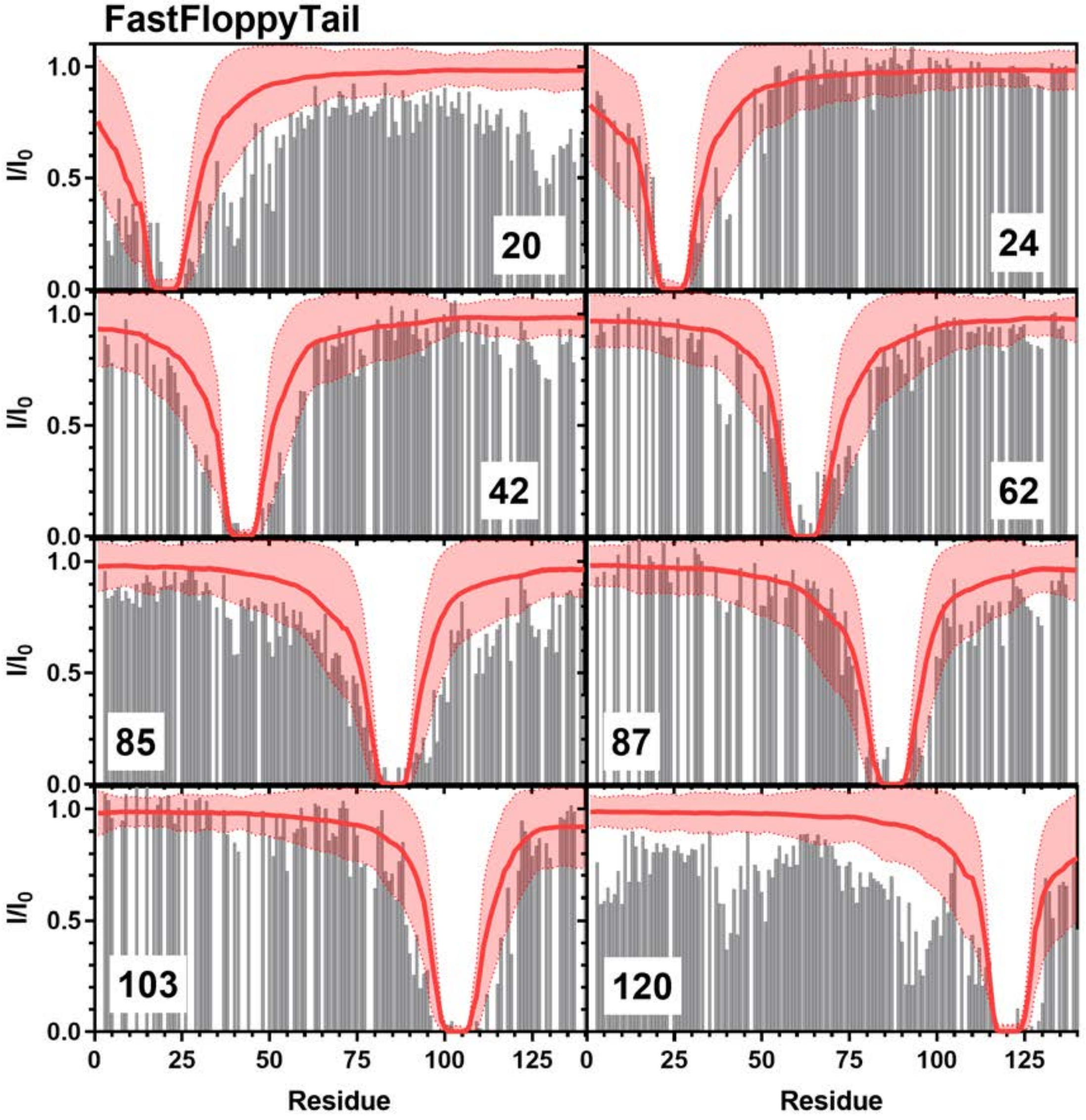


**Figure S34:** Comparison of Simulated PRE Values and Experimental PRE Values: Simulated PRE values from FastFloppyTail (red line) overlayed on top of experimental data (grey bars) from positions 20 (Top Left), 24 (Top Right), 42 (Upper Middle Left), 62 (Upper Middle Right), 85 (Lower Middle Left), 87 (Lower Middle Right), 103 (Bottom Left), 120 (Bottom Right). Experimental data for positions 20, 85, and 120 are from Sung *et al*. [8] and data for positions 24, 42, 62, 87, and 103 are from Dedmon *et al*. [9].

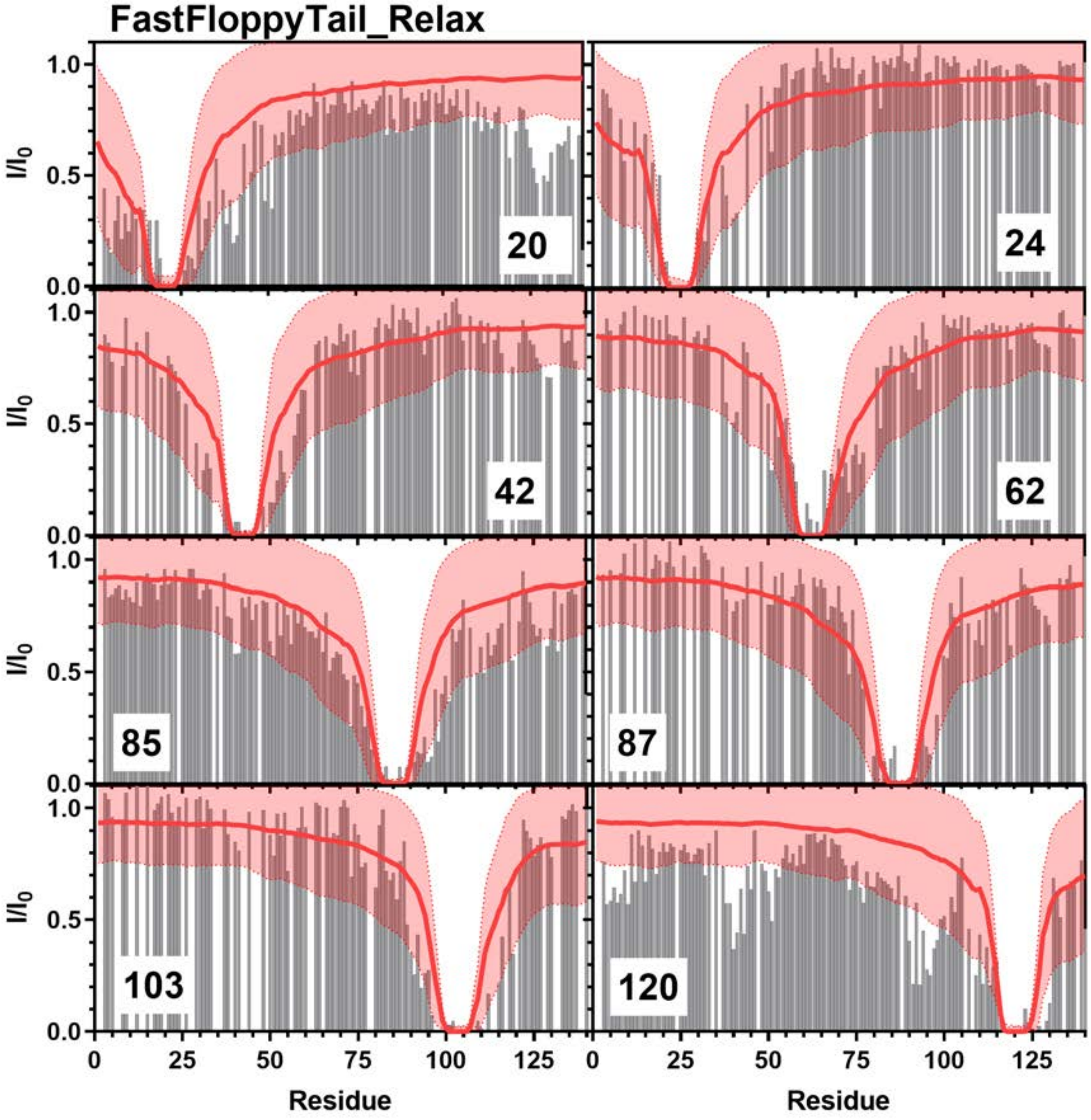


**Figure S35:** Comparison of Simulated PRE Values and Experimental PRE Values: Simulated PRE values from FastFloppyTail (red line) overlayed on top of experimental data (grey bars) from positions 20 (Top Left), 24 (Top Right), 42 (Upper Middle Left), 62 (Upper Middle Right), 85 (Lower Middle Left), 87 (Lower Middle Right), 103 (Bottom Left), 120 (Bottom Right). Experimental data for positions 20, 85, and 120 are from Sung *et al*. [8] and data for positions 24, 42, 62, 87, and 103 are from Dedmon *et al*. [9].

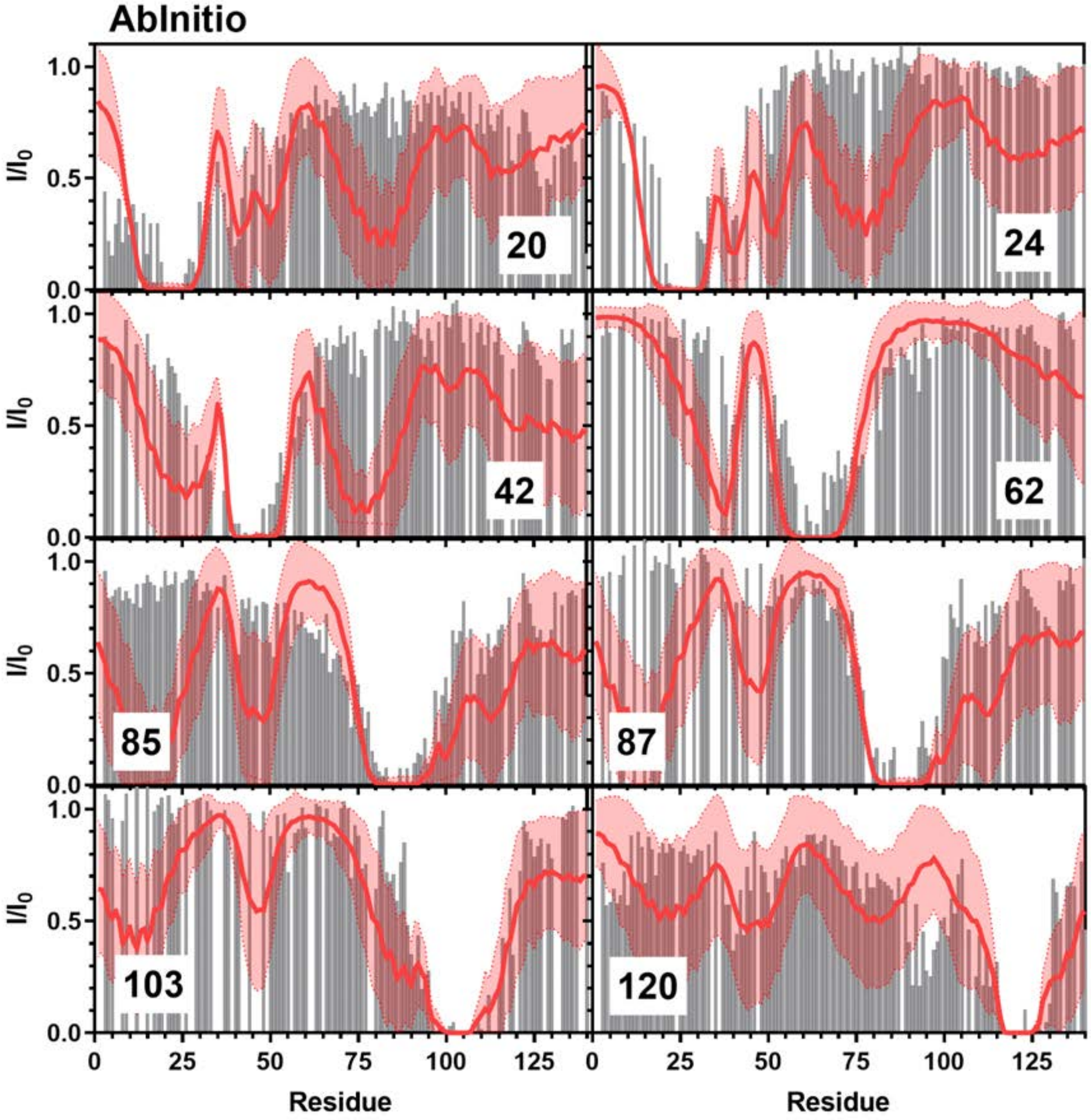


**Figure S36:** Comparison of Simulated PRE Values and Experimental PRE Values: Simulated PRE values from AbInitio (red line) overlayed on top of experimental data (grey bars) from positions 20 (Top Left), 24 (Top Right), 42 (Upper Middle Left), 62 (Upper Middle Right), 85 (Lower Middle Left), 87 (Lower Middle Right), 103 (Bottom Left), 120 (Bottom Right). Experimental data for positions 20, 85, and 120 are from Sung *et al.* [8] and data for positions 24, 42, 62, 87, and 103 are from Dedmon *et al.* [9].

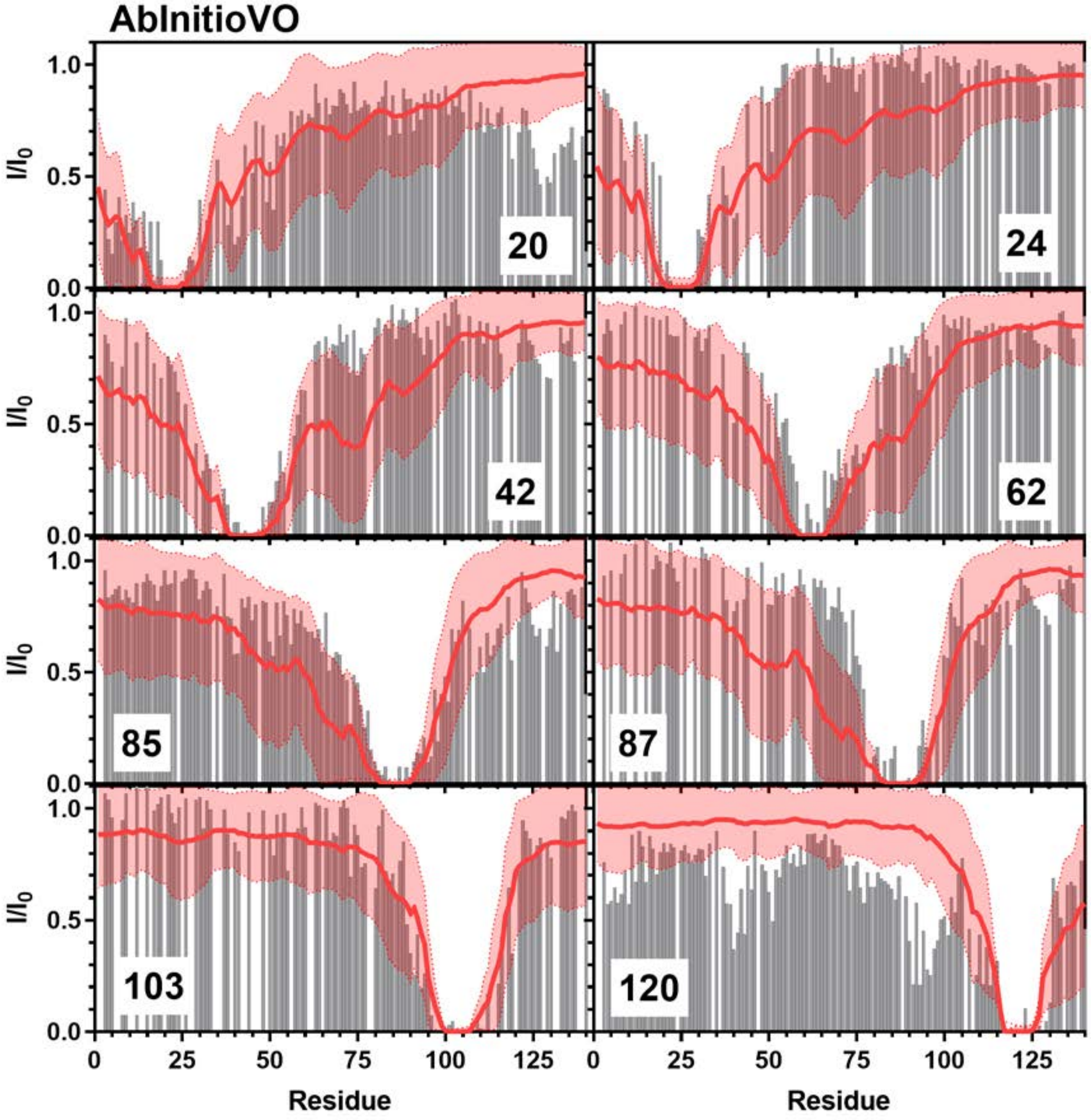


**Figure S37:** Comparison of Simulated PRE Values and Experimental PRE Values: Simulated PRE values from AbInitioVO (red line) overlayed on top of experimental data (grey bars) from positions 20 (Top Left), 24 (Top Right), 42 (Upper Middle Left), 62 (Upper Middle Right), 85 (Lower Middle Left), 87 (Lower Middle Right), 103 (Bottom Left), 120 (Bottom Right). Experimental data for positions 20, 85, and 120 are from Sung *et al*. [8] and data for positions 24, 42, 62, 87, and 103 are from Dedmon *et al*. [9].

**Comparison with αS Chemical Shift Data**

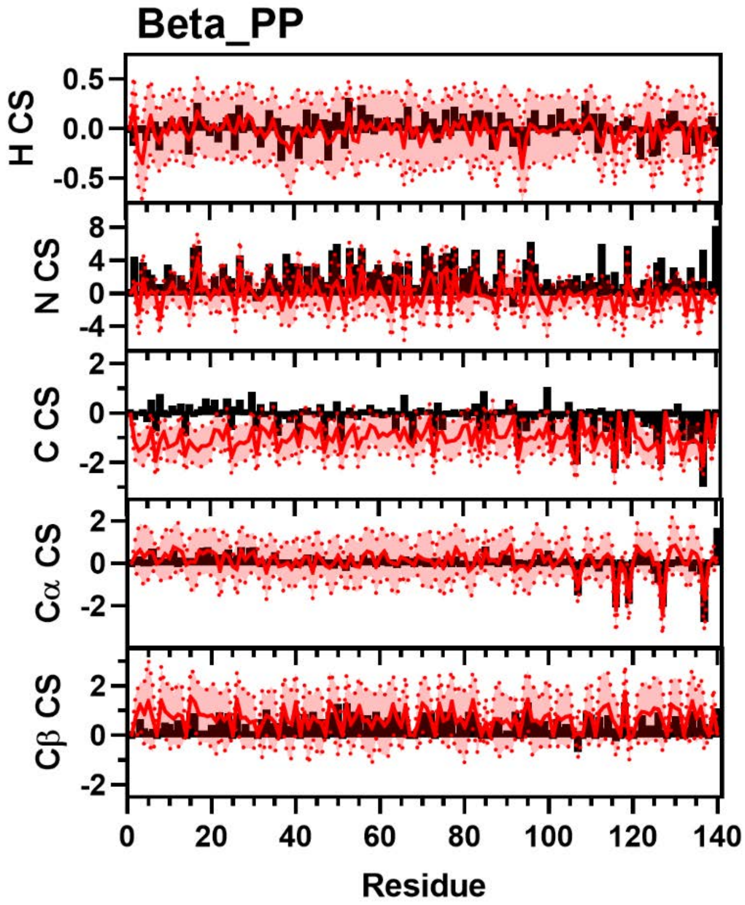


**Figure S38:** Comparison of Simulated and Experimental NMR Chemical Shift Data. Simulated Chemical Shift values from the Beta_PP ensemble (red) overlayed on experimental data (black bars) of N (Top), H (Upper Middle), C (Middle), Cα (Lower Middle), and Cβ (Bottom) chemical shifts from Sung *et al*. [8] Neighbor corrected random coil chemical shift values have been subtracted from both simulated and experimental data [11].

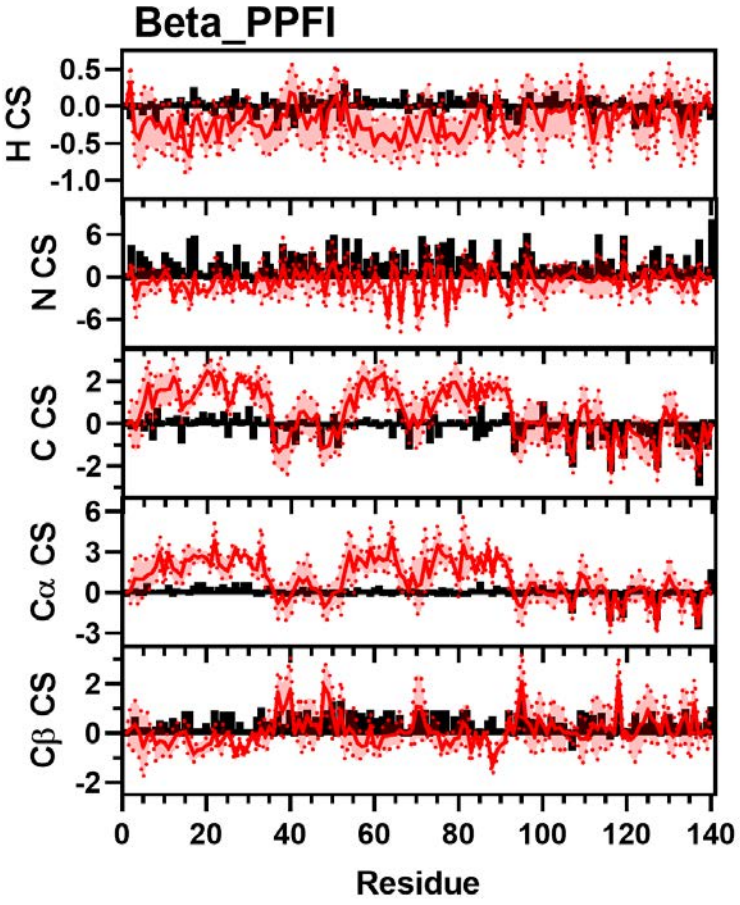


**Figure S39:** Comparison of Simulated and Experimental NMR Chemical Shift Data. Simulated Chemical Shift values from the Beta_PPFI ensemble (red) overlayed on experimental data (black bars) of N (Top), H (Upper Middle), C (Middle), Cα (Lower Middle), and Cβ (Bottom) chemical shifts from Sung *et al*. [8] Neighbor corrected random coil chemical shift values have been subtracted from both simulated and experimental data [11].

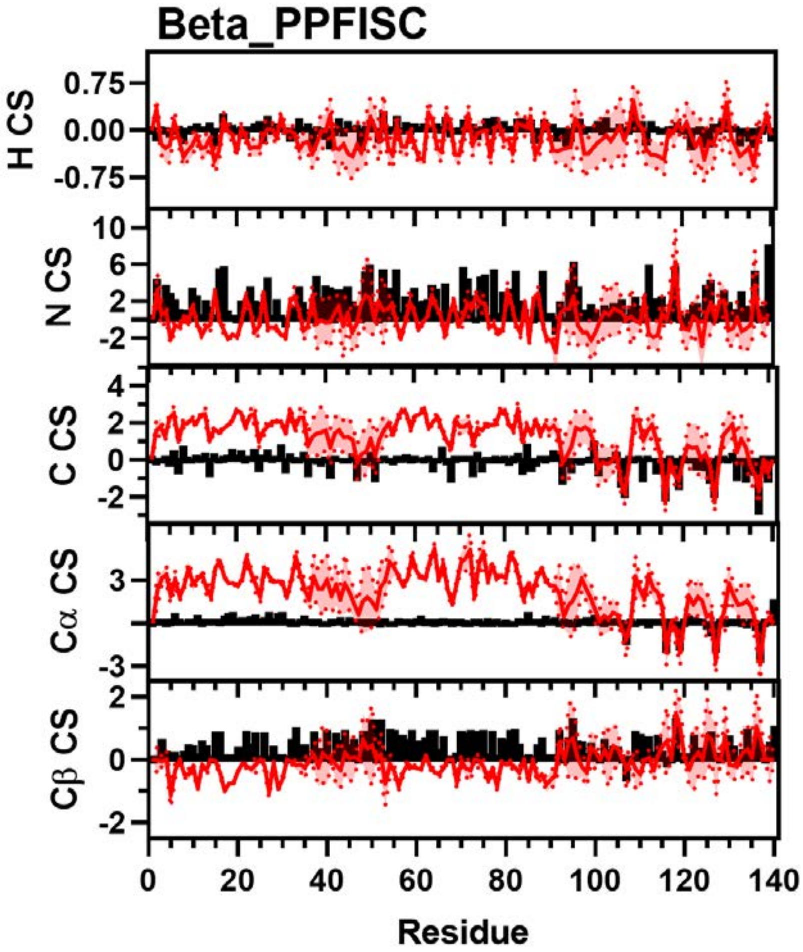


**Figure S40:** Comparison of Simulated and Experimental NMR Chemical Shift Data. Simulated Chemical Shift values from the Beta_PPFISC ensemble (red) overlayed on experimental data (black bars) of N (Top), H (Upper Middle), C (Middle), Cα (Lower Middle), and Cβ (Bottom) chemical shifts from Sung *et al*. [8] Neighbor corrected random coil chemical shift values have been subtracted from both simulated and experimental data [11].

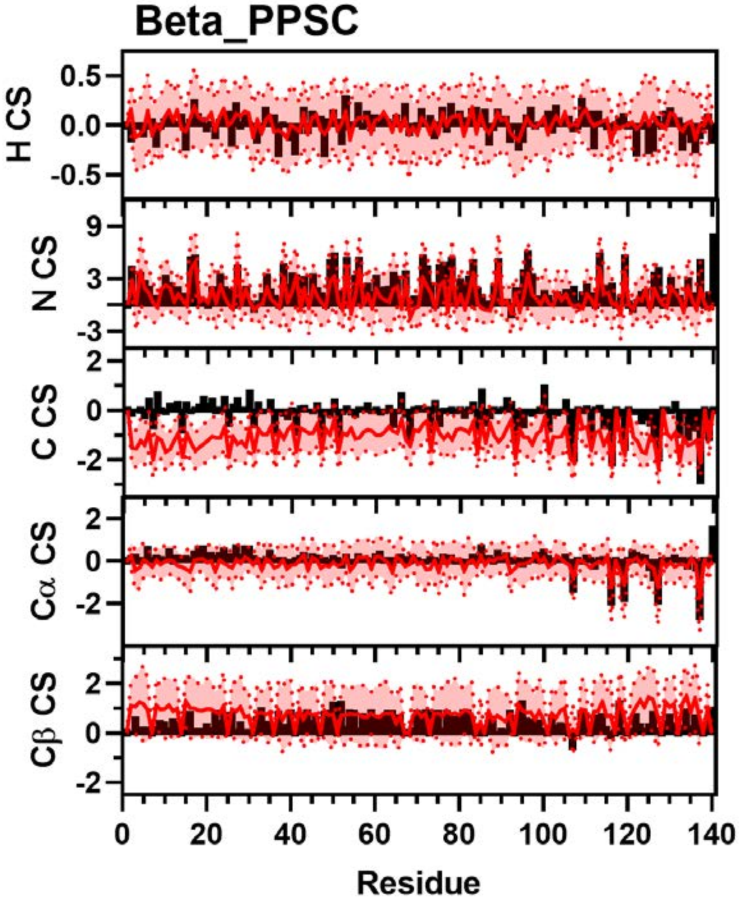


**Figure S41:** Comparison of Simulated and Experimental NMR Chemical Shift Data. Simulated Chemical Shift values from the Beta_PPSC ensemble (red) overlayed on experimental data (black bars) of N (Top), H (Upper Middle), C (Middle), Cα (Lower Middle), and Cβ (Bottom) chemical shifts from Sung *et al*. [8] Neighbor corrected random coil chemical shift values have been subtracted from both simulated and experimental data [11].

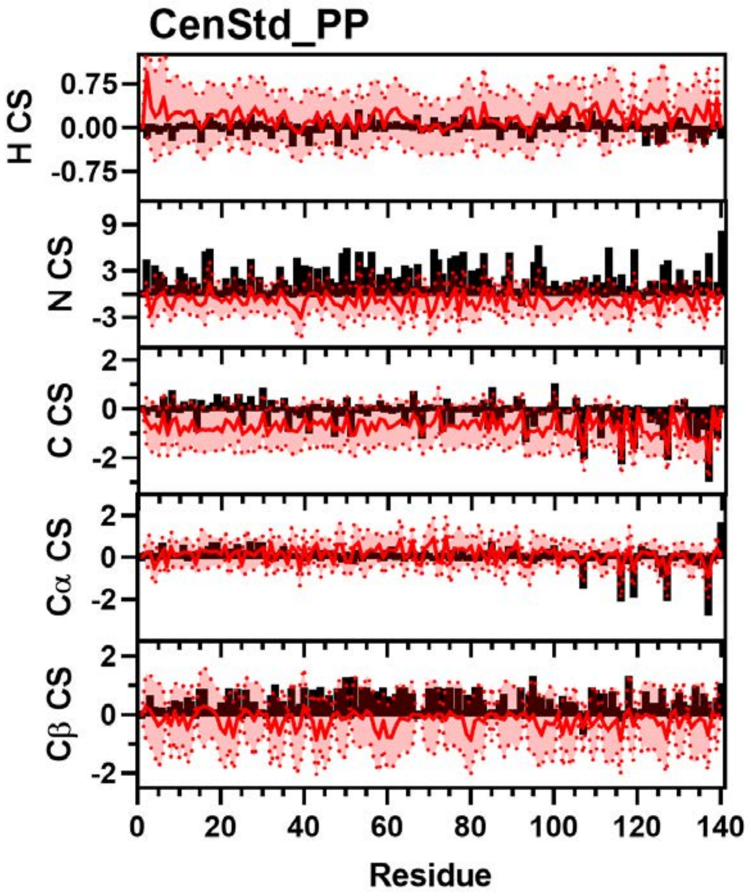


**Figure S42:** Comparison of Simulated and Experimental NMR Chemical Shift Data. Simulated Chemical Shift values from the CenStd_PP ensemble (red) overlayed on experimental data (black bars) of N (Top), H (Upper Middle), C (Middle), Cα (Lower Middle), and Cβ (Bottom) chemical shifts from Sung *et al*. [8] Neighbor corrected random coil chemical shift values have been subtracted from both simulated and experimental data [11].

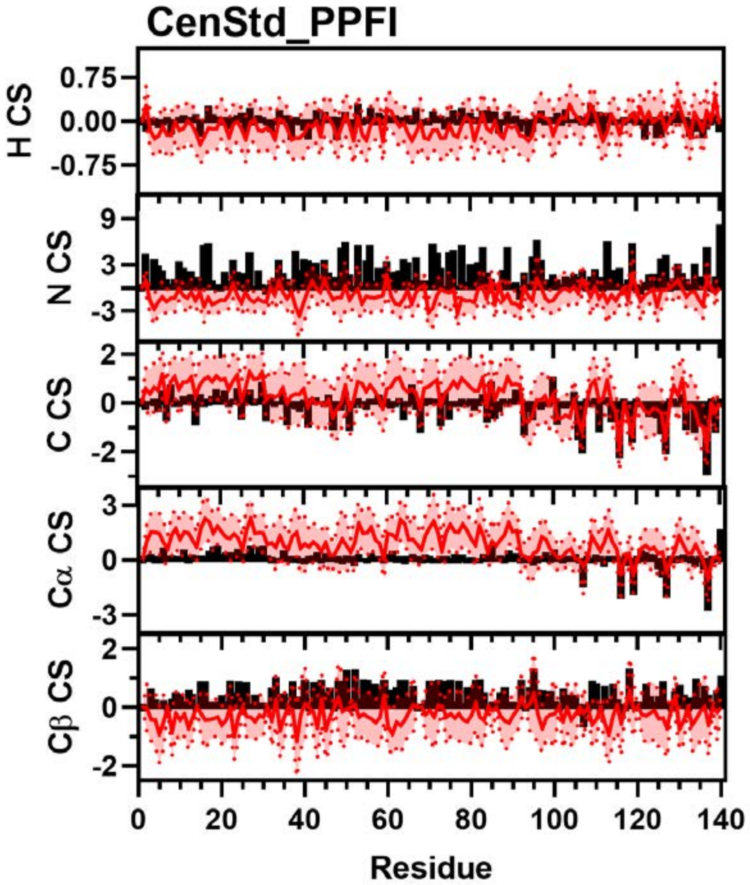


**Figure S43:** Comparison of Simulated and Experimental NMR Chemical Shift Data. Simulated Chemical Shift values from the CenStd_PPFI ensemble (red) overlayed on experimental data (black bars) of N (Top), H (Upper Middle), C (Middle), Cα (Lower Middle), and Cβ (Bottom) chemical shifts from Sung *et al*. [8] Neighbor corrected random coil chemical shift values have been subtracted from both simulated and experimental data [11].

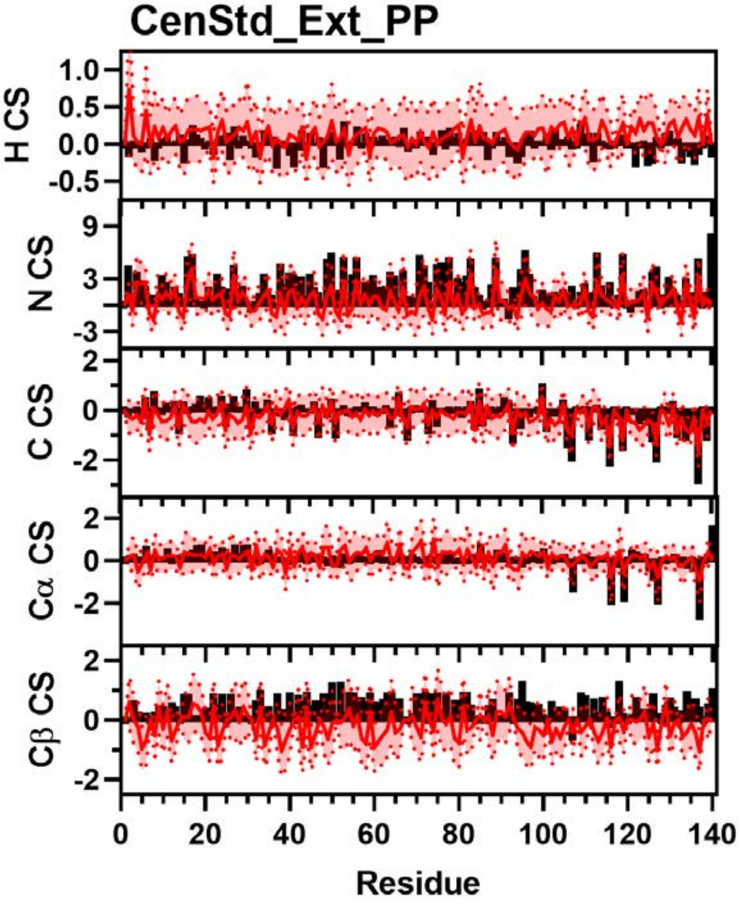


**Figure S44:** Comparison of Simulated and Experimental NMR Chemical Shift Data. Simulated Chemical Shift values from the CenStd_Ext_PP ensemble (red) overlayed on experimental data (black bars) of N (Top), H (Upper Middle), C (Middle), Cα (Lower Middle), and Cβ (Bottom) chemical shifts from Sung *et al*. [8] Neighbor corrected random coil chemical shift values have been subtracted from both simulated and experimental data [11].

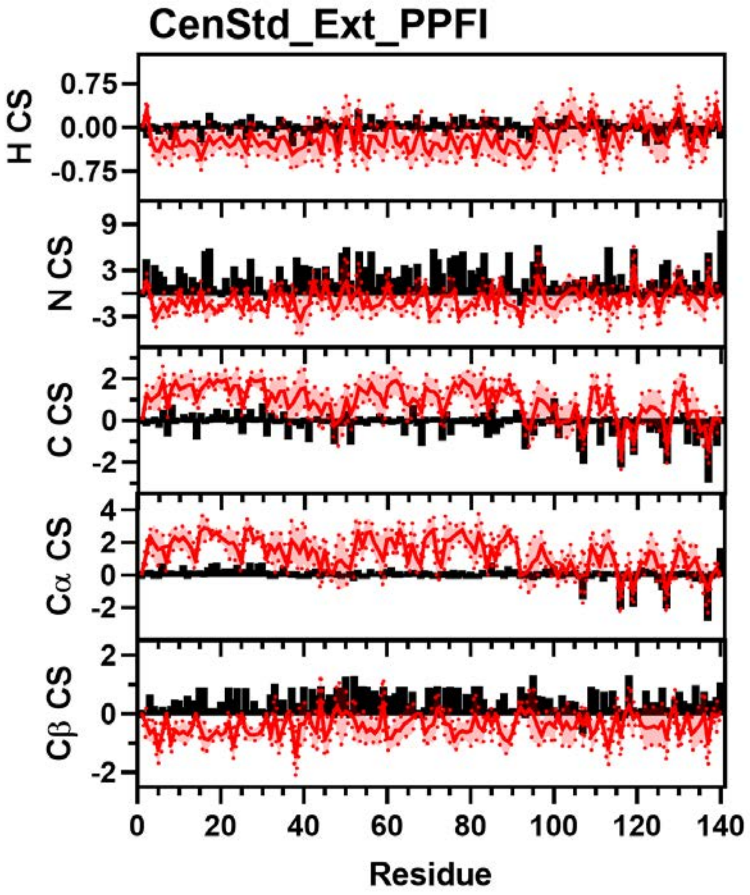


**Figure S45:** Comparison of Simulated and Experimental NMR Chemical Shift Data. Simulated Chemical Shift values from the CenStd_Ext_PPFI ensemble (red) overlayed on experimental data (black bars) of N (Top), H (Upper Middle), C (Middle), Cα (Lower Middle), and Cβ (Bottom) chemical shifts from Sung *et al*. [8] Neighbor corrected random coil chemical shift values have been subtracted from both simulated and experimental data [11].

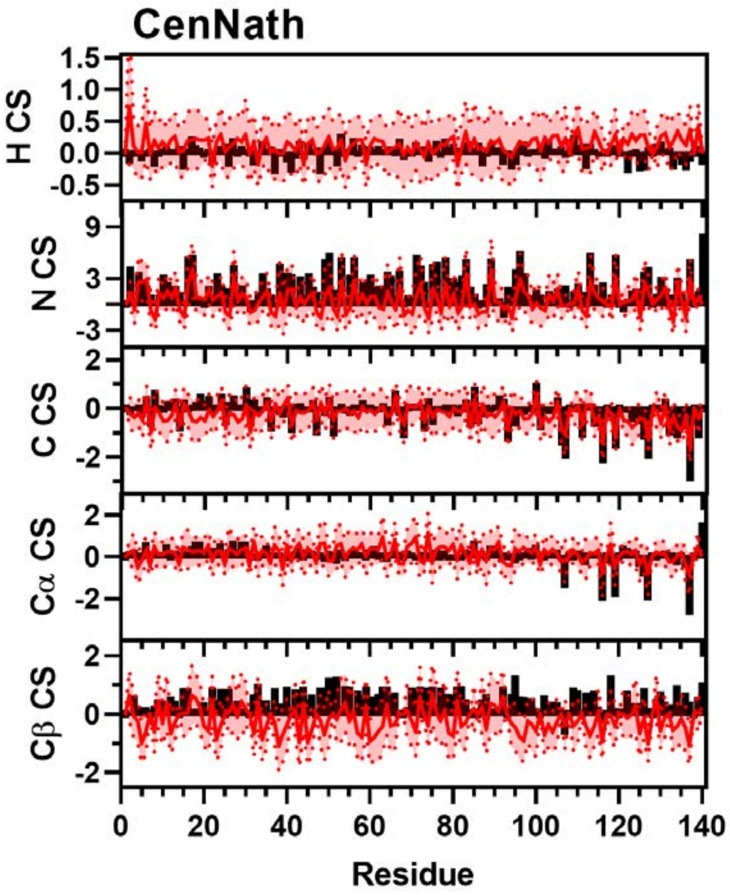


**Figure S46:** Comparison of Simulated and Experimental NMR Chemical Shift Data. Simulated Chemical Shift values from the CenNath ensemble (red) overlayed on experimental data (black bars) of N (Top), H (Upper Middle), C (Middle), Cα (Lower Middle), and Cβ (Bottom) chemical shifts from Sung *et al*. [8] Neighbor corrected random coil chemical shift values have been subtracted from both simulated and experimental data [11].

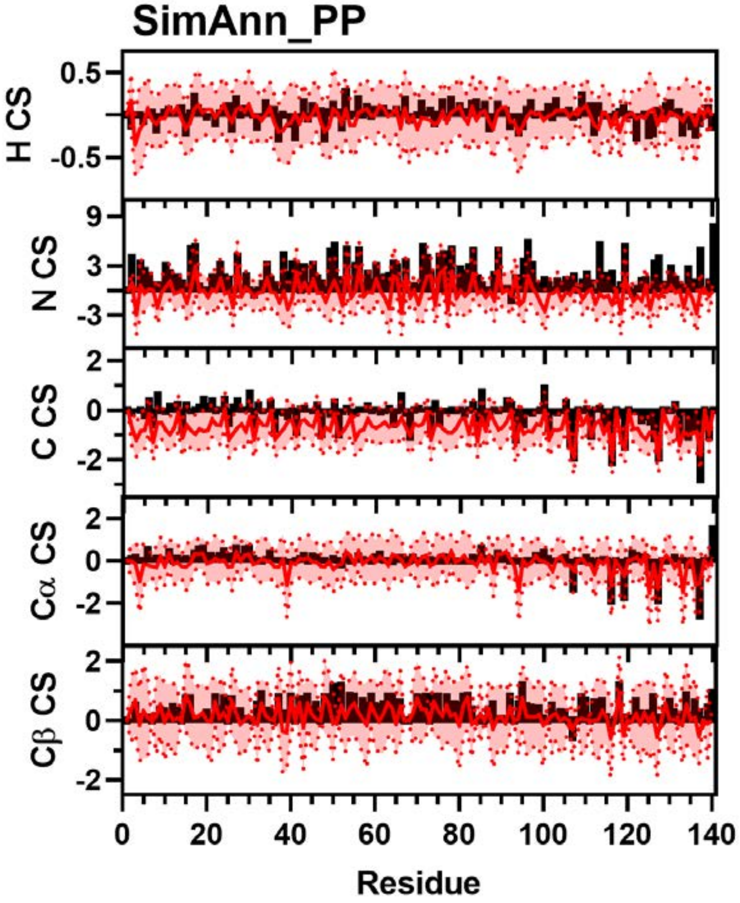


**Figure S47:** Comparison of Simulated and Experimental NMR Chemical Shift Data. Simulated Chemical Shift values from the SimAnn_PP ensemble (red) overlayed on experimental data (black bars) of N (Top), H (Upper Middle), C (Middle), Cα (Lower Middle), and Cβ (Bottom) chemical shifts from Sung *et al*. [8] Neighbor corrected random coil chemical shift values have been subtracted from both simulated and experimental data [11].

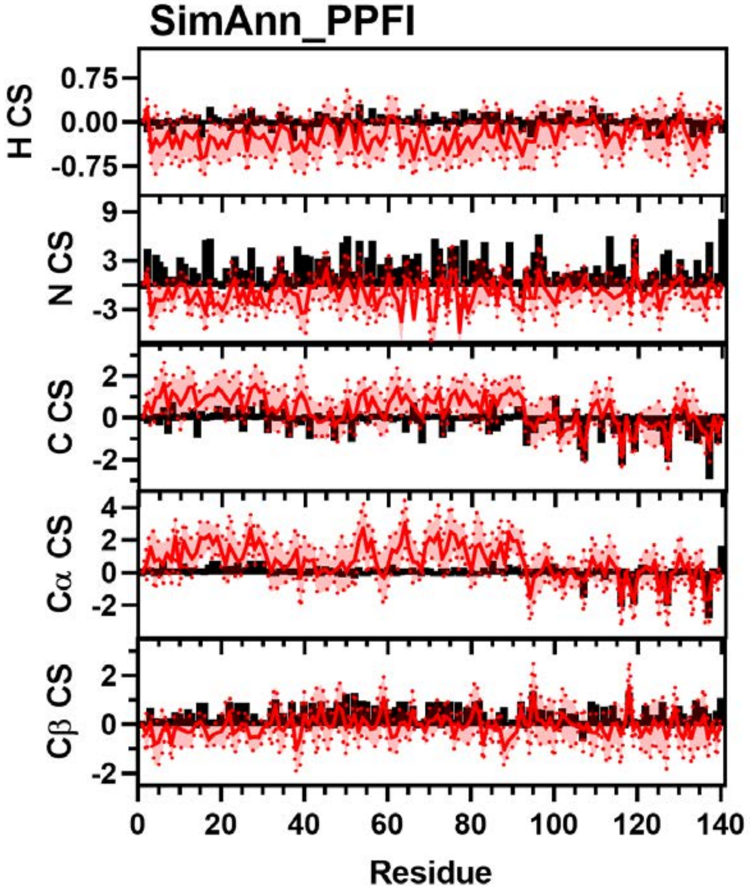


**Figure S48:** Comparison of Simulated and Experimental NMR Chemical Shift Data. Simulated Chemical Shift values from the SimAnn_PPFI ensemble (red) overlayed on experimental data (black bars) of N (Top), H (Upper Middle), C (Middle), Cα (Lower Middle), and Cβ (Bottom) chemical shifts from Sung *et al*. [8] Neighbor corrected random coil chemical shift values have been subtracted from both simulated and experimental data [11].

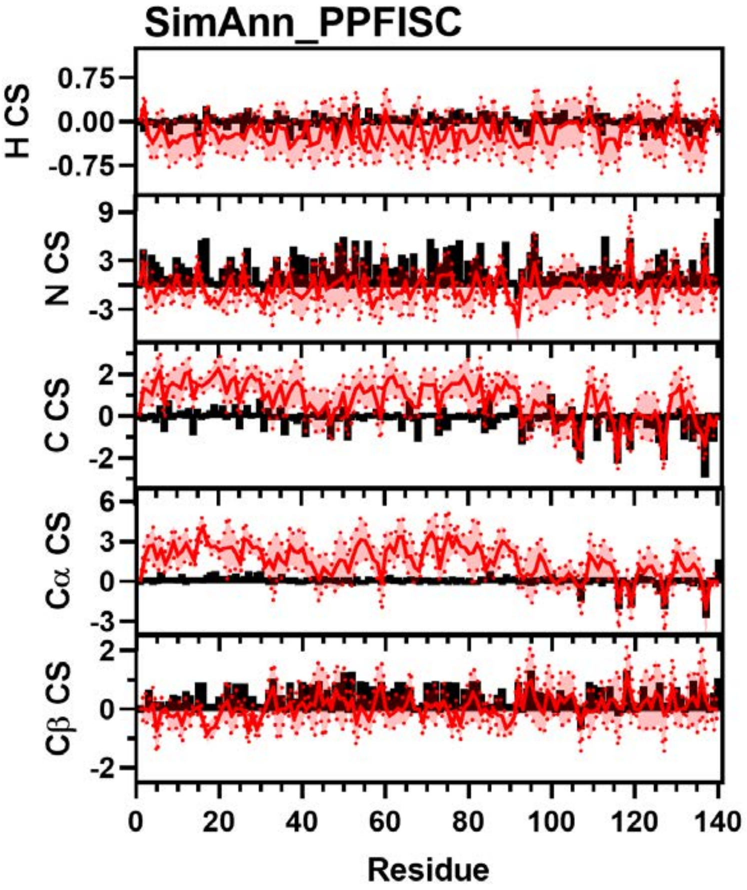


**Figure S49:** Comparison of Simulated and Experimental NMR Chemical Shift Data. Simulated Chemical Shift values from the SimAnn_PPFISC ensemble (red) overlayed on experimental data (black bars) of N (Top), H (Upper Middle), C (Middle), Cα (Lower Middle), and Cβ (Bottom) chemical shifts from Sung *et al*. [8] Neighbor corrected random coil chemical shift values have been subtracted from both simulated and experimental data [11].

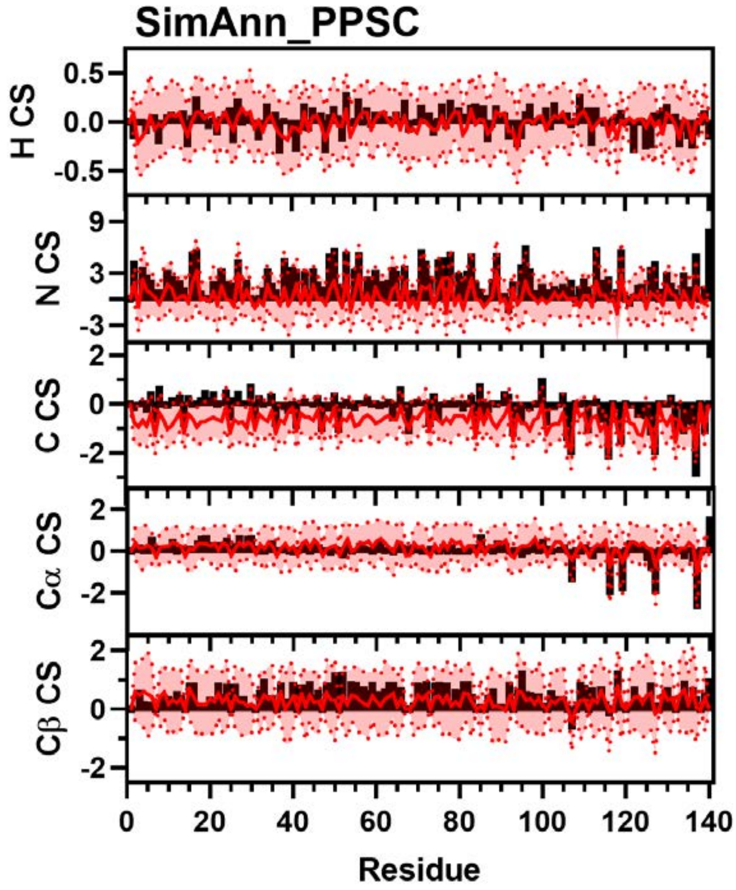


**Figure S50:** Comparison of Simulated and Experimental NMR Chemical Shift Data. Simulated Chemical Shift values from the SimAnn_PPSC ensemble (red) overlayed on experimental data (black bars) of N (Top), H (Upper Middle), C (Middle), Cα (Lower Middle), and Cβ (Bottom) chemical shifts from Sung *et al*. [8] Neighbor corrected random coil chemical shift values have been subtracted from both simulated and experimental data [11].

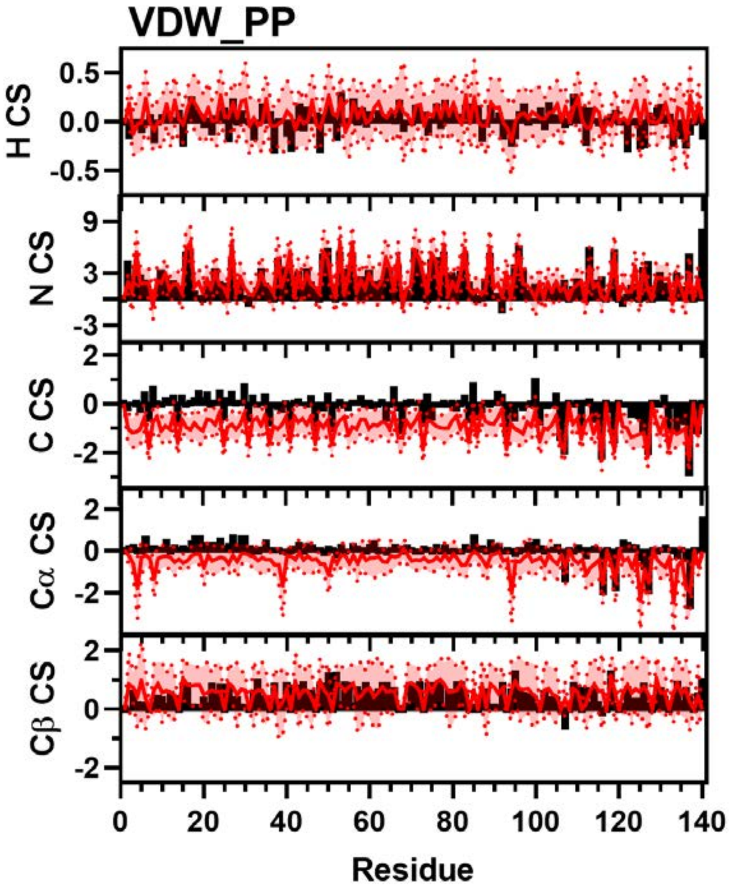


**Figure S51:** Comparison of Simulated and Experimental NMR Chemical Shift Data. Simulated Chemical Shift values from the VDW_PP ensemble (red) overlayed on experimental data (black bars) of N (Top), H (Upper Middle), C (Middle), Cα (Lower Middle), and Cβ (Bottom) chemical shifts from Sung *et al*. [8] Neighbor corrected random coil chemical shift values have been subtracted from both simulated and experimental data [11].

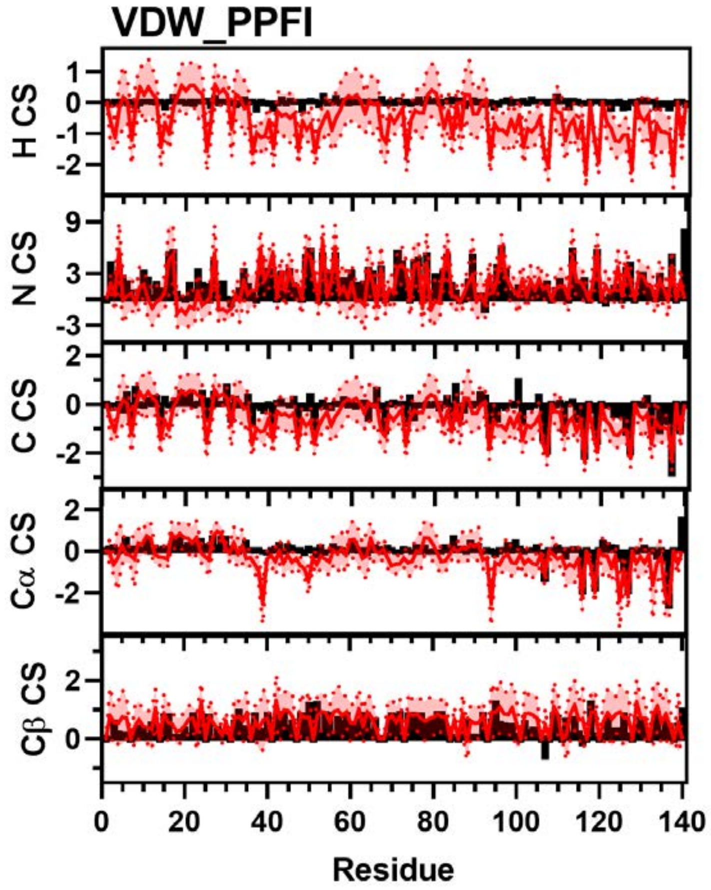


**Figure S52:** Comparison of Simulated and Experimental NMR Chemical Shift Data. Simulated Chemical Shift values from the VDW_PPFI ensemble (red) overlayed on experimental data (black bars) of N (Top), H (Upper Middle), C (Middle), Cα (Lower Middle), and Cβ (Bottom) chemical shifts from Sung *et al*. [8] Neighbor corrected random coil chemical shift values have been subtracted from both simulated and experimental data [11].

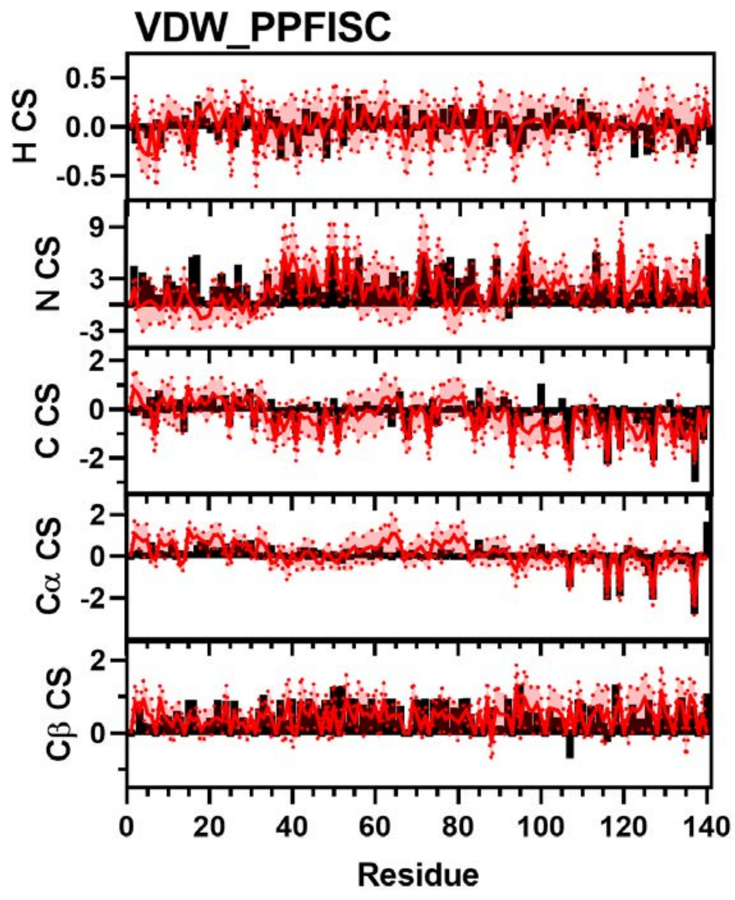


**Figure S53:** Comparison of Simulated and Experimental NMR Chemical Shift Data. Simulated Chemical Shift values from the VDW_PPFISC ensemble (red) overlayed on experimental data (black bars) of N (Top), H (Upper Middle), C (Middle), Cα (Lower Middle), and Cβ (Bottom) chemical shifts from Sung *et al*. [8] Neighbor corrected random coil chemical shift values have been subtracted from both simulated and experimental data [11].

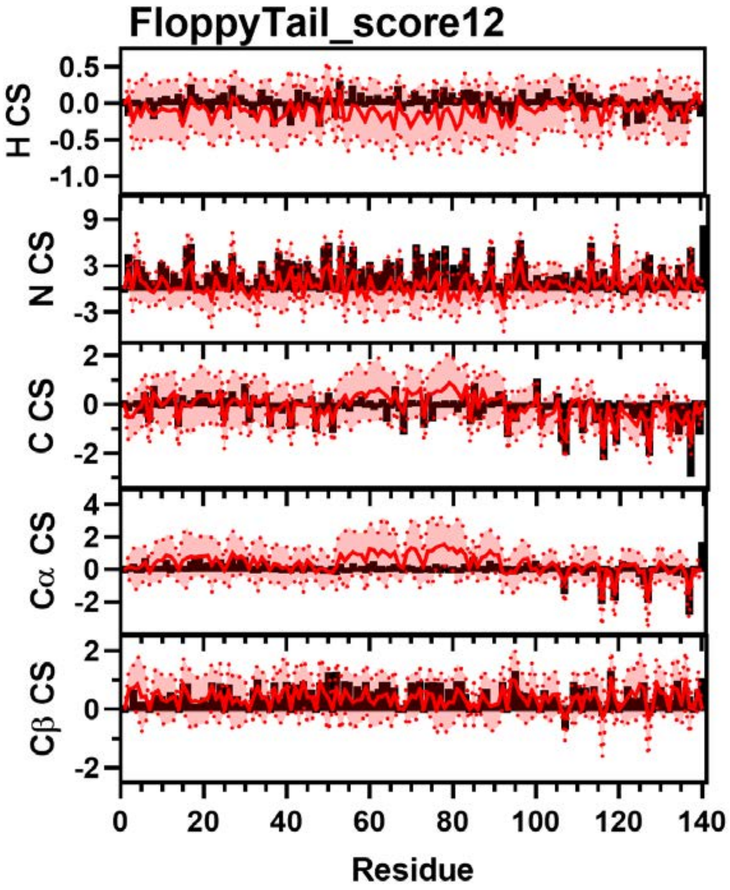


**Figure S54:** Comparison of Simulated and Experimental NMR Chemical Shift Data. Simulated Chemical Shift values from the FloppyTail_score12 ensemble (red) overlayed on experimental data (black bars) of N (Top), H (Upper Middle), C (Middle), Cα (Lower Middle), and Cβ (Bottom) chemical shifts from Sung *et al*. [8] Neighbor corrected random coil chemical shift values have been subtracted from both simulated and experimental data [11].

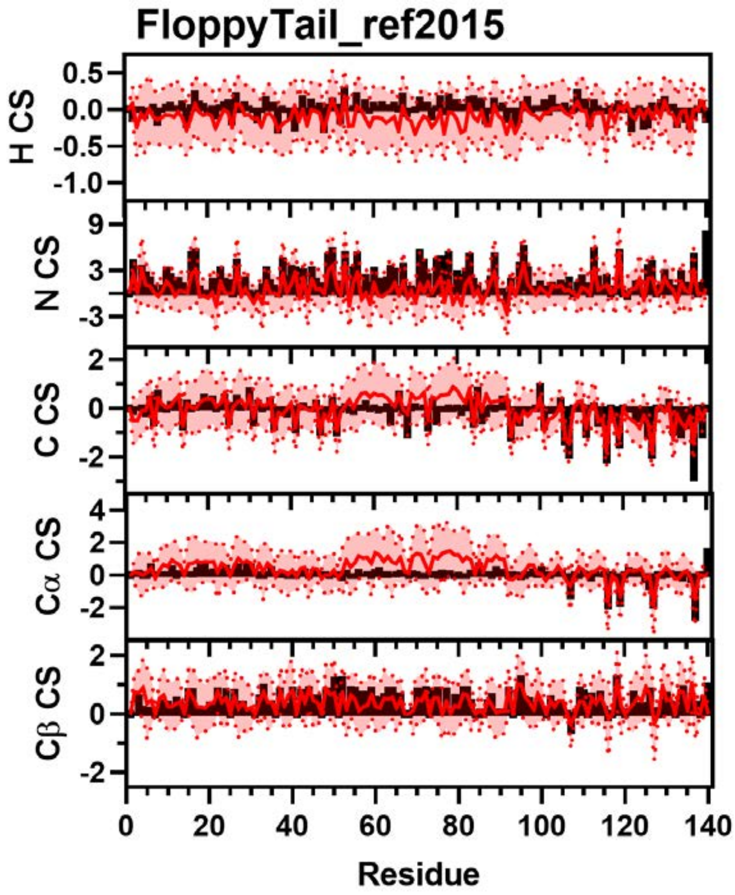


**Figure S55:** Comparison of Simulated and Experimental NMR Chemical Shift Data. Simulated Chemical Shift values from the FloppyTail_ref2015 ensemble (red) overlayed on experimental data (black bars) of N (Top), H (Upper Middle), C (Middle), Cα (Lower Middle), and Cβ (Bottom) chemical shifts from Sung *et al*. [8] Neighbor corrected random coil chemical shift values have been subtracted from both simulated and experimental data [11].

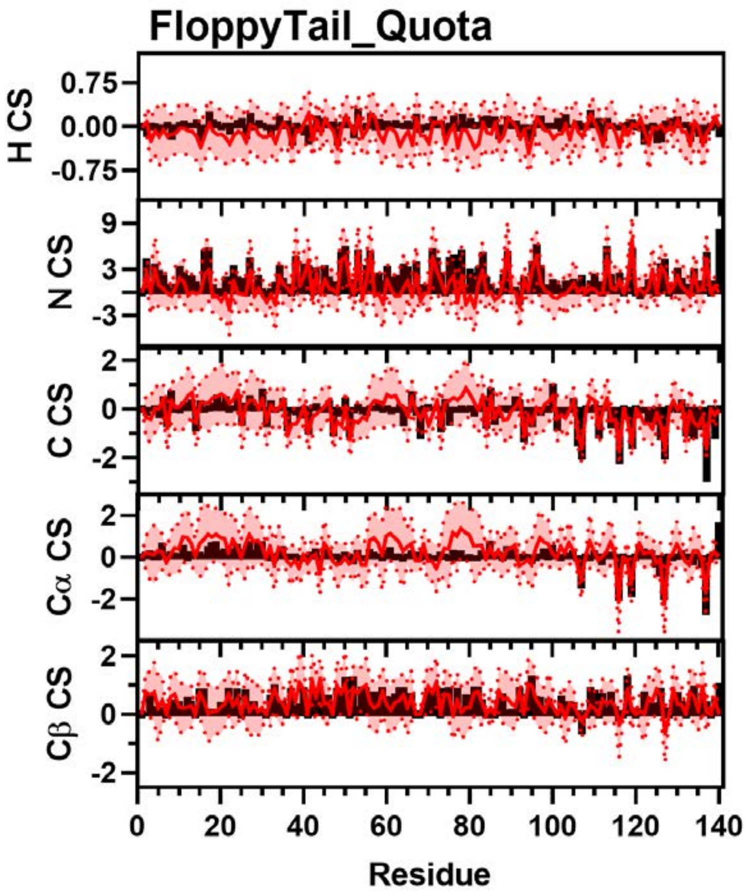


**Figure S56:** Comparison of Simulated and Experimental NMR Chemical Shift Data. Simulated Chemical Shift values from the FloppyTail_Quota ensemble (red) overlayed on experimental data (black bars) of N (Top), H (Upper Middle), C (Middle), Cα (Lower Middle), and Cβ (Bottom) chemical shifts from Sung *et al*. [8] Neighbor corrected random coil chemical shift values have been subtracted from both simulated and experimental data [11].

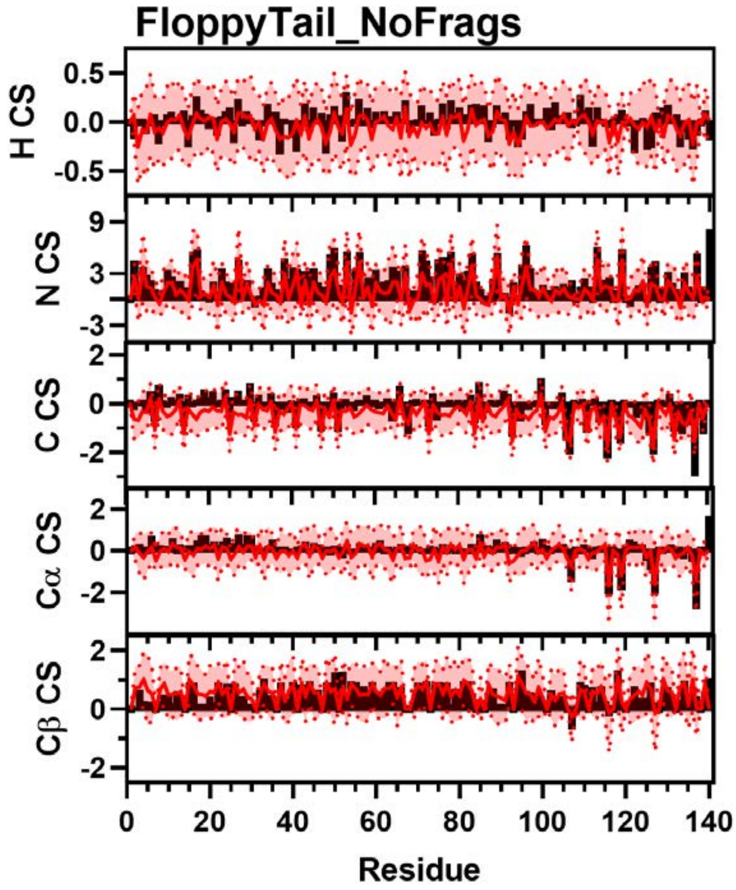


**Figure S57:** Comparison of Simulated and Experimental NMR Chemical Shift Data. Simulated Chemical Shift values from the FloppyTail_NoFrags ensemble (red) overlayed on experimental data (black bars) of N (Top), H (Upper Middle), C (Middle), Cα (Lower Middle), and Cβ (Bottom) chemical shifts from Sung *et al*. [8] Neighbor corrected random coil chemical shift values have been subtracted from both simulated and experimental data [11].

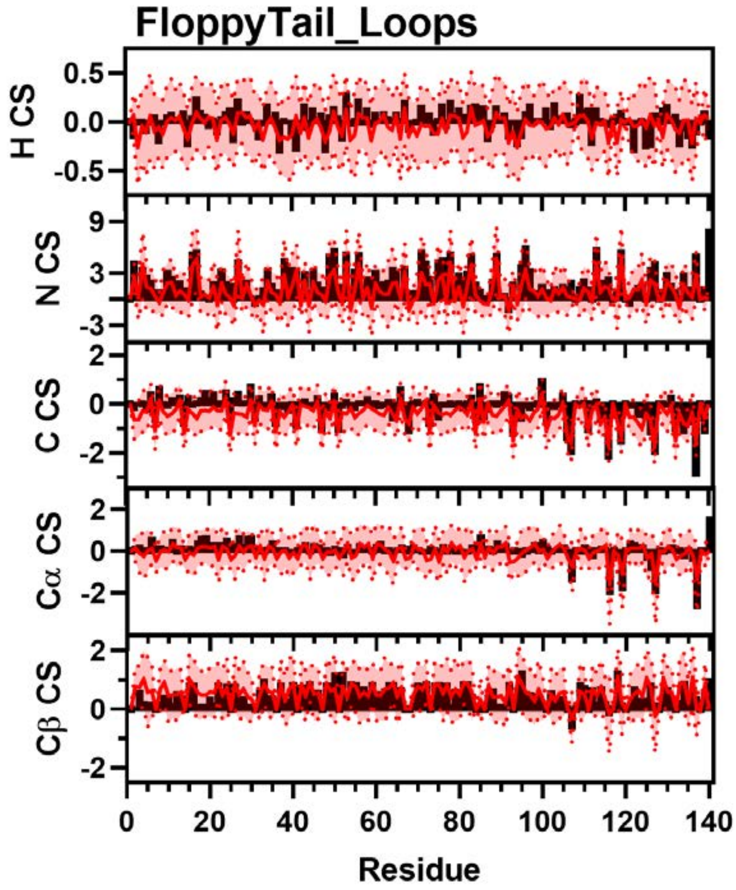


**Figure S58:** Comparison of Simulated and Experimental NMR Chemical Shift Data. Simulated Chemical Shift values from FloppyTail_Loops ensemble (red) overlayed on experimental data (black bars) of N (Top), H (Upper Middle), C (Middle), Cα (Lower Middle), and Cβ (Bottom) chemical shifts from Sung *et al*. [8] Neighbor corrected random coil chemical shift values have been subtracted from both simulated and experimental data [11].

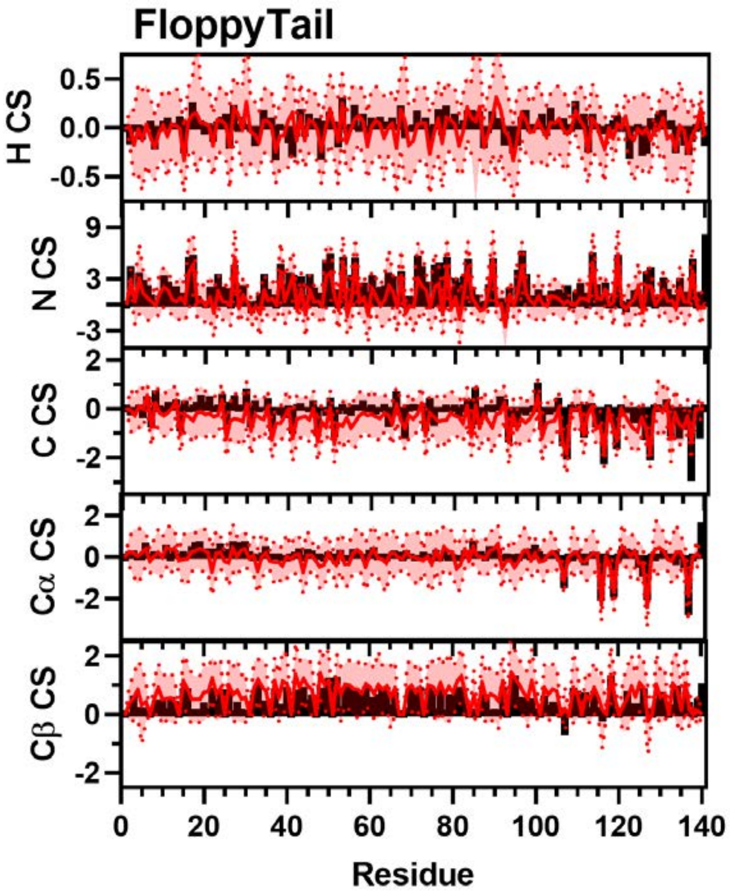


**Figure S59:** Comparison of Simulated and Experimental NMR Chemical Shift Data. Simulated Chemical Shift values from the FloppyTail ensemble (red) overlayed on experimental data (black bars) of N (Top), H (Upper Middle), C (Middle), Cα (Lower Middle), and Cβ (Bottom) chemical shifts from Sung *et al*. [8] Neighbor corrected random coil chemical shift values have been subtracted from both simulated and experimental data [11].

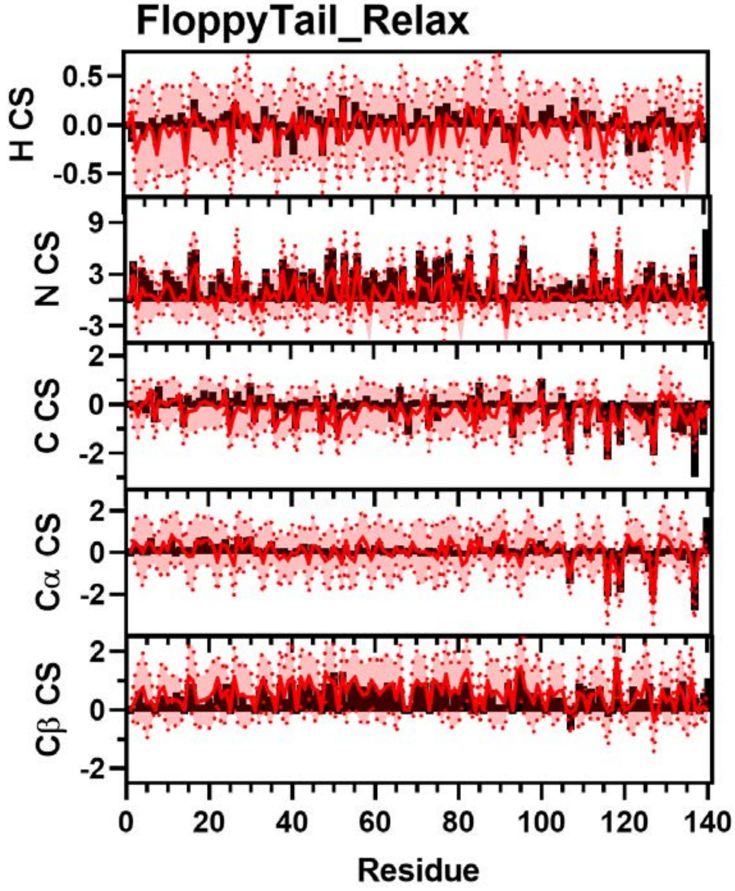


**Figure S60:** Comparison of Simulated and Experimental NMR Chemical Shift Data. Simulated Chemical Shift values from the FloppyTail_Relax ensemble (red) overlayed on experimental data (black bars) of N (Top), H (Upper Middle), C (Middle), Cα (Lower Middle), and Cβ (Bottom) chemical shifts from Sung *et al*. [8] Neighbor corrected random coil chemical shift values have been subtracted from both simulated and experimental data [11].

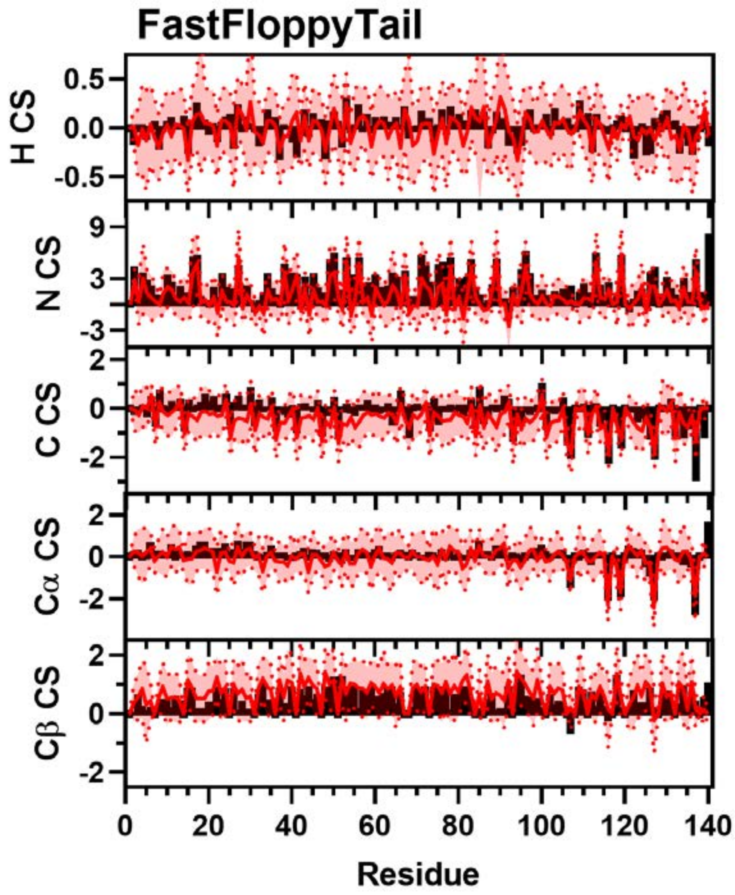


**Figure S61:** Comparison of Simulated and Experimental NMR Chemical Shift Data. Simulated Chemical Shift values from the FastFloppyTail ensemble (red) overlayed on experimental data (black bars) of N (Top), H (Upper Middle), C (Middle), Cα (Lower Middle), and Cβ (Bottom) chemical shifts from Sung *et al*. [8] Neighbor corrected random coil chemical shift values have been subtracted from both simulated and experimental data [11].

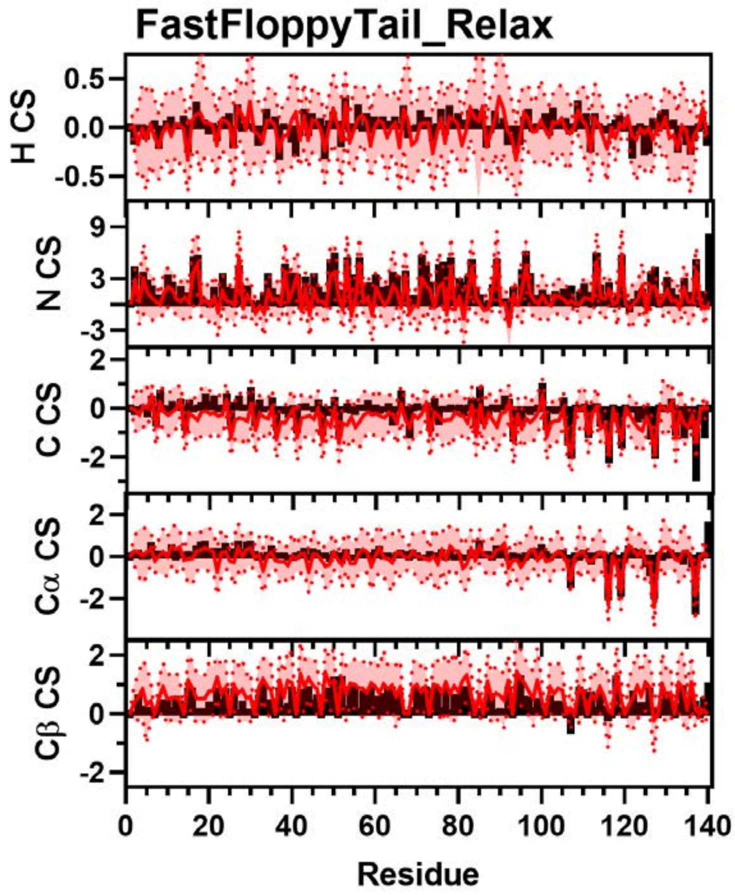


**Figure S62:** Comparison of Simulated and Experimental NMR Chemical Shift Data. Simulated Chemical Shift values from the FastFloppyTail_Relax ensemble (red) overlayed on experimental data (black bars) of N (Top), H (Upper Middle), C (Middle), Cα (Lower Middle), and Cβ (Bottom) chemical shifts from Sung *et al*. [8] Neighbor corrected random coil chemical shift values have been subtracted from both simulated and experimental data [11].

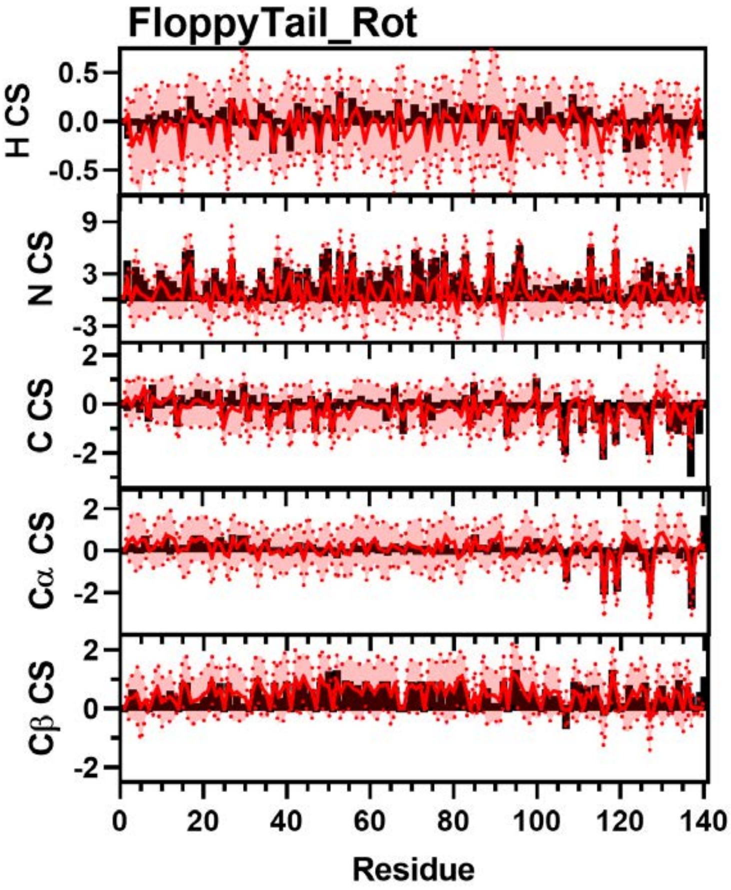


**Figure S63:** Comparison of Simulated and Experimental NMR Chemical Shift Data. Simulated Chemical Shift values from the FloppyTail_Rot ensemble (red) overlayed on experimental data (black bars) of N (Top), H (Upper Middle), C (Middle), Cα (Lower Middle), and Cβ (Bottom) chemical shifts from Sung *et al*. [8] Neighbor corrected random coil chemical shift values have been subtracted from both simulated and experimental data [11].

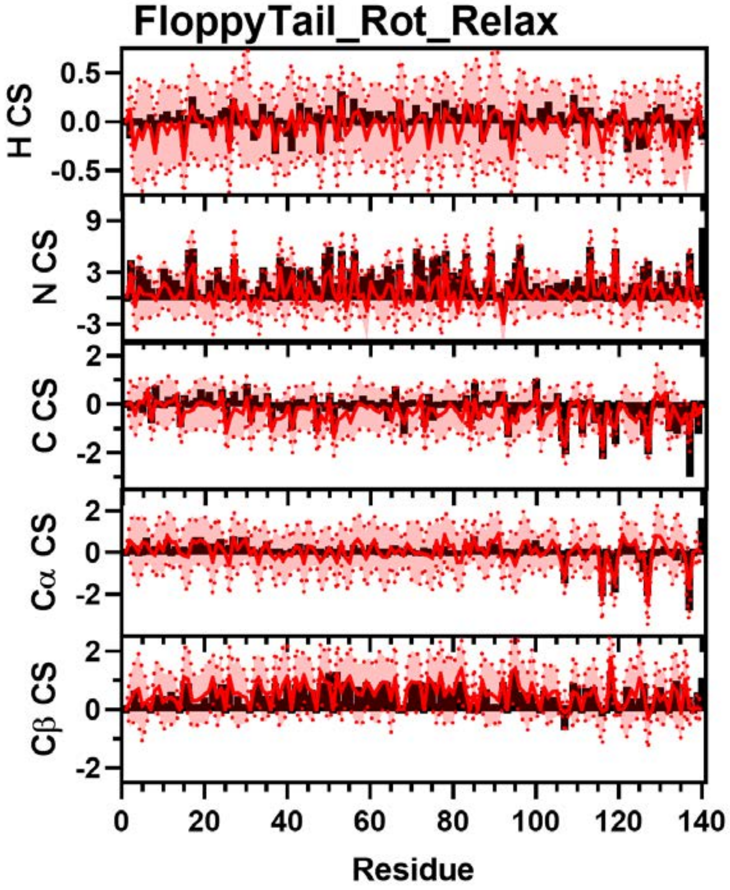


**Figure S64:** Comparison of Simulated and Experimental NMR Chemical Shift Data. Simulated Chemical Shift values from the FloppyTail_Rot_Relax ensemble (red) overlayed on experimental data (black bars) of N (Top), H (Upper Middle), C (Middle), Cα (Lower Middle), and Cβ (Bottom) chemical shifts from Sung *et al*. [8] Neighbor corrected random coil chemical shift values have been subtracted from both simulated and experimental data [11].

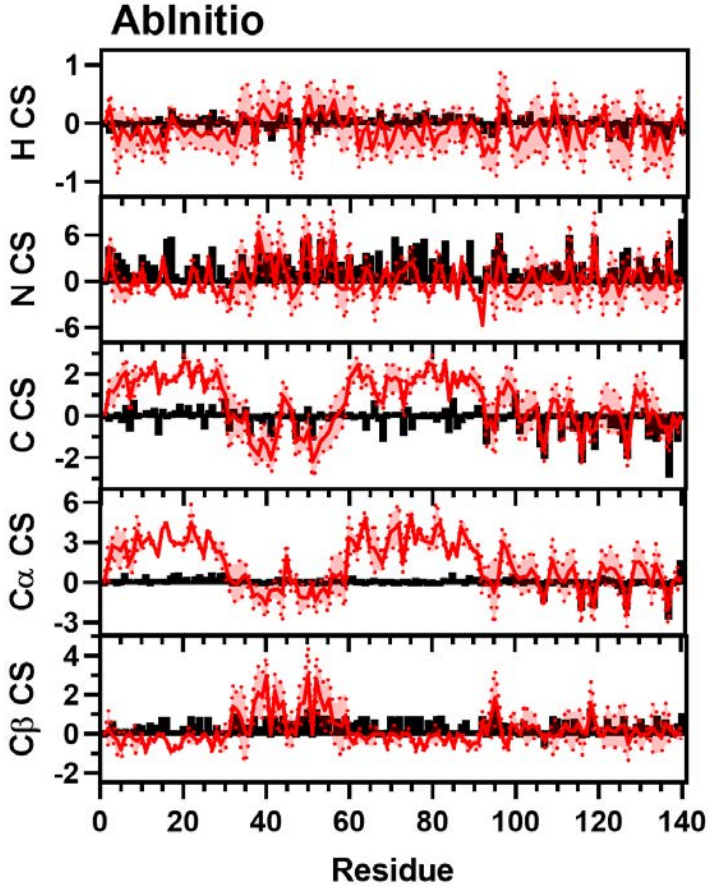


**Figure S65:** Comparison of Simulated and Experimental NMR Chemical Shift Data. Simulated Chemical Shift values from the AbInitio ensemble (red) overlayed on experimental data (black bars) of N (Top), H (Upper Middle), C (Middle), Cα (Lower Middle), and Cβ (Bottom) chemical shifts from Sung *et al*. [8] Neighbor corrected random coil chemical shift values have been subtracted from both simulated and experimental data [11].

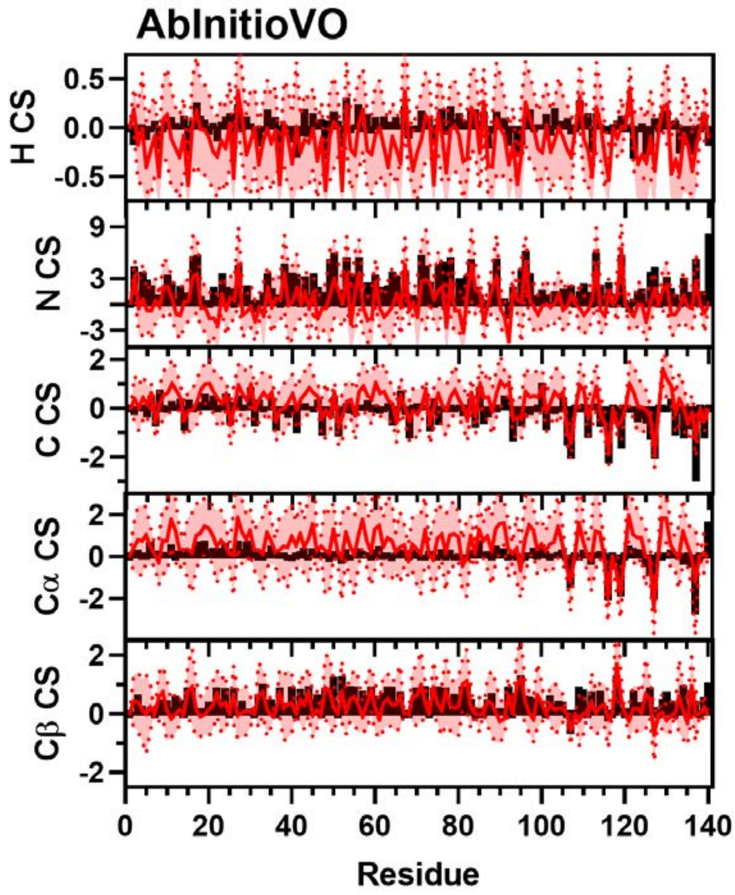


**Figure S66:** Comparison of Simulated and Experimental NMR Chemical Shift Data. Simulated Chemical Shift values from the AbInitioVO ensemble (red) overlayed on experimental data (black bars) of N (Top), H (Upper Middle), C (Middle), Cα (Lower Middle), and Cβ (Bottom) chemical shifts from Sung *et al*. [8] Neighbor corrected random coil chemical shift values have been subtracted from both simulated and experimental data [11].

**Comparison with αS Residual Dipolar Coupling Data**

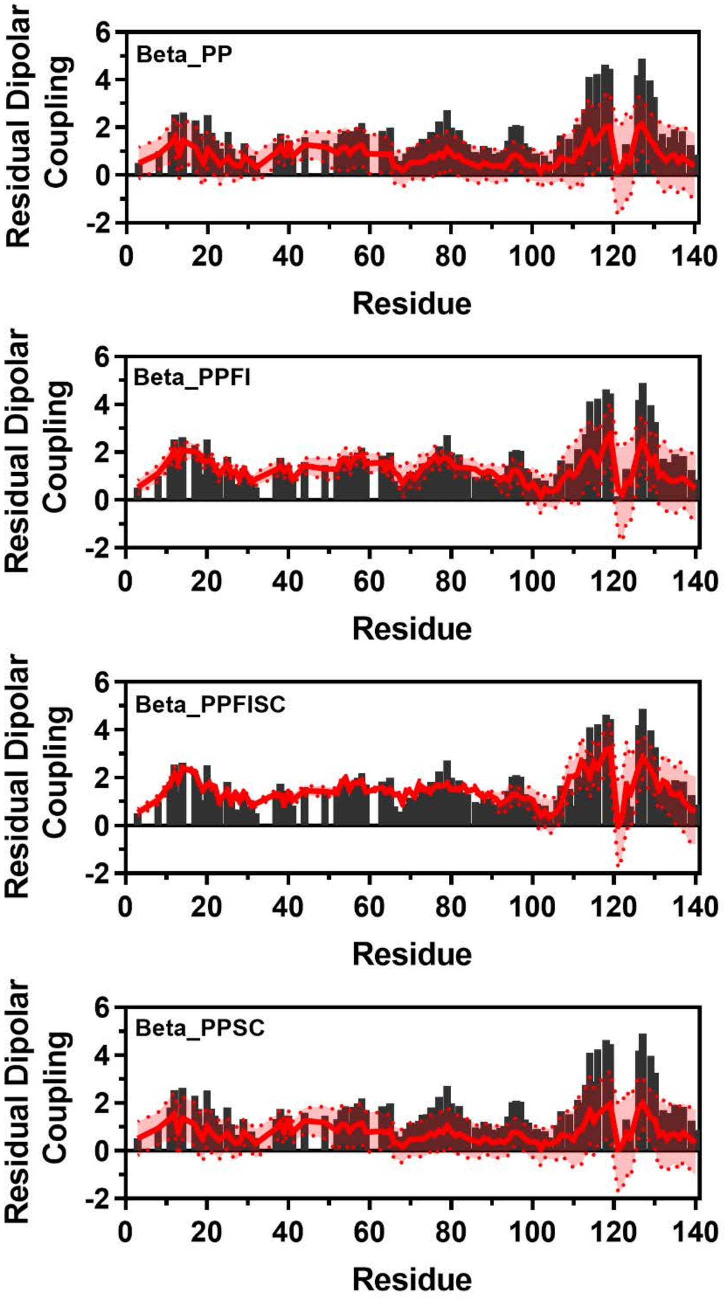


**Figure S67:** Comparison of Simulated and Experimental Residual Dipolar Coupling Data. Simulated RDC values (red line) from Beta_PP (Top), Beta_PPFI (Upper Middle), Beta_PPFISC (Lower Middle), and Beta_PPSC (Bottom) overlayed on experimental data (black bars) from Bertoncini *et al*. [25].

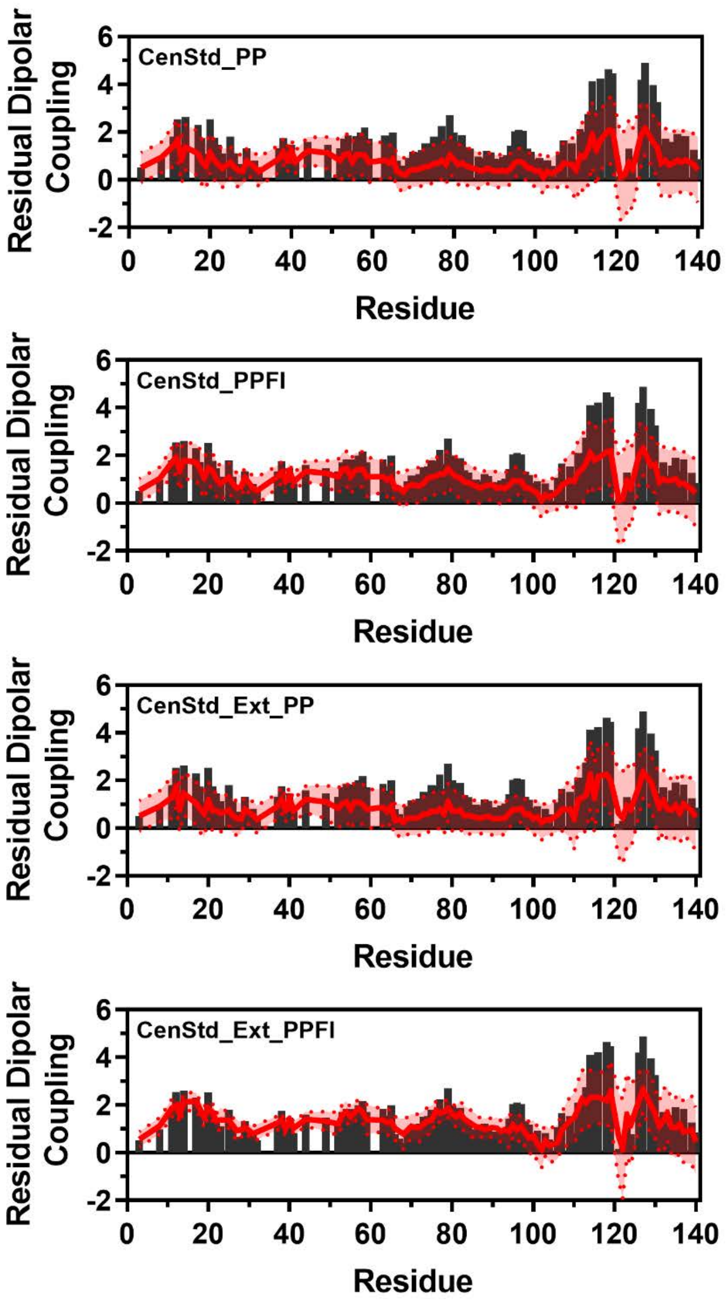


**Figure S68:** Comparison of Simulated and Experimental Residual Dipolar Coupling Data. Simulated RDC values (red line) from CenStd_PP (Top), CenStd_PPFI (Upper Middle), CenStd_Ext_PP (Lower Middle), and CenStd_Ext_PPFI (Bottom) overlayed on experimental data (black bars) from Bertoncini *et al*. [25].

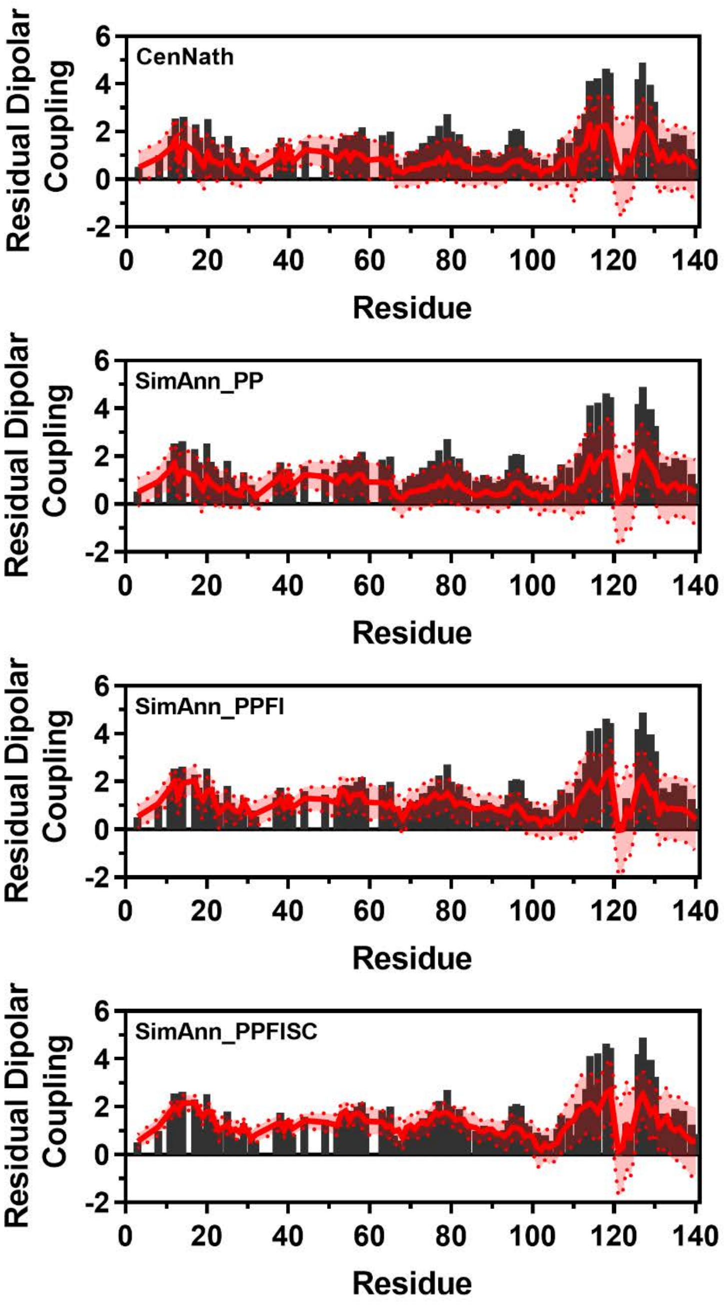


**Figure S69:** Comparison of Simulated and Experimental Residual Dipolar Coupling Data. Simulated RDC values (red line) from SimAnn_PP (Top), SimAnn_PPFI (Upper Middle), SimAnn_PPFISC (Lower Middle), and SimAnn_PPSC (Bottom) overlayed on experimental data (black bars) from Bertoncini *et al*. [25].

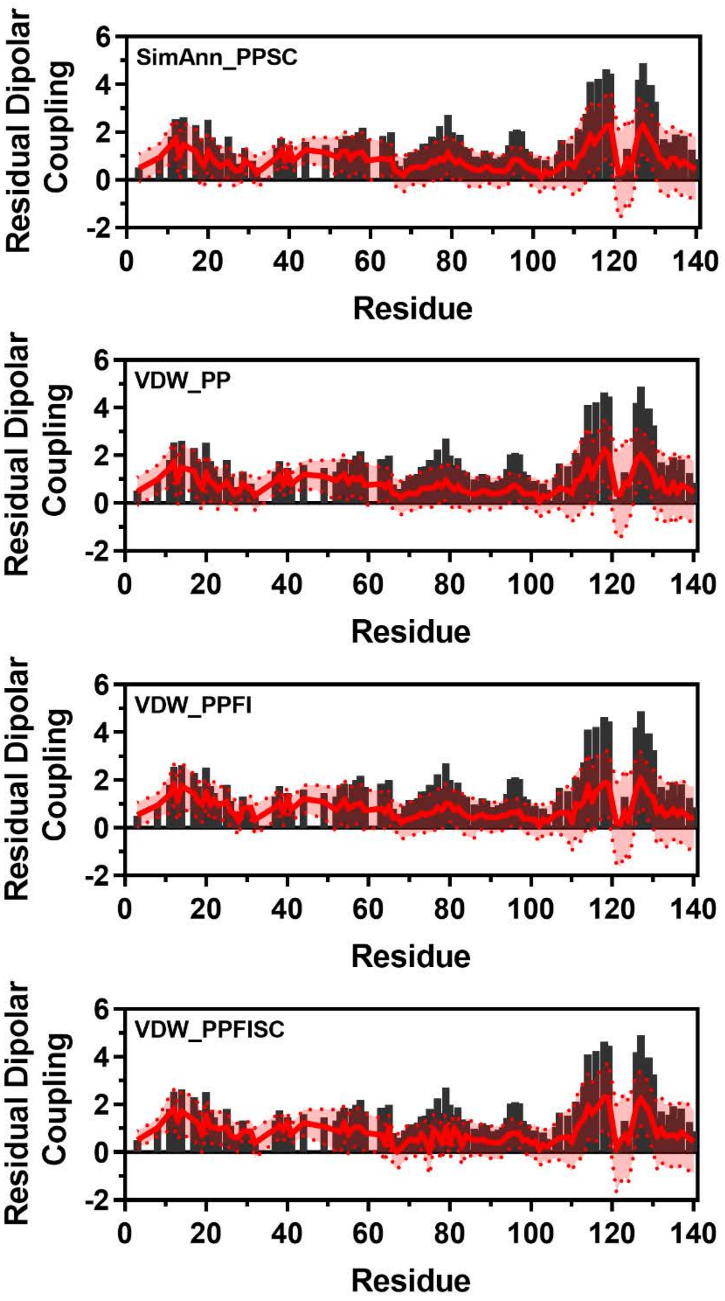


**Figure S70:** Comparison of Simulated and Experimental Residual Dipolar Coupling Data. Simulated RDC values (red line) from CenNath (Top), VDW_PP (Upper Middle), VDW_PPFI (Lower Middle), and VDW_PPFISC (Bottom) overlayed on experimental data (black bars) from Bertoncini *et al*. [25].

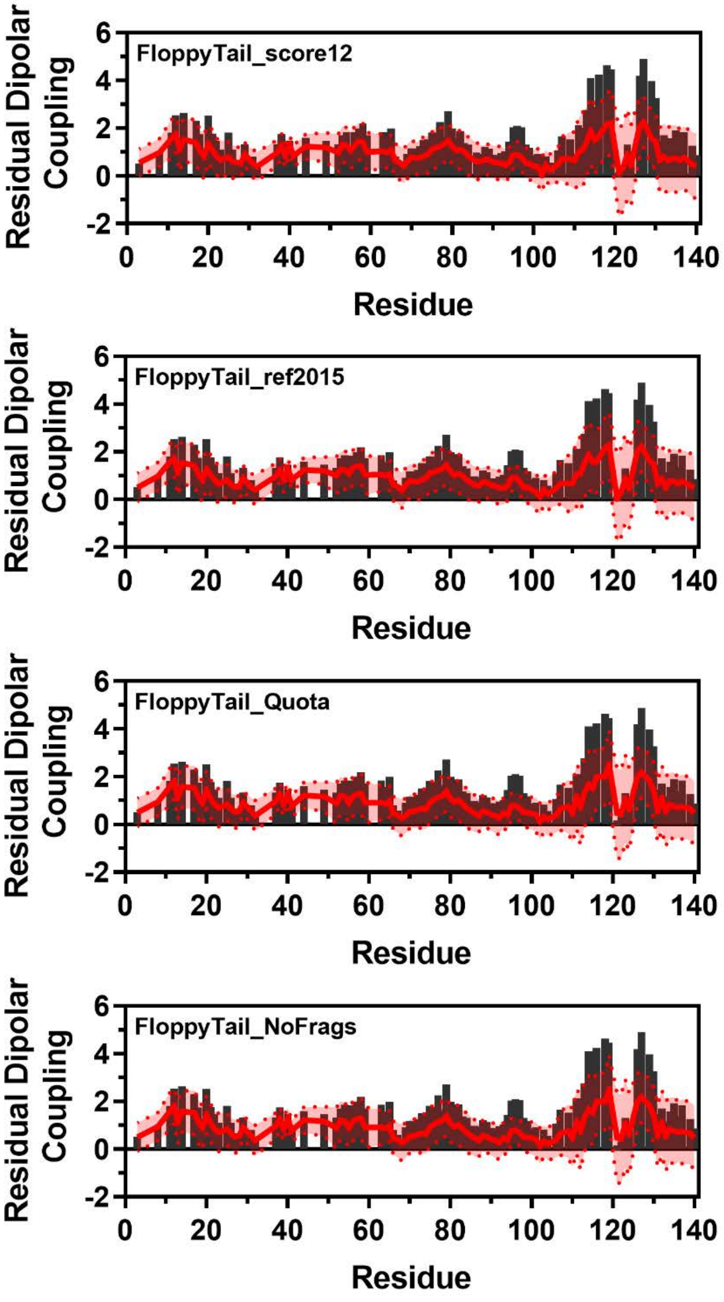


**Figure S71:** Comparison of Simulated and Experimental Residual Dipolar Coupling Data. Simulated RDC values (red line) from FloppyTail_score12 (Top), FloppyTail_ref2015 (Upper Middle), FloppyTail_Quota (Lower Middle), and FloppyTail_NoFrags (Bottom) overlayed on experimental data (black bars) from Bertoncini *et al*. [25].

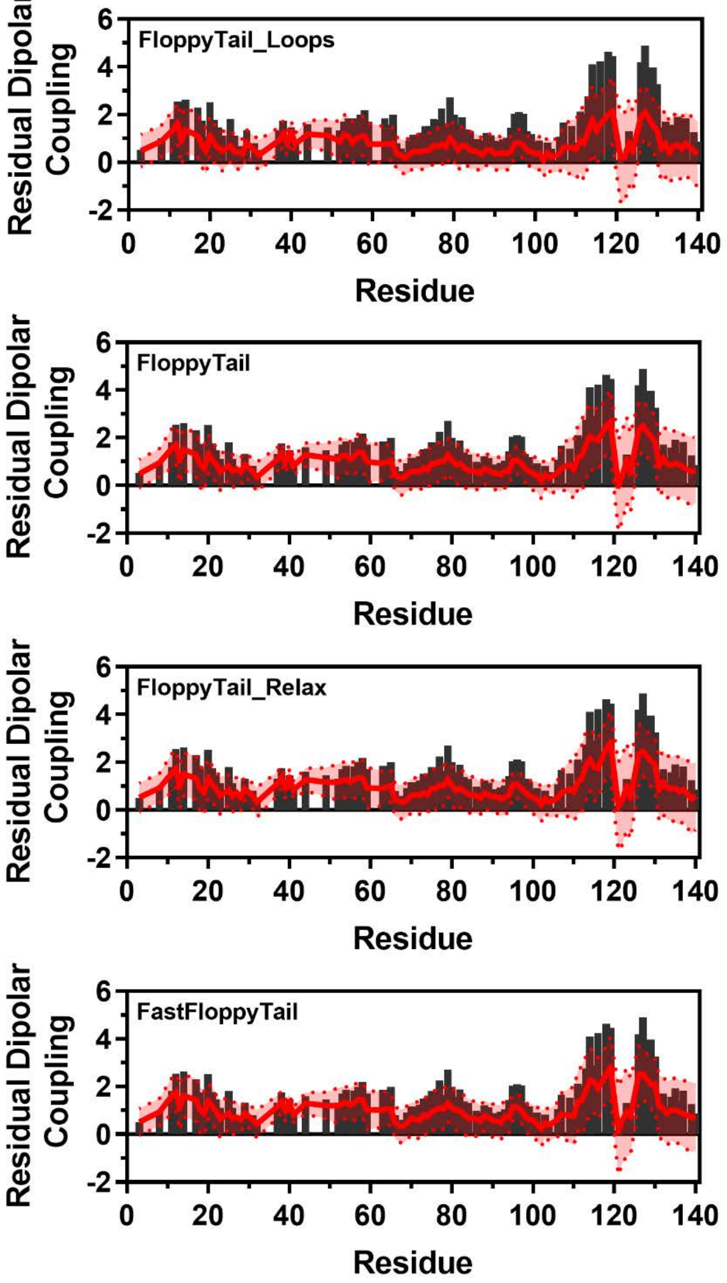


**Figure S72:** Comparison of Simulated and Experimental Residual Dipolar Coupling Data. Simulated RDC values (red line) from FloppyTail_Loops (Top), FloppyTail (Upper Middle), FloppyTail_Relax (Lower Middle), and FastFloppyTail (Bottom) overlayed on experimental data (black bars) from Bertoncini *et al*. [25].

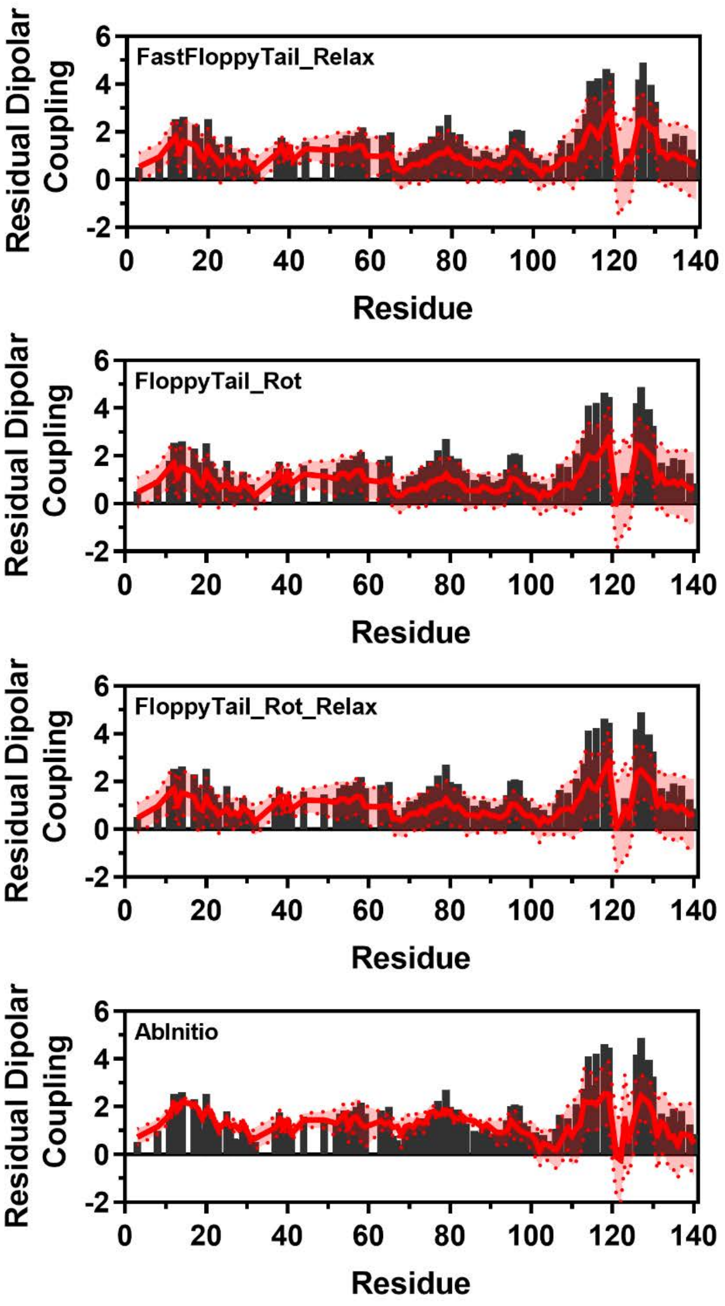


**Figure S73:** Comparison of Simulated and Experimental Residual Dipolar Coupling Data. Simulated RDC values (red line) from FastFloppyTail_Relax (Top), FloppyTail_Rot (Upper Middle), FloppyTail_Rot_Relax (Lower Middle), and AbInitio (Bottom) overlayed on experimental data (black bars) from Bertoncini *et al*. [25].

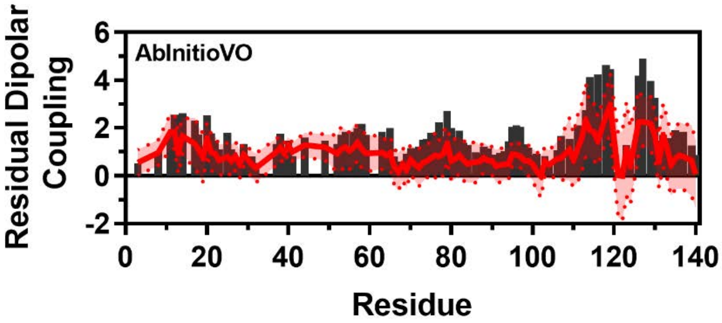


**Figure S74:** Comparison of Simulated and Experimental Residual Dipolar Coupling Data. Simulated RDC values (red line) from AbInitioVO (Top) overlayed on experimental data (black bars) from Bertoncini *et al*. [25].

### Comparison with αS J-Coupling Data

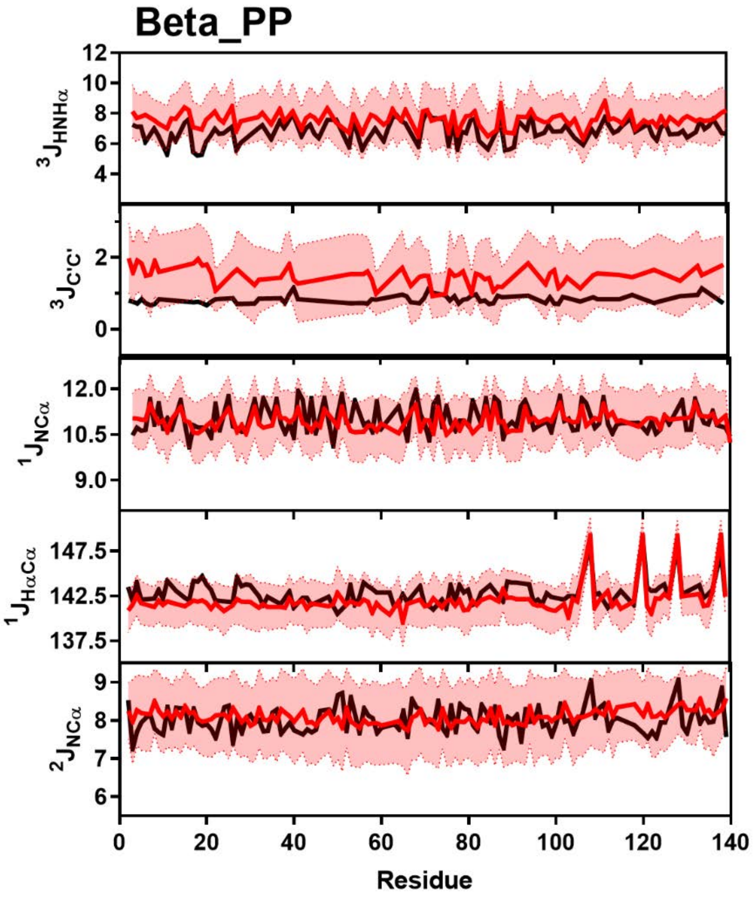


**Figure S75:** Comparison of Simulated and Experimental J-Coupling Data. J-Coupling values from the Beta_PP ensemble (red) overlayed on experimental data (black) of $^3J_{HNH\alpha}$ (Top), $^3J_{CC}$ (Upper Middle), $^1J_{H\alpha C\alpha}$ (Middle), $^1J_{NC\alpha}$ (Lower Middle) and $^2J_{NC\alpha}$ (Bottom) from Mantsyzov *et al.* and Lee *et al*. [13,18].

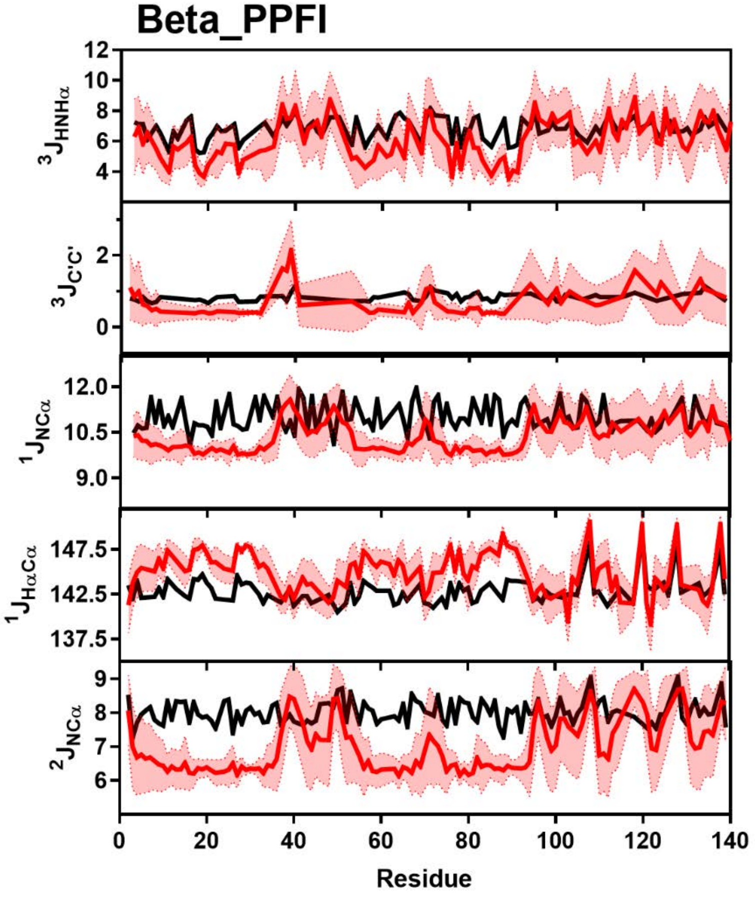


**Figure S76:** Comparison of Simulated and Experimental J-Coupling Data. Simulated J-Coupling values from the Beta_PPFI ensemble (red) overlayed on experimental data (black) of $^{3}J_{HNH\alpha}$ (Top), $^{3}J_{CC}$ (Upper Middle), $^{1}J_{H\alpha C\alpha}$ (Middle), $^{1}J_{NC\alpha}$ (Lower Middle) and $^{2}J_{NC\alpha}$ (Bottom) from Mantsyzov *et al.* and Lee *et al*. [13,18].

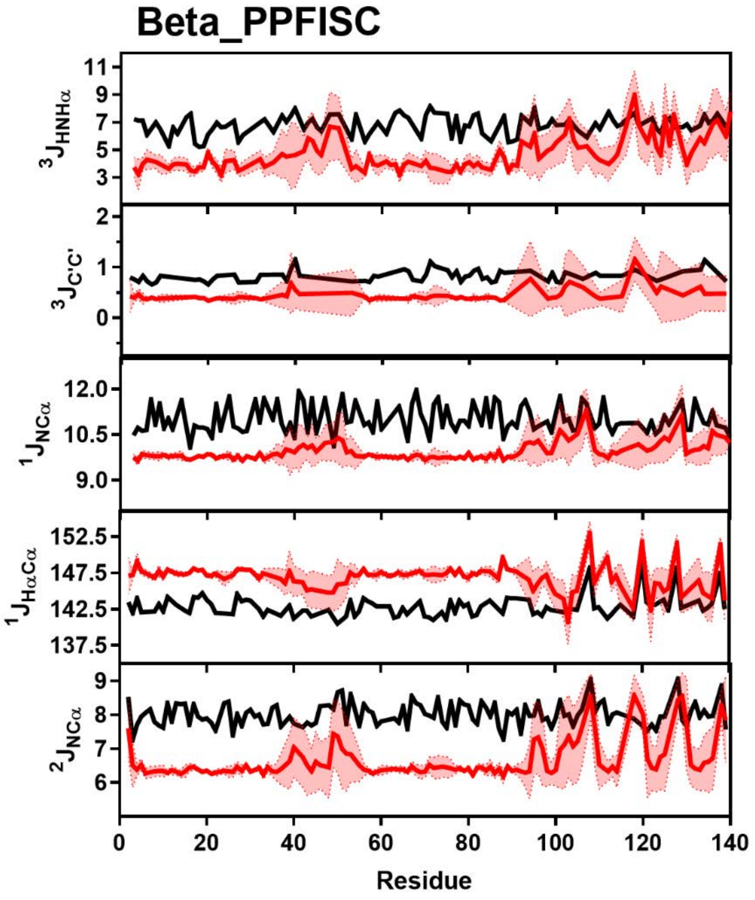


**Figure S77:** Comparison of Simulated and Experimental J-Coupling Data. Simulated J-Coupling values from the Beta_PPFISC ensemble (red) overlayed on experimental data (black) of $^3J_{HNH\alpha}$ (Top), $^3J_{CC}$ (Upper Middle), $^1J_{H\alpha C\alpha}$ (Middle), $^1J_{NC\alpha}$ (Lower Middle) and $^2J_{NC\alpha}$ (Bottom) from Mantsyzov *et al.* and Lee *et al*. [13,18].

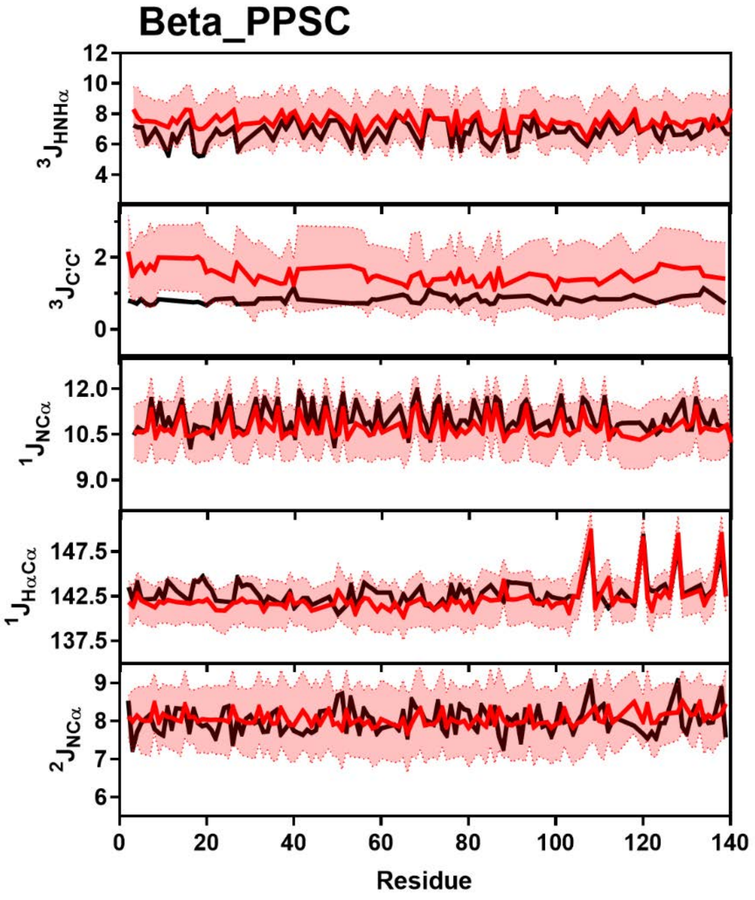


**Figure S78:** Comparison of Simulated and Experimental J-Coupling Data. Simulated J-Coupling values from the Beta_PPSC ensemble (red) overlayed on experimental data (black) of $^{3}J_{HNH\alpha}$ (Top), $^{3}J_{CC}$ (Upper Middle), $^{1}J_{H\alpha C\alpha}$ (Middle), $^{1}J_{NC\alpha}$ (Lower Middle) and $^{2}J_{NC\alpha}$ (Bottom) from Mantsyzov *et al.* and Lee *et al*. [13,18].

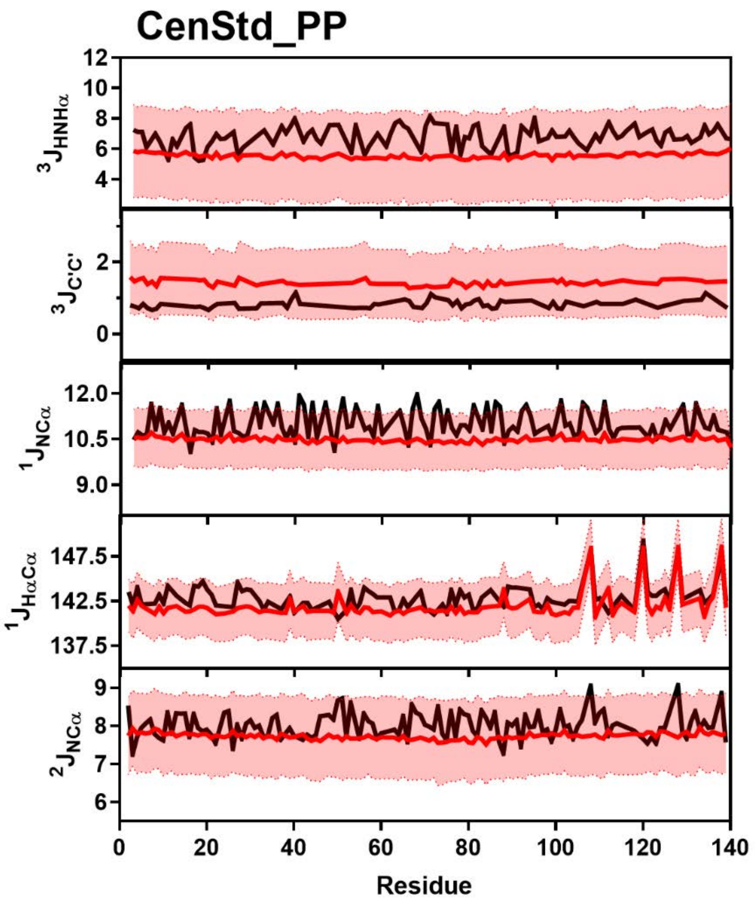


**Figure S79:** Comparison of Simulated and Experimental J-Coupling Data. Simulated J-Coupling values from the CenStd_PP ensemble (red) overlayed on experimental data (black) of $^3J_{HNH\alpha}$ (Top), $^3J_{CC}$ (Upper Middle), $^1J_{H\alpha C\alpha}$ (Middle), $^1J_{NC\alpha}$ (Lower Middle) and $^2J_{NC\alpha}$ (Bottom) from Mantsyzov *et al.* and Lee *et al*. [13,18].

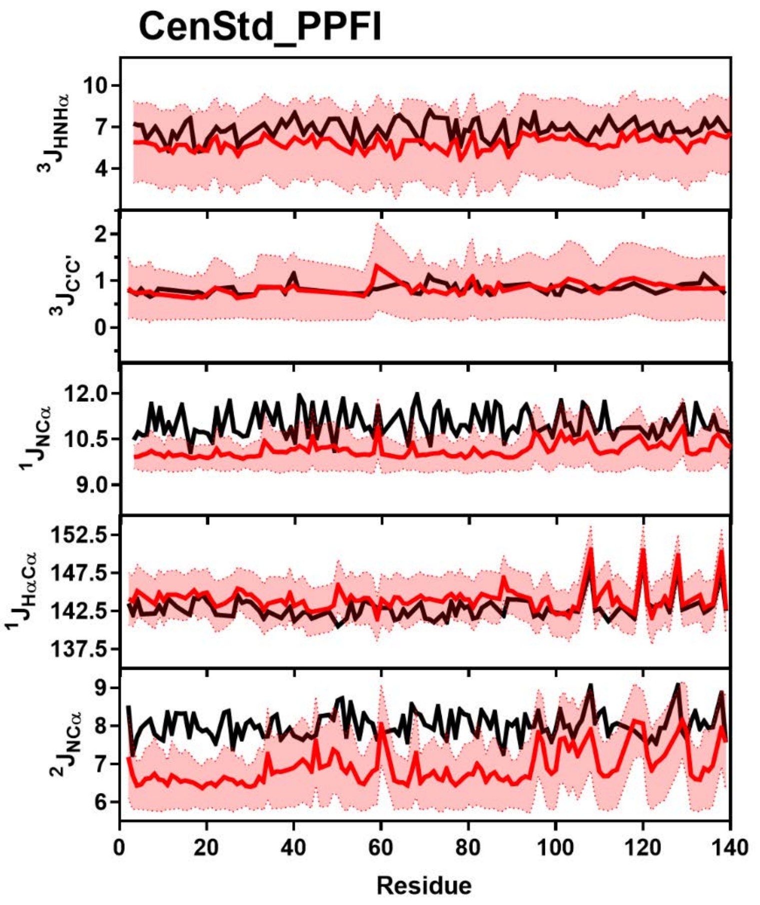


**Figure S80:** Comparison of Simulated and Experimental J-Coupling Data. Simulated J-Coupling values from the CenStd_PPFI ensemble (red) overlayed on experimental data (black) of $^3J_{HNH\alpha}$ (Top), $^3J_{CC}$ (Upper Middle), $^1J_{H\alpha C\alpha}$ (Middle), $^1J_{NC\alpha}$ (Lower Middle) and $^2J_{NC\alpha}$ (Bottom) from Mantsyzov *et al.* and Lee *et al*. [13,18].

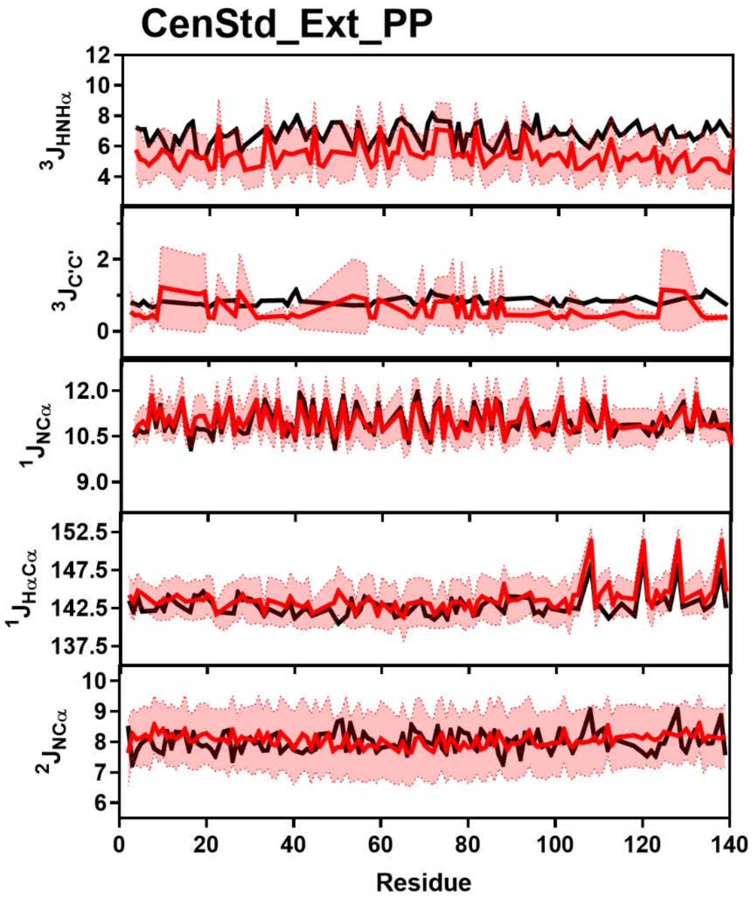


**Figure S81:** Comparison of Simulated and Experimental J-Coupling Data. Simulated J-Coupling values from the CenStd_Ext_PP ensemble (red) overlayed on experimental data (black) of $^3J_{HNH\alpha}$ (Top), $^3J_{CC}$ (Upper Middle), $^1J_{H\alpha C\alpha}$ (Middle), $^1J_{NC\alpha}$ (Lower Middle) and $^2J_{NC\alpha}$ (Bottom) from Mantsyzov *et al.* and Lee *et al*. [13,18].

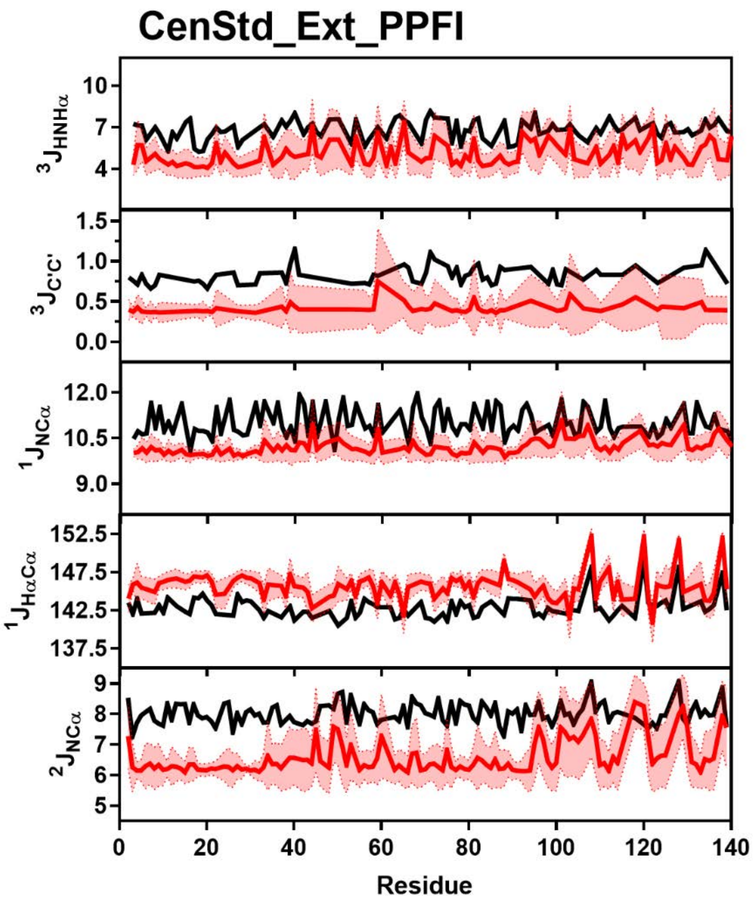


**Figure S82:** Comparison of Simulated and Experimental J-Coupling Data. Simulated J-Coupling values from the CenStd_Ext_PPFI ensemble (red) overlayed on experimental data (black) of $^3J_{HNH\alpha}$ (Top), $^3J_{CC}$ (Upper Middle), $^1J_{H\alpha C\alpha}$ (Middle), $^1J_{NC\alpha}$ (Lower Middle) and $^2J_{NC\alpha}$ (Bottom) from Mantsyzov *et al.* and Lee *et al*. [13,18].

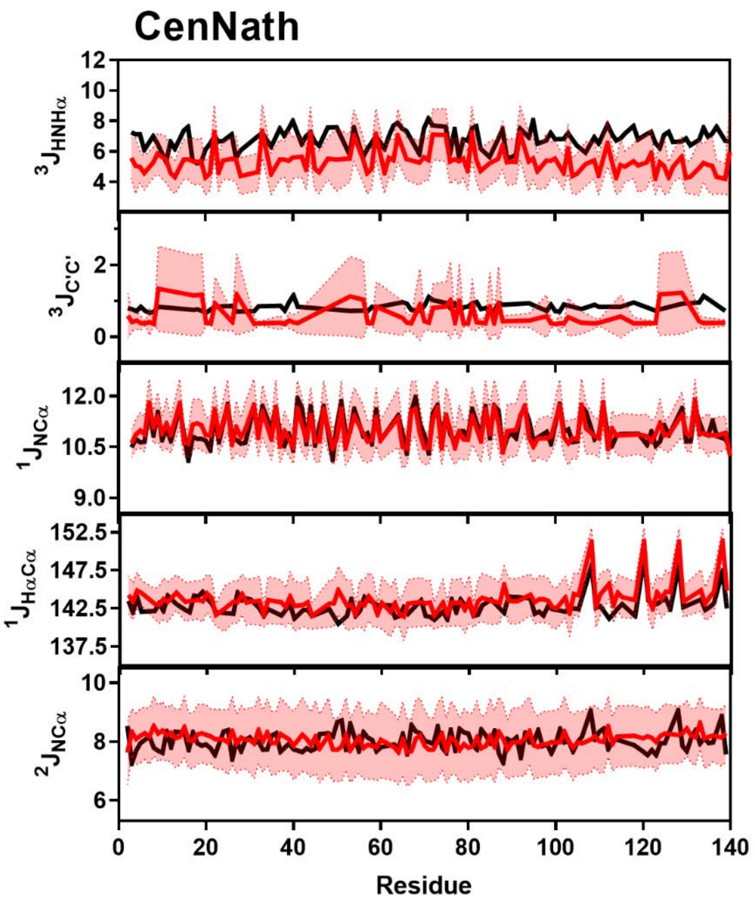


**Figure S83:** Comparison of Simulated and Experimental J-Coupling Data. Simulated J-Coupling values from the CenNath ensemble (red) overlayed on experimental data (black) of $^3J_{HNH\alpha}$ (Top), $^3J_{CC}$ (Upper Middle), $^1J_{H\alpha C\alpha}$ (Middle), $^1J_{NC\alpha}$ (Lower Middle) and $^2J_{NC\alpha}$ (Bottom) from Mantsyzov *et al.* and Lee *et al*. [13,18].

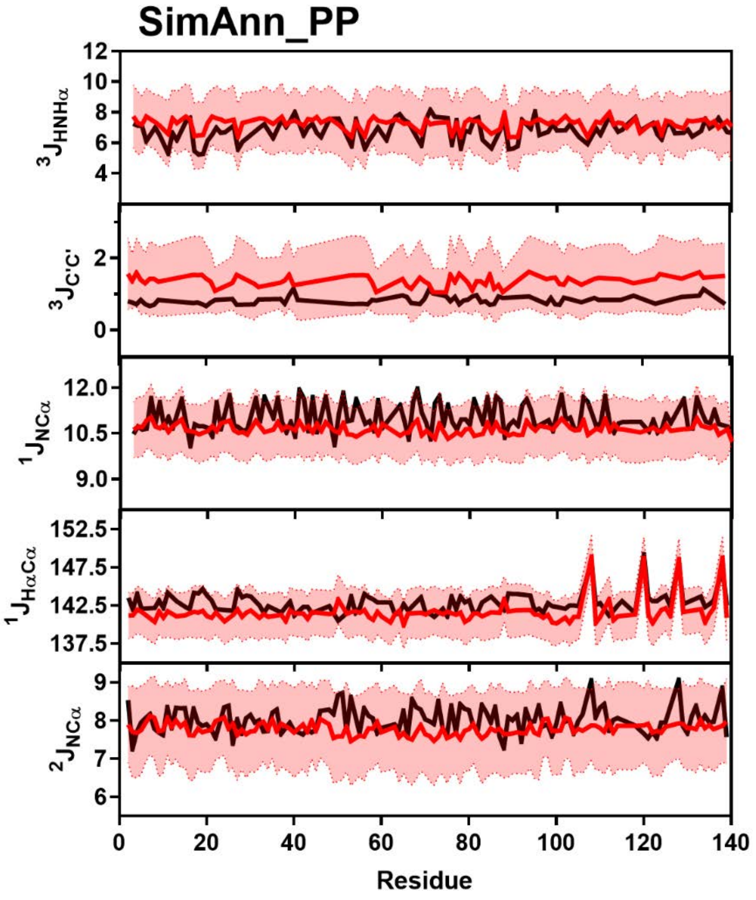


**Figure S84:** Comparison of Simulated and Experimental J-Coupling Data. Simulated J-Coupling values from the SimAnn_PP ensemble (red) overlayed on experimental data (black) of $^{3}J_{HNH\alpha}$ (Top), $^{3}J_{CC}$ (Upper Middle), $^{1}J_{H\alpha C\alpha}$ (Middle), $^{1}J_{NC\alpha}$ (Lower Middle) and $^{2}J_{NC\alpha}$ (Bottom) from Mantsyzov *et al.* and Lee *et al*. [13,18].

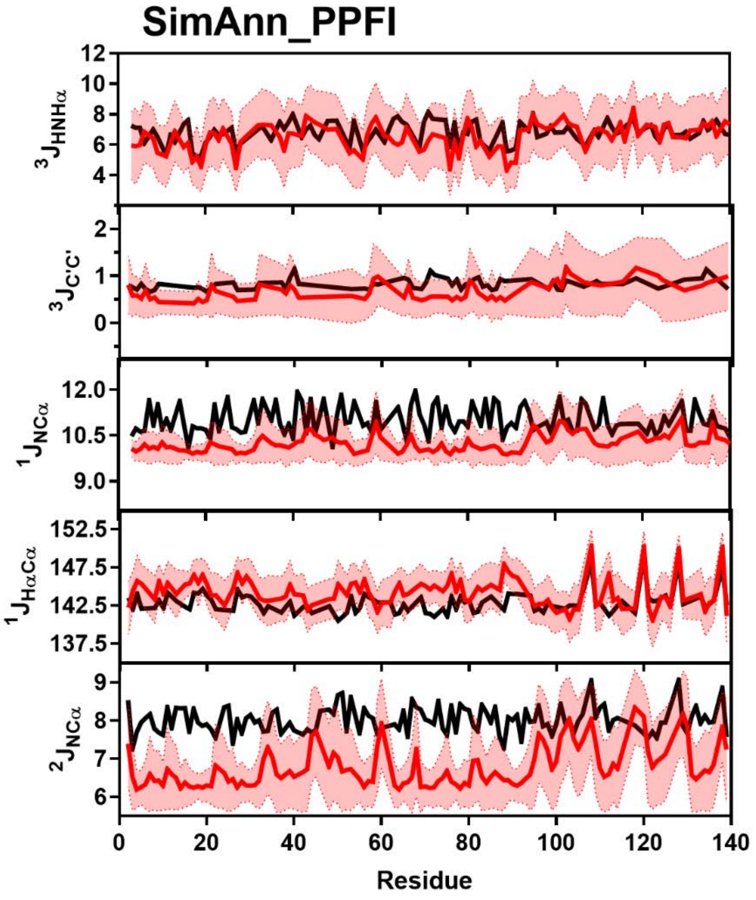


**Figure S85:** Comparison of Simulated and Experimental J-Coupling Data. Simulated J-Coupling values from the SimAnn_PPFI ensemble (red) overlayed on experimental data (black) of $^3J_{HNH\alpha}$ (Top), $^3J_{CC}$ (Upper Middle), $^1J_{H\alpha C\alpha}$ (Middle), $^1J_{NC\alpha}$ (Lower Middle) and $^2J_{NC\alpha}$ (Bottom) from Mantsyzov *et al.* and Lee *et al*. [13,18].

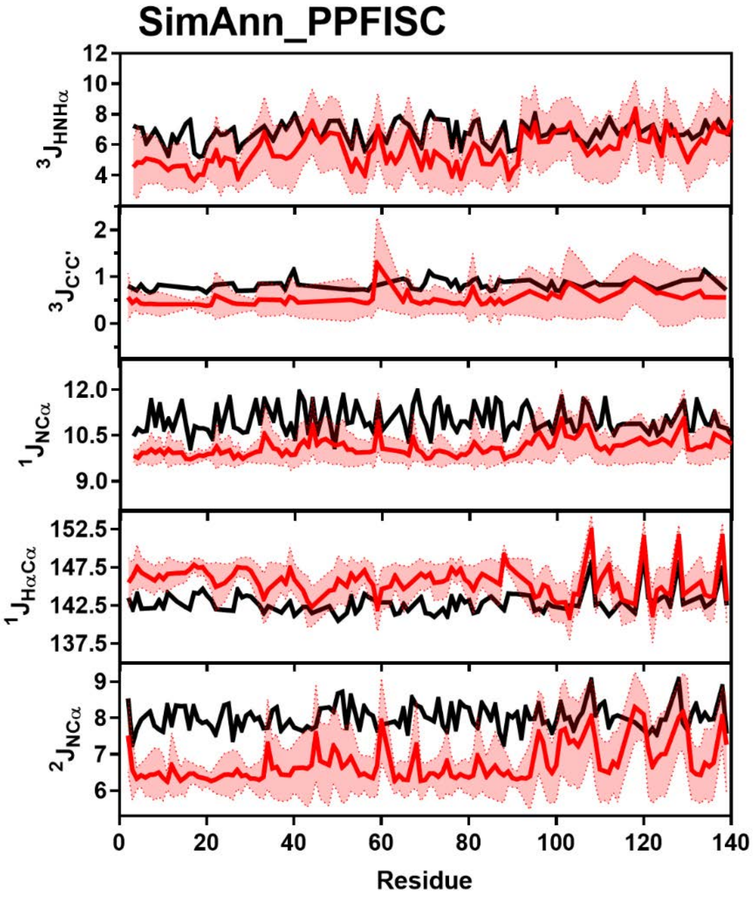


**Figure S86:** Comparison of Simulated and Experimental J-Coupling Data. Simulated J-Coupling values from the SimAnn_PPFISC ensemble (red) overlayed on experimental data (black) of $^3J_{HNH\alpha}$ (Top), $^3J_{CC}$ (Upper Middle), $^1J_{H\alpha C\alpha}$ (Middle), $^1J_{NC\alpha}$ (Lower Middle) and $^2J_{NC\alpha}$ (Bottom) from Mantsyzov *et al.* and Lee *et al*. [13,18].

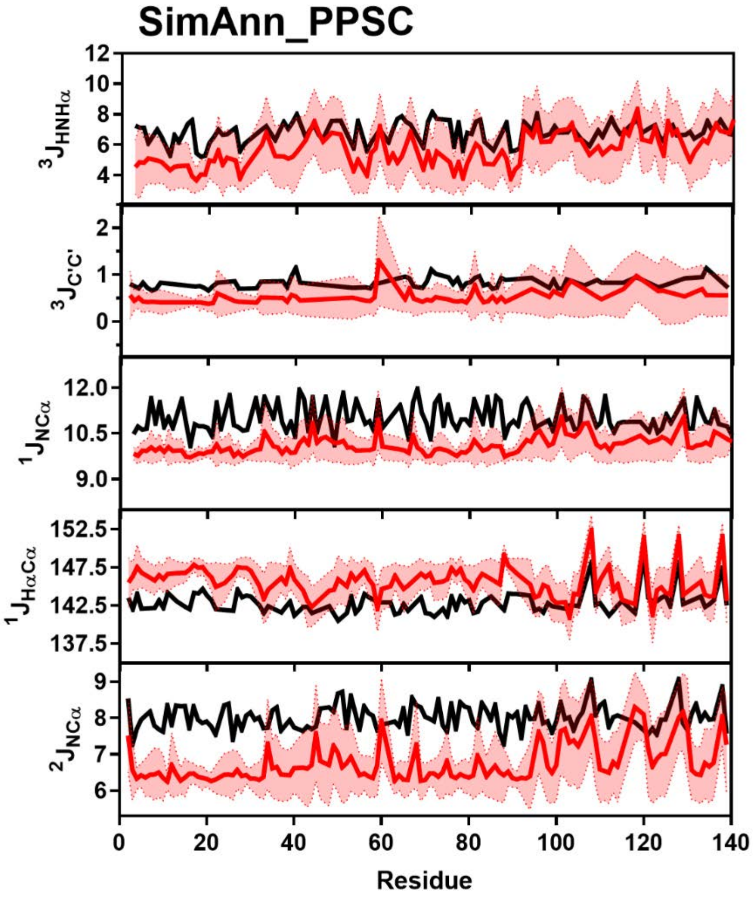


**Figure S87:** Comparison of Simulated and Experimental J-Coupling Data. Simulated J-Coupling values from the SimAnn_PPSC ensemble (red) overlayed on experimental data (black) of $^3J_{HNH\alpha}$ (Top), $^3J_{CC}$ (Upper Middle), $^1J_{H\alpha C\alpha}$ (Middle), $^1J_{NC\alpha}$ (Lower Middle) and $^2J_{NC\alpha}$ (Bottom) from Mantsyzov *et al.* and Lee *et al*. [13,18].

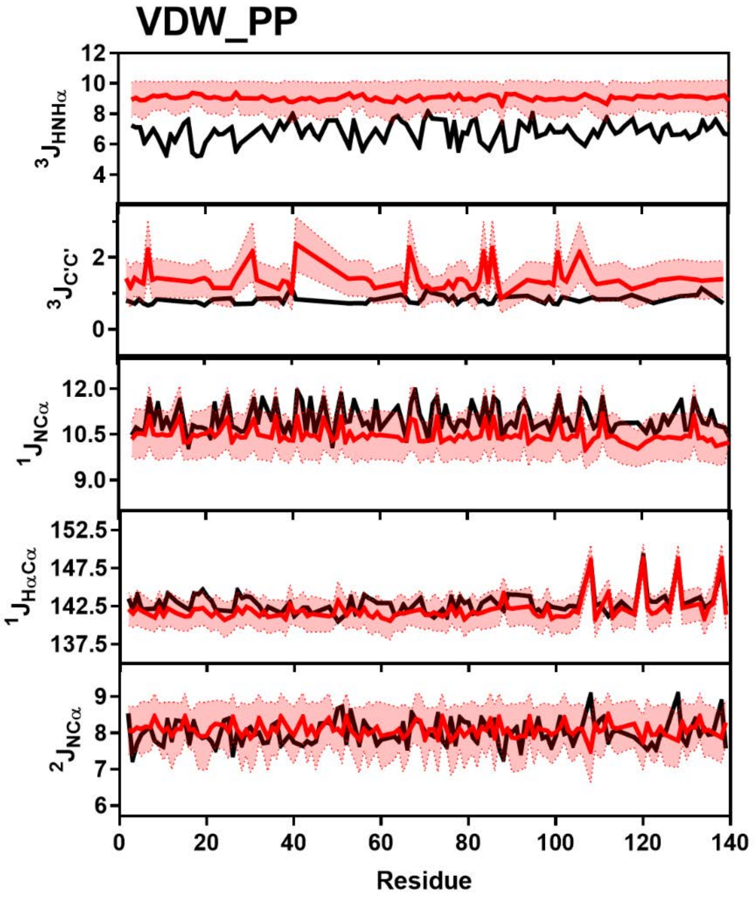


**Figure S88:** Comparison of Simulated and Experimental J-Coupling Data. Simulated J-Coupling values from the VDW_PP ensemble (red) overlayed on experimental data (black) of $^3J_{HNH\alpha}$ (Top), $^3J_{CC}$ (Upper Middle), $^1J_{H\alpha C\alpha}$ (Middle), $^1J_{NC\alpha}$ (Lower Middle) and $^2J_{NC\alpha}$ (Bottom) from Mantsyzov *et al.* and Lee *et al*. [13,18].

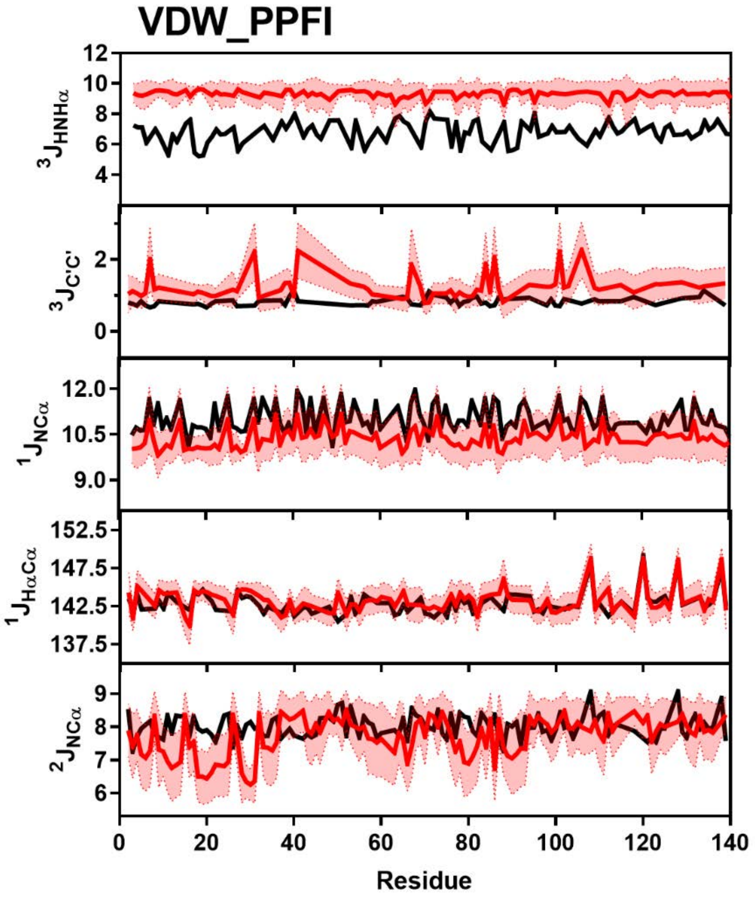


**Figure S89:** Comparison of Simulated and Experimental J-Coupling Data. Simulated J-Coupling values from the VDW_PPFI ensemble (red) overlayed on experimental data (black) of $^3J_{HNH\alpha}$ (Top), $^3J_{CC}$ (Upper Middle), $^1J_{H\alpha C\alpha}$ (Middle), $^1J_{NC\alpha}$ (Lower Middle) and $^2J_{NC\alpha}$ (Bottom) from Mantsyzov *et al.* and Lee *et al*. [13,18].

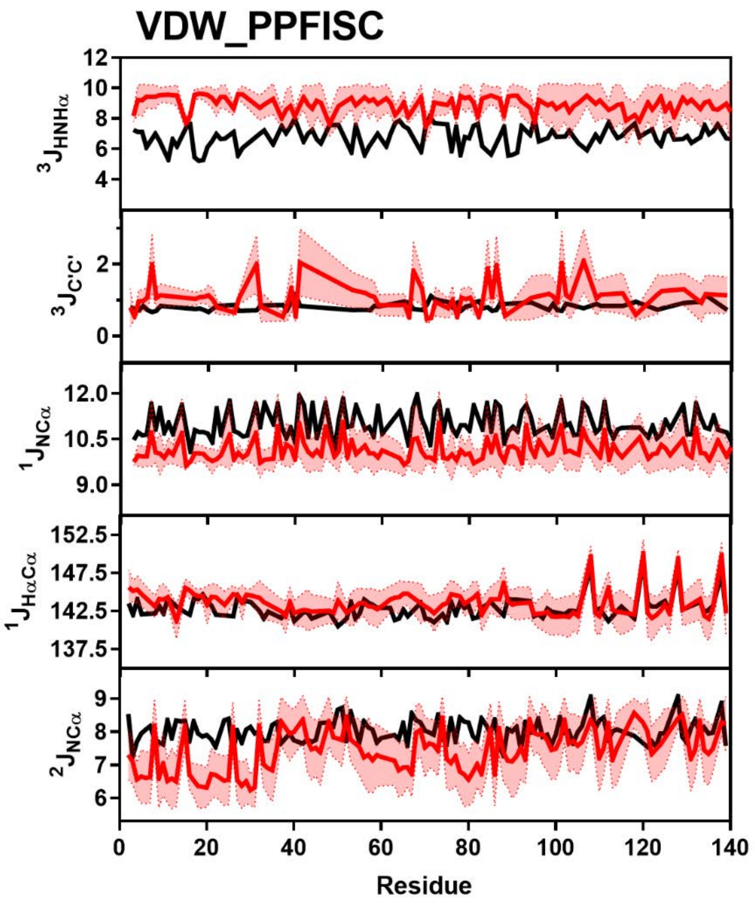


**Figure S90:** Comparison of Simulated and Experimental J-Coupling Data. Simulated J-Coupling values from the VDW_PPFISC ensemble (red) overlayed on experimental data (black) of $^3J_{HNH\alpha}$ (Top), $^3J_{CC}$ (Upper Middle), $^1J_{H\alpha C\alpha}$ (Middle), $^1J_{NC\alpha}$ (Lower Middle) and $^2J_{NC\alpha}$ (Bottom) from Mantsyzov *et al.* and Lee *et al*. [13,18].

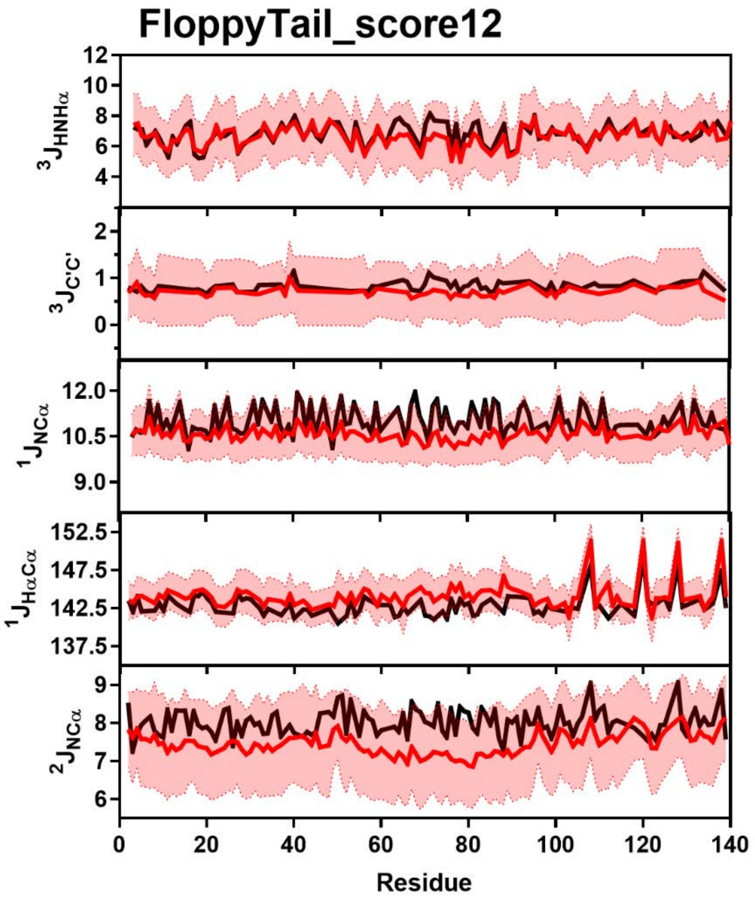


**Figure S91:** Comparison of Simulated and Experimental J-Coupling Data. Simulated J-Coupling values from the FloppyTail_score12 ensemble (red) overlayed on experimental data (black) of $^3J_{HNH\alpha}$ (Top), $^3J_{CC}$ (Upper Middle), $^1J_{H\alpha C\alpha}$ (Middle), $^1J_{NC\alpha}$ (Lower Middle) and $^2J_{NC\alpha}$ (Bottom) from Mantsyzov *et al.* and Lee *et al*. [13,18].

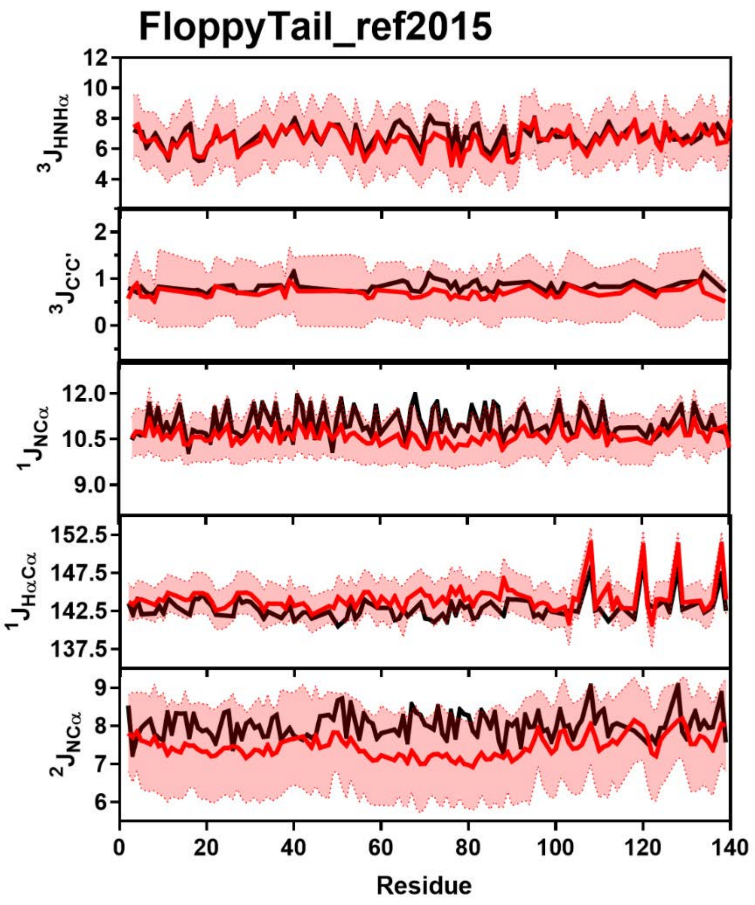


**Figure S92:** Comparison of Simulated and Experimental J-Coupling Data. Simulated J-Coupling values from the FloppyTail_ref2015 ensemble (red) overlayed on experimental data (black) of $^3J_{HNH\alpha}$ (Top), $^3J_{CC}$ (Upper Middle), $^1J_{H\alpha C\alpha}$ (Middle), $^1J_{NC\alpha}$ (Lower Middle) and $^2J_{NC\alpha}$ (Bottom) from Mantsyzov *et al.* and Lee *et al*. [13,18].

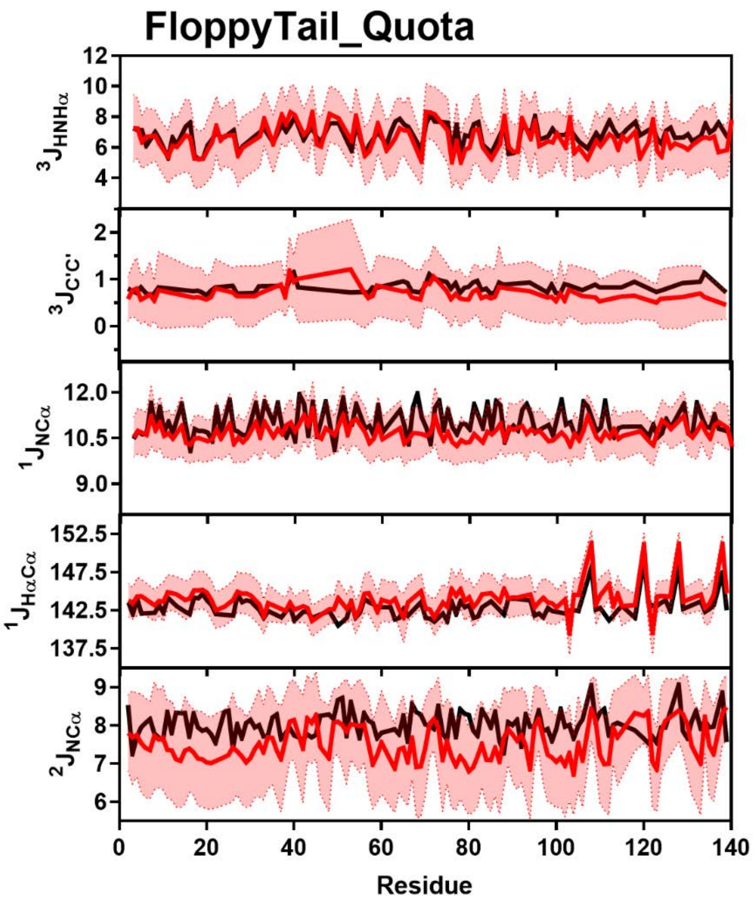


**Figure S93:** Comparison of Simulated and Experimental J-Coupling Data. Simulated J-Coupling values from the FloppyTail_Quota ensemble (red) overlayed on experimental data (black) of $^3J_{HNH\alpha}$ (Top), $^3J_{CC}$ (Upper Middle), $^1J_{H\alpha C\alpha}$ (Middle), $^1J_{NC\alpha}$ (Lower Middle) and $^2J_{NC\alpha}$ (Bottom) from Mantsyzov *et al.* and Lee *et al*. [13,18].

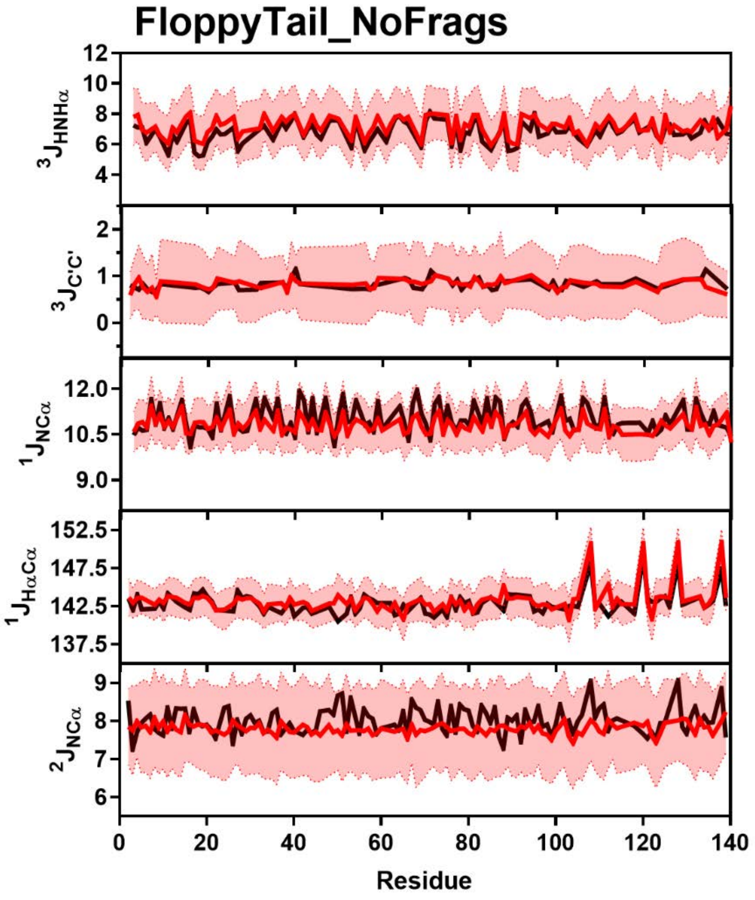


**Figure S94:** Comparison of Simulated and Experimental J-Coupling Data. Simulated J-Coupling values from the FloppyTail_NoFrags ensemble (red) overlayed on experimental data (black) of $^3J_{HNH\alpha}$ (Top), $^3J_{CC}$ (Upper Middle), $^1J_{H\alpha C\alpha}$ (Middle), $^1J_{NC\alpha}$ (Lower Middle) and $^2J_{NC\alpha}$ (Bottom) from Mantsyzov *et al.* and Lee *et al*. [13,18].

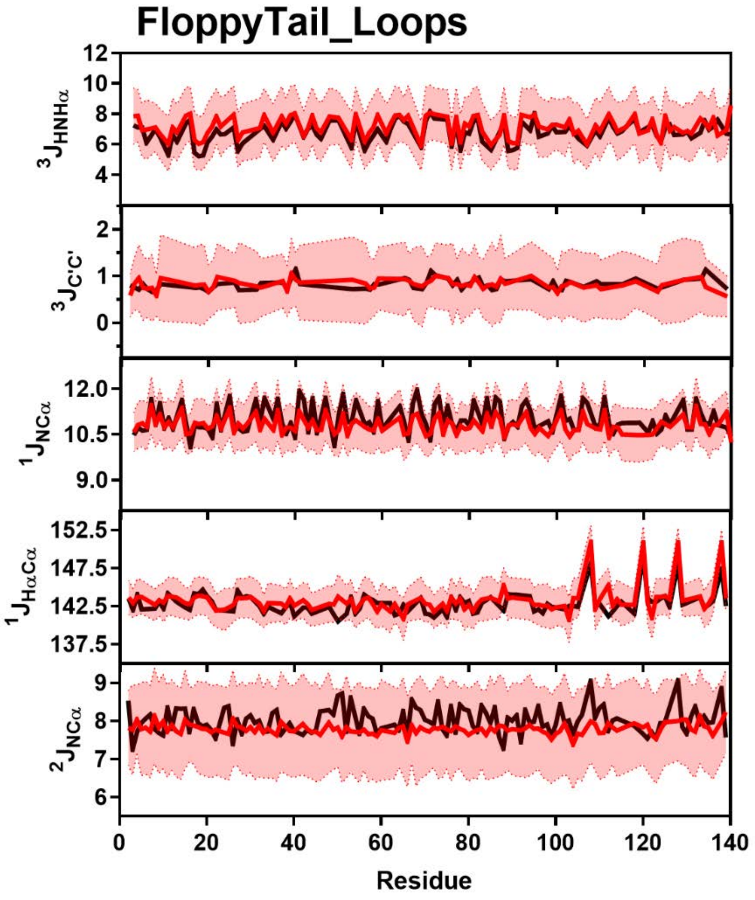


**Figure S95:** Comparison of Simulated and Experimental J-Coupling Data. Simulated J-Coupling values from the FloppyTail_Loops ensemble (red) overlayed on experimental data (black) of $^3J_{HNH\alpha}$ (Top), $^3J_{CC}$ (Upper Middle), $^1J_{H\alpha C\alpha}$ (Middle), $^1J_{NC\alpha}$ (Lower Middle) and $^2J_{NC\alpha}$ (Bottom) from Mantsyzov *et al.* and Lee *et al*. [13,18].

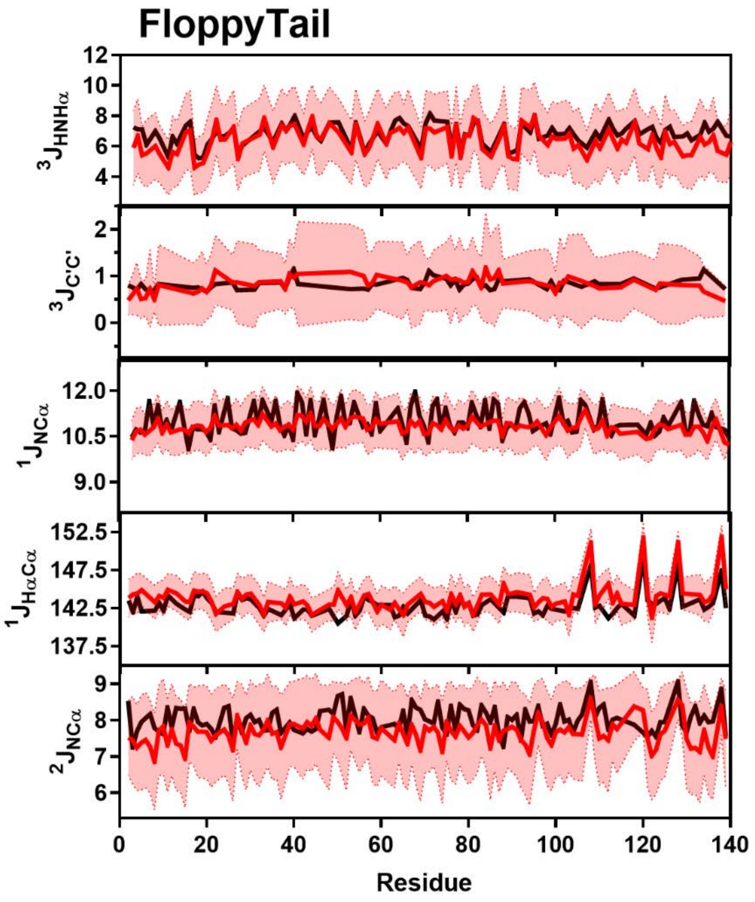


**Figure S96:** Comparison of Simulated and Experimental J-Coupling Data. Simulated J-Coupling values from the FloppyTail ensemble (red) overlayed on experimental data (black) of $^3J_{HNH\alpha}$ (Top), $^3J_{CC}$ (Upper Middle), $^1J_{H\alpha C\alpha}$ (Middle), $^1J_{NC\alpha}$ (Lower Middle) and $^2J_{NC\alpha}$ (Bottom) from Mantsyzov *et al.* and Lee *et al*. [13,18].

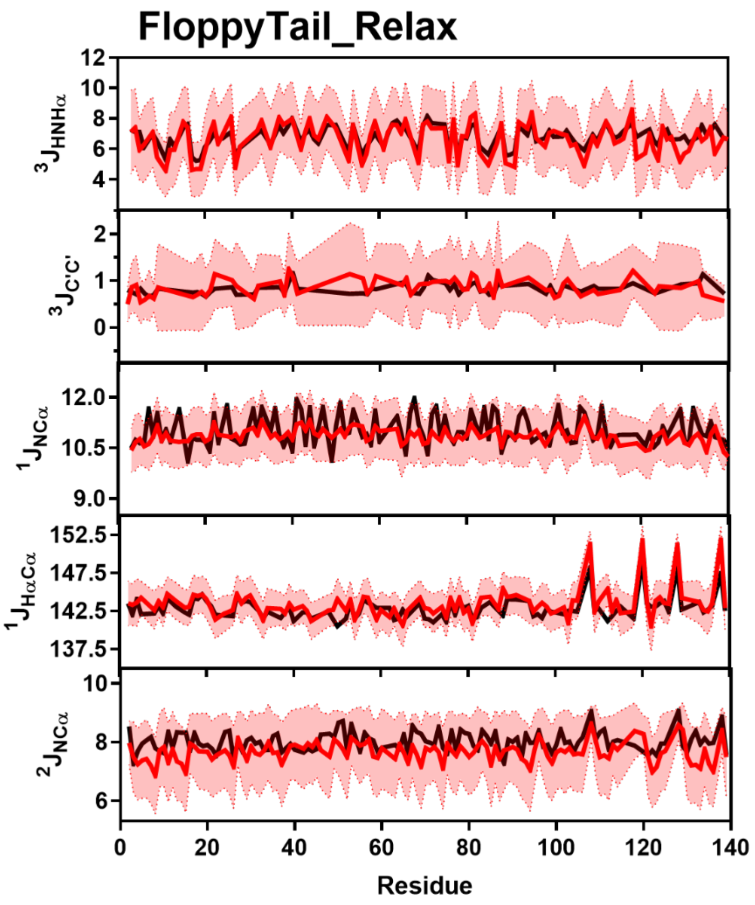


**Figure S97:** Comparison of Simulated and Experimental J-Coupling Data. Simulated J-Coupling values from the FloppyTail_Relax ensemble (red) overlayed on experimental data (black) of $^3J_{HNH\alpha}$ (Top), $^3J_{CC}$ (Upper Middle), $^1J_{H\alpha C\alpha}$ (Middle), $^1J_{NC\alpha}$ (Lower Middle) and $^2J_{NC\alpha}$ (Bottom) from Mantsyzov *et al.* and Lee *et al.* [13,18].

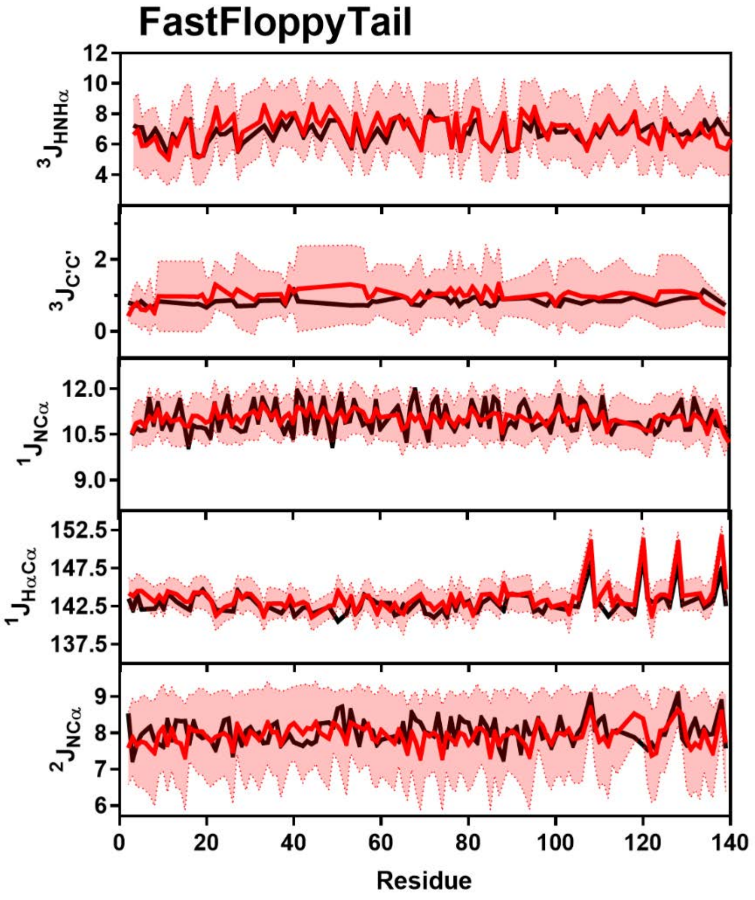


**Figure S98:** Comparison of Simulated and Experimental J-Coupling Data. Simulated J-Coupling values from the FastFloppyTail ensemble (red) overlayed on experimental data (black) of $^3J_{HNH\alpha}$ (Top), $^3J_{CC}$ (Upper Middle), $^1J_{H\alpha C\alpha}$ (Middle), $^1J_{NC\alpha}$ (Lower Middle) and $^2J_{NC\alpha}$ (Bottom) from Mantsyzov *et al.* and Lee *et al*. [13,18].

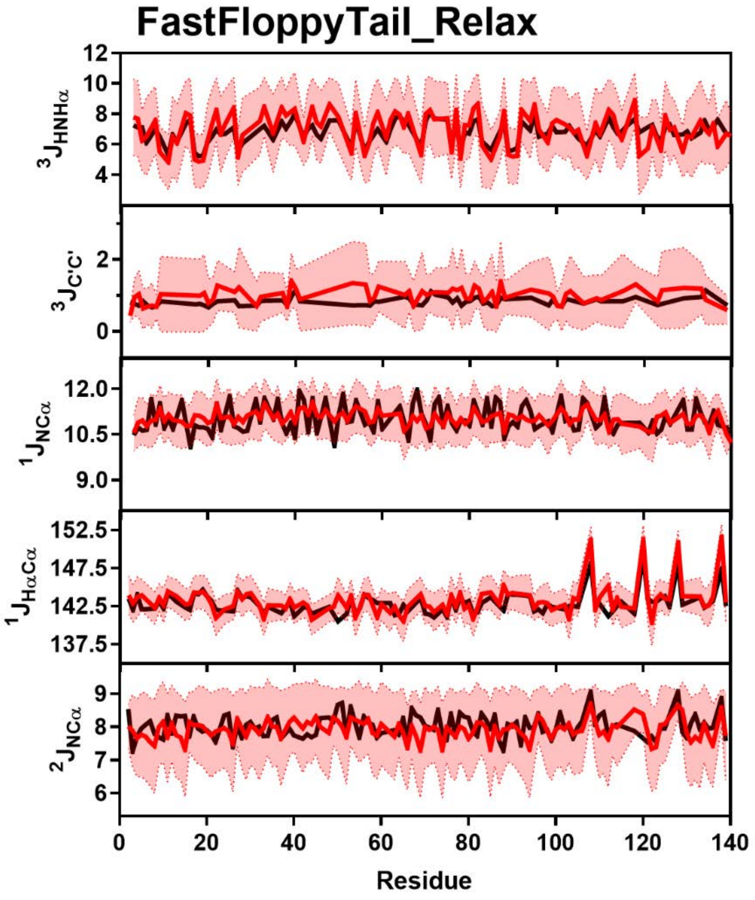


**Figure S99:** Comparison of Simulated and Experimental J-Coupling Data. Simulated J-Coupling values from the FastFloppyTail_Relax ensemble (red) overlayed on experimental data (black) of $^3J_{HNH\alpha}$ (Top), $^3J_{CC}$ (Upper Middle), $^1J_{H\alpha C\alpha}$ (Middle), $^1J_{NC\alpha}$ (Lower Middle) and $^2J_{NC\alpha}$ (Bottom) from Mantsyzov *et al.* and Lee *et al*. [13,18].

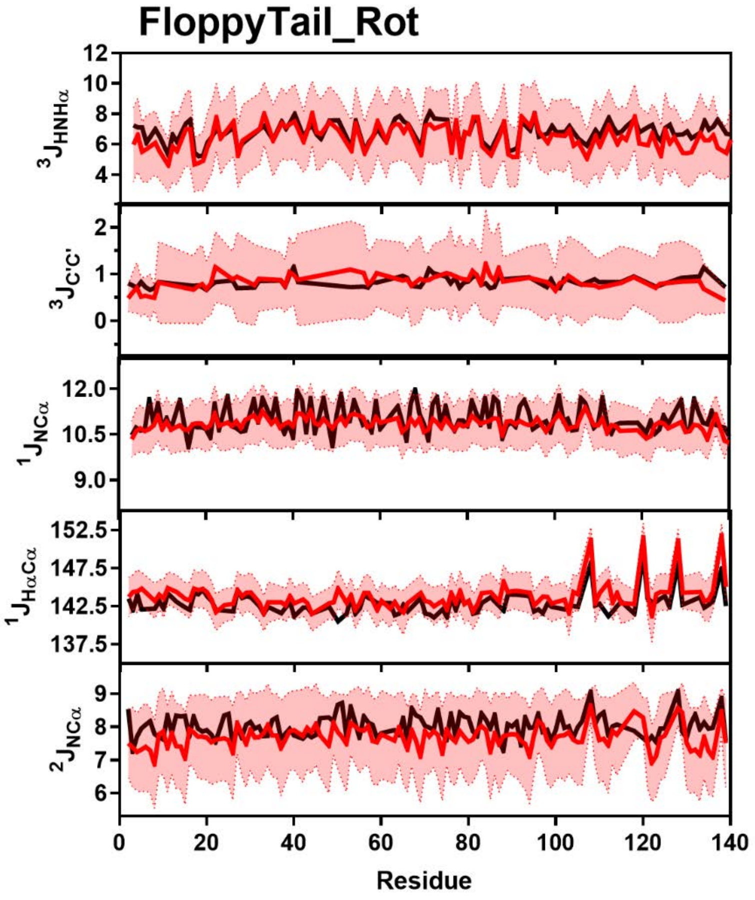


**Figure S100:** Comparison of Simulated and Experimental J-Coupling Data. Simulated J-Coupling values from the FloppyTail_Rot ensemble (red) overlayed on experimental data (black) of $^3J_{HNH\alpha}$ (Top), $^3J_{CC}$ (Upper Middle), $^1J_{H\alpha C\alpha}$ (Middle), $^1J_{NC\alpha}$ (Lower Middle) and $^2J_{NC\alpha}$ (Bottom) from Mantsyzov *et al.* and Lee *et al*. [13,18].

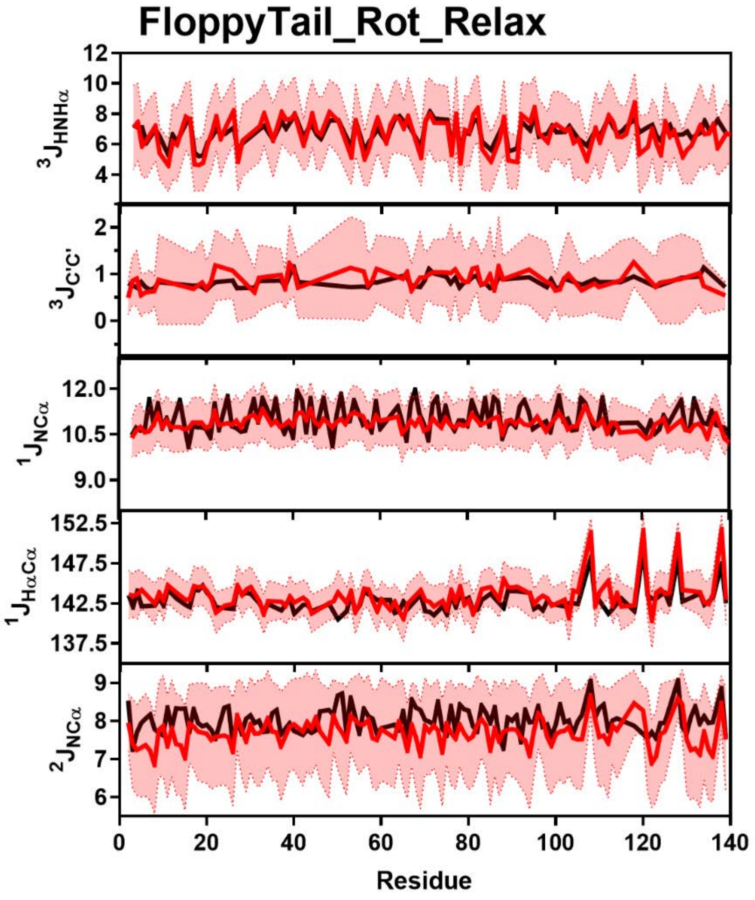


**Figure S101:** Comparison of Simulated and Experimental J-Coupling Data. Simulated J-Coupling values from the FloppyTail_Rot_Relax ensemble (red) overlayed on experimental data (black) of $^3J_{HNH\alpha}$ (Top), $^3J_{CC}$ (Upper Middle), $^1J_{H\alpha C\alpha}$ (Middle), $^1J_{NC\alpha}$ (Lower Middle) and $^2J_{NC\alpha}$ (Bottom) from Mantsyzov *et al.* and Lee *et al*. [13,18].

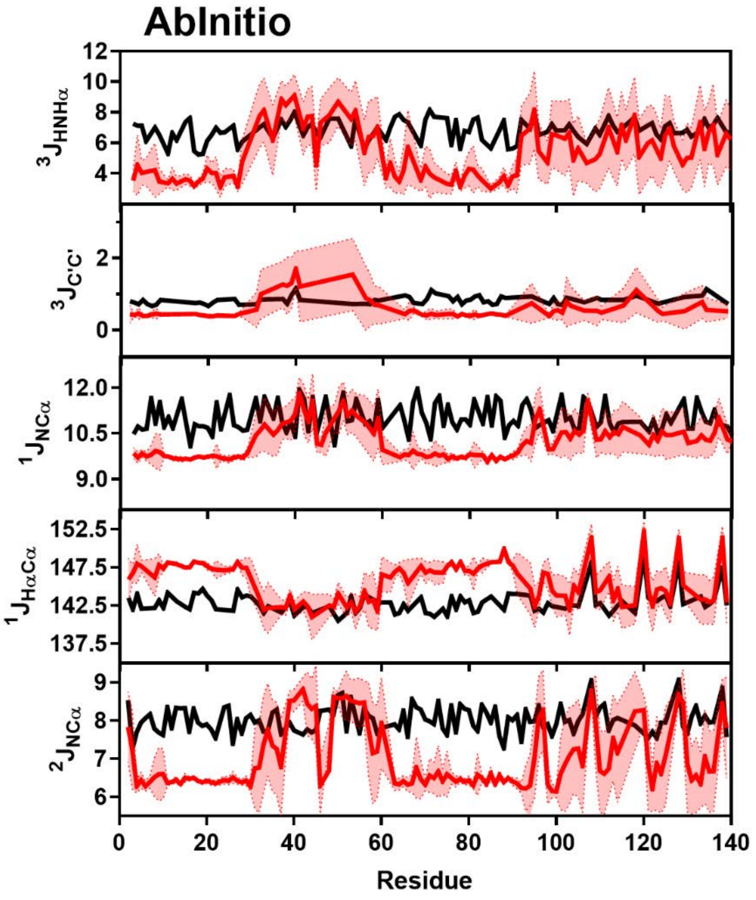


**Figure S102:** Comparison of Simulated and Experimental J-Coupling Data. Simulated J-Coupling values from the AbInitio ensemble (red) overlayed on experimental data (black) of $^3J_{HNH\alpha}$ (Top), $^3J_{CC}$ (Upper Middle), $^1J_{H\alpha C\alpha}$ (Middle), $^1J_{NC\alpha}$ (Lower Middle) and $^2J_{NC\alpha}$ (Bottom) from Mantsyzov *et al.* and Lee *et al*. [13,18].

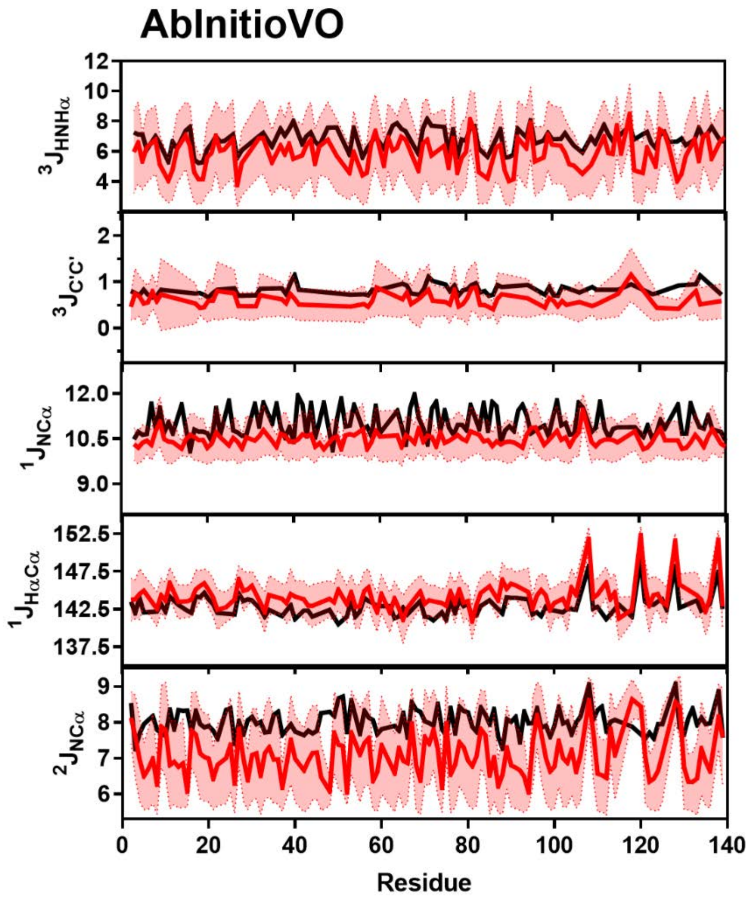


**Figure S103:** Comparison of Simulated and Experimental J-Coupling Data. Simulated J-Coupling values from the AbInitioVO ensemble (red) overlayed on experimental data (black) of $^3J_{HNH\alpha}$ (Top), $^3J_{CC}$ (Upper Middle), $^1J_{H\alpha C\alpha}$ (Middle), $^1J_{NC\alpha}$ (Lower Middle) and $^2J_{NC\alpha}$ (Bottom) from Mantsyzov *et al.* and Lee *et al*. [13,18].

**Construction of the New Rg Score Term:** The previous "rg" centroid score term used by Rosetta is equal to a constant multiplied by the value of the radius of gyration of the protein at each step of the simulation. Therefore, even when a protein approaches its appropriate radius of gyration, this term continues to force compaction energetically. In our effort to abrogate this effect, which leads to the production of overly-compact conformations of disordered proteins, we have chosen to craft the potential in the form of a self-avoiding walk (SAW) probability distribution.

$$P(r) = A\frac{4\pi}{\overline{R_g}}\left(\frac{r_g}{\overline{R_g}}\right)^{2+g} \exp\left[-\alpha\left(\frac{r_g}{\overline{R_g}}\right)^{\delta}\right] \quad \text{(Eqn. S8)}$$ [27]

$$g = {}^{(\gamma - 1)}/_{\nu} \quad \text{(Eqn. S9)}$$ [27]

$$\delta = {}^{1}/_{(1-\nu)} \quad \text{(Eqn. S10)}$$ [27]

$$\overline{R_g} = \sqrt{\frac{2l_p b}{(2\nu+1)(2\nu+2)}}N^{\nu} \quad \text{(Eqn. S11)}$$ [28]

Here, $r_g$ is the radius of gyration of the protein in a particular conformation, $\overline{R_g}$ is the average radius of gyration for a given protein, $\nu$ is the scaling exponent that, as exemplified in Eqn. S11, relates the number of residues in a protein $N$ to an average $Rg$. The constants $b$ (0.38 nm), $l_p$ (0.53 nm), and $\gamma$ (1.1615) have been previously estimated for proteins and the constants $A$ and $\alpha$ are determined for each set of $\nu$ and $\overline{R_g}$ through the conditions $\int_0^{\infty} P(r_g)dr_g = 1$ and $\int_0^{\infty} P(r_g){r_g}^2 dr = \overline{R_g}^2$ [27,28]. This provides for both a broad distribution of radii of gyration as well as an energetic barrier for overly compact states. The energy potential is defined as:

$$E_{\overline{R_g}}(r_g) = 1 - {}^{P(r_g)}/_{max[P(r_g)]} \quad \text{(Eqn. S12)}$$

Through this form, the energy value scales from zero to one for any average radius of gyration allowing the weight of this score term within our overall score function to determine the depth of the potential.

To utilize this potential, a radius of gyration needs to be predicted from the primary sequence. Here, we utilize functional forms described in Hofmann *et al.* to predict the scaling exponent for a given protein based on the primary sequence composition and predict the radius of gyration using Eqn. S11 [28]. The scaling exponent can be predicted from either the net charge or hydrophobicity of the protein via the equations:

$$\nu(Q) = \frac{1}{3} + a\left[1 + \frac{\exp(x_0 - Q)}{z}\right]^{-1} \quad \text{(Eqn. S13)}$$ [28]

$$\nu(H) = \frac{1}{3} + a\left[1 + \frac{\exp(x_0 + cH - d)}{z}\right]^{-1} \quad \text{(Eqn. S14)}$$ [28]

The parameters $a$ = 0.394, $z$ = 0.009, $x_0$ = 0.114, $c$ = 1.72 and $d$ = 0.9, were determined by fitting observed radii of gyration for a set of proteins in guanidinium hydrochloride to the above equations [28].

To select between the scaling factors computed by the net charge and hydrophobicity when computing an expected radius of gyration, the polyampholyte theory was applied to determine the effect of the net charge on the excluded volume of the system by computing the excess volume:

$$v^* = \frac{4\pi l_B (f-g)^2}{\kappa^2} - \frac{\pi {l_B}^2 (f+g)^2}{\kappa} \qquad \text{(Eqn. S15)}^{28}$$

Here, $f$ and $g$ are the fraction of positively (Arg/Lys) and negatively (Glu/Asp) charged residues, respectively, $\kappa$ is the Debye length defined by $\kappa^{-1} = {0.304}/{\sqrt{I}}$, where $I$ is the ionic strength and $l_B$ is the Bjerrum length defined by:

$$l_B = {e^2}/{(4\pi \varepsilon_0 \varepsilon_r k_B T)} \qquad \text{(Eqn. S16)}^{28}$$

In the above equation, $e$ is the elementary charge ($1.602 \times 10^{-19}$), $\varepsilon_0$ is the dielectric constant ($8.854 \times 10^{-12}$), $\varepsilon_r$ is the permittivity of water (78.7) and $k_B$ is the Boltzmann constant. In the general case for the calculation of $v^*$, the ionic strength and the temperature are set to 0.15 M and 298 K respectively. Since values of $v^*$ greater than zero correspond to net electrostatic repulsion, when $v^*$ is greater than zero, the scaling term computed using the net charge is applied when predicting the radius of gyration. Conversely, when $v^*$ is less than zero, there is a net electrostatic attraction, forcing the scaling based on the hydrophobicity to dominate. In cases where both the fractions of positively and negatively charged residues are zero, the scaling is determined using the net hydrophobicity. Lastly, in the cases where either the fraction of positively or negatively charged residues equals zero, the scaling due to the net charge is selected.

Upon further inspection of the above method proposed by Hofmann *et al.*, we noted that when testing this function for lysozyme (PDB: 2LZM) the predicted scaling value ($v$ = 0.519) is significantly higher than the expected value for a folded protein [28]. Since the fitting was performed on denatured proteins, the work by Hofmann *et al.* shows that when determining the radius of gyration for a suite of proteins from the Protein Data Bank, the minimum observed scaling factor is not equal to 0.33, which is the value for well-folded proteins, but instead is ~ 0.4 [28]. Therefore, we utilized the per residue disordered probability of the protein to determine if a given segment of the protein of interest was folded or unfolded. The RaptorX server was used to obtain per residue disordered probabilities, where residues with a disordered probability >0.5 were determined to be disordered [6]. Segments were defined as any stretch of residues greater than 10 in which all residues were of the same classification, ordered or disordered. Segments deemed ordered were assigned a scaling value of 0.33 while the scaling value for disordered regions was computed using the above equations.

**Implementation of the New Rg Score Term in PyRosetta:** The updated "rg" term was implemented within PyRosetta by producing a new score term using the WholeStructureEnergy object. The annotated script (available upon email request to JJF or EJP) demonstrates how this is accomplished in detail, and the method is explained in brief below. After producing a pose from the input sequence, the net disordered probability was computed from the per residue disordered probability from RaptorX [6]. For net disordered probabilities below 0.5, a scaling constant value of 0.33 was assigned. For all others, scaling values were computed from the net charge and hydrophobicitiy and a scaling constant value was selected using the effect of the net charge on the excluded volume as detailed above. After determining the value of the scaling constant, an expected average radius of gyration and radii of gyration probability distribution described by a self-avoiding random walk were computed. The values for $A$ and $\alpha$ were obtained using a discrete sum of distances from zero to $5 \times R_g$ Å at a spacing of 0.01 Å. The probability distribution was converted in a potential energy function as described above and stored as the interpolated function after conversion using the scipy interp1d functionality [29]. Each time the pose was scored, the radius of gyration was computed using the neighbor atom coordinates and the value of the new radius of gyration score term value was assigned at each residue as the energy determined from the interpolated potential divided by the total number of residues. The weight of the score term within the score1-3 score functions was set at 80.0, which was determined to be optimal via the process detailed below.

**Determination of the Optimal Weight for the New Rg Score Term:** To determine the optimal weight of the new $R_g$ score term, αS was simulated using a range of weighting values. We identified the optimal weighting value as that which maximized the impact of the score term on the average radius of gyration while minimizing the restriction on the conformational diversity.

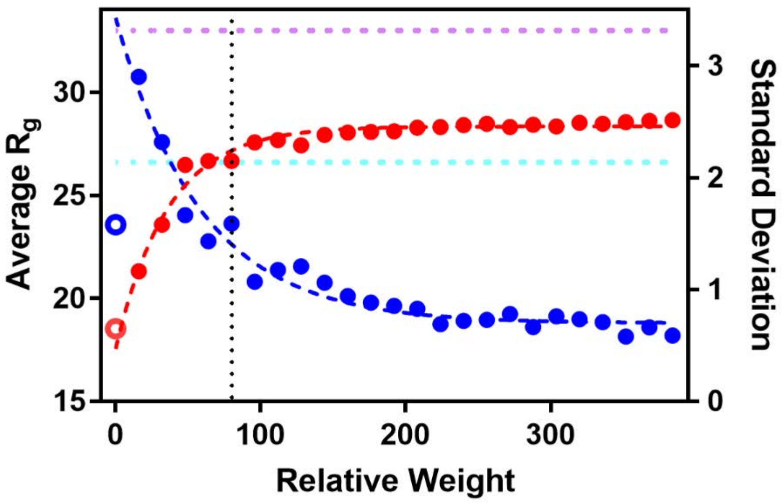


**Figure S104:** Determination of optimal new $R_g$ score term weight. Plot shows the average (filled blue circles) and standard deviation (filled red circles) radius of gyration resultant from simulations with various relative weights of the new $R_g$ score term compared to the simulation lacking the traditional rg term (open red and blue circles) and experimental data from SAXS (pink dashed line) and NMR (cyan dashed line). The vertical black dashed line illustrates the chosen optimal score term weight, as this pushes the average *Rg* closest to the experimental value without over-constraining the sampling. Therefore, the weights of all terms in the new score3 term is equivalent to the traditional score3 term, with the canonical rg term set to weight a zero and the weight of this new term set to 84.

### Comparison of AbInitio and AbInitioVO Simulations to Experimental Data

#### Simulated Proteins and Abbreviations

The 25-protein test set to assessing the accuracy of AbInitioVO compared to AbInito consisted of anti-HIV protein AOP-RANTES (1b3a), α-spectrin SH3 domain D46G mutant (1bk2), Rubredoxin from Pyrococcus Furiosus (1bq9), Drosophila engrailed homeodomain (1enh), Protein L B1 domain from Peptosteptococcus Magnus (1hz6), B2 immunoglobulin-binding domain of Streptococcal Protein G (1pgx), amino-terminal domain of phage 434 repressor (1r69), Human FYN SH3 domain (1shf), ubiquitin (1ubi), CRO repressor protein from bacteriophage lambda (5cro), Human translation initiation factor EIF1A (1d7q), Human co-chaperone P23 (1ejf), C-terminal domain of 50S ribosomal protein L11 (1fox), histone H1 (1ghc), HMGB protein NHP6A (1lwm), subunit F6 from the peripheral stalk region of ATP synthase (1vzs), Rap1 BRCT domain from Saccharomyces Cerevisiae (2l42), yeast Tah1 (2lsu), α-synuclein (αS or asyn), Drosophila adaptor protein Drk N-terminal SH3 domain (dsh3), C-terminal domain of nucleoprotein of measles virus (ntal), PaaA2 antitoxin from human pathogen *E. coli* O157 (paaa), cyclin-dependent kinase inhibitor Sic1 (sic1), and microtubule-associated protein tau K18 isoform (tauk).

### Summary of Simulated Proteins

**Table S5:** Comparison of AbInitio and AbInitioVO Outputs to PDB Structures

| PDB ID | # of Residues | Ordered Region | Fold Type | Min RMSD AbInitio | Min RMSD AbInitioVO | <Rg> AbInitio | <Rg> AbInitioVO | Exp Rg |
|---|---|---|---|---|---|---|---|---|
| | | | | ***Ordered*** | | | | |
| 1b3a | 67 | - | α/β | 2.68 | 3.31 | 12.0 | 11.8 | - |
| 1bk2 | 57 | - | β | 2.33 | 2.16 | 11.4 | 11.1 | - |
| 1bq9 | 54 | - | β | 2.64 | 3.91 | 11.0 | 10.7 | - |
| 1enh | 54 | - | α | 0.77 | 1.18 | 10.7 | 10.5 | - |
| 1hz6 | 64 | - | α/β | 2.16 | 1.99 | 12.4 | 12.1 | - |
| 1pgx | 70 | - | α | 3.71 | 3.54 | 13.0 | 13.0 | - |
| 1r69 | 63 | - | α | 0.99 | 1.14 | 10.7 | 10.5 | - |
| 1shf | 59 | - | β | 3.30 | 1.52 | 11.8 | 11.2 | - |
| 1ubi | 76 | - | α/β | 2.72 | 2.59 | 12.5 | 12.3 | - |
| 5cro | 61 | - | α/β | 4.35 | 4.56 | 11.3 | 11.0 | - |
| | | | | ***Partially-ordered*** | | | | |
| 1d7q | 144 | 27-115 | α/β | 6.38 | 6.41 | 17.4 | 21.8 | - |
| 1ejf | 160 | 1-110 | β | 6.97 | 7.73 | 18.4 | 26.5 | - |
| 1fox | 76 | 1-76 | α/β | 1.10 | 1.22 | 12.7 | 12.1 | - |
| 1ghc | 225 | 41-114 | α/β | 3.04 | 2.33 | 21.6 | 47.6 | - |
| 1lwm | 93 | 26-93 | α | 1.43 | 3.61 | 15.2 | 17.5 | - |
| 1vzs | 76 | 1-49 | α | 2.02 | 1.98 | 13.3 | 12.8 | - |
| 2l42 | 100 | 14-90 | α/β | 6.70 | 5.32 | 14.3 | 14.0 | - |
| 2lsu | 110 | 1-89 | α | 1.06 | 1.64 | 14.9 | 13.9 | - |
| | | | | ***Disordered*** | | | | |
| asyn | 140 | - | - | - | - | 17.4 | 28.4 | 33.0 / 26.6 |
| dsh3 | 59 | - | - | - | - | 12.3 | 11.5 | 18.3 |
| ntal | 125 | - | - | - | - | 15.9 | 27.0 | 27.5 |
| paaa | 71 | - | - | - | - | 18.4 | 19.6 | 22.4 |
| sic1 | 92 | - | - | - | - | 14.0 | 30.6 | 32.1 |
| tauk | 130 | - | - | - | - | 16.0 | 30.7 | 33.4 |

* Experimental values for the radius of gyration were determined for asyn from SAXS [19] and NMR [20] experiments, for dsh3 from NMR experiments capturing the Rg of the folded [30] and unfolded [31] states, for ntal from NMR experiments [32], for paaa from a simulated ensemble generated from SAXS and NMR experiments [33], for sic1 from simulated ensembles from SAXS and NMR experiments [34] and was determined from simulated ensembles based on NMR data for tauk [35].

**Comparison of Radii of Gyration:**

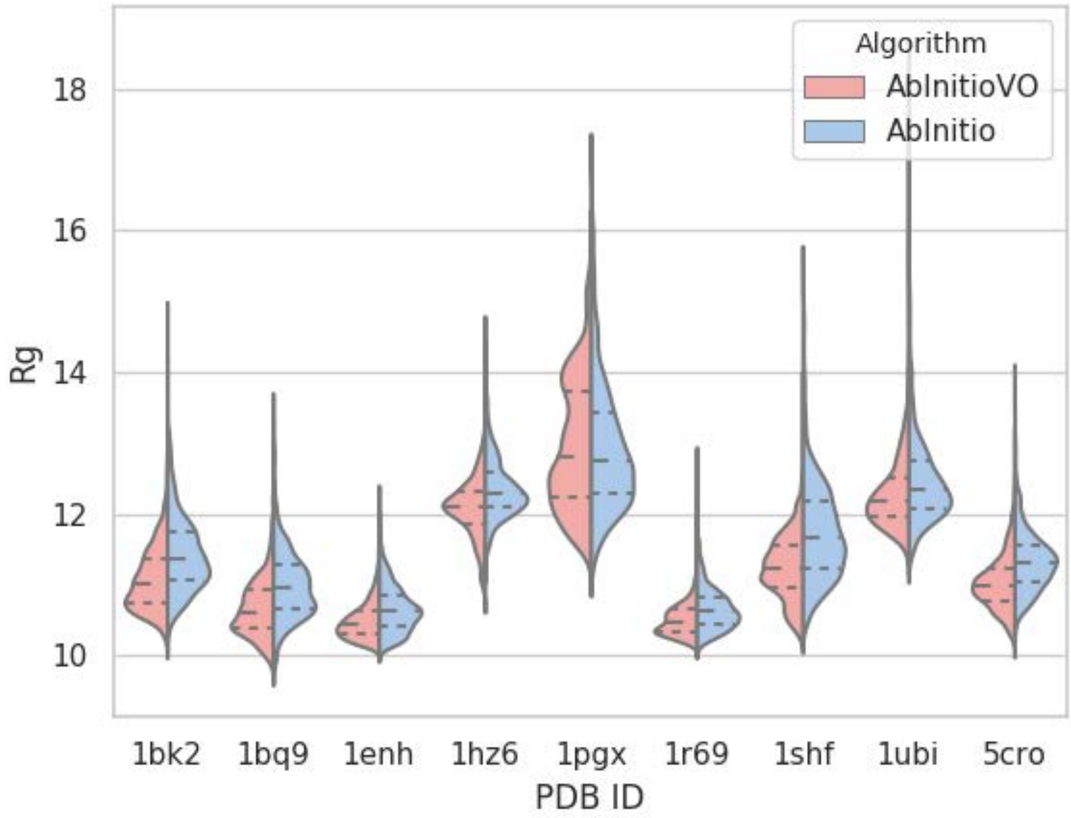


**Figure S105:** Radii of Gyration of Ordered Proteins. Violin plots show histogram for each simulation protein, as indicated by the x-axis, for all output from the AbInitioVO (red, left) and AbInitio (blue, right) simulations. Dashed lines on each violin represent the upper and lower quartile and the mean Rg.

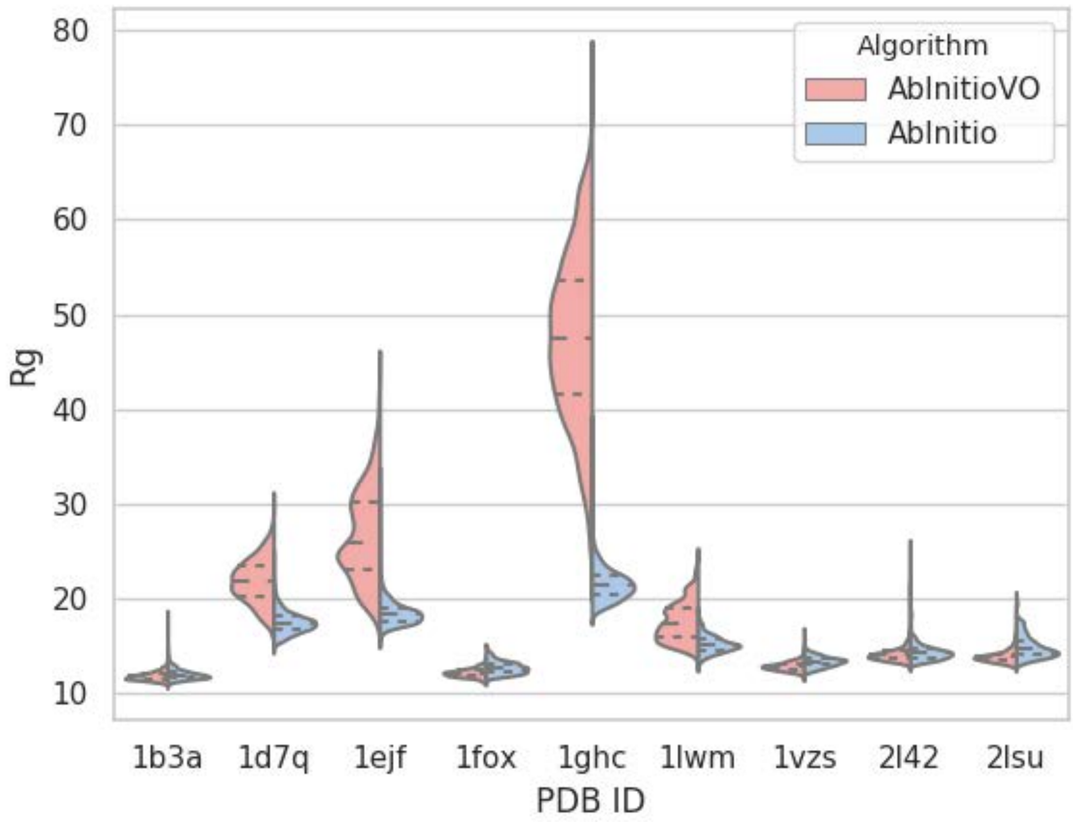


**Figure S106:** Radii of Gyration of Partially-Ordered Proteins. Violin plots show histogram for each simulation protein, as indicated by the x-axis, for all output from the AbInitioVO (red, left) and AbInitio (blue, right) simulations. Dashed lines on each violin represent the upper and lower quartile and the mean Rg.

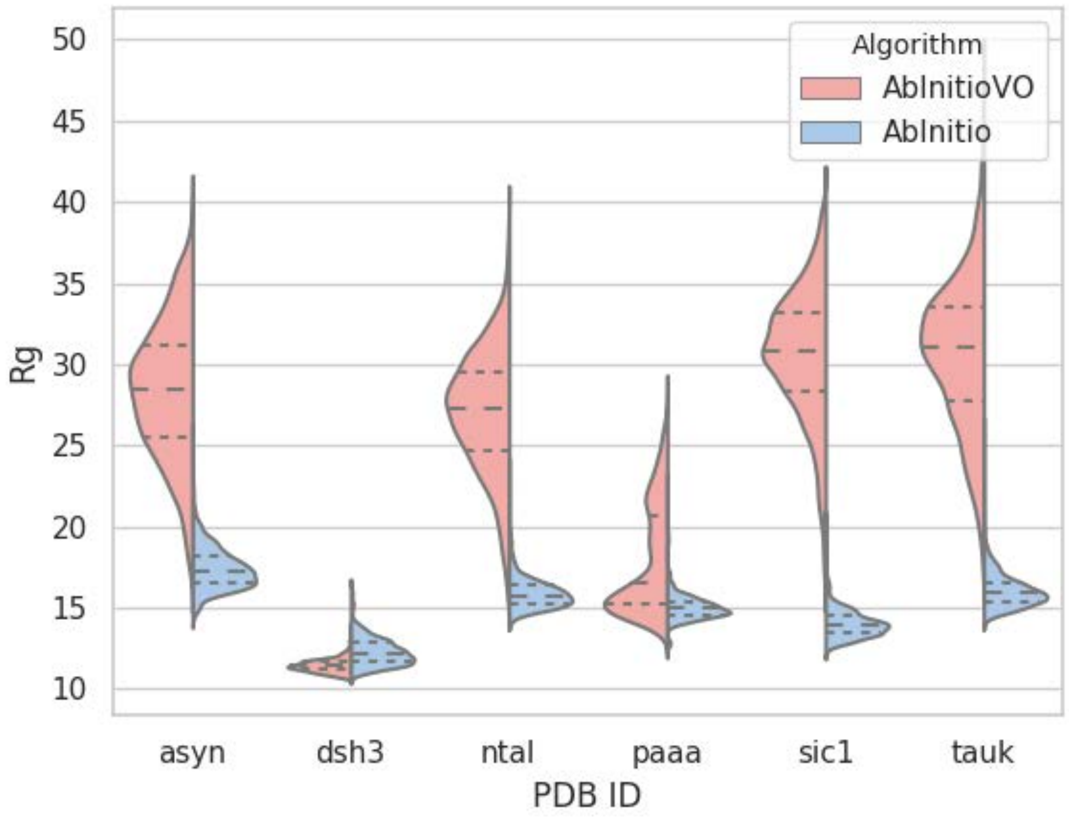


**Figure S107:** Radii of Gyration of Disordered Proteins. Violin plots show histogram for each simulation protein, as indicated by the x-axis, for all output from the AbInitioVO (red, left) and AbInitio (blue, right) simulations. Dashed lines on each violin represent the upper and lower quartile and the mean Rg.

### Comparison of Folding Funnels

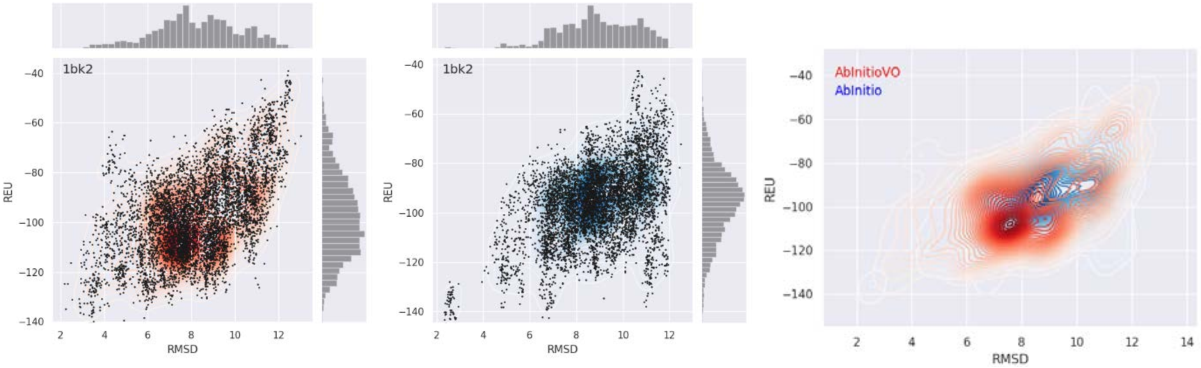


**Figure S108:** Folding Funnel Comparison for 1bka. Folding funnel from AbInitioVO (left) and AbInitio (middle) simulations plotting Cα RMSD versus Rosetta Energy Units (REU) from each structure compared to the PDB structure. For each folding funnel, histograms of the computed RMSD (top) and REU (right) are shown. KDE plot (right) showing overlay of AbInitiovo (red) and AbInitio (blue) folding funnels.

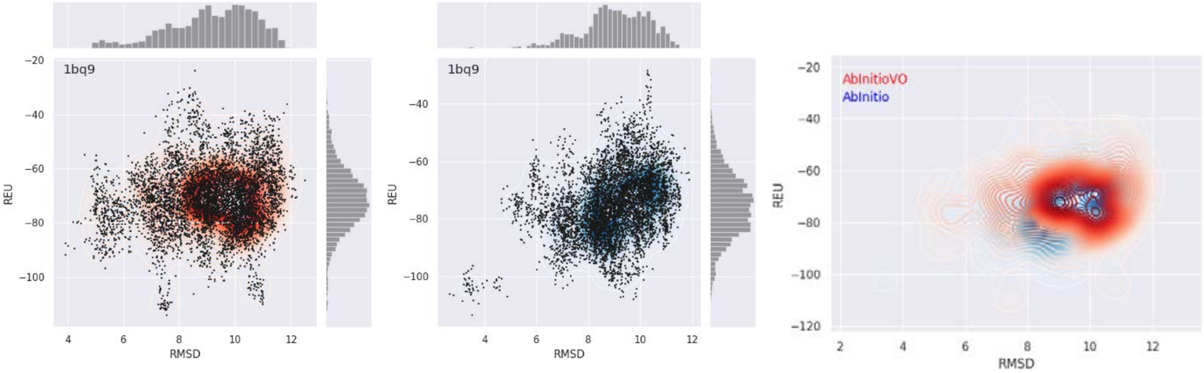


**Figure S109:** Folding Funnel Comparison for 1bq9. Folding funnel from AbInitioVO (left) and AbInitio (middle) simulations plotting Cα RMSD versus Rosetta Energy Units (REU) from each structure compared to the PDB structure. For each folding funnel, histograms of the computed RMSD (top) and REU (right) are shown. KDE plot (right) showing overlay of AbInitiovo (red) and AbInitio (blue) folding funnels.

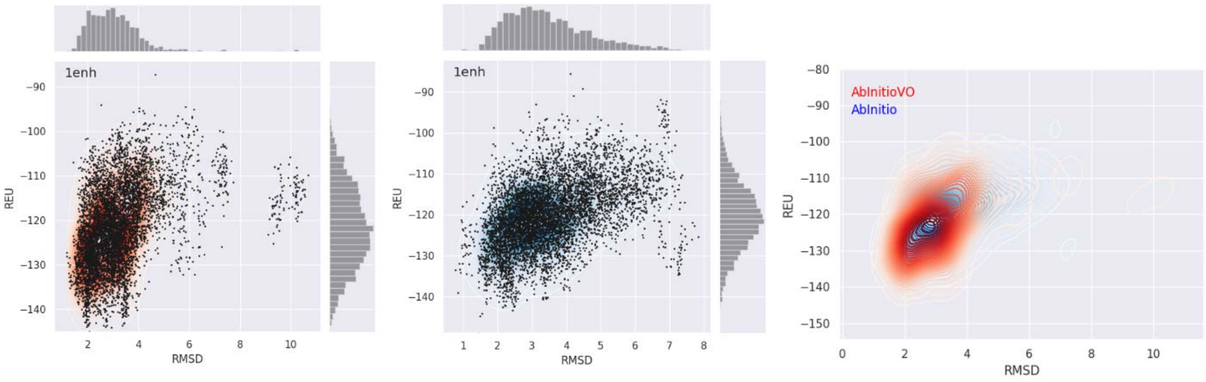


**Figure S110:** Folding Funnel Comparison for 1enh. Folding funnel from AbInitioVO (left) and AbInitio (middle) simulations plotting Cα RMSD versus Rosetta Energy Units (REU) from each structure compared to the PDB structure. For each folding funnel, histograms of the computed RMSD (top) and REU (right) are shown. KDE plot (right) showing overlay of AbInitiovo (red) and AbInitio (blue) folding funnels.

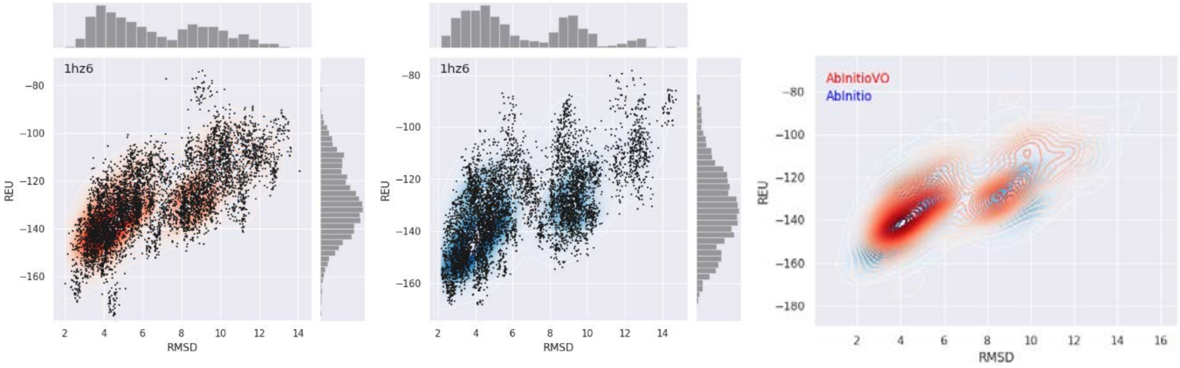


**Figure S111:** Folding Funnel Comparison for 1hz6. Folding funnel from AbInitioVO (left) and AbInitio (middle) simulations plotting Cα RMSD versus Rosetta Energy Units (REU) from each structure compared to the PDB structure. For each folding funnel, histograms of the computed RMSD (top) and REU (right) are shown. KDE plot (right) showing overlay of AbInitiovo (red) and AbInitio (blue) folding funnels.

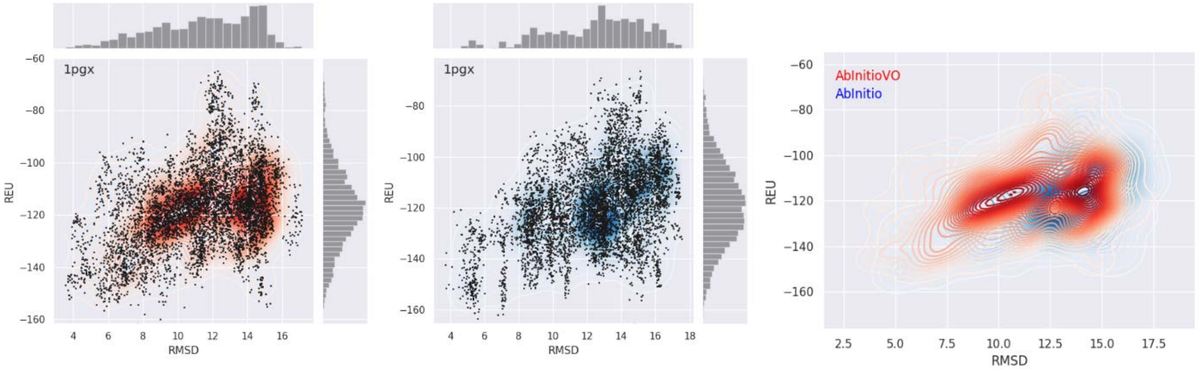


**Figure S112:** Folding Funnel Comparison for 1pgx. Folding funnel from AbInitioVO (left) and AbInitio (middle) simulations plotting Cα RMSD versus Rosetta Energy Units (REU) from each structure compared to the PDB structure. For each folding funnel, histograms of the computed RMSD (top) and REU (right) are shown. KDE plot (right) showing overlay of AbInitiovo (red) and AbInitio (blue) folding funnels.

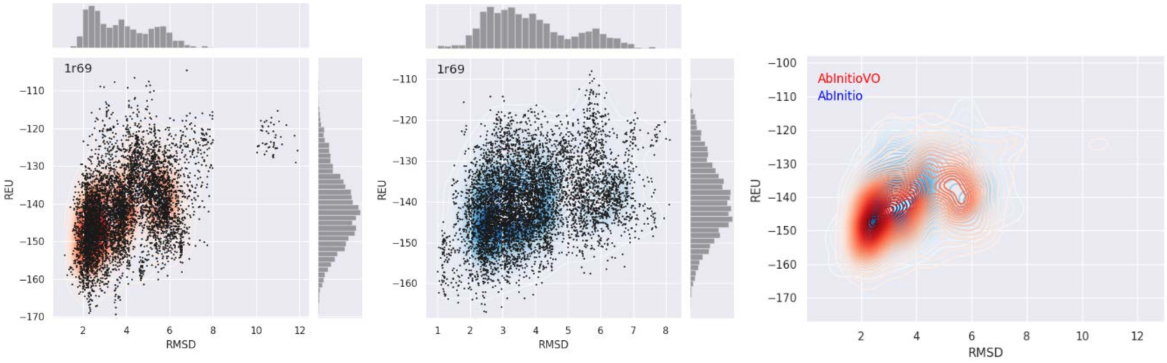


**Figure S113:** Folding Funnel Comparison for 1r69. Folding funnel from AbInitioVO (left) and AbInitio (middle) simulations plotting Cα RMSD versus Rosetta Energy Units (REU) from each structure compared to the PDB structure. For each folding funnel, histograms of the computed RMSD (top) and REU (right) are shown. KDE plot (right) showing overlay of AbInitiovo (red) and AbInitio (blue) folding funnels.

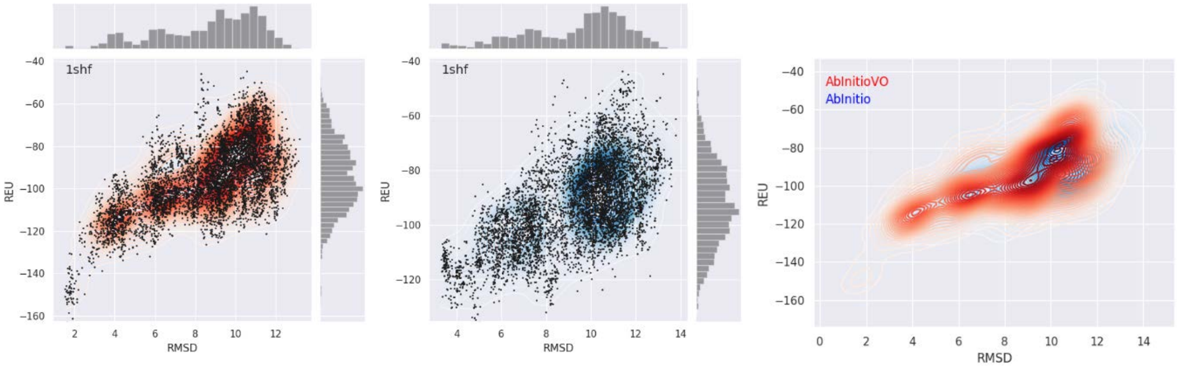


**Figure S114:** Folding Funnel Comparison for 1shf. Folding funnel from AbInitioVO (left) and AbInitio (middle) simulations plotting Cα RMSD versus Rosetta Energy Units (REU) from each structure compared to the PDB structure. For each folding funnel, histograms of the computed RMSD (top) and REU (right) are shown. KDE plot (right) showing overlay of AbInitiovo (red) and AbInitio (blue) folding funnels.

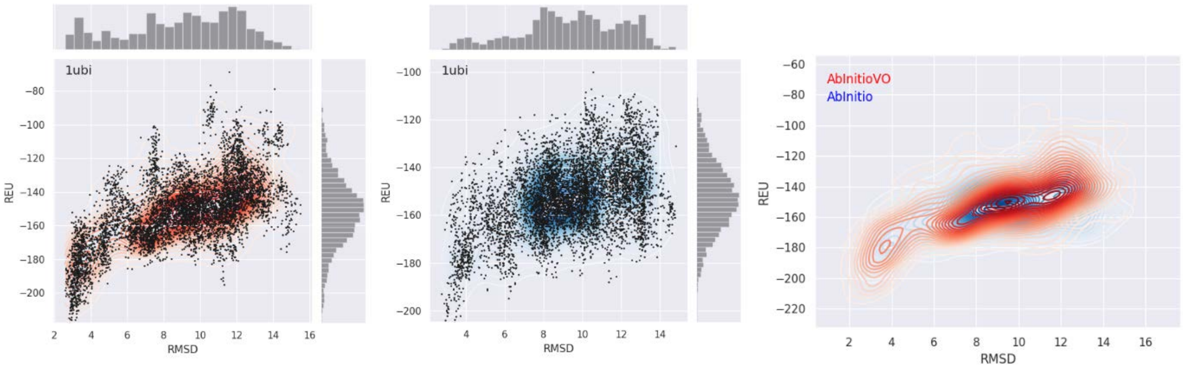


**Figure S115:** Folding Funnel Comparison for 1ubi. Folding funnel from AbInitioVO (left) and AbInitio (middle) simulations plotting Cα RMSD versus Rosetta Energy Units (REU) from each structure compared to the PDB structure. For each folding funnel, histograms of the computed RMSD (top) and REU (right) are shown. KDE plot (right) showing overlay of AbInitiovo (red) and AbInitio (blue) folding funnels.

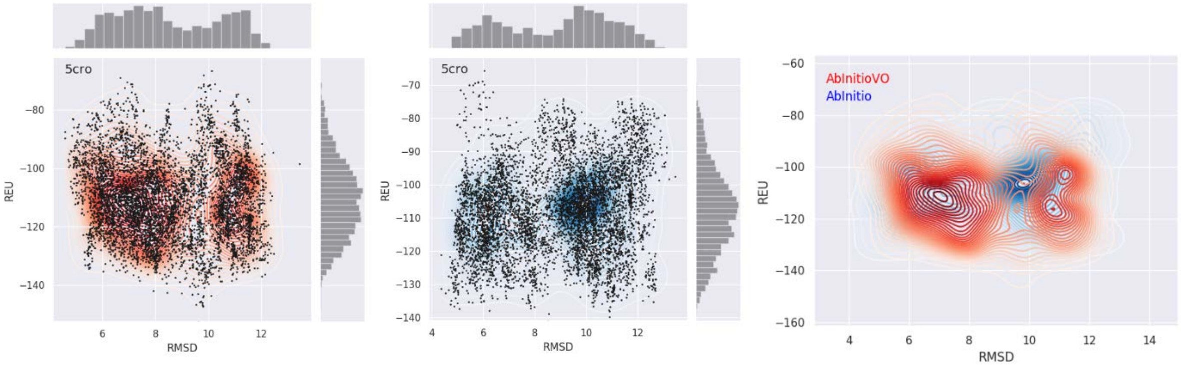


**Figure S116:** Folding Funnel Comparison for 5cro. Folding funnel from AbInitioVO (left) and AbInitio (middle) simulations plotting Cα RMSD versus Rosetta Energy Units (REU) from each structure compared to the PDB structure. For each folding funnel, histograms of the computed RMSD (top) and REU (right) are shown. KDE plot (right) showing overlay of AbInitiovo (red) and AbInitio (blue) folding funnels.

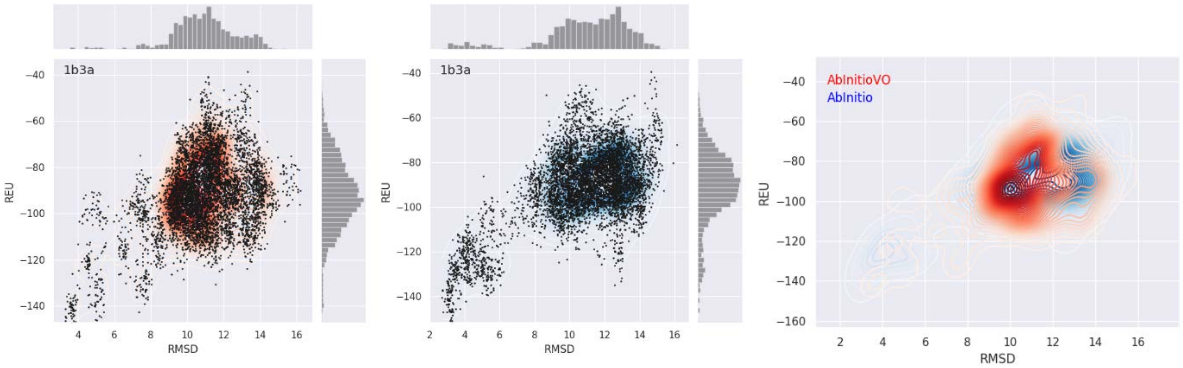


**Figure S117:** Folding Funnel Comparison for 1b3a. Folding funnel from AbInitioVO (left) and AbInitio (middle) simulations plotting Cα RMSD versus Rosetta Energy Units (REU) from each structure compared to the folded domain of the PDB structure. For each folding funnel, histograms of the computed RMSD (top) and REU (right) are shown. KDE plot (right) showing overlay of AbInitiovo (red) and AbInitio (blue) folding funnels.

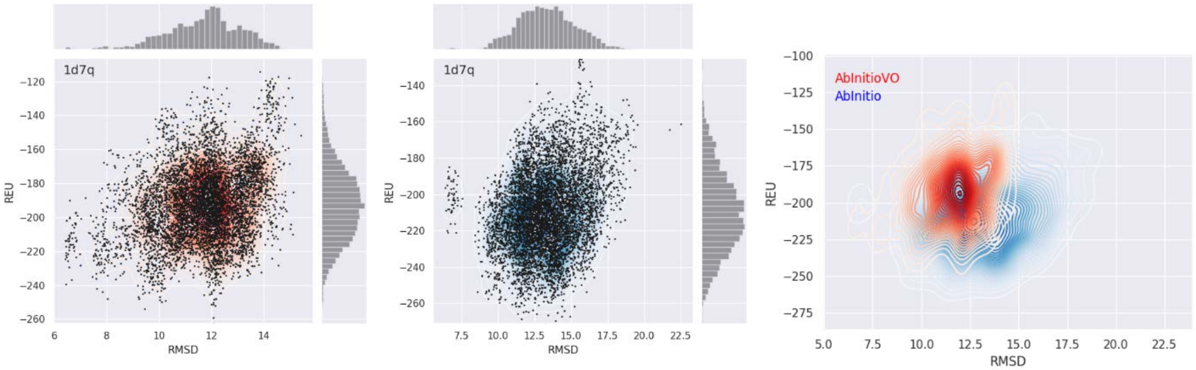


**Figure S118:** Folding Funnel Comparison for 1d7q. Folding funnel from AbInitioVO (left) and AbInitio (middle) simulations plotting Cα RMSD versus Rosetta Energy Units (REU) from each structure compared to the folded domain of the PDB structure. For each folding funnel, histograms of the computed RMSD (top) and REU (right) are shown. KDE plot (right) showing overlay of AbInitiovo (red) and AbInitio (blue) folding funnels.

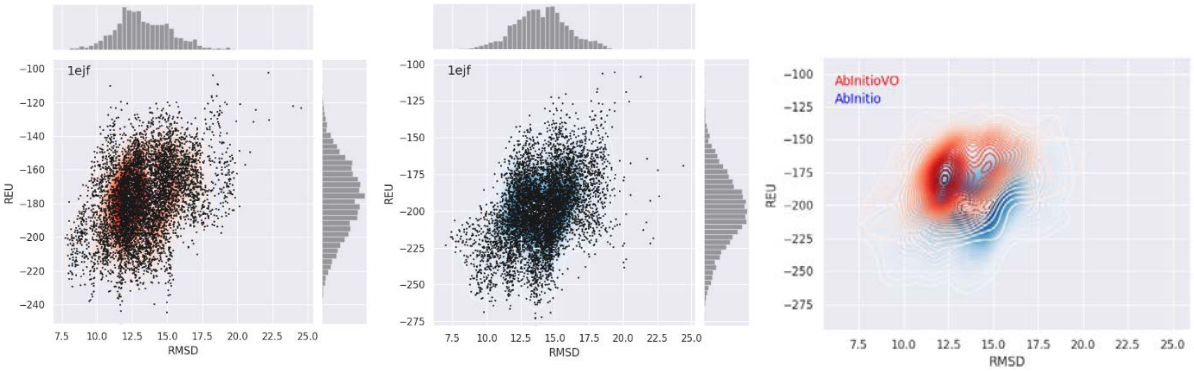


**Figure S119:** Folding Funnel Comparison for 1ejf. Folding funnel from AbInitioVO (left) and AbInitio (middle) simulations plotting Cα RMSD versus Rosetta Energy Units (REU) from each structure compared to the folded domain of the PDB structure. For each folding funnel, histograms of the computed RMSD (top) and REU (right) are shown. KDE plot (right) showing overlay of AbInitiovo (red) and AbInitio (blue) folding funnels.

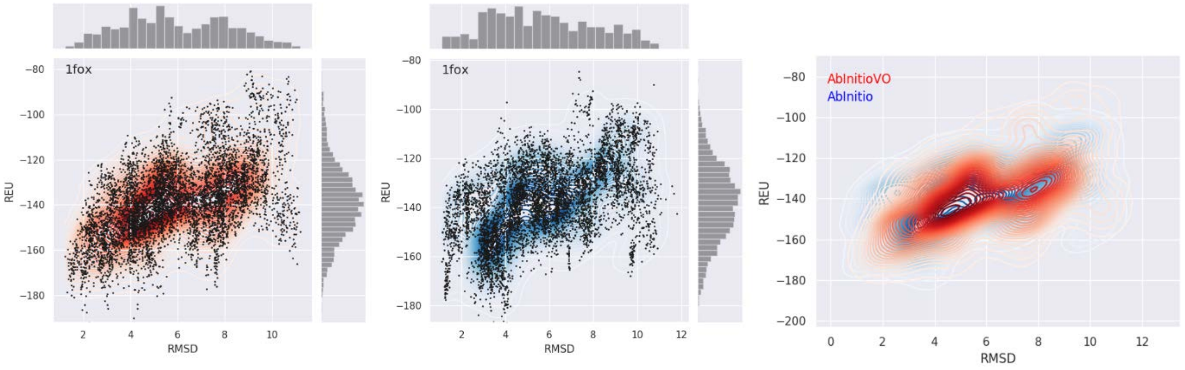


**Figure S120:** Folding Funnel Comparison for 1fox. Folding funnel from AbInitioVO (left) and AbInitio (middle) simulations plotting Cα RMSD versus Rosetta Energy Units (REU) from each structure compared to the folded domain of the PDB structure. For each folding funnel, histograms of the computed RMSD (top) and REU (right) are shown. KDE plot (right) showing overlay of AbInitiovo (red) and AbInitio (blue) folding funnels.

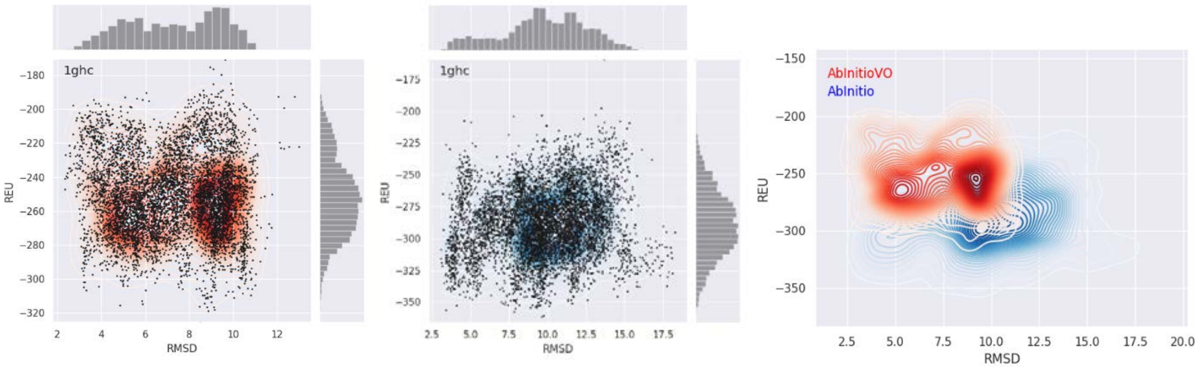


**Figure S121:** Folding Funnel Comparison for 1ghc. Folding funnel from AbInitioVO (left) and AbInitio (middle) simulations plotting Cα RMSD versus Rosetta Energy Units (REU) from each structure compared to the folded domain of the PDB structure. For each folding funnel, histograms of the computed RMSD (top) and REU (right) are shown. KDE plot (right) showing overlay of AbInitiovo (red) and AbInitio (blue) folding funnels.

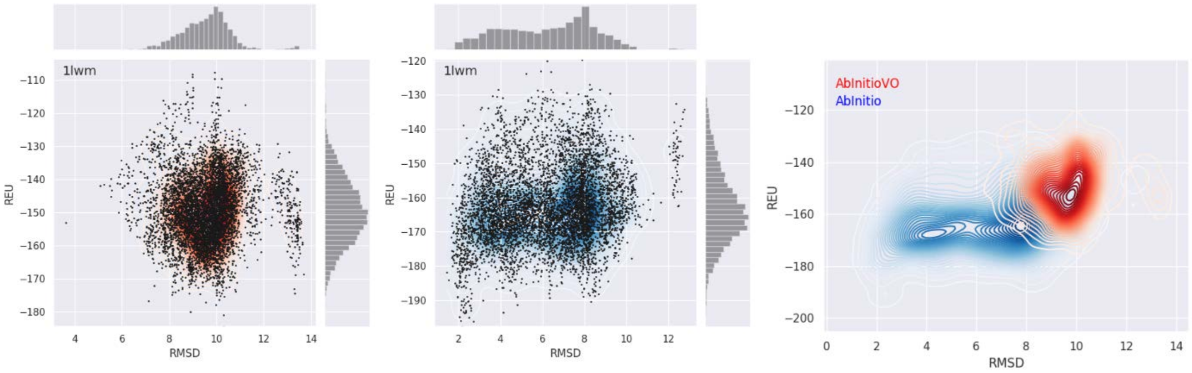


**Figure S122:** Folding Funnel Comparison for 1lwm. Folding funnel from AbInitioVO (left) and AbInitio (middle) simulations plotting Cα RMSD versus Rosetta Energy Units (REU) from each structure compared to the folded domain of the PDB structure. For each folding funnel, histograms of the computed RMSD (top) and REU (right) are shown. KDE plot (right) showing overlay of AbInitiovo (red) and AbInitio (blue) folding funnels.

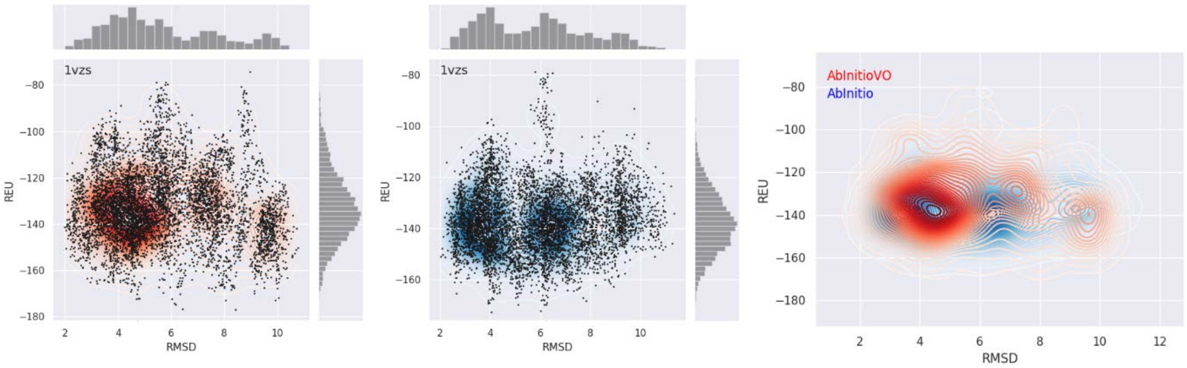


**Figure S123:** Folding Funnel Comparison for 1vzs. Folding funnel from AbInitioVO (left) and AbInitio (middle) simulations plotting Cα RMSD versus Rosetta Energy Units (REU) from each structure compared to the folded domain of the PDB structure. For each folding funnel, histograms of the computed RMSD (top) and REU (right) are shown. KDE plot (right) showing overlay of AbInitiovo (red) and AbInitio (blue) folding funnels.

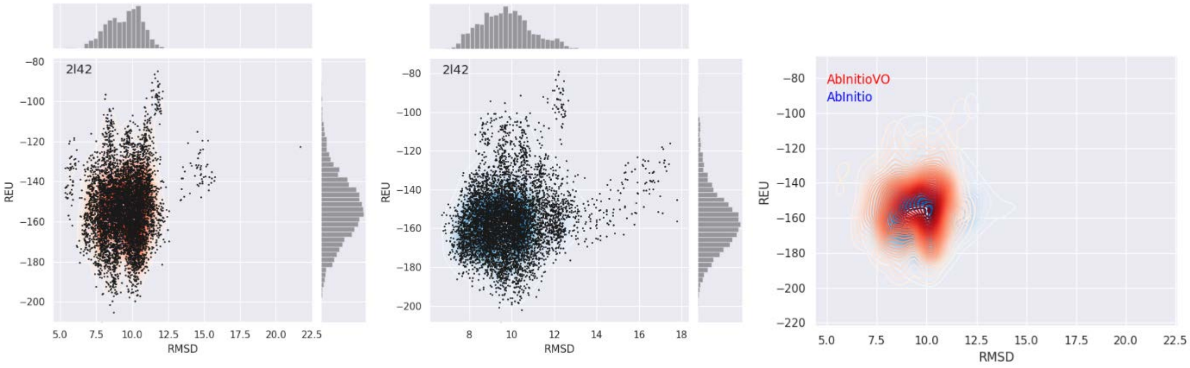


**Figure S124:** Folding Funnel Comparison for 2l42. Folding funnel from AbInitioVO (left) and AbInitio (middle) simulations plotting Cα RMSD versus Rosetta Energy Units (REU) from each structure compared to the folded domain of the PDB structure. For each folding funnel, histograms of the computed RMSD (top) and REU (right) are shown. KDE plot (right) showing overlay of AbInitiovo (red) and AbInitio (blue) folding funnels.

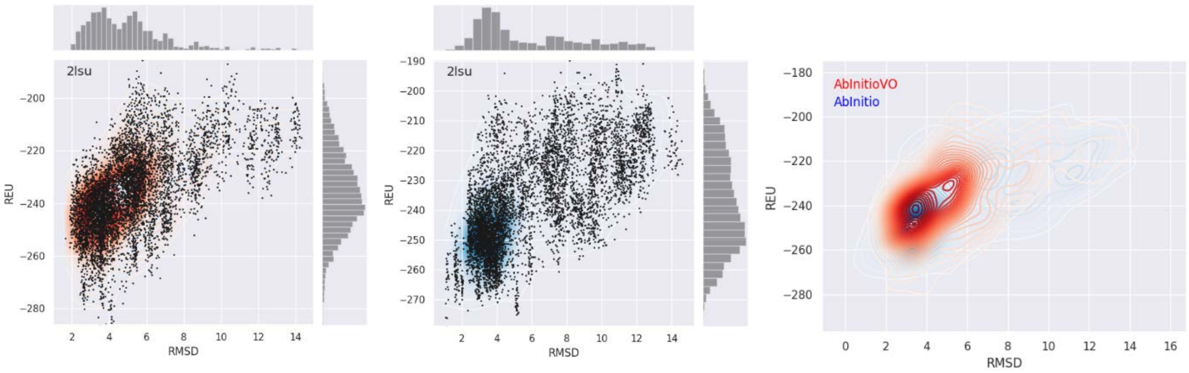


**Figure S125:** Folding Funnel Comparison for 2lsu. Folding funnel from AbInitioVO (left) and AbInitio (middle) simulations plotting Cα RMSD versus Rosetta Energy Units (REU) from each structure compared to the folded domain of the PDB structure. For each folding funnel, histograms of the computed RMSD (top) and REU (right) are shown. KDE plot (right) showing overlay of AbInitiovo (red) and AbInitio (blue) folding funnels.

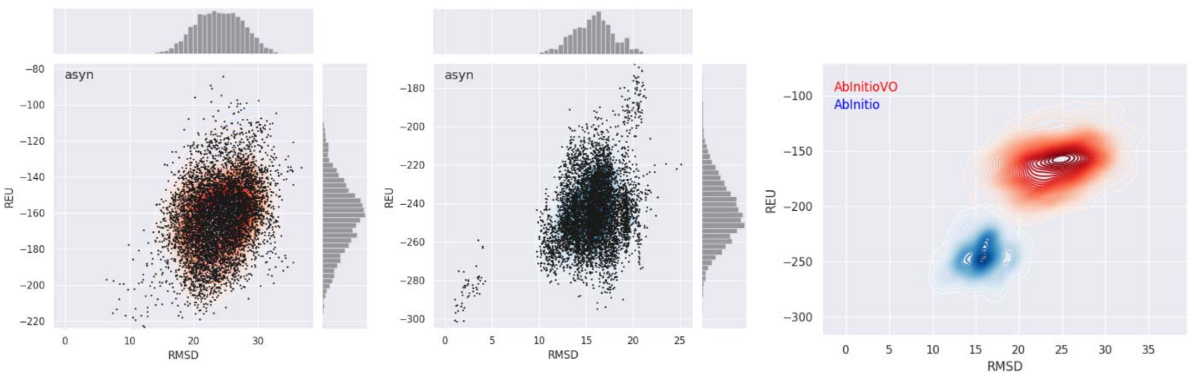


**Figure S126:** Folding Funnel Comparison for asyn. Folding funnel from AbInitioVO (left) and AbInitio (middle) simulations plotting Cα RMSD versus Rosetta Energy Units (REU) from each structure compared to the lowest energy structure. For each folding funnel, histograms of the computed RMSD (top) and REU (right) are shown. KDE plot (right) showing overlay of AbInitiovo (red) and AbInitio (blue) folding funnels.

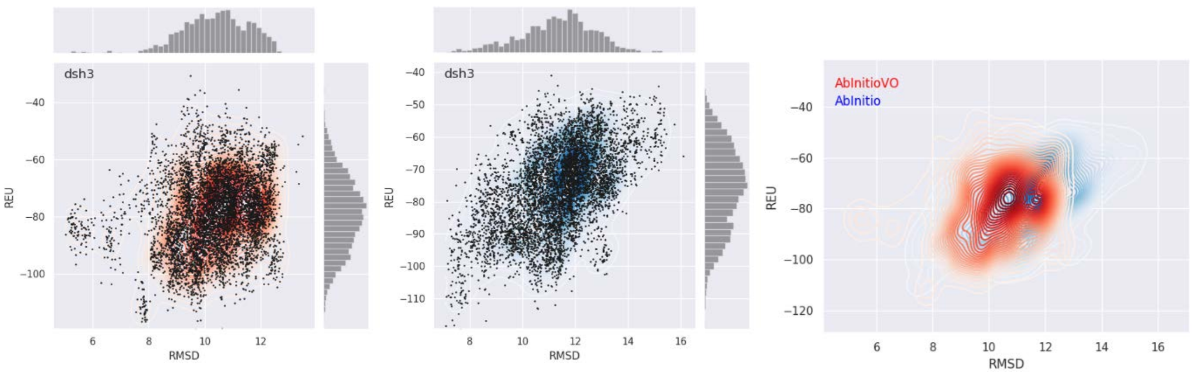


**Figure S127:** Folding Funnel Comparison for dsh3. Folding funnel from AbInitioVO (left) and AbInitio (middle) simulations plotting Cα RMSD versus Rosetta Energy Units (REU) from each structure compared to the PDB structure 2A37. For each folding funnel, histograms of the computed RMSD (top) and REU (right) are shown. KDE plot (right) showing overlay of AbInitiovo (red) and AbInitio (blue) folding funnels.

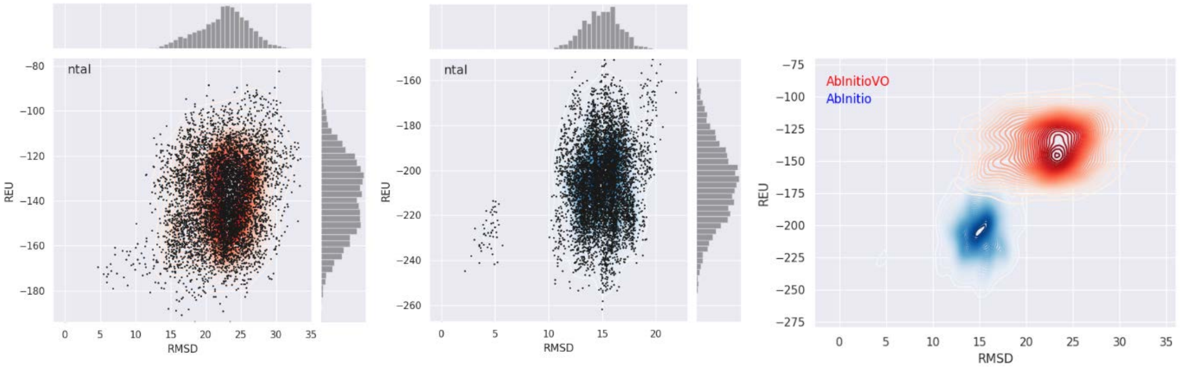


**Figure S128:** Folding Funnel Comparison for ntal. Folding funnel from AbInitioVO (left) and AbInitio (middle) simulations plotting Cα RMSD versus Rosetta Energy Units (REU) from each structure compared to the lowest energy structure. For each folding funnel, histograms of the computed RMSD (top) and REU (right) are shown. KDE plot (right) showing overlay of AbInitiovo (red) and AbInitio (blue) folding funnels.

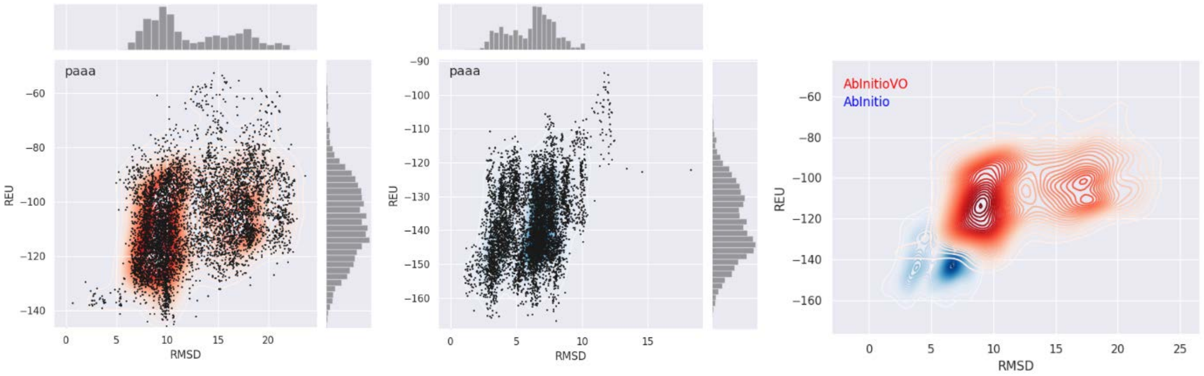


**Figure S129:** Folding Funnel Comparison for paaa. Folding funnel from AbInitioVO (left) and AbInitio (middle) simulations plotting Cα RMSD versus Rosetta Energy Units (REU) from each structure compared to the lowest energy structure. For each folding funnel, histograms of the computed RMSD (top) and REU (right) are shown. KDE plot (right) showing overlay of AbInitiovo (red) and AbInitio (blue) folding funnels.

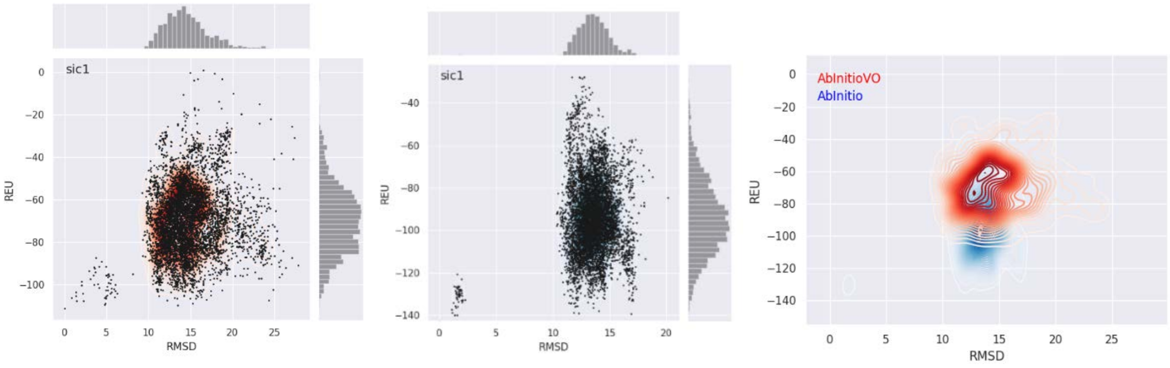


**Figure S130:** Folding Funnel Comparison for sic1. Folding funnel from AbInitioVO (left) and AbInitio (middle) simulations plotting Cα RMSD versus Rosetta Energy Units (REU) from each structure compared to the lowest energy structure. For each folding funnel, histograms of the computed RMSD (top) and REU (right) are shown. KDE plot (right) showing overlay of AbInitiovo (red) and AbInitio (blue) folding funnels.

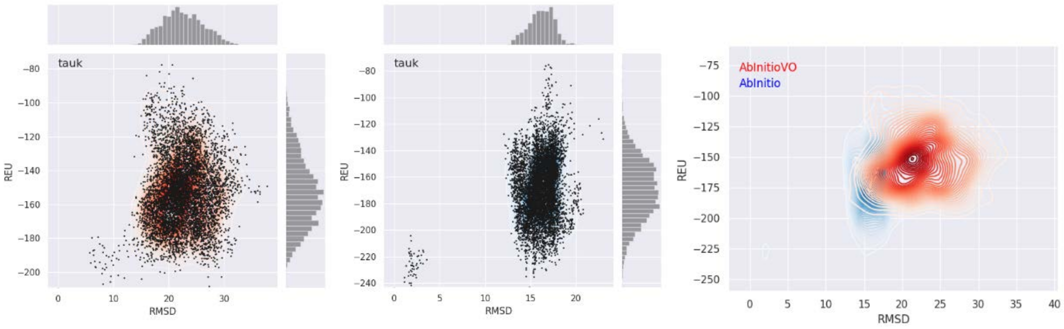


**Figure S131:** Folding Funnel Comparison for tauk. Folding funnel from AbInitioVO (left) and AbInitio (middle) simulations plotting Cα RMSD versus Rosetta Energy Units (REU) from each structure compared to the lowest energy structure. For each folding funnel, histograms of the computed RMSD (top) and REU (right) are shown. KDE plot (right) showing overlay of AbInitiovo (red) and AbInitio (blue) folding funnels.

**Comparisons of Folded Domain Structures of Ordered and Partially-Ordered Proteins**

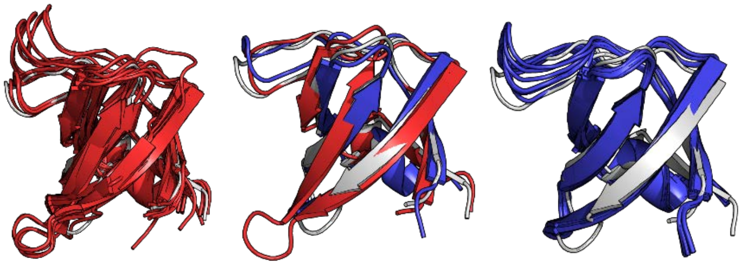

**Figure S132:** 1bk2 Lowest RMSD Structures. 10 lowest RMSD structures from AbInitioVO (left) and AbInitio (right) along with the overlay (center) of PDB 1bk2 (white) and the single lowest RMSD structure from AbInitioVO (red) and AbInitio (blue)

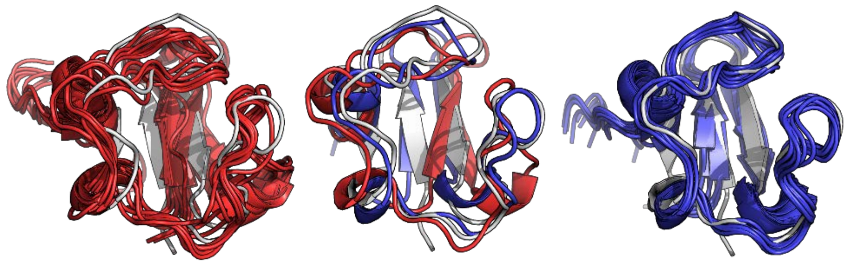

**Figure S133:** 1bq9 Lowest RMSD Structures. 10 lowest RMSD structures from AbInitioVO (left) and AbInitio (right) along with the overlay (center) of PDB 1bq9 (white) and the single lowest RMSD structure from AbInitioVO (red) and AbInitio (blue)

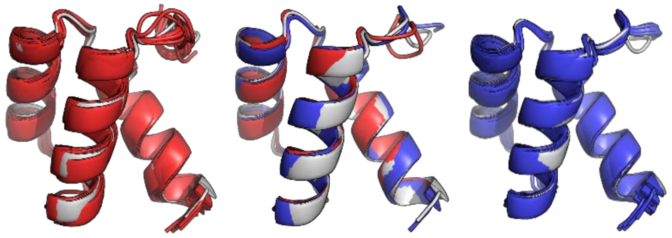

**Figure S134:** 1enh Lowest RMSD Structures. 10 lowest RMSD structures from AbInitioVO (left) and AbInitio (right) along with the overlay (center) of PDB 1enh (white) and the single lowest RMSD structure from AbInitioVO (red) and AbInitio (blue)

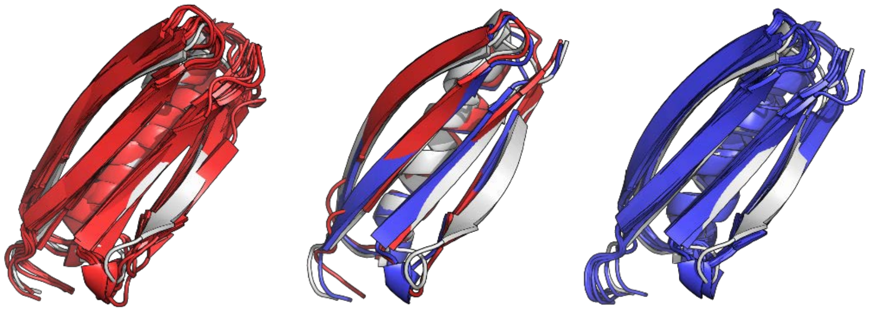

**Figure S135:** 1hz6 Lowest RMSD Structures. 10 lowest RMSD structures from AbInitioVO (left) and AbInitio (right) along with the overlay (center) of PDB 1hz6 (white) and the single lowest RMSD structure from AbInitioVO (red) and AbInitio (blue)

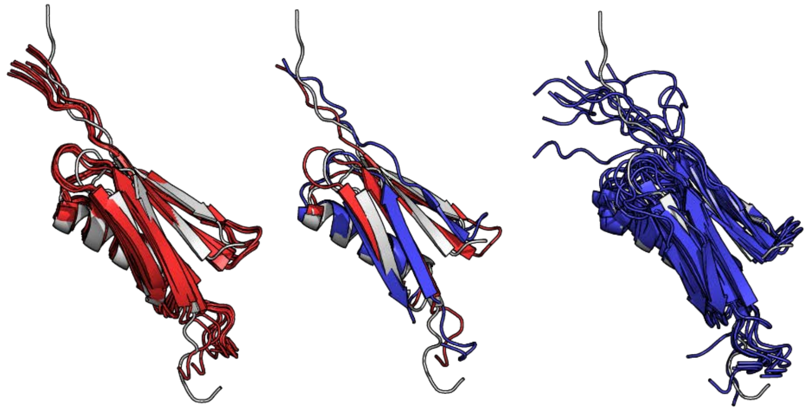

**Figure S136:** 1pgx Lowest RMSD Structures. 10 lowest RMSD structures from AbInitioVO (left) and AbInitio (right) along with the overlay (center) of PDB 1pgx (white) and the single lowest RMSD structure from AbInitioVO (red) and AbInitio (blue)

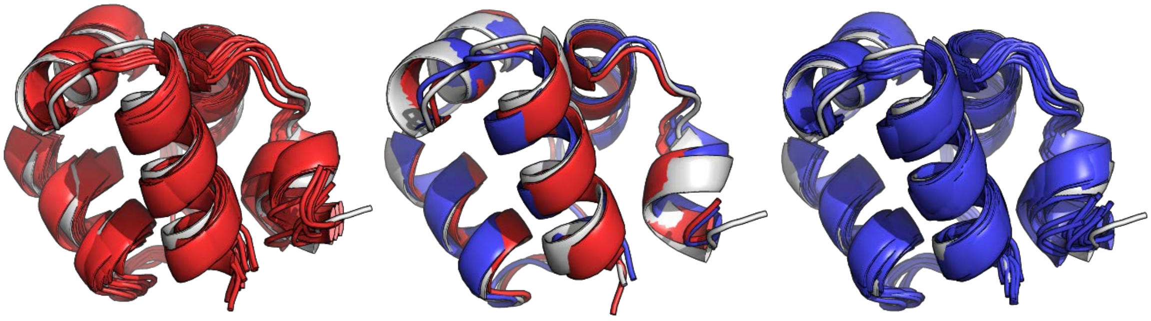

**Figure S137:** 1r69 Lowest RMSD Structures. 10 lowest RMSD structures from AbInitioVO (left) and AbInitio (right) along with the overlay (center) of PDB 1r69 (white) and the single lowest RMSD structure from AbInitioVO (red) and AbInitio (blue)

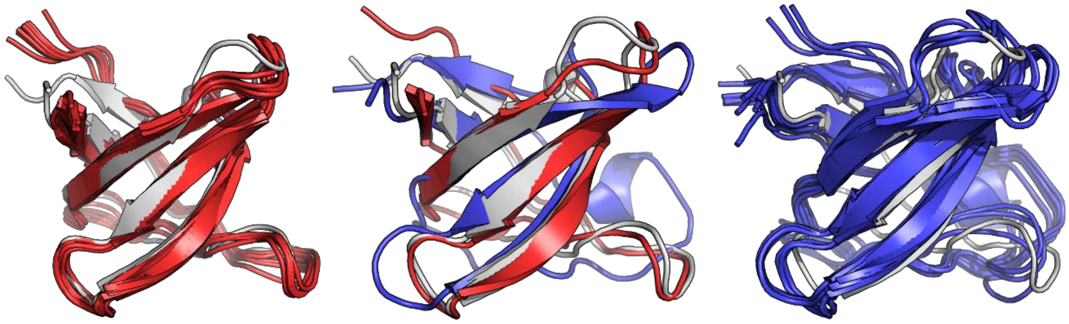

**Figure S138:** 1shf Lowest RMSD Structures. 10 lowest RMSD structures from AbInitioVO (left) and AbInitio (right) along with the overlay (center) of PDB 1shf (white) and the single lowest RMSD structure from AbInitioVO (red) and AbInitio (blue)

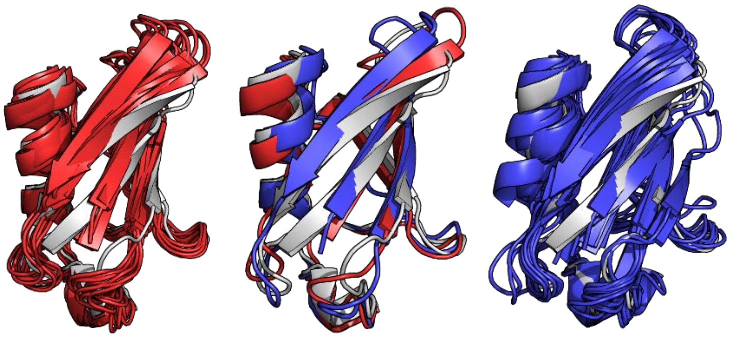

**Figure S139:** 1ubi Lowest RMSD Structures. 10 lowest RMSD structures from AbInitioVO (left) and AbInitio (right) along with the overlay (center) of PDB 1ubi (white) and the single lowest RMSD structure from AbInitioVO (red) and AbInitio (blue)

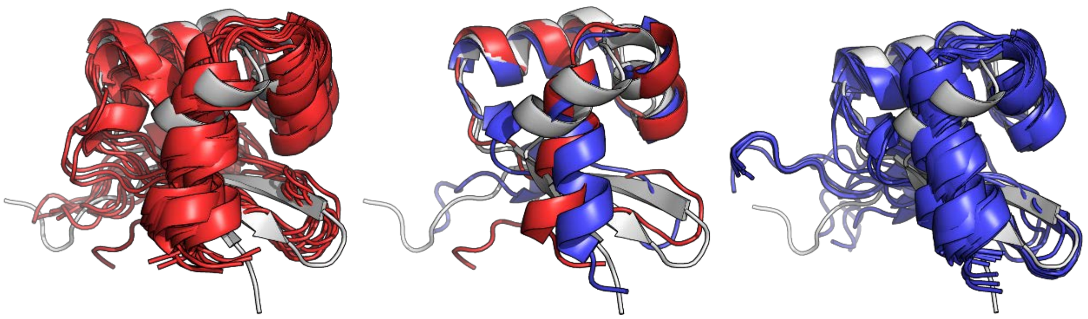

**Figure S140:** 5cro Lowest RMSD Structures. 10 lowest RMSD structures from AbInitioVO (left) and AbInitio (right) along with the overlay (center) of PDB 5cro (white) and the single lowest RMSD structure from AbInitioVO (red) and AbInitio (blue)

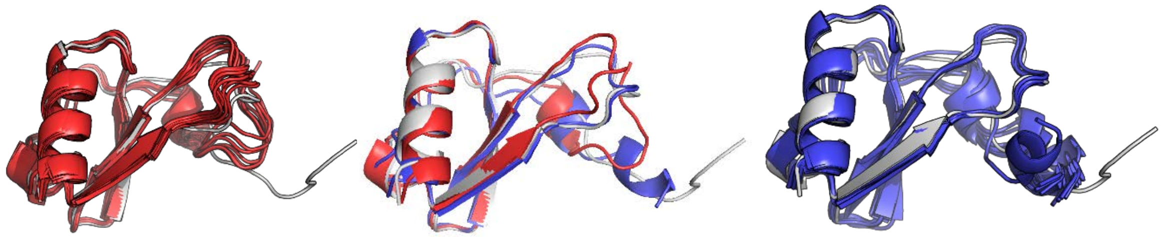

**Figure S141:** 1b3a Lowest RMSD Structures. 10 lowest RMSD structures from AbInitioVO (left) and AbInitio (right) along with the overlay (center) of PDB 1b3a (white) and the single lowest RMSD structure from AbInitioVO (red) and AbInitio (blue)

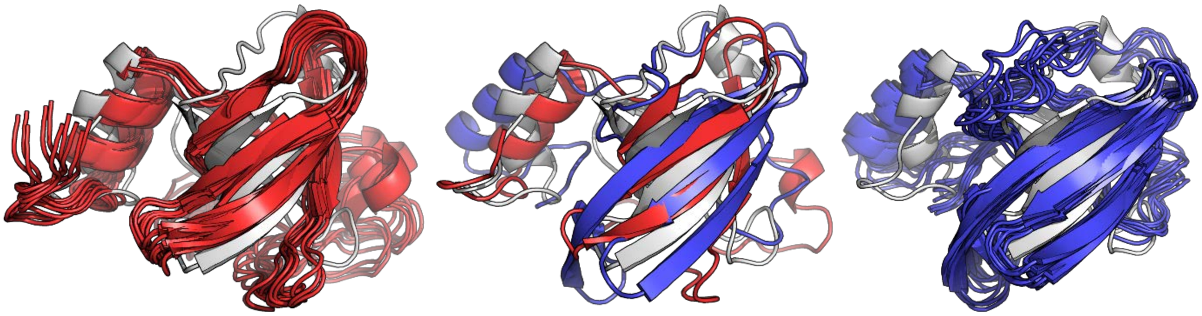

**Figure S142:** 1d7q Lowest RMSD Structures. 10 lowest RMSD structures from AbInitioVO (left) and AbInitio (right) along with the overlay (center) of PDB 1d7q (white) and the single lowest RMSD structure from AbInitioVO (red) and AbInitio (blue)

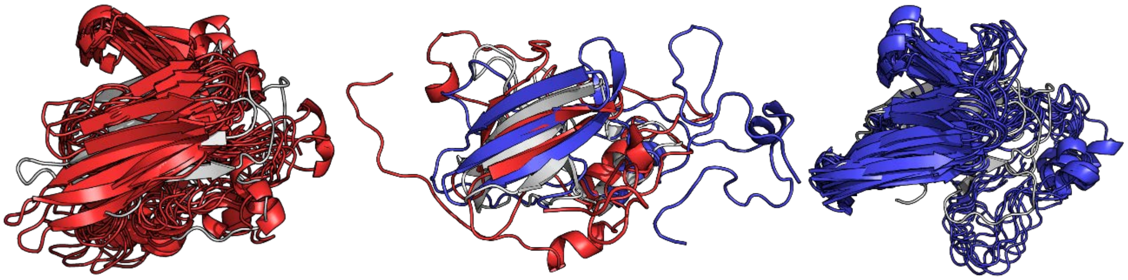

**Figure S143:** 1ejf Lowest RMSD Structures. 10 lowest RMSD structures from AbInitioVO (left) and AbInitio (right) along with the overlay (center) of PDB 1ejf (white) and the single lowest RMSD structure from AbInitioVO (red) and AbInitio (blue)

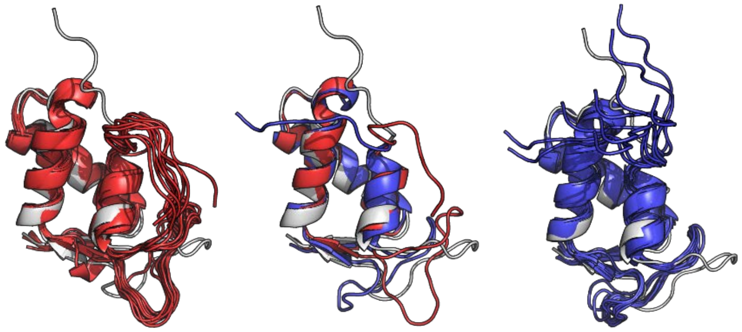

**Figure S144:** 1fox Lowest RMSD Structures. 10 lowest RMSD structures from AbInitioVO (left) and AbInitio (right) along with the overlay (center) of PDB 1fox (white) and the single lowest RMSD structure from AbInitioVO (red) and AbInitio (blue)

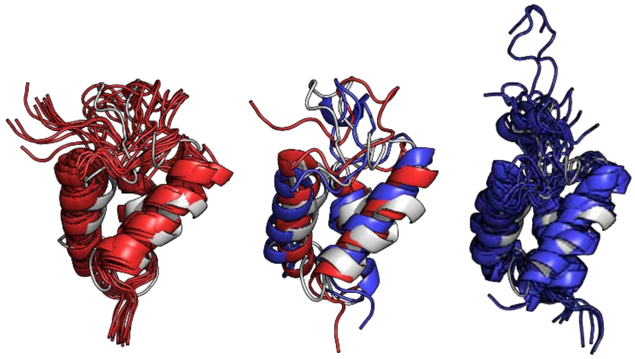

**Figure S145:** 1ghc Lowest RMSD Structures. 10 lowest RMSD structures from AbInitioVO (left) and AbInitio (right) along with the overlay (center) of PDB 1ghc (white) and the single lowest RMSD structure from AbInitioVO (red) and AbInitio (blue)

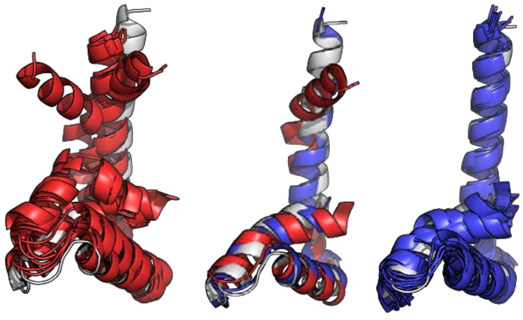

**Figure S146:** 1lwm Lowest RMSD Structures. 10 lowest RMSD structures from AbInitioVO (left) and AbInitio (right) along with the overlay (center) of PDB 1lwm (white) and the single lowest RMSD structure from AbInitioVO (red) and AbInitio (blue)

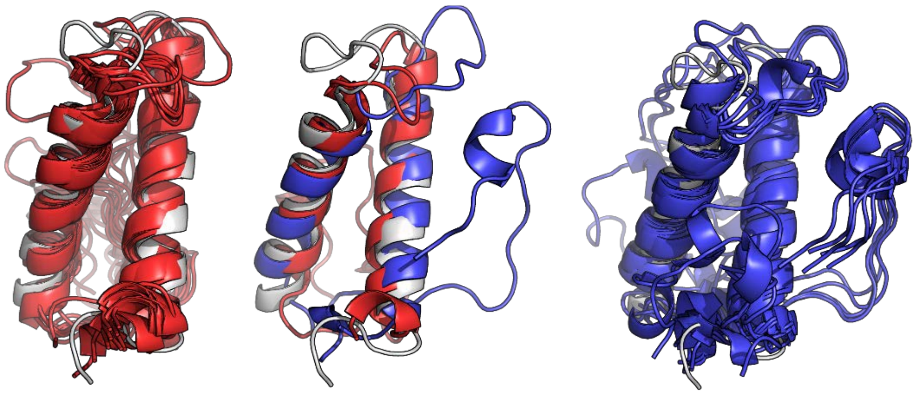

**Figure S147:** 1vzs Lowest RMSD Structures. 10 lowest RMSD structures from AbInitioVO (left) and AbInitio (right) along with the overlay (center) of PDB 1vzs (white) and the single lowest RMSD structure from AbInitioVO (red) and AbInitio (blue)

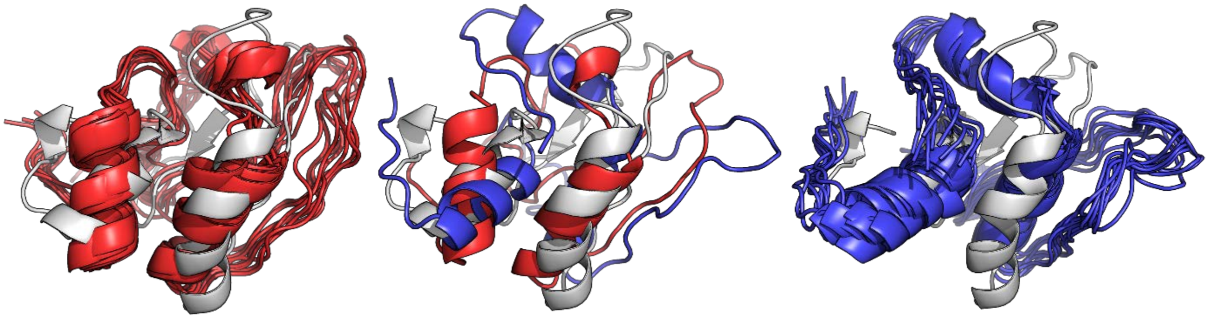

**Figure S148:** 2l42 Lowest RMSD Structures. 10 lowest RMSD structures from AbInitioVO (left) and AbInitio (right) along with the overlay (center) of PDB 2l42 (white) and the single lowest RMSD structure from AbInitioVO (red) and AbInitio (blue)

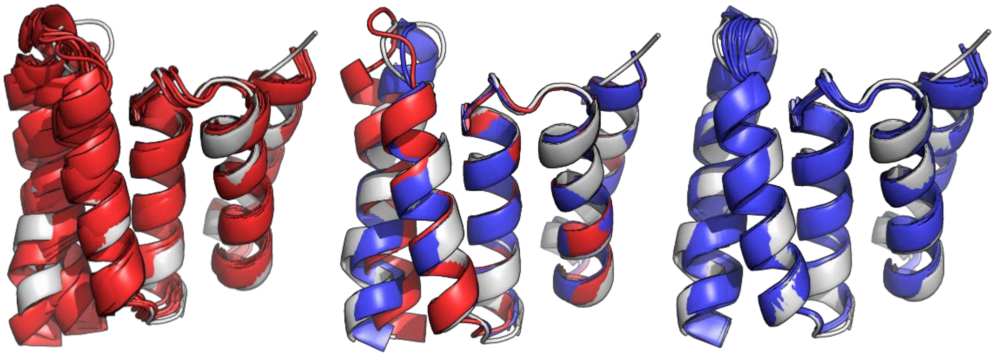

**Figure S149:** 2lsu Lowest RMSD Structures. 10 lowest RMSD structures from AbInitioVO (left) and AbInitio (right) along with the overlay (center) of PDB 2lsu (white) and the single lowest RMSD structure from AbInitioVO (red) and AbInitio (blue)

**Structural Comparisons of Disordered Domains of Partially-Ordered and Disordered Proteins**

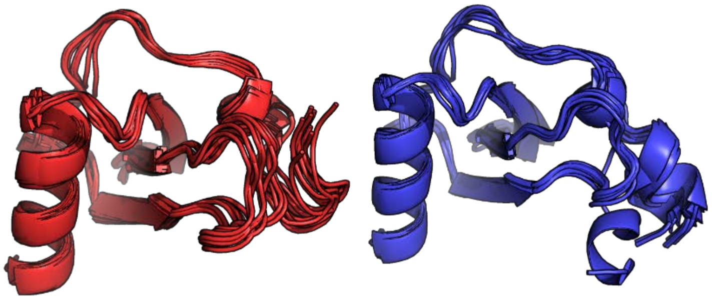

**Figure S150:** 1b3a Lowest RMSD Full Structures. 10 lowest RMSD structures from AbInitioVO (left) and AbInitio (right)

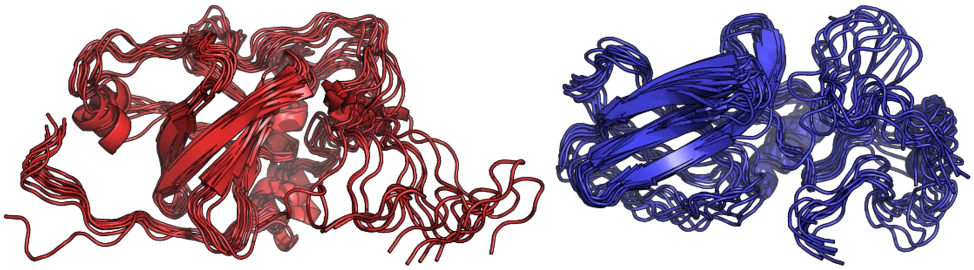

**Figure S151:** 1d7q Lowest RMSD Full Structures. 10 lowest RMSD structures from AbInitioVO (left) and AbInitio (right)

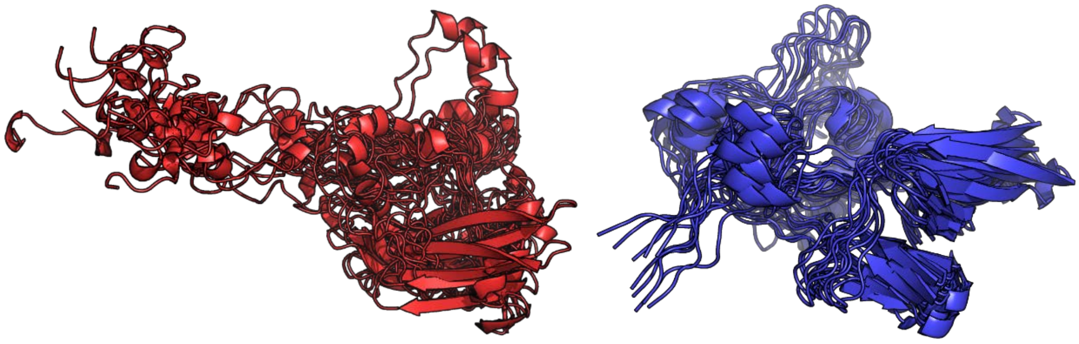

**Figure S152:** 1ejf Lowest RMSD Full Structures. 10 lowest RMSD structures from AbInitioVO (left) and AbInitio (right)

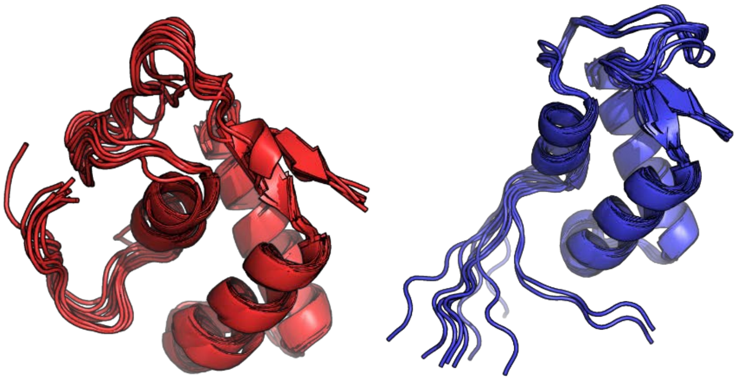

**Figure S153:** 1fox Lowest RMSD Full Structures. 10 lowest RMSD structures from AbInitioVO (left) and AbInitio (right)

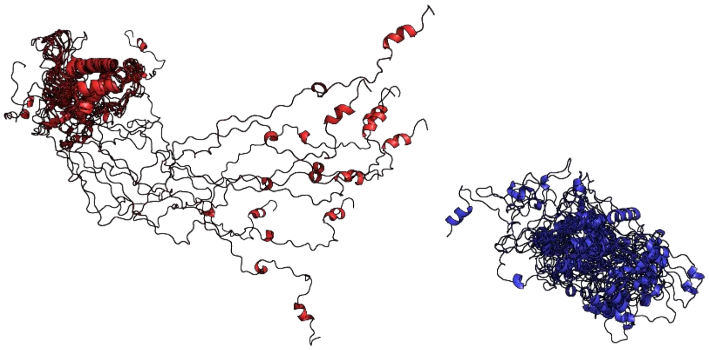

**Figure S154:** 1ghc Lowest RMSD Full Structures. 10 lowest RMSD structures from AbInitioVO (left) and AbInitio (right)

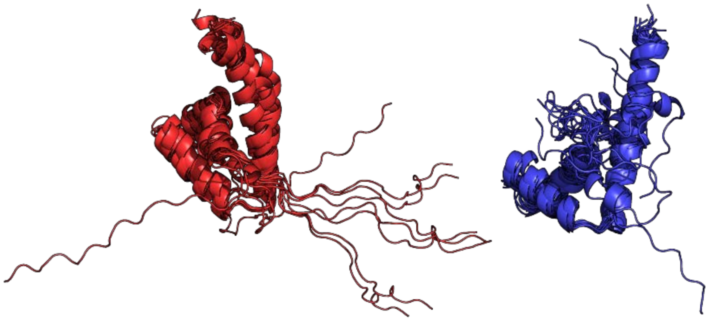

**Figure S155:** 1lwm Lowest RMSD Full Structures. 10 lowest RMSD structures from AbInitioVO (left) and AbInitio (right)

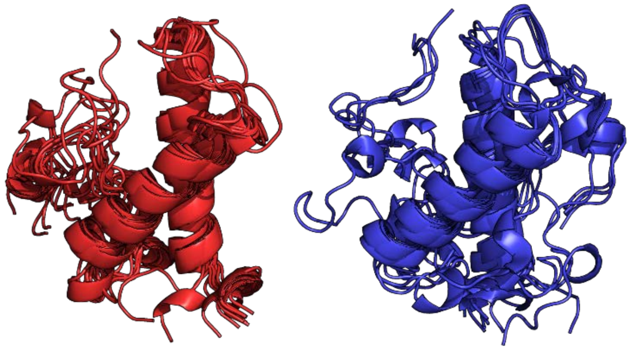

**Figure S156:** 1vzs Lowest RMSD Full Structures. 10 lowest RMSD structures from AbInitioVO (left) and AbInitio (right)

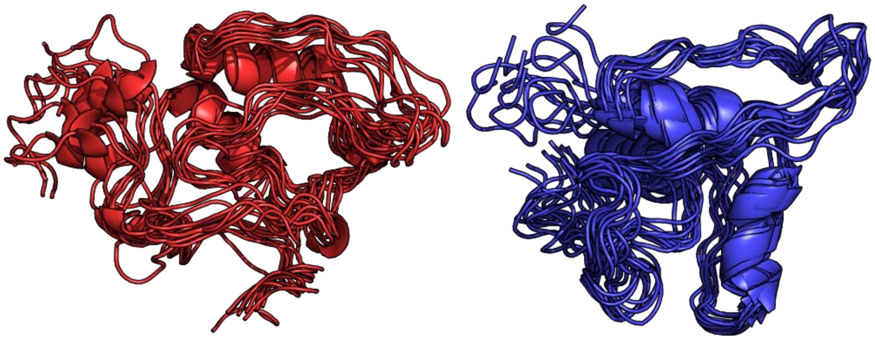

**Figure S157:** 2l42 Lowest RMSD Full Structures. 10 lowest RMSD structures from AbInitioVO (left) and AbInitio (right)

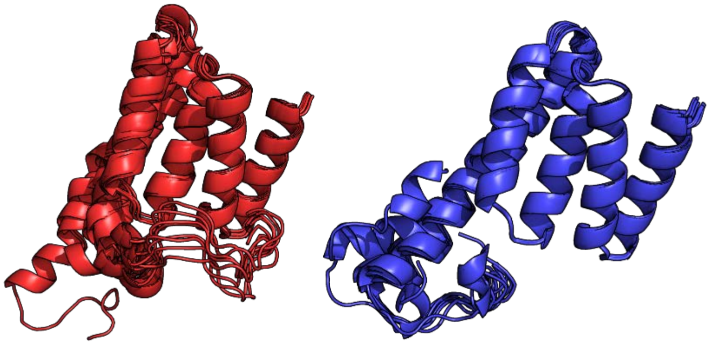

**Figure S158:** 2lsu Lowest RMSD Full Structures. 10 lowest RMSD structures from AbInitioVO (left) and AbInitio (right)

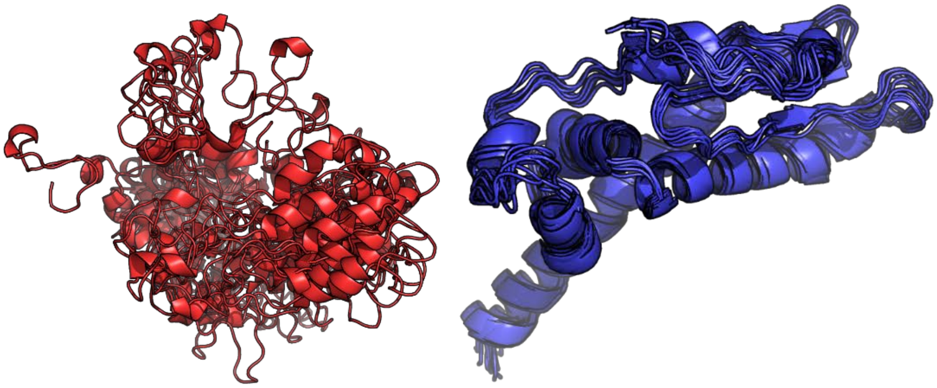

**Figure S159:** Asyn Lowest RMSD Full Structures. 10 lowest RMSD structures from AbInitioVO (left) and AbInitio (right)

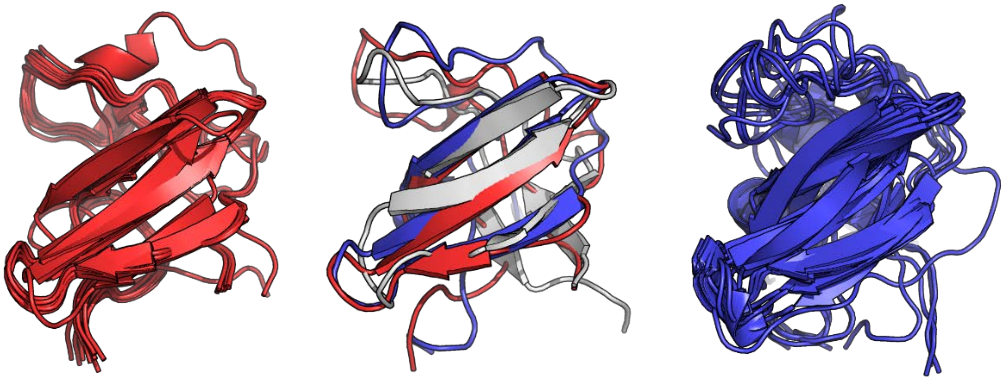

**Figure S160:** Dsh3 Lowest RMSD Full Structures. 10 lowest RMSD structures from AbInitioVO (left) and AbInitio (right) along with the overlay (center) of PDB 2A37 (white) and the single lowest RMSD structure from AbInitioVO (red) and AbInitio (blue)

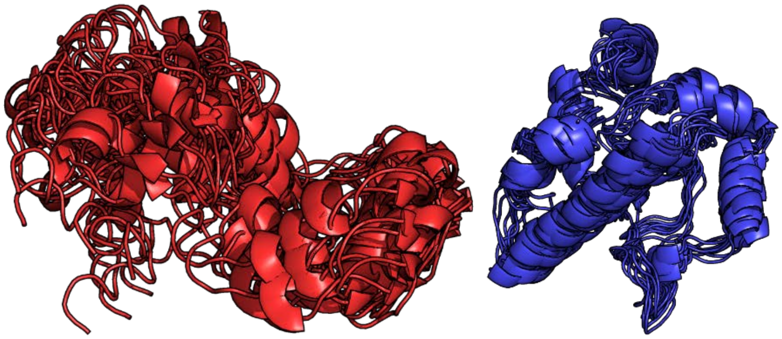

**Figure S161:** Ntal Lowest RMSD Full Structures. 10 lowest RMSD structures from AbInitioVO (left) and AbInitio (right)

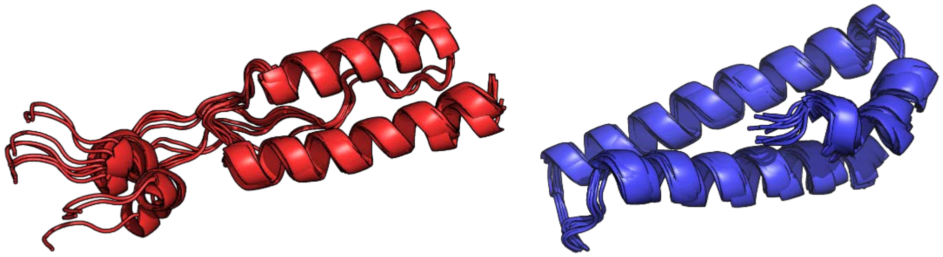

**Figure S162:** Paaa Lowest RMSD Full Structures. 10 lowest RMSD structures from AbInitioVO (left) and AbInitio (right)

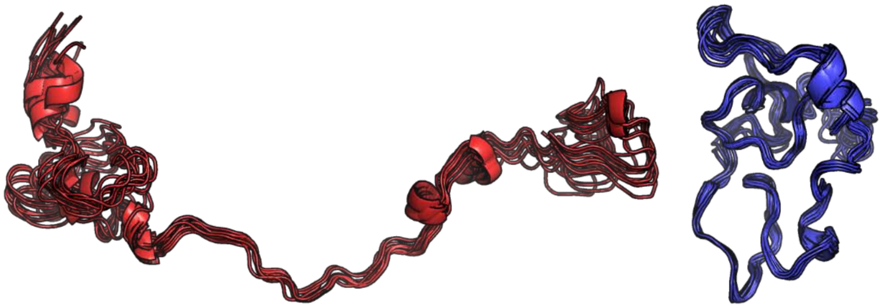

**Figure S163:** Sic1 Lowest RMSD Full Structures. 10 lowest RMSD structures from AbInitioVO (left) and AbInitio (right)

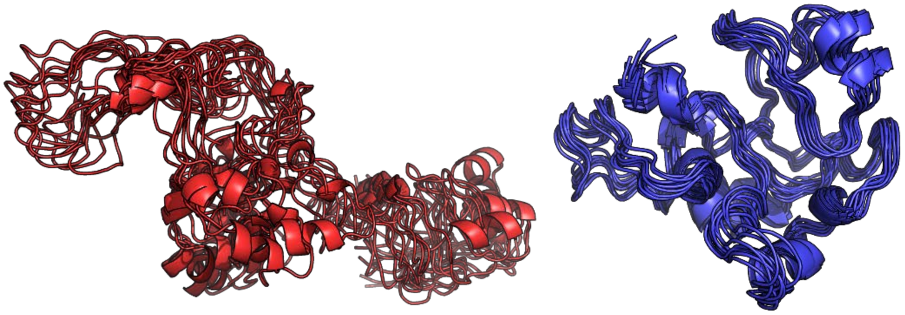

**Figure S164:** Tauk Lowest RMSD Full Structures. 10 lowest RMSD structures from AbInitioVO (left) and AbInitio (right)

### Comparison of FastFloppyTail and FastFloppyTail-Relax Simulations to Experimental Data

#### Summary of Simulated Proteins

**Table S6:** Comparison of FastFloppyTail and FastFloppyTail-Relax Outputs to Experimental Data

| *Protein*<br>Experiment | FastFloppyTail | FastFloppyTail Relaxed | Shaw et. al.[36] |
|---|---|---|---|
| *α-synuclein (αS, asyn)* | *Best / Self* | *Best / Self* | |
| **EFRET** | 0.13 / 0.12 | 0.10 / 0.11 | - |
| **PRE** | 0.20 / 0.20 | 0.16 / 0.16 | 0.17 |
| **DISTANCES** | 11.5 / 10.8 | 6.00 / 5.41 | - |
| **J-COUPLINGS** | | | |
| $^{3}J_{HNH\alpha}$ | 0.58 / 0.58 | 0.67 / 0.67 | 1.11 |
| $^{1}J_{C\alpha H\alpha}$ | 1.21 / 1.21 | 1.14 / 1.13 | - |
| $^{1}J_{NC\alpha}$ | 0.35 / 0.35 | 0.37 / 0.36 | - |
| $^{2}J_{NC\alpha}$ | 0.40 / 0.39 | 0.39 / 0.39 | - |
| $^{3}J_{C'C'}$ | 0.25 / 0.26 | 0.27 / 0.27 | 0.18 |
| **CHEMICAL SHIFTS** | | | |
| **C** | 0.43 / 0.44 | 0.48 / 0.49 | 0.31 |
| **CA** | 0.34 / 0.34 | 0.38 / 0.38 | 0.51 |
| **CB** | 1.02 / 1.02 | 1.03 / 1.03 | 1.04 |
| **N** | 1.54 / 1.56 | 1.62 / 1.63 | 1.46 |
| **NH** | 0.12 / 0.12 | 0.12 / 0.11 | 0.14 |
| **RDC** | 0.41 / 0.40 | 0.41 / 0.41 | 0.41 |
| **$R_g$ (33.0 ± 3.0, 26.6 ± 0.5)** | 39.3 / 38.6 | 32.3 / 31.3 | 36.76 |
| *Ntail (ntal)* | *Best / Self* | *Best / Self* | |
| **CHEMICAL SHIFTS** | | | |
| **CA** | 0.57 / 0.40 | 0.58 / 0.40 | 0.44 |
| **HN** | 0.13 / 0.11 | 0.13 / 0.11 | 0.12 |
| **N** | 0.90 / 0.83 | 0.94 / 0.87 | 0.89 |
| **C** | 0.66 / 0.49 | 0.69 / 0.52 | 0.41 |
| **CB** | 0.46 / 0.36 | 0.48 / 0.38 | 0.37 |
| **RDC** | 0.48 / 0.39 | 0.48 / 0.40 | 0.95 |
| **$R_g$ (27.5 ± 0.70)** | 37.5 / 36.9 | 32.9 / 32.3 | 26.65 |
| *Sic1 (sic1)* | *Best / Self* | *Best / Self* | |
| **CHEMICAL SHIFTS** | | | |
| **CA** | 0.71 / 0.71 | 0.68 / 0.67 | 0.77 |
| **HN** | 0.15 / 0.15 | 0.15 / 0.15 | 0.12 |
| **HA** | 0.24 / 0.24 | 0.25 / 0.25 | 0.25 |
| **N** | 1.60 / 1.57 | 1.69 / 1.65 | 1.16 |
| **CB** | 0.51 / 0.52 | 0.55 / 0.56 | 0.52 |
| **RDC** | 0.44 / 0.44 | 0.44 / 0.44 | 0.43 |
| **$R_g$ (31.2 ± 0.8)** | 28.4 / 28.5 | 24.9 / 25.3 | 24.42 |

| *Tau K18 (tauk)* | ***Best / Self*** | ***Best / Self*** | |
|---|---|---|---|
| **CHEMICAL SHIFTS** | | | |
| **C** | 0.41 / 0.47 | 0.45 / 0.51 | - |
| **CB** | 0.34 / 0.37 | 0.34 / 0.37 | - |
| **CA** | 0.33 / 0.42 | 0.38 / 0.47 | - |
| **N** | 1.67 / 1.69 | 1.68 / 1.69 | - |
| **HN** | 0.13 / 0.14 | 0.13 / 0.15 | - |
| **RDC** | 0.39 / 0.41 | 0.40 / 0.41 | - |
| **$R_g$ (33.4 ± 7.2)** | 40.1 / 39.7 | 34.7 / 34.5 | - |
| *DrkN Sh3 (dsh3)* | ***Best / Self*** | ***Best / Self*** | |
| **CHEMICAL SHIFTS** | | | |
| **C** | 0.98 / 0.77 | 0.93 / 0.74 | 0.65 |
| **CB** | 0.48 / 0.51 | 0.47 / 0.51 | 0.50 |
| **CA** | 0.74 / 0.68 | 0.70 / 0.71 | 0.50 |
| **HN** | 0.29 / 0.22 | 0.28 / 0.22 | 0.23 |
| **HA** | 0.12 / 0.13 | 0.11 / 0.14 | 0.10 |
| **RDC** | 0.38 / 0.39 | 0.37 / 0.39 | 0.91 |
| **$^3J_{HNH\alpha}$** | 0.89 / 1.01 | 0.86 / 1.13 | 1.08 |
| **PRE** | 0.33 / 0.33 | 0.30 / 0.30 | 0.3 |
| **$R_g$ (18.3)** | 20.6 / 23.6 | 19.2 / 22.2 | 19.46 |
| *PaaA2 (paaa)* | ***Best / Self*** | ***Best / Self*** | |
| **CHEMICAL SHIFTS** | | | |
| **C** | 0.57 / 0.71 | 0.55 / 0.68 | 0.66 |
| **CB** | 0.46 / 0.55 | 0.44 / 0.53 | 0.37 |
| **CA** | 0.79 / 0.99 | 0.79 / 0.97 | 0.64 |
| **N** | 1.38 / 1.44 | 1.39 / 1.55 | 0.99 |
| **HN** | 0.18 / 0.21 | 0.18 / 0.20 | 0.17 |
| **HA** | 0.10 / 0.12 | 0.10 / 0.12 | 0.10 |
| **RDC** | 0.35 / 0.36 | 0.35 / 0.36 | 0.76 |
| **$R_g$ (22.4 ± 4.0)** | 23.6 / 23.5 | 22.4 / 22.4 | 21.39 |

*All values reported above are computed at RMSD values except the reported Rg values which represent the average Rg (Å) and RDC values which are computed as Q-values, as described in Zweckstetter *et. al.* [12].

### Comparisons of αS Data

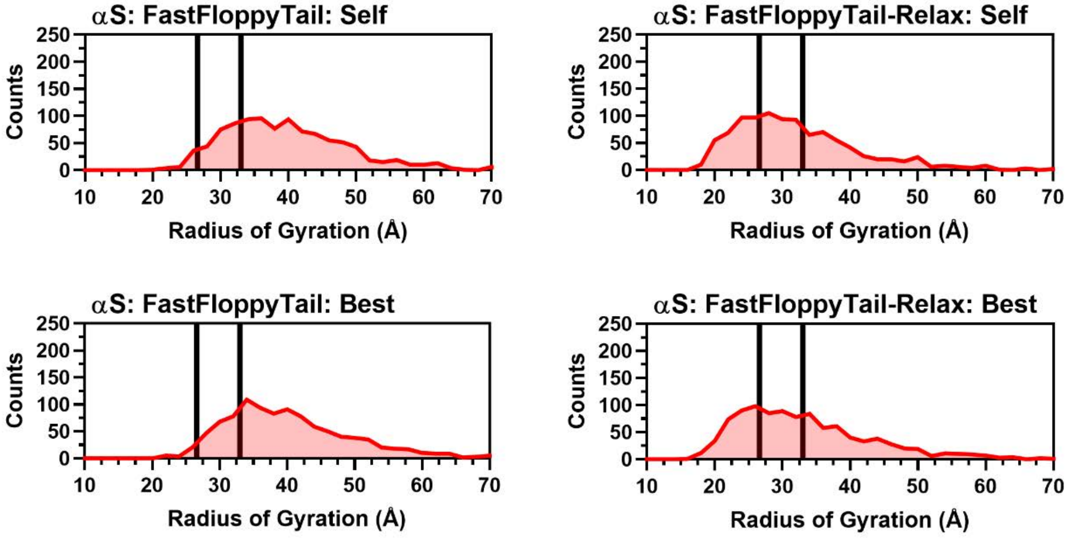


**Figure S165:** Histograms of Radii of Gyration for αS. Histograms of the radius of gyration from self (Top Left) and best (Top Right) reweighting in FastFloppyTail and self (Bottom Left) and best (Bottom Right) reweighting in FastFloppyTail-Relax compared to experimental values from SAXS [19] and NMR [20] (black).

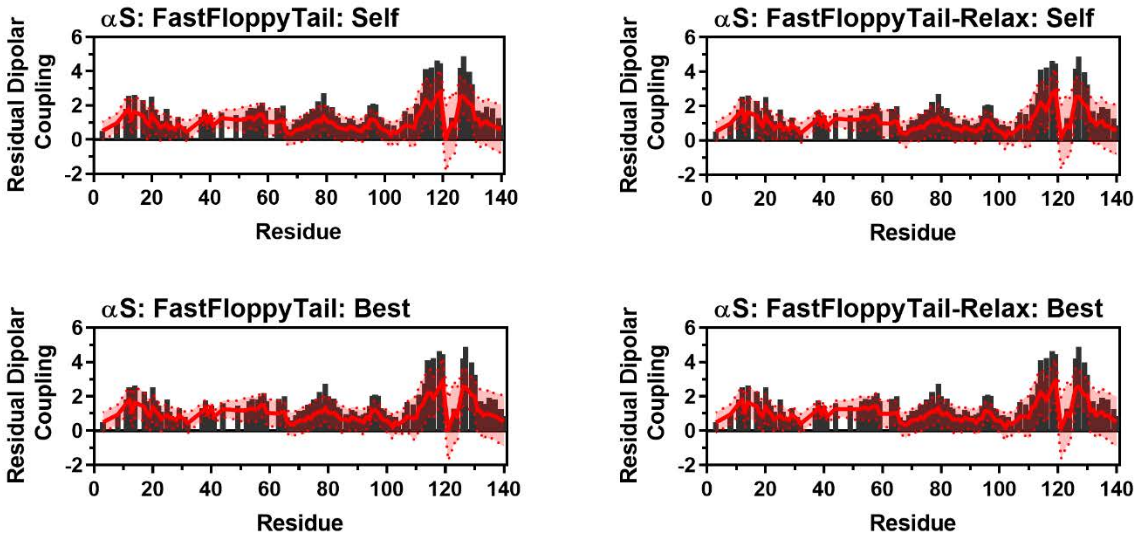


**Figure S166:** Comparison of Simulated and Experimental Residual Dipolar Coupling Data. Simulated RDC values (red line) from self (Top Left) and best (Top Right) reweighting in FastFloppyTail and self (Bottom Left) and best (Bottom Right) reweighting in FastFloppyTail-Relax overlayed on experimental data (black bars) from Bertoncini *et al.* [25].

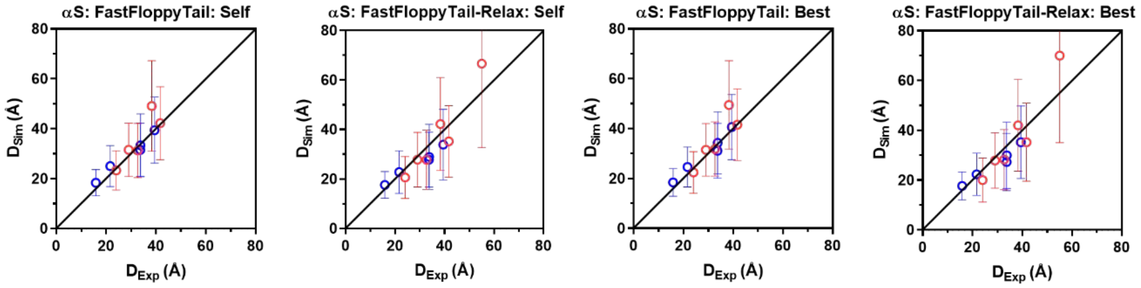


**Figure S167:** Comparison of Simulated Distance with Experimental Distances. Simulated distances for self-reweighting in FastFloppyTail (Left) and FastFloppyTail-Relax (Middle Left) and best-reweighting in FastFloppyTail (Middle Right) and FastFloppyTail-Relax (Far Right) with data from Lee *et al*. [24] (Red) and Grupi *et al*. [23] (Blue).

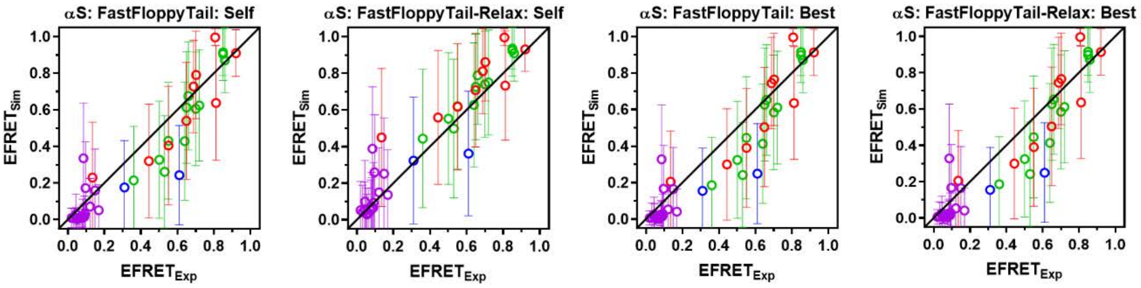


**Figure S168:** Comparison of Simulated $E_{FRET}$ with Experimental $E_{FRET}$. Simulated $E_{FRET}$s for self-reweighting in FastFloppyTail (Left) and FastFloppyTail-Relax (Middle Left) and best-reweighting in FastFloppyTail (Middle Right) and FastFloppyTail-Relax (Far Right) with data from Ferrie *et al*. Cnf-Trp [21] (Purple) and Fam-Raz [21] (Red) Pairs, Ferrie *et al*. Mcm-Acd pair [22] (Blue), and Nath *et al*. [1] (Green).

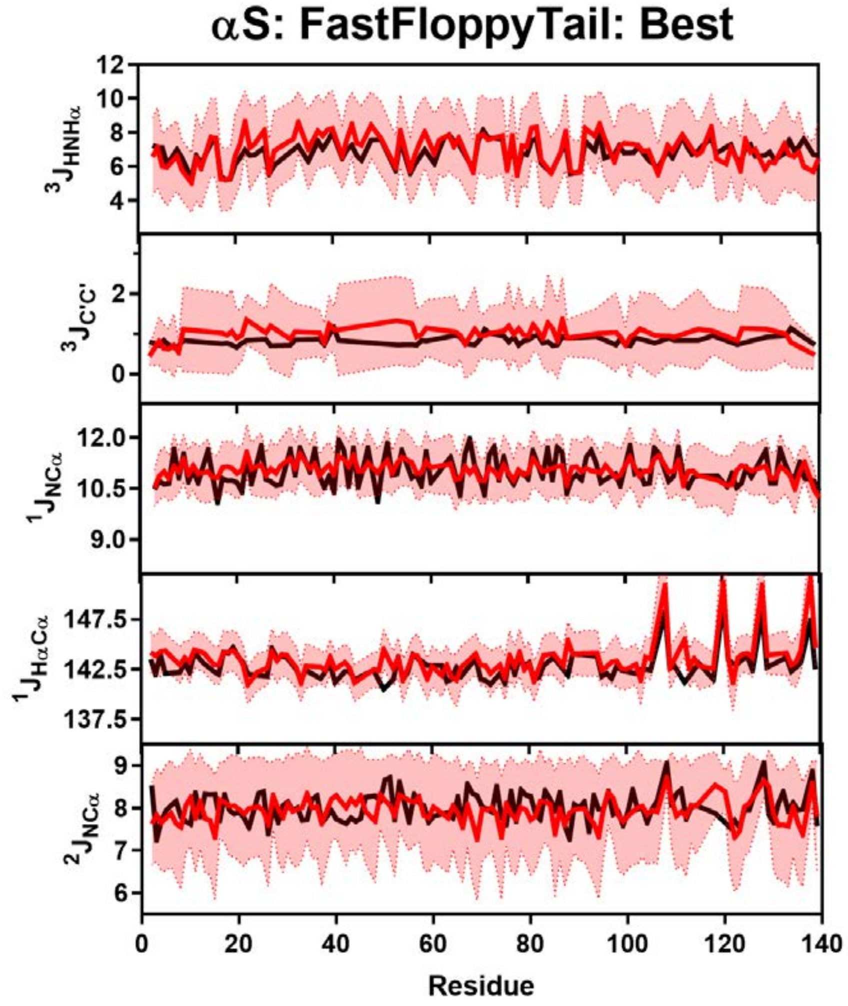


**Figure S169:** Comparison of Simulated and Experimental J-Coupling Data. Simulated J-Coupling values from the FastFloppyTail simulations with best-reweighting (red) overlayed on experimental data (black) of $^3J_{HNH\alpha}$ (Row 1), $^3J_{CC}$ (Row 2), $^1J_{H\alpha C\alpha}$ (Row 3), $^1J_{NC\alpha}$ (Row 4) and $^2J_{NC\alpha}$ (Row 5) from Mantsyzov *et al.* and Lee *et. al*. [13,18].

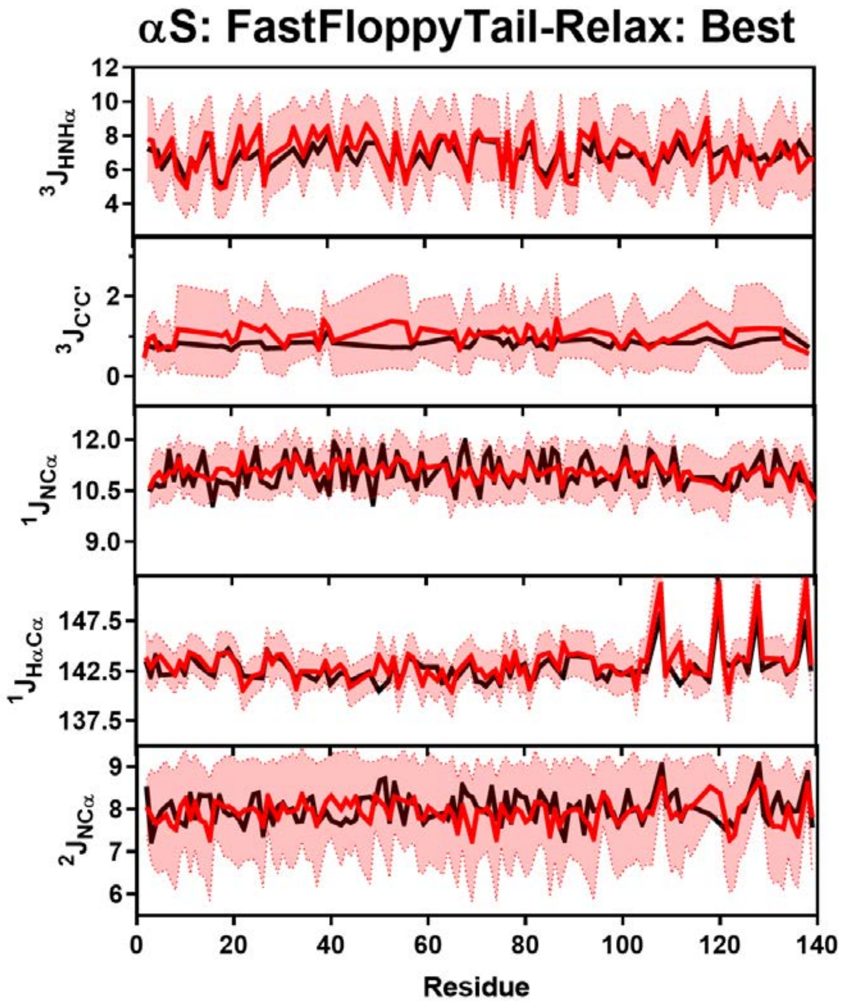


**Figure S170:** Comparison of Simulated and Experimental J-Coupling Data. Simulated J-Coupling values from the FastFloppyTail-Relax simulations with best-reweighting (red) overlayed on experimental data (black) of $^3J_{HNH\alpha}$ (Row 1), $^3J_{CC}$ (Row 2), $^1J_{H\alpha C\alpha}$ (Row 3), $^1J_{NC\alpha}$ (Row 4) and $^2J_{NC\alpha}$ (Row 5) from Mantsyzov *et al.* and Lee *et. al*. [13,18].

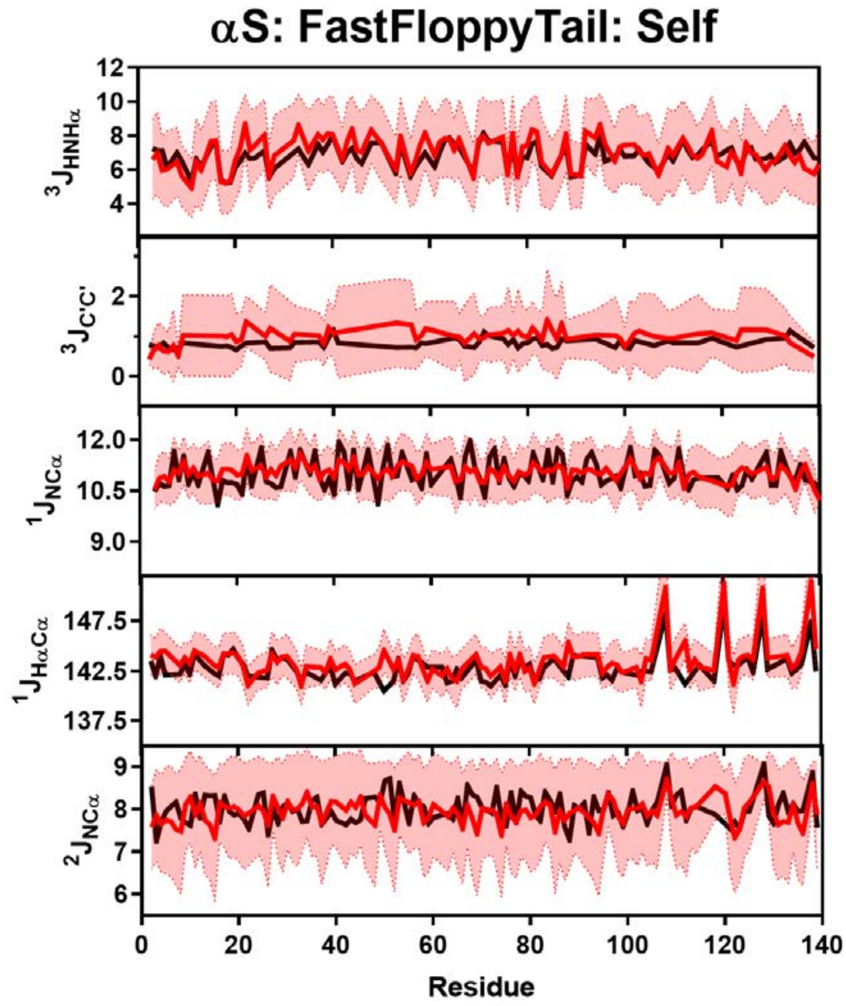


**Figure S171:** Comparison of Simulated and Experimental J-Coupling Data. Simulated J-Coupling values from the FastFloppyTail simulations with self-reweighting (red) overlayed on experimental data (black) of $^{3}J_{HNH\alpha}$ (Row 1), $^{3}J_{CC}$ (Row 2), $^{1}J_{H\alpha C\alpha}$ (Row 3), $^{1}J_{NC\alpha}$ (Row 4) and $^{2}J_{NC\alpha}$ (Row 5) from Mantsyzov *et al.* and Lee *et. al*. [13,18].

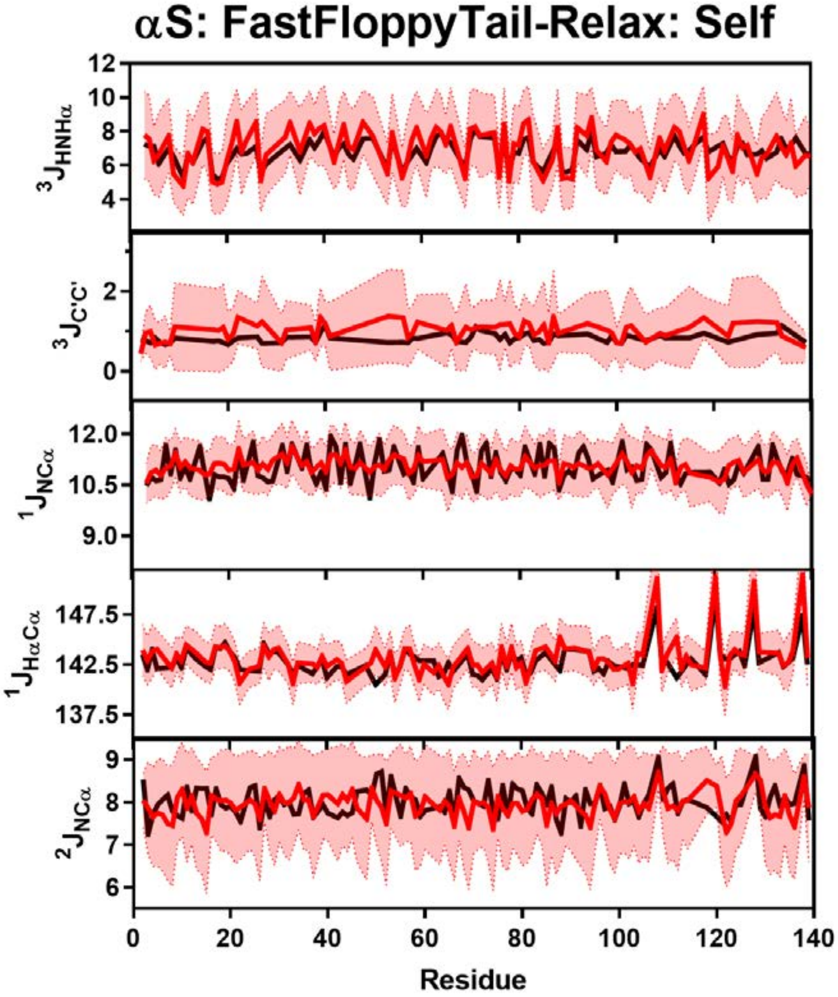


**Figure S172:** Comparison of Simulated and Experimental J-Coupling Data. Simulated J-Coupling values from the FastFloppyTail-Relax simulations with self-reweighting (red) overlayed on experimental data (black) of $^{3}J_{HNH\alpha}$ (Row 1), $^{3}J_{CC}$ (Row 2), $^{1}J_{H\alpha C\alpha}$ (Row 3), $^{1}J_{NC\alpha}$ (Row 4) and $^{2}J_{NC\alpha}$ (Row 5) from Mantsyzov *et al.* and Lee *et. al*. [13,18].

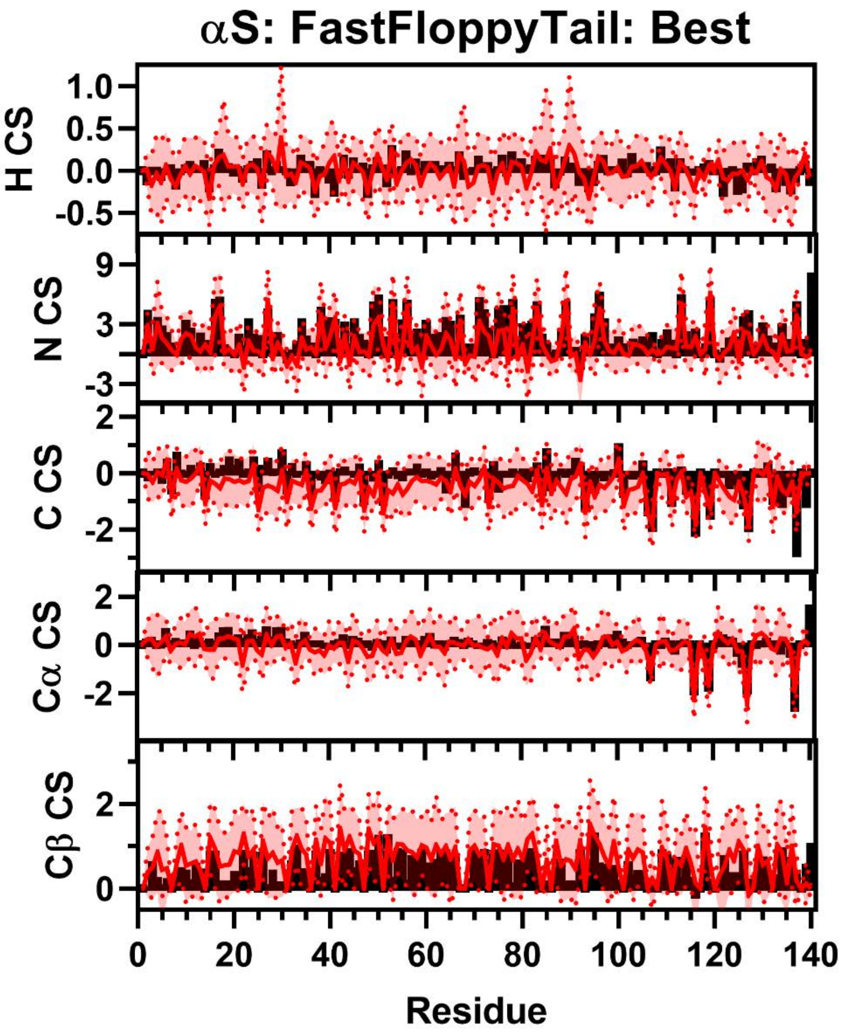


**Figure S173:** Comparison of Simulated and Experimental NMR Secondary Chemical Shift Data for αS. Simulated Chemical Shift values from the FastFloppyTail ensemble using best-reweighted fragment selection (red) overlayed on experimental data (black bars) of N (Row 1), H (Row 2), C (Row 3), Cα (Row 4), and Cβ (Row 5) chemical shifts from Sung *et al.* [8]. Neighbor corrected random coil chemical shift values from SPARTA+ have been subtracted from both simulated and experimental chemical shifts to generate secondary chemical shift data [11].

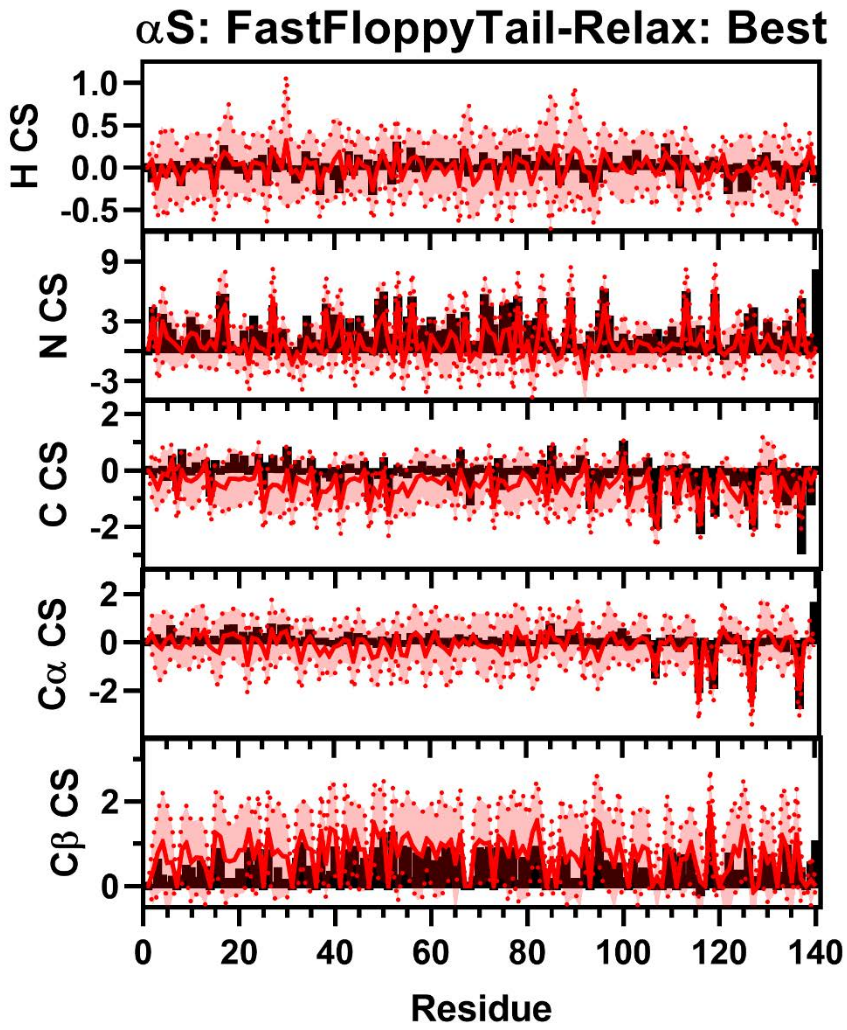


**Figure S174:** Comparison of Simulated and Experimental NMR Secondary Chemical Shift Data for αS. Simulated Chemical Shift values from the FastFloppyTail-Relax ensemble using best-reweighted fragment selection (red) overlayed on experimental data (black bars) of N (Row 1), H (Row 2), C (Row 3), Cα (Row 4), and Cβ (Row 5) chemical shifts from Sung *et al.* [8] Neighbor corrected random coil chemical shift values from SPARTA+ have been subtracted from both simulated and experimental chemical shifts to generate secondary chemical shift data [11].

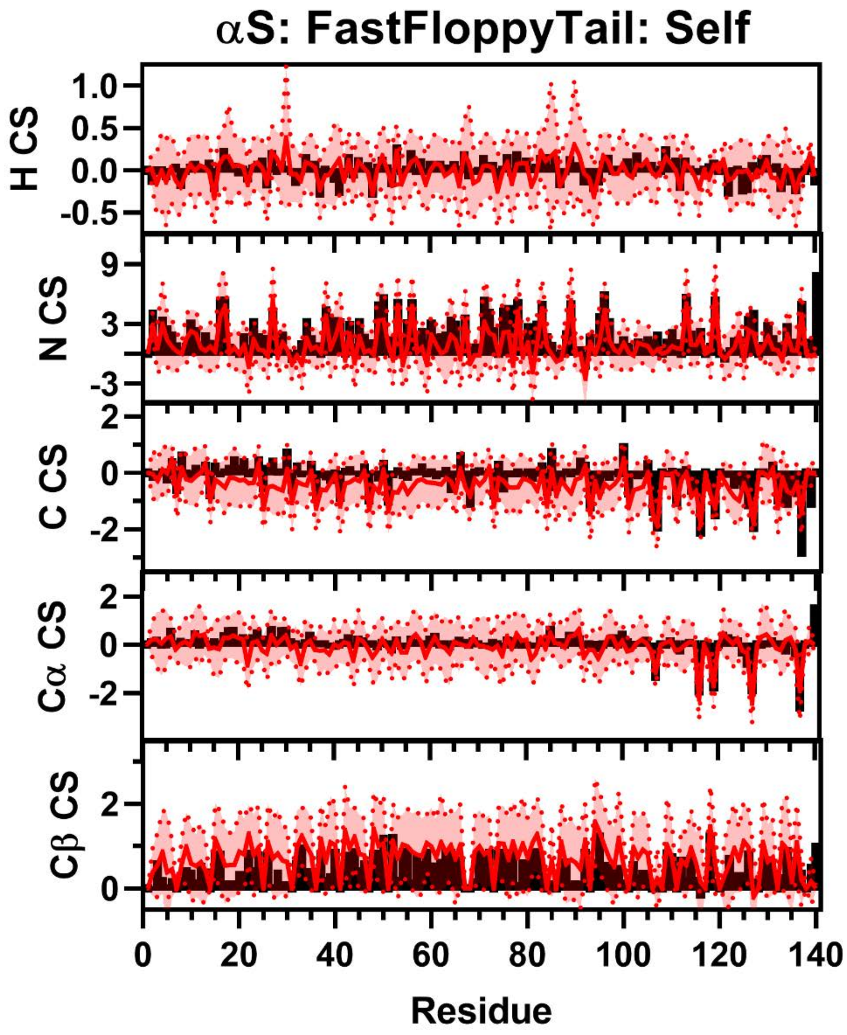


**Figure S175:** Comparison of Simulated and Experimental NMR Secondary Chemical Shift Data for αS. Simulated Chemical Shift values from the FastFloppyTail ensemble using self-reweighted fragment selection (red) overlayed on experimental data (black bars) of N (Row 1), H (Row 2), C (Row 3), Cα (Row 4), and Cβ (Row 5) chemical shifts from Sung *et al*. [8] Neighbor corrected random coil chemical shift values from SPARTA+ have been subtracted from both simulated and experimental chemical shifts to generate secondary chemical shift data [11].

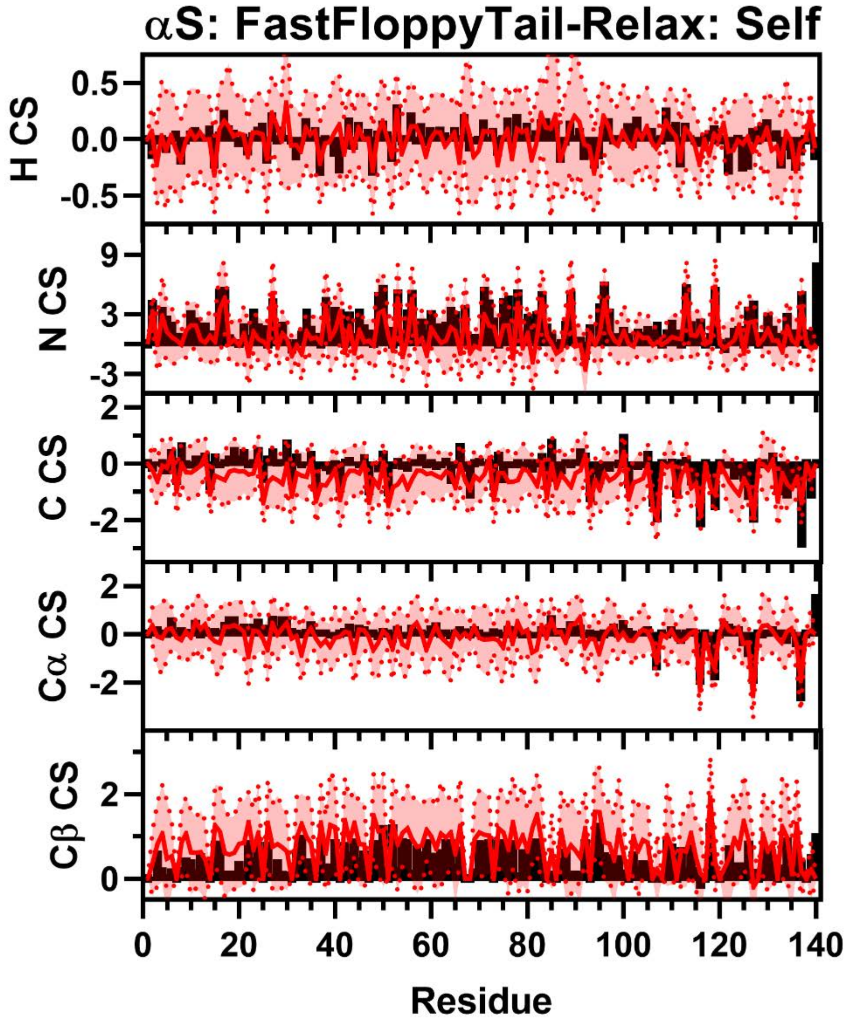


**Figure S176:** Comparison of Simulated and Experimental NMR Secondary Chemical Shift Data for αS. Simulated Chemical Shift values from the FastFloppyTail-Relax ensemble using self-reweighted fragment selection (red) overlayed on experimental data (black bars) of N (Row 1), H (Row 2), C (Row 3), Cα (Row 4), and Cβ (Row 5) chemical shifts from Sung *et al.* [8] Neighbor corrected random coil chemical shift values from Sparta+ have been subtracted from both simulated and experimental chemical shifts to generate secondary chemical shift data [11].

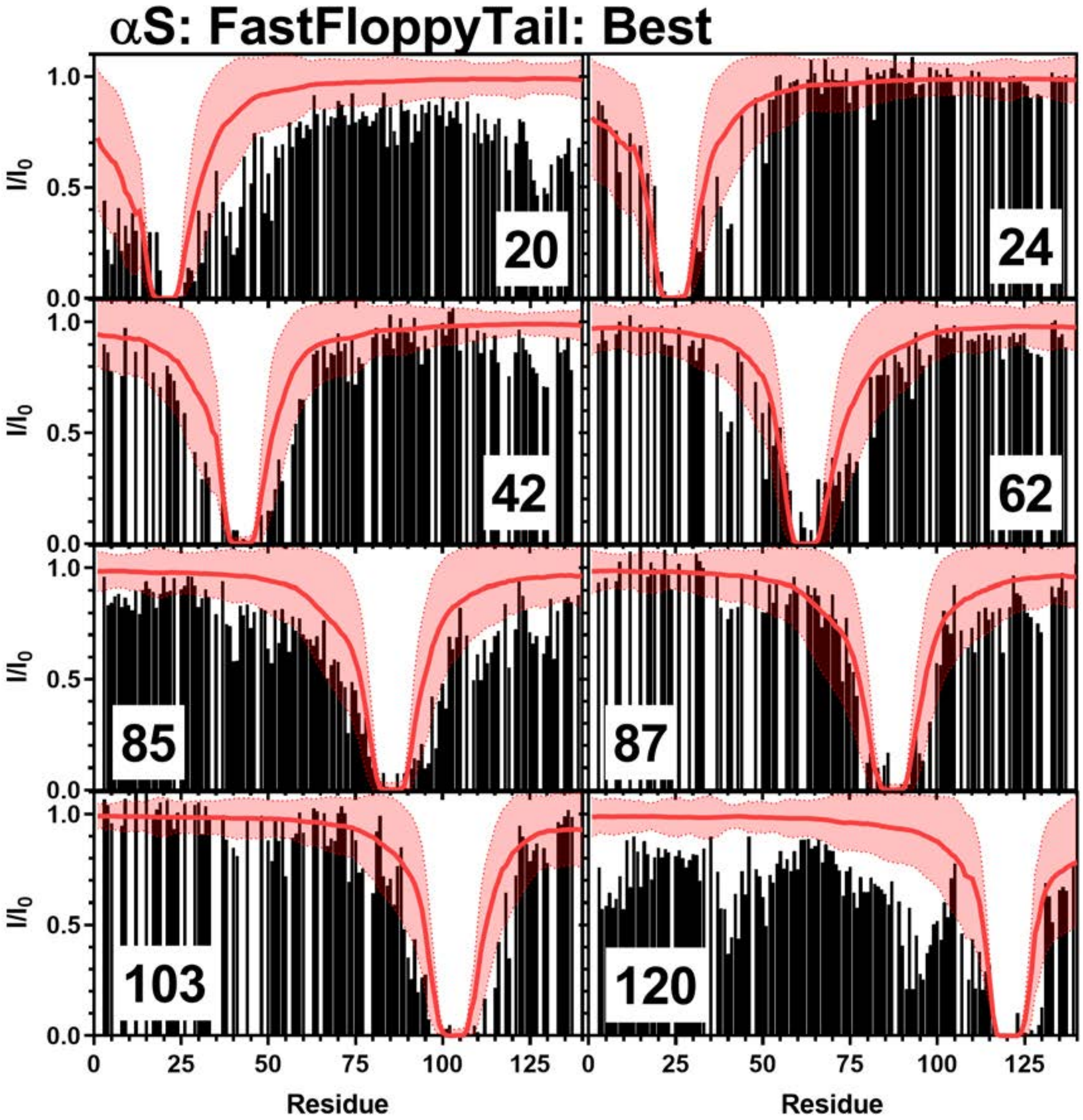


**Figure S177:** Comparison of Simulated PRE Values and Experimental PRE Values: Simulated PRE values from FastFloppyTail using best-reweighting (red line) overlayed on top of experimental data (black bars) from positions 20 (Top Left), 24 (Top Right), 42 (Upper Middle Left), 62 (Upper Middle Right), 85 (Lower Middle Left), 87 (Lower Middle Right), 103 (Bottom Left), 120 (Bottom Right). Experimental data for positions 20, 85, and 120 are from Sung *et al.* [8] and data for positions 24, 42, 62, 87, and 103 are from Dedmon *et al.* [9].

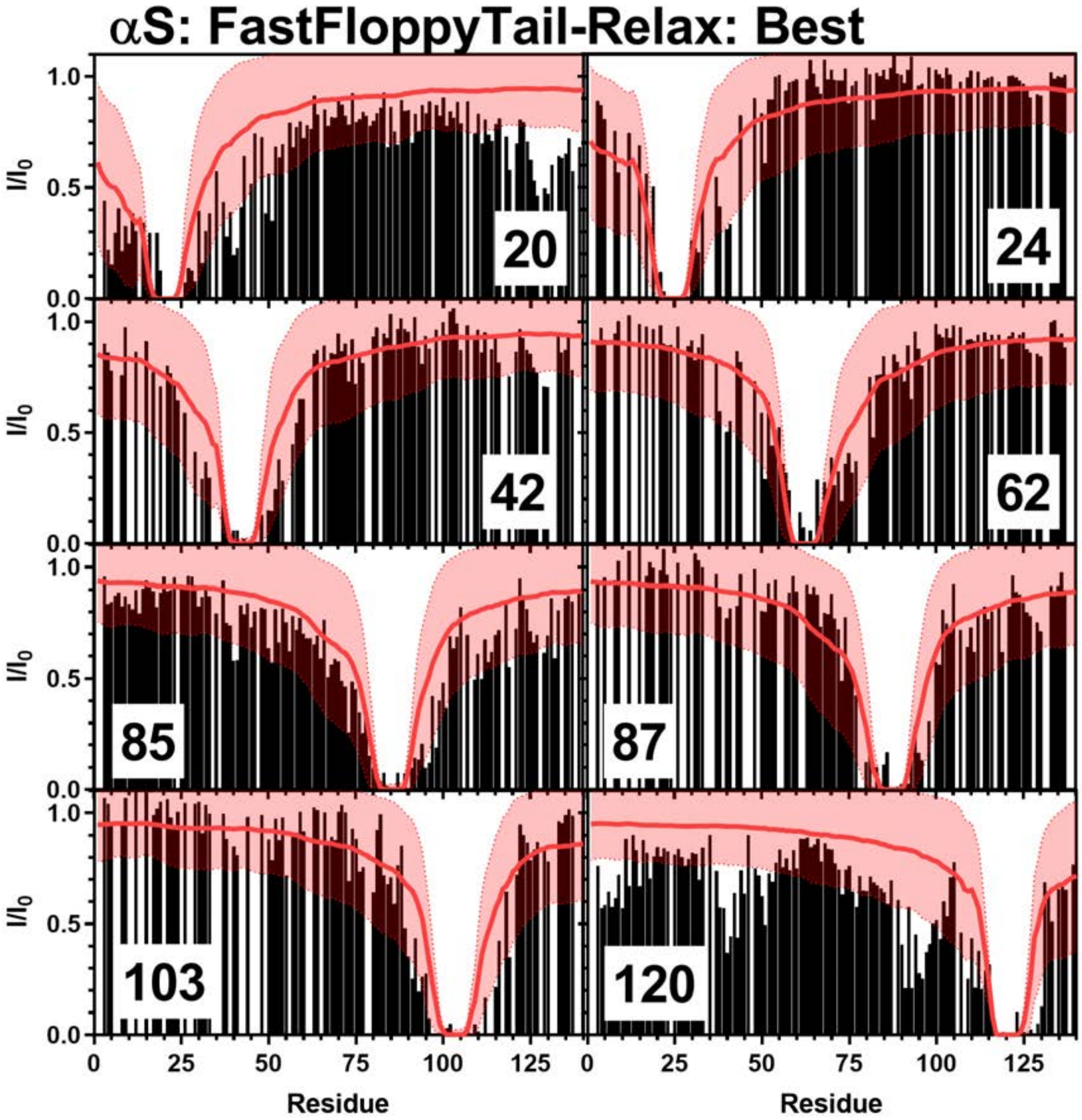


**Figure S178:** Comparison of Simulated PRE Values and Experimental PRE Values: Simulated PRE values from FastFloppyTail-Relax using best-reweighting (red line) overlayed on top of experimental data (black bars) from positions 20 (Top Left), 24 (Top Right), 42 (Upper Middle Left), 62 (Upper Middle Right), 85 (Lower Middle Left), 87 (Lower Middle Right), 103 (Bottom Left), 120 (Bottom Right). Experimental data for positions 20, 85, and 120 are from Sung *et al.* [8] and data for positions 24, 42, 62, 87, and 103 are from Dedmon *et al.* [9].

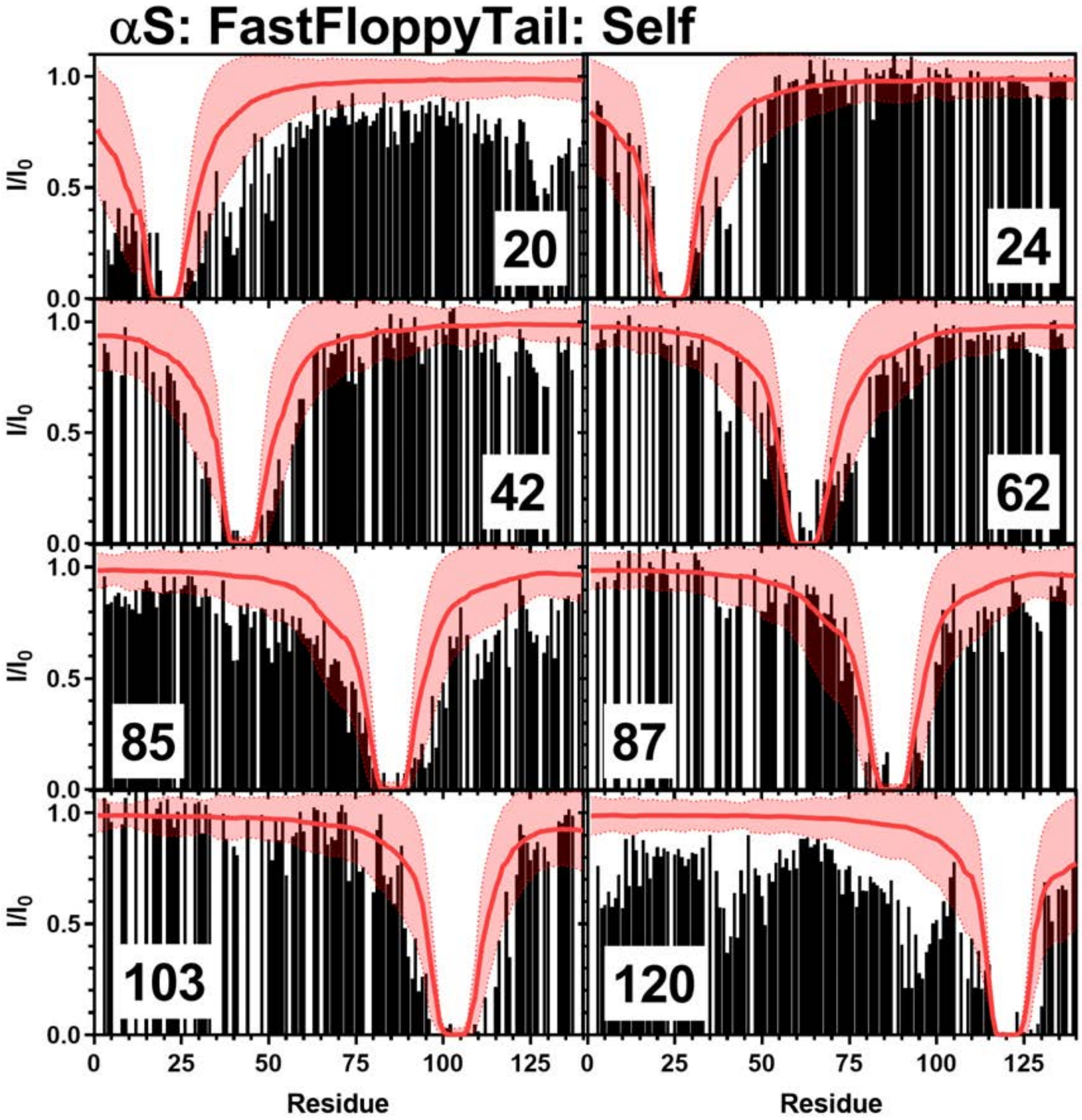


**Figure S179:** Comparison of Simulated PRE Values and Experimental PRE Values: Simulated PRE values from FastFloppyTail using self-reweighting (red line) overlayed on top of experimental data (black bars) from positions 20 (Top Left), 24 (Top Right), 42 (Upper Middle Left), 62 (Upper Middle Right), 85 (Lower Middle Left), 87 (Lower Middle Right), 103 (Bottom Left), 120 (Bottom Right). Experimental data for positions 20, 85, and 120 are from Sung *et al.* [8] and data for positions 24, 42, 62, 87, and 103 are from Dedmon *et al.* [9].

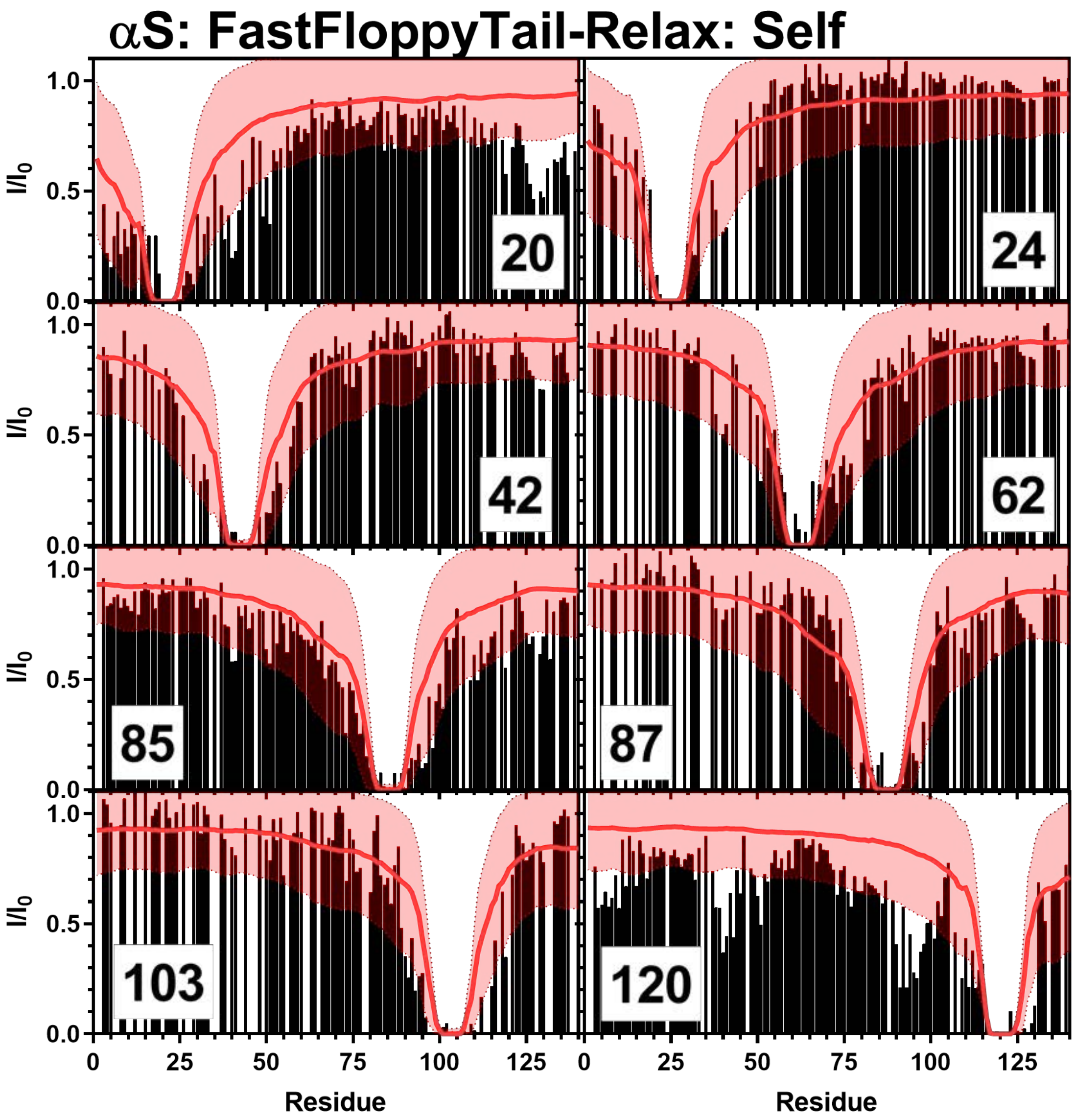


**Figure S180:** Comparison of Simulated PRE Values and Experimental PRE Values: Simulated PRE values from FastFloppyTail-Relax using self-reweighting (red line) overlayed on top of experimental data (black bars) from positions 20 (Top Left), 24 (Top Right), 42 (Upper Middle Left), 62 (Upper Middle Right), 85 (Lower Middle Left), 87 (Lower Middle Right), 103 (Bottom Left), 120 (Bottom Right). Experimental data for positions 20, 85, and 120 are from Sung *et al.* [8] and data for positions 24, 42, 62, 87, and 103 are from Dedmon *et al.* [9].

### Comparisons of DSH3 Data

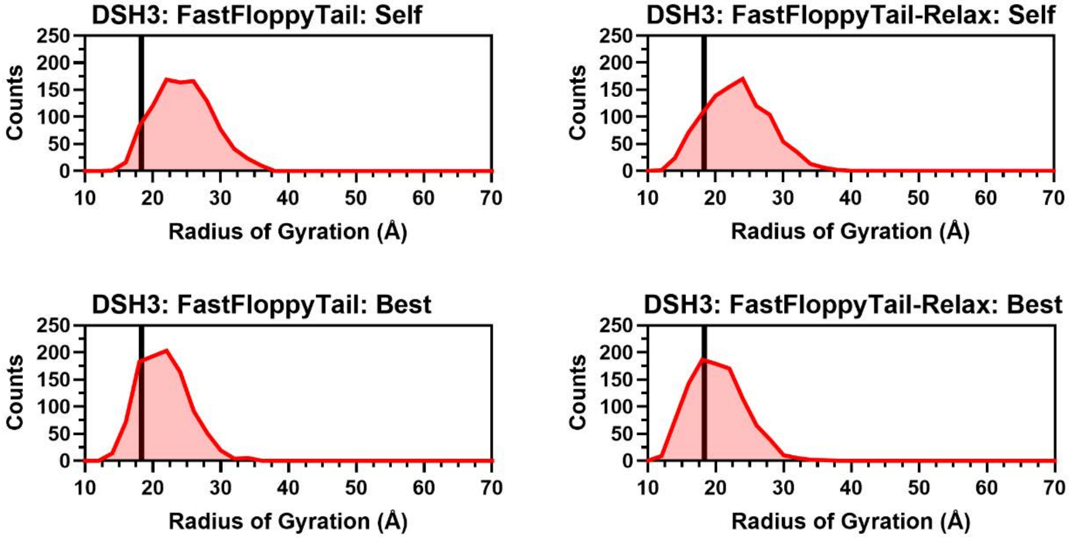


**Figure S190:** Histograms of Radii of Gyration for DSH3. Histograms of the radius of gyration from self (Top Left) and best (Top Right) reweighting in FastFloppyTail and self (Bottom Left) and best (Bottom Right) reweighting in FastFloppyTail-Relax compared to experimental values reported by Marsh *et al.* (black) [31].

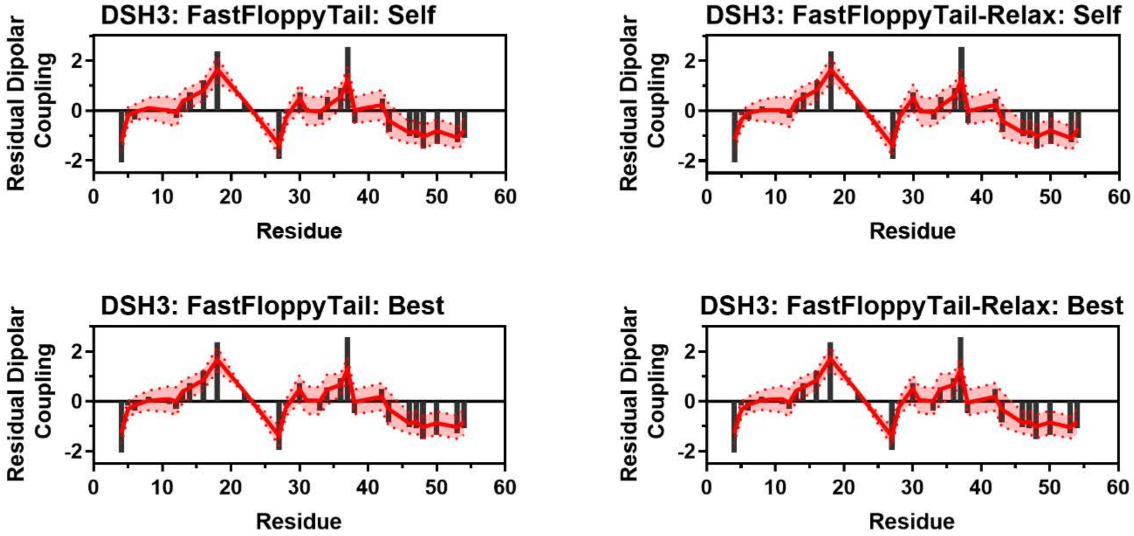


**Figure S191:** Comparison of Simulated and Experimental Residual Dipolar Coupling Data. Simulated RDC values (red line) from self (Top Left) and best (Top Right) reweighting in FastFloppyTail and self (Bottom Left) and best (Bottom Right) reweighting in FastFloppyTail-Relax overlayed on experimental data (black bars) from Marsh *et al.* [31].

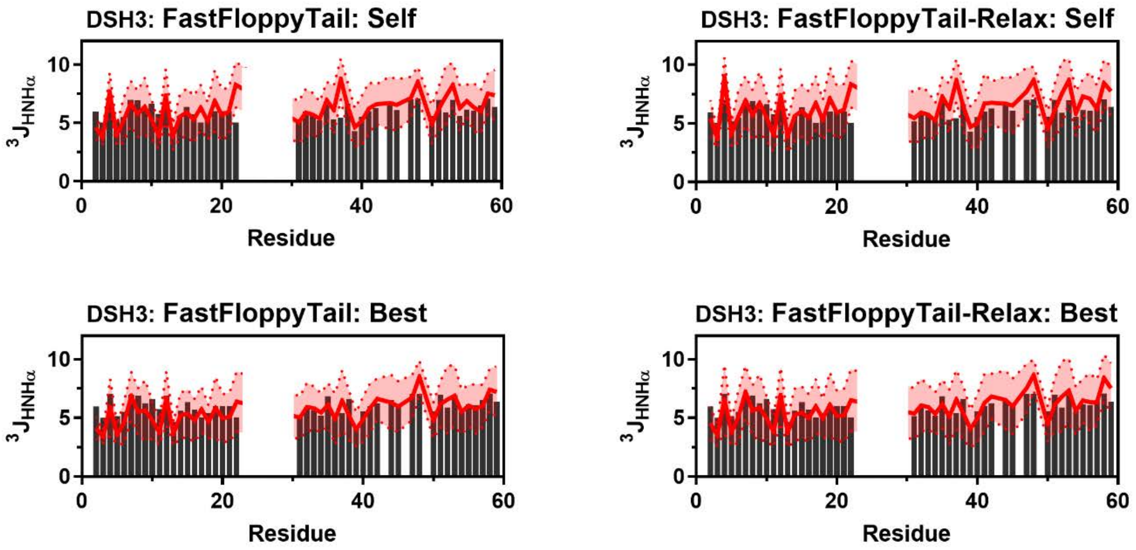


**Figure S192:** Comparison of Simulated and Experimental J-Coupling Data. Simulated $^3J_{HNH\alpha}$ values (red lines) from self (Top Left) and best (Top Right) reweighting in FastFloppyTail and self (Bottom Left) and best (Bottom Right) reweighting in FastFloppyTail-Relax compared to experimental values from Zhang *et al.* (black bars) [37].

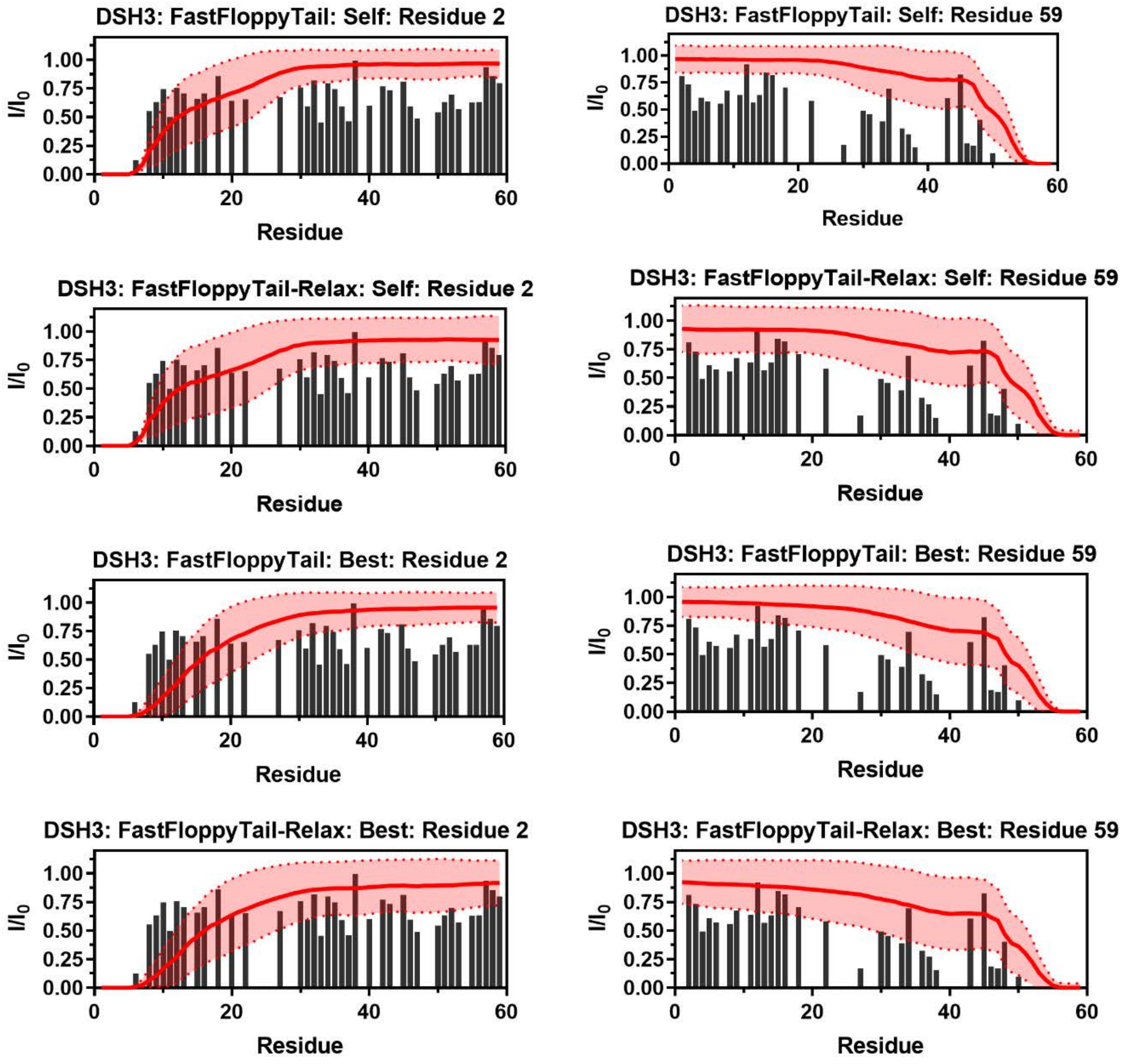


**Figure S193:** Comparison of Simulated PRE Values and Experimental PRE Values: Simulated PRE values (red lines) from FastFloppyTail (Row 1) and FastFloppyTail-Relax (Row 2) using self-reweighting and FastFloppyTail (Row 3) and FastFloppyTail-Relax (Row 4) using best-reweighting overlayed on top of experimental data (black bars) from positions 2 (Left) and 59 (Right) from Marsh *et al.* (black) [31].

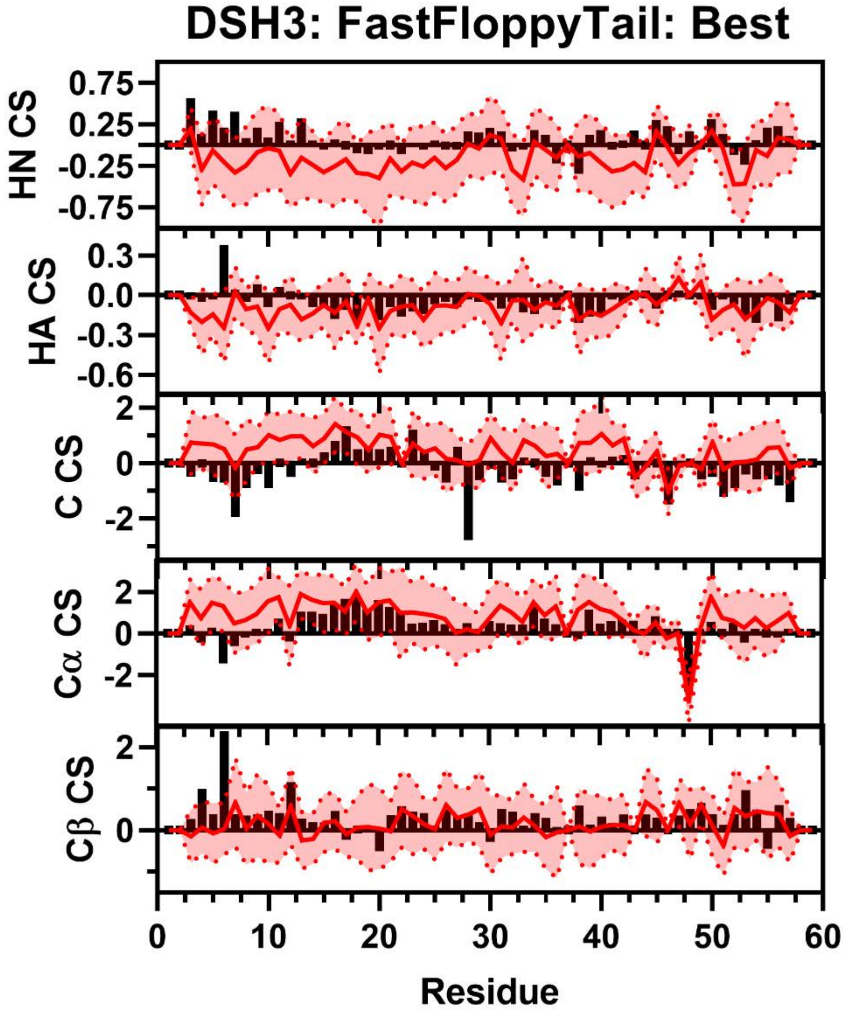


**Figure S194:** Comparison of Simulated and Experimental NMR Secondary Chemical Shift Data for DSH3. Simulated Chemical Shift values from the FastFloppyTail ensemble using best-reweighted fragment selection (red) overlayed on experimental data (black bars) of HN (Row 1), HA (Row 2), C (Row 3), Cα (Row 4), and Cβ (Row 5) chemical shifts from Zhang *et al*.[37] Neighbor corrected random coil chemical shift values from Sparta+ have been subtracted from both simulated and experimental chemical shifts to generate secondary chemical shift data [11].

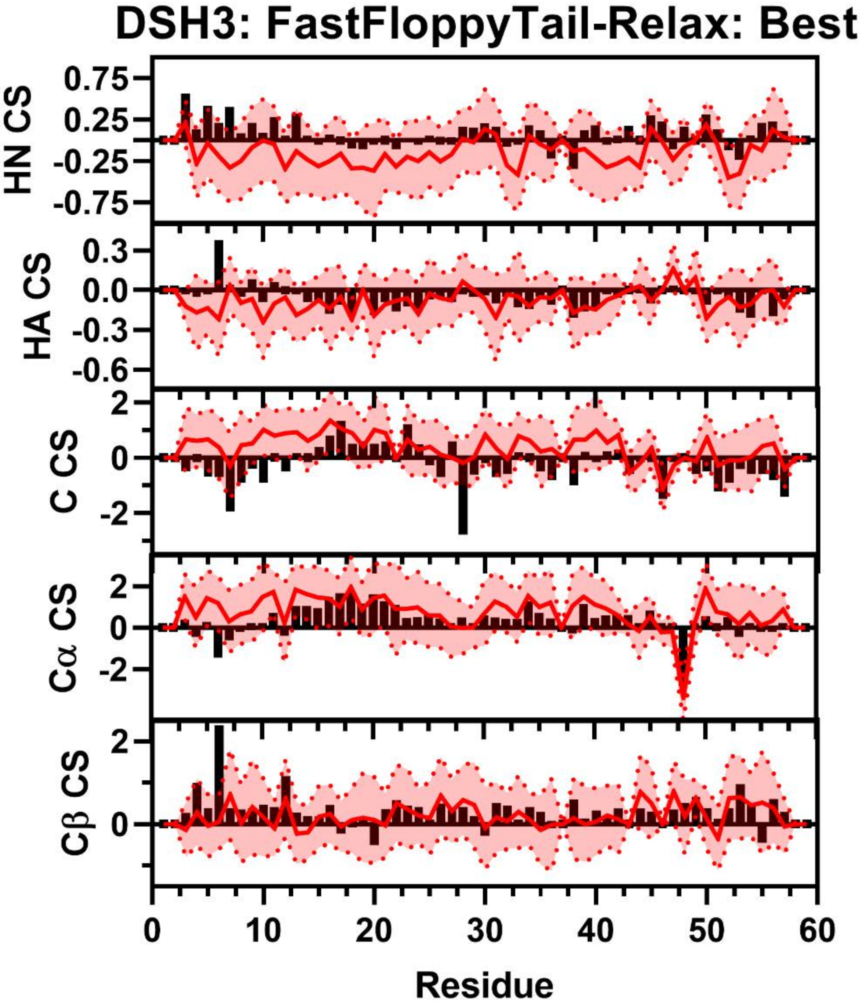


**Figure S195:** Comparison of Simulated and Experimental NMR Secondary Chemical Shift Data for DSH3. Simulated Chemical Shift values from the FastFloppyTail-Relax ensemble using best-reweighted fragment selection (red) overlayed on experimental data (black bars) of HN (Row 1), HA (Row 2), C (Row 3), Cα (Row 4), and Cβ (Row 5) chemical shifts from Zhang *et al*. [37] Neighbor corrected random coil chemical shift values from SPARTA+ have been subtracted from both simulated and experimental chemical shifts to generate secondary chemical shift data [11].

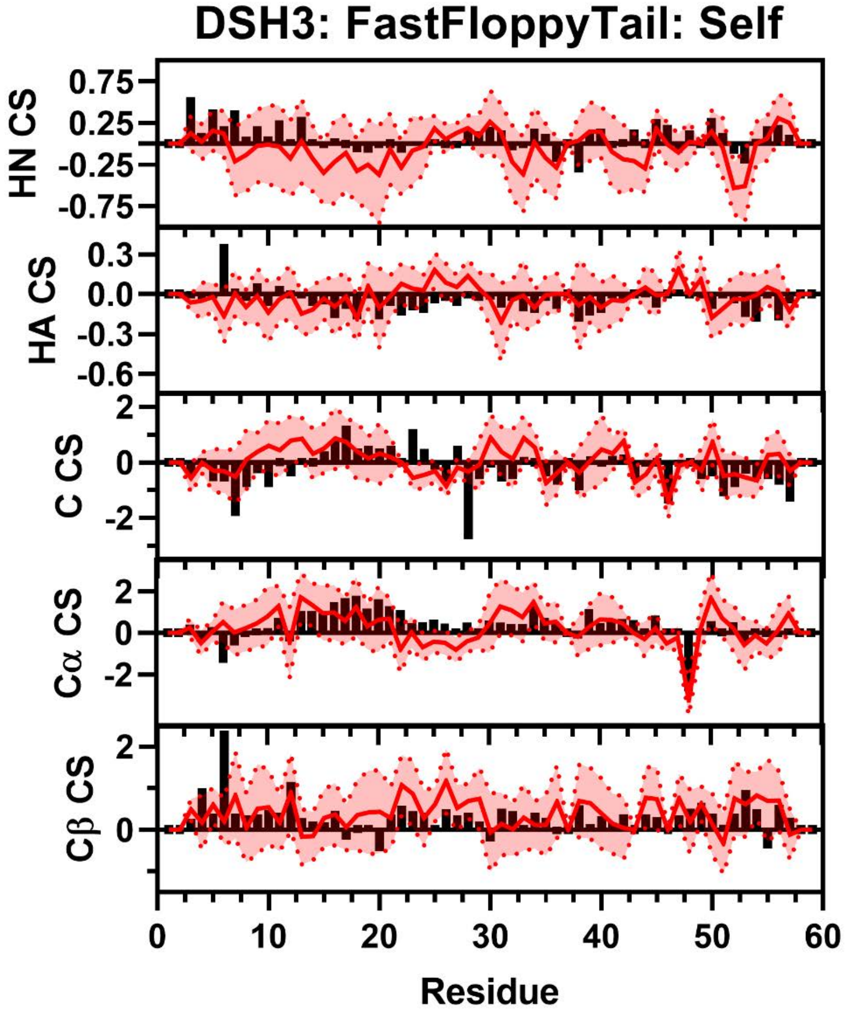


**Figure S196:** Comparison of Simulated and Experimental NMR Secondary Chemical Shift Data for DSH3. Simulated Chemical Shift values from the FastFloppyTail ensemble using self-reweighted fragment selection (red) overlayed on experimental data (black bars) of HN (Row 1), HA (Row 2), C (Row 3), Cα (Row 4), and Cβ (Row 5) chemical shifts from Zhang *et al*. [37] Neighbor corrected random coil chemical shift values from Sparta+ have been subtracted from both simulated and experimental chemical shifts to generate secondary chemical shift data [11].

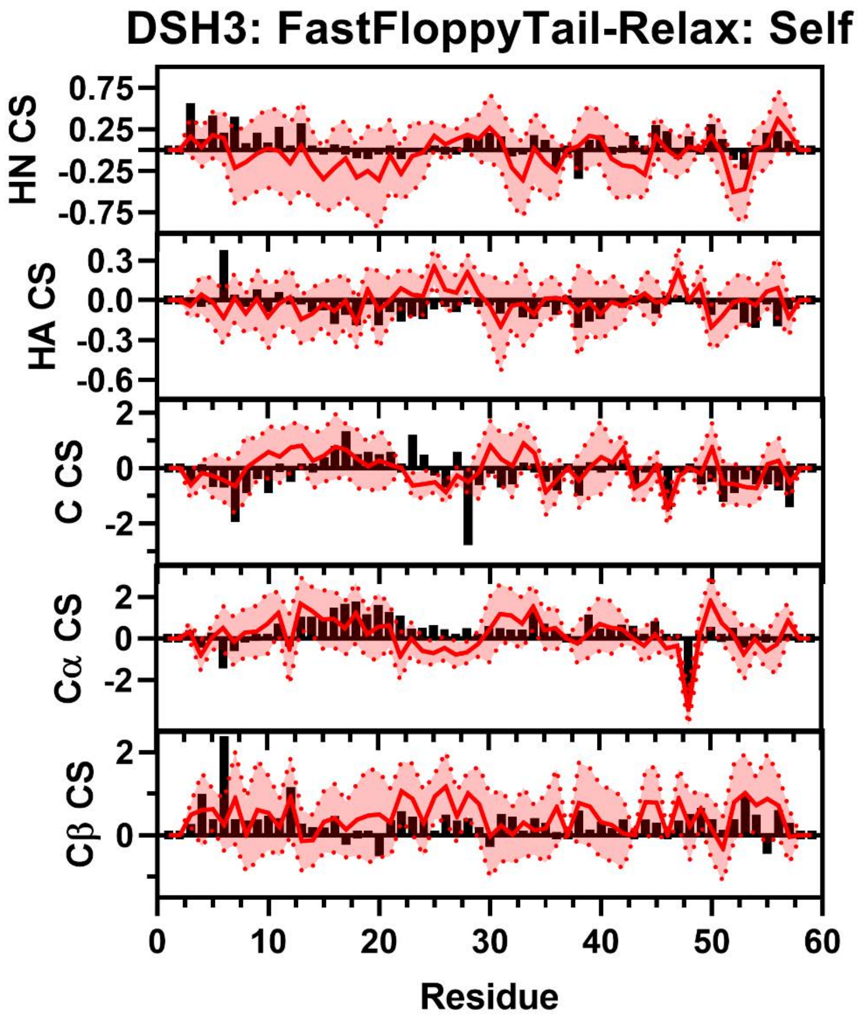


**Figure S197:** Comparison of Simulated and Experimental NMR Secondary Chemical Shift Data for DSH3. Simulated Chemical Shift values from the FastFloppyTail-Relax ensemble using self-reweighted fragment selection (red) overlayed on experimental data (black bars) of HN (Row 1), HA (Row 2), C (Row 3), Cα (Row 4), and Cβ (Row 5) chemical shifts from Zhang *et al*. [37] Neighbor corrected random coil chemical shift values from Sparta+ have been subtracted from both simulated and experimental chemical shifts to generate secondary chemical shift data [11].

**Comparisons of NTAL Data**

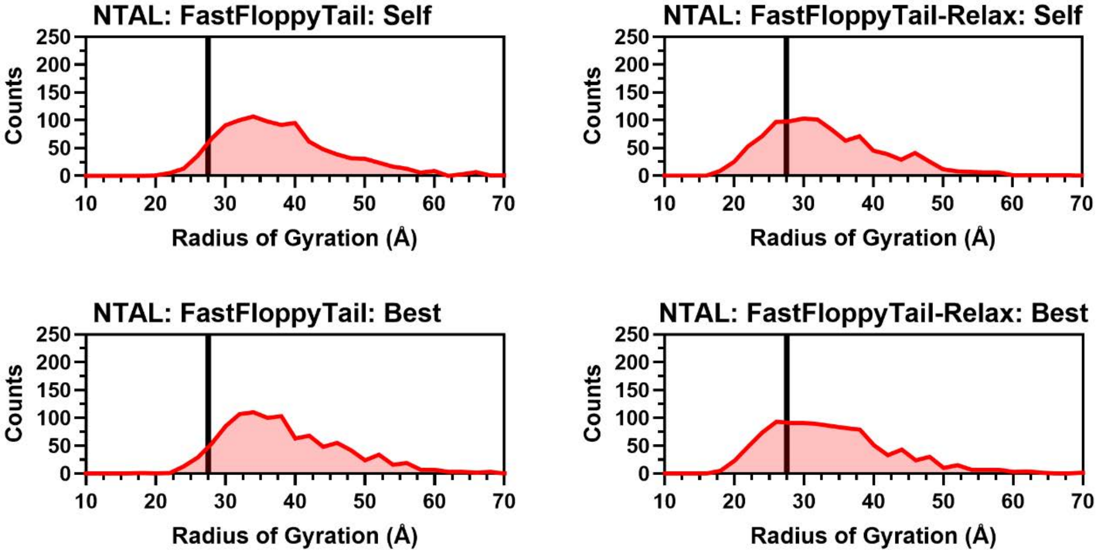


**Figure S198:** Histograms of Radii of Gyration for DSH3. Histograms of the radius of gyration from self (Top Left) and best (Top Right) reweighting in FastFloppyTail and self (Bottom Left) and best (Bottom Right) reweighting in FastFloppyTail-Relax compared to experimental values reported by Jensen *et al.* (black) [32].

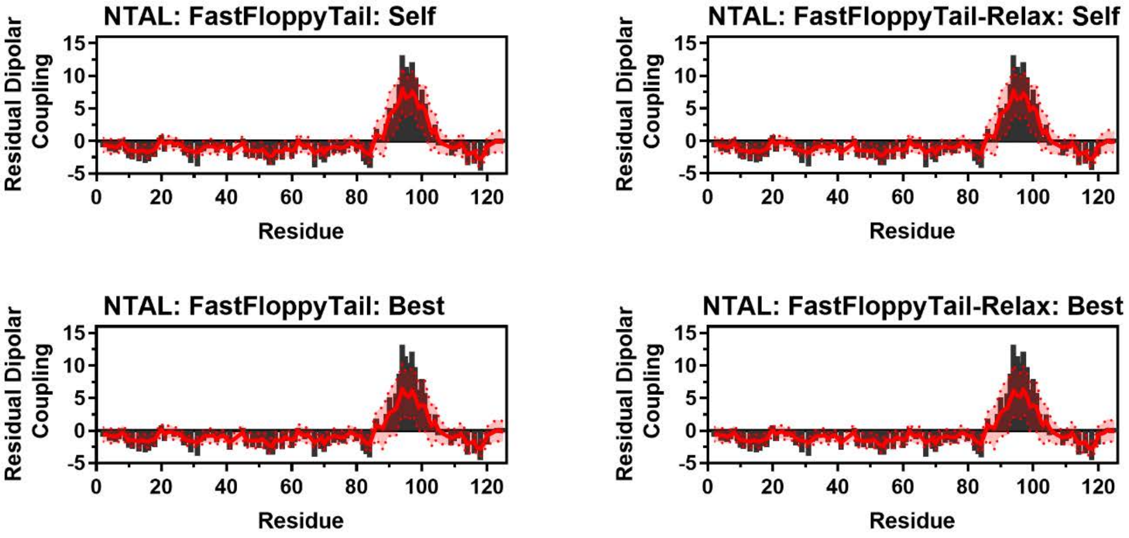


**Figure S199:** Comparison of Simulated and Experimental Residual Dipolar Coupling Data. Simulated RDC values (red line) from self (Top Left) and best (Top Right) reweighting in FastFloppyTail and self (Bottom Left) and best (Bottom Right) reweighting in FastFloppyTail-Relax overlayed on experimental data (black bars) from Jensen *et al.* [32].

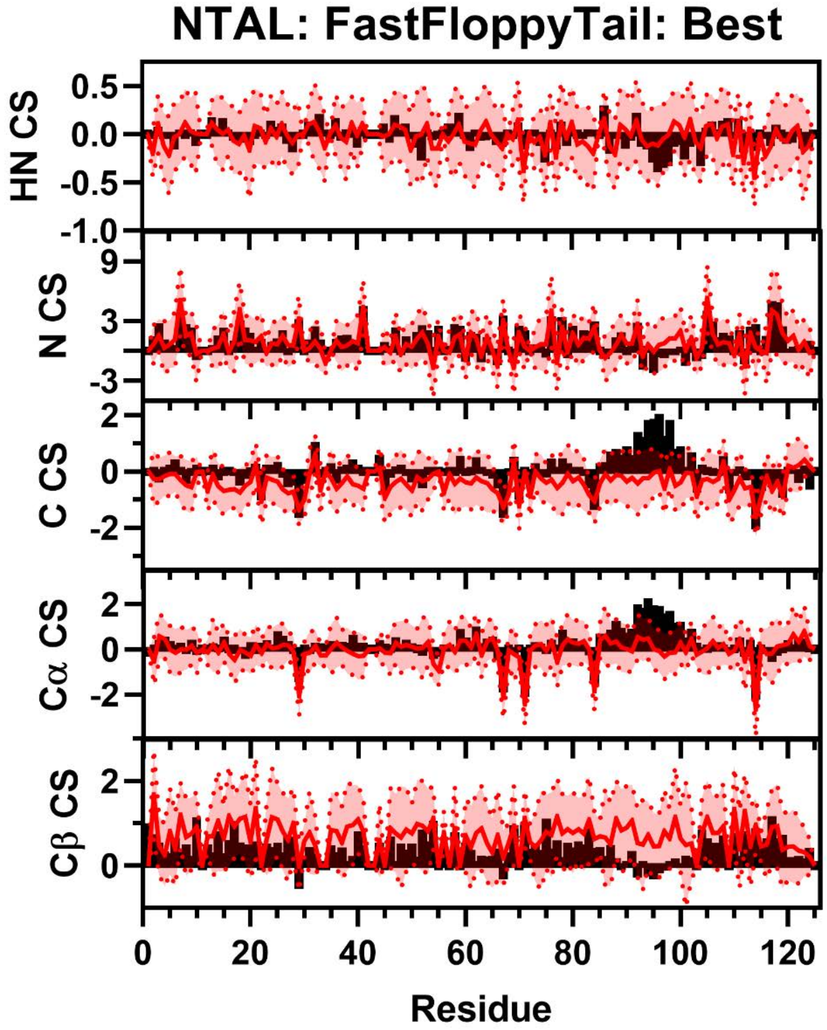


**Figure S200:** Comparison of Simulated and Experimental NMR Secondary Chemical Shift Data for NTAL. Simulated Chemical Shift values from the FastFloppyTail ensemble using best-reweighted fragment selection (red) overlayed on experimental data (black bars) of HN (Row 1), HA (Row 2), C (Row 3), Cα (Row 4), and Cβ (Row 5) chemical shifts from Jensen *et. al*.[32] Neighbor corrected random coil chemical shift values from SPARTA+ have been subtracted from both simulated and experimental chemical shifts to generate secondary chemical shift data [11].

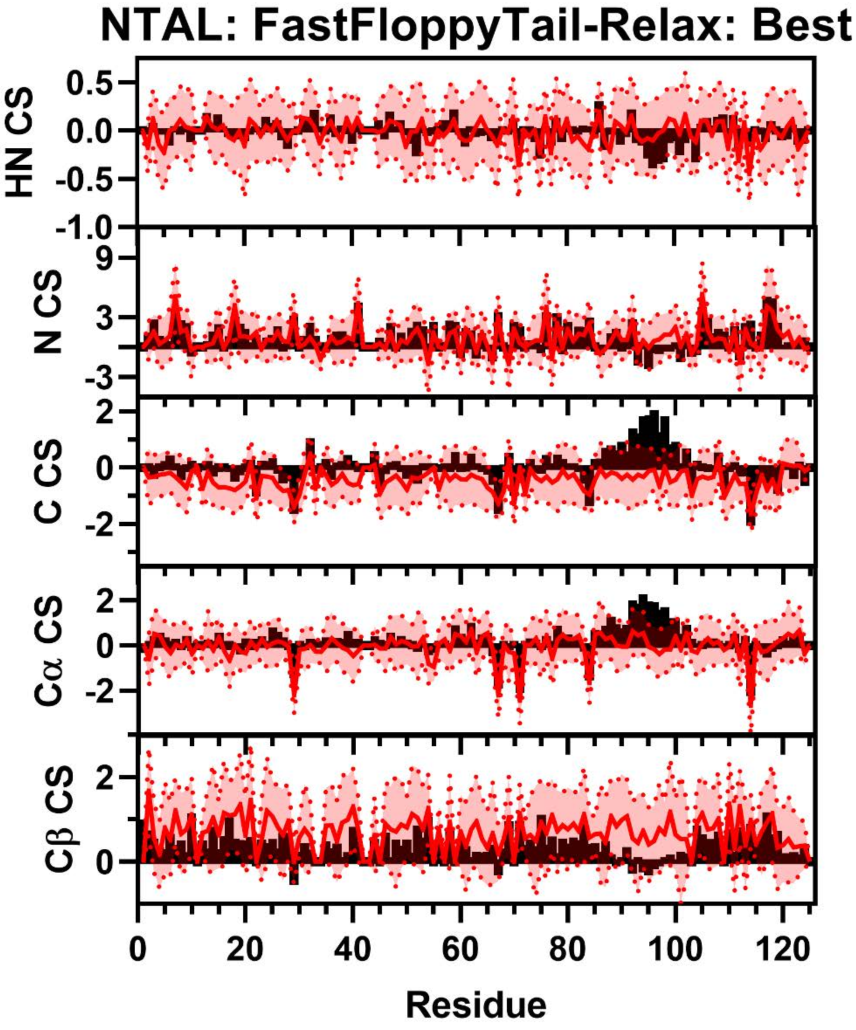


**Figure S201:** Comparison of Simulated and Experimental NMR Secondary Chemical Shift Data for NTAL. Simulated Chemical Shift values from the FastFloppyTail-Relax ensemble using best-reweighted fragment selection (red) overlayed on experimental data (black bars) of HN (Row 1), HA (Row 2), C (Row 3), Cα (Row 4), and Cβ (Row 5) chemical shifts from Jensen *et al*. [32] Neighbor corrected random coil chemical shift values from SPARTA+ have been subtracted from both simulated and experimental chemical shifts to generate secondary chemical shift data [11].

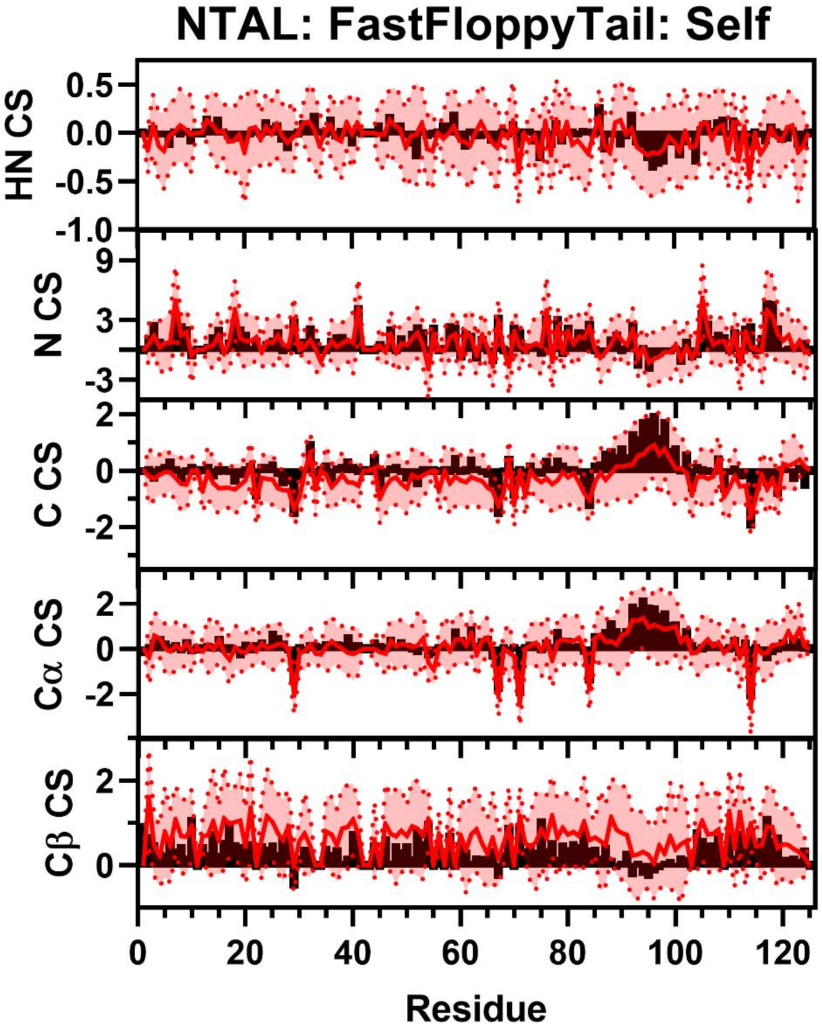


**Figure S202:** Comparison of Simulated and Experimental NMR Secondary Chemical Shift Data for NTAL. Simulated Chemical Shift values from the FastFloppyTail ensemble using self-reweighted fragment selection (red) overlayed on experimental data (black bars) of HN (Row 1), HA (Row 2), C (Row 3), Cα (Row 4), and Cβ (Row 5) chemical shifts from Jensen *et al.* [32] Neighbor corrected random coil chemical shift values from Sparta+ have been subtracted from both simulated and experimental chemical shifts to generate secondary chemical shift data [11].

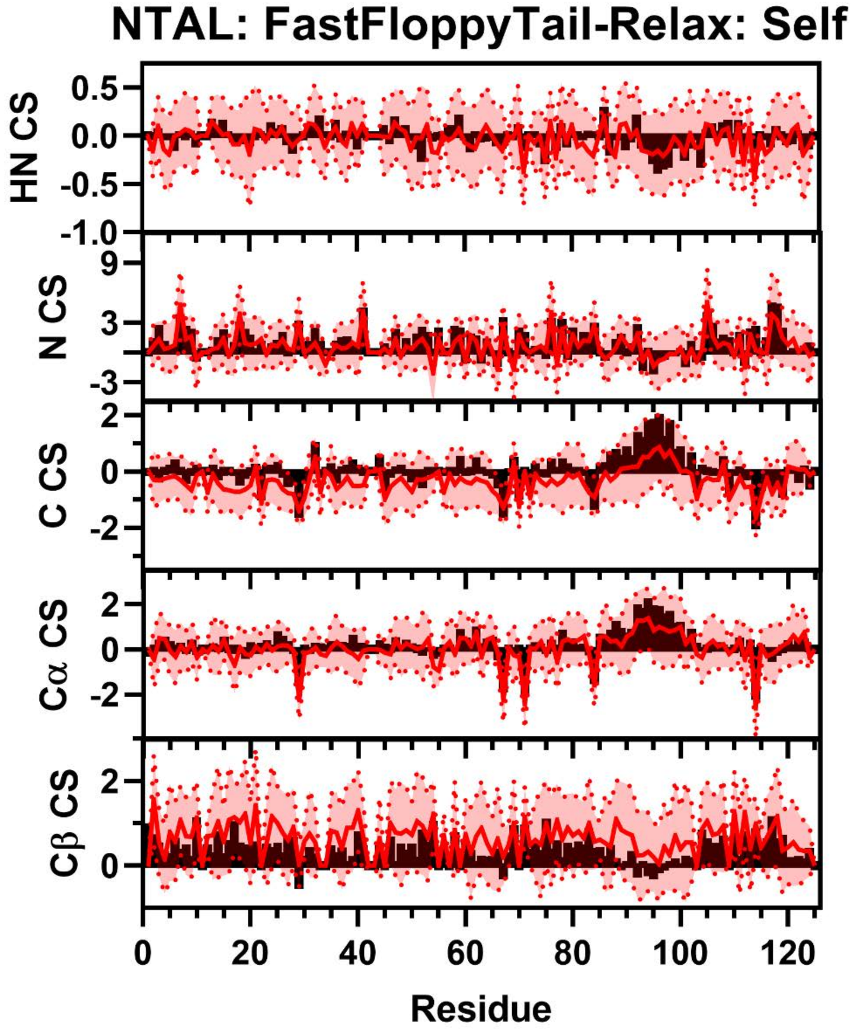


**Figure S203:** Comparison of Simulated and Experimental NMR Secondary Chemical Shift Data for NTAL. Simulated Chemical Shift values from the FastFloppyTail-Relax ensemble using self-reweighted fragment selection (red) overlayed on experimental data (black bars) of HN (Row 1), HA (Row 2), C (Row 3), Cα (Row 4), and Cβ (Row 5) chemical shifts from Jensen *et al*. [32] Neighbor corrected random coil chemical shift values from Sparta+ have been subtracted from both simulated and experimental chemical shifts to generate secondary chemical shift data [11].

### Comparisons of PAAA Data

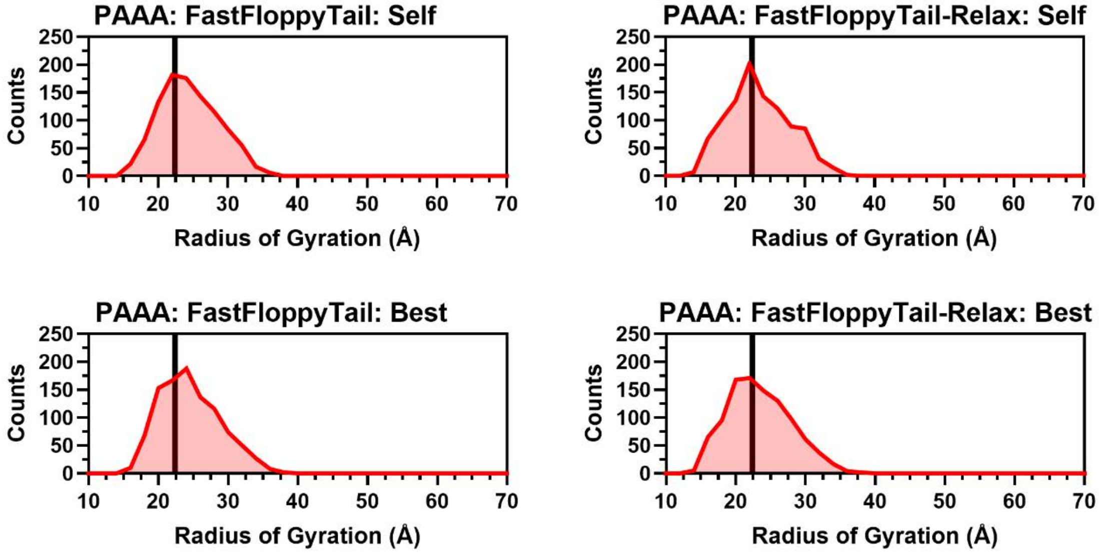


**Figure S204:** Histograms of Radii of Gyration for PAAA. Histograms of the radius of gyration from self (Top Left) and best (Top Right) reweighting in FastFloppyTail and self (Bottom Left) and best (Bottom Right) reweighting in FastFloppyTail-Relax compared to experimental values from NMR (black) [33].

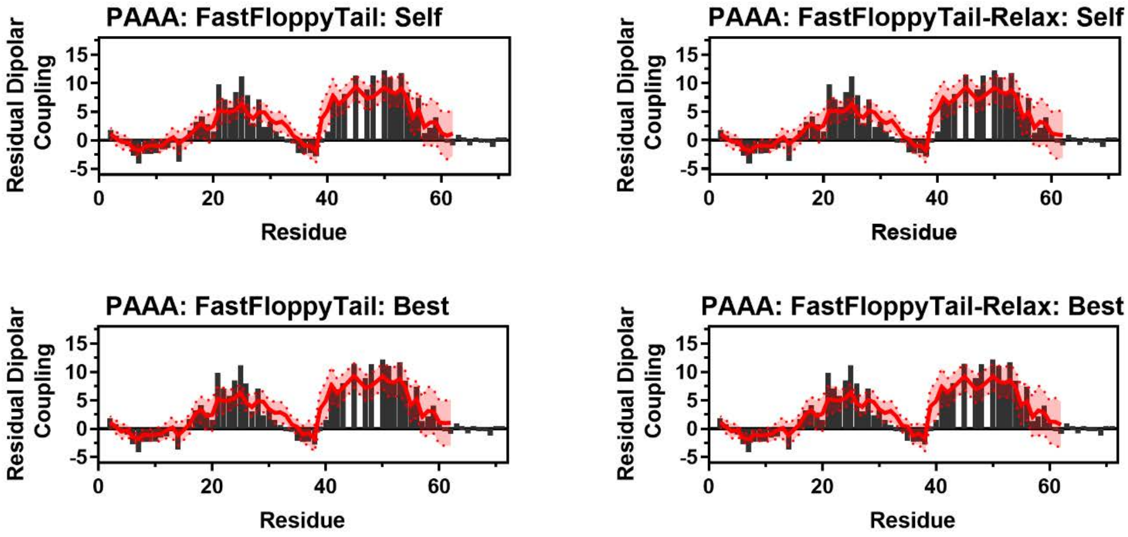


**Figure S205:** Comparison of Simulated and Experimental Residual Dipolar Coupling Data. Simulated RDC values (red line) from self (Top Left) and best (Top Right) reweighting in FastFloppyTail and self (Bottom Left) and best (Bottom Right) reweighting in FastFloppyTail-Relax overlayed on experimental data (black bars) from Streckx *et al*. [33].

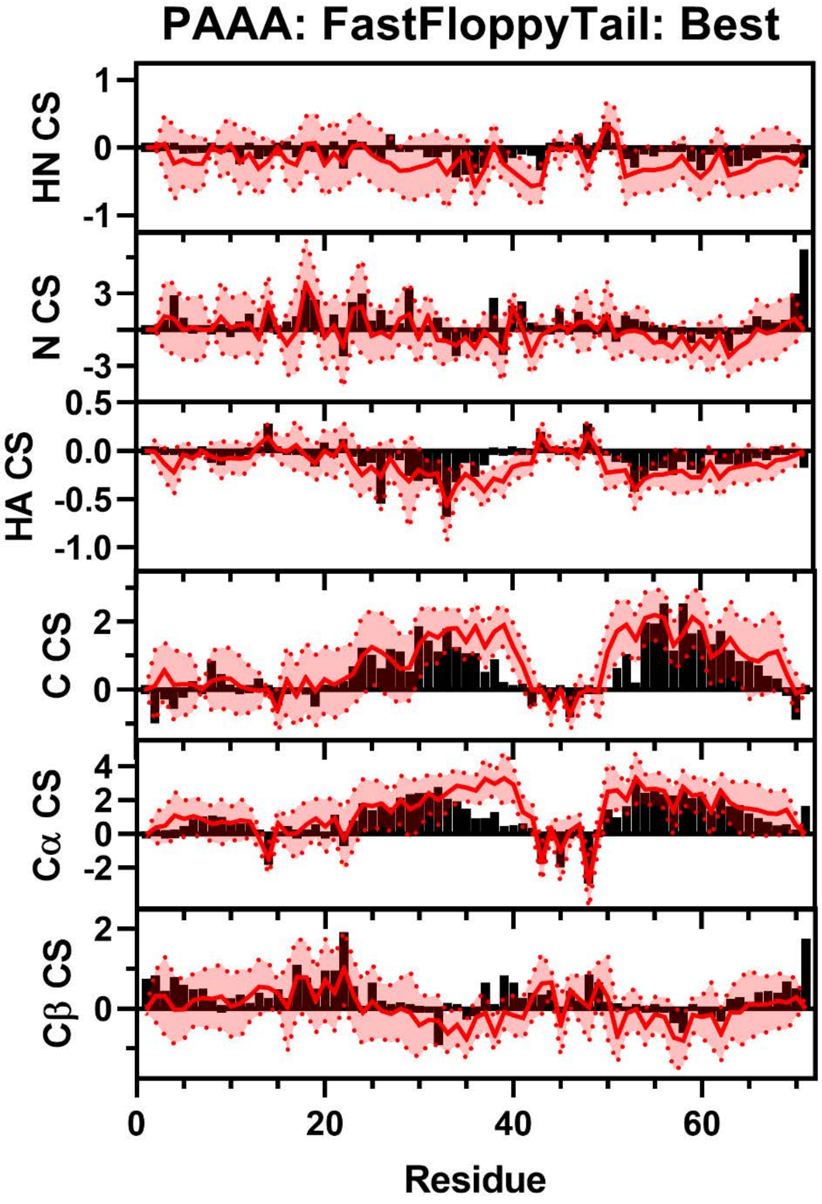


**Figure S206:** Comparison of Simulated and Experimental NMR Secondary Chemical Shift Data for NTAL. Simulated Chemical Shift values from the FastFloppyTail ensemble using best-reweighted fragment selection (red) overlayed on experimental data (black bars) of HN (Row 1), N (Row 2), HA (Row 3), C, (Row 4), Cα (Row 5), and Cβ (Row 6) chemical shifts from Streckx *et al*. [33] Neighbor corrected random coil chemical shift values from Sparta+ have been subtracted from both simulated and experimental chemical shifts to generate secondary chemical shift data [11].

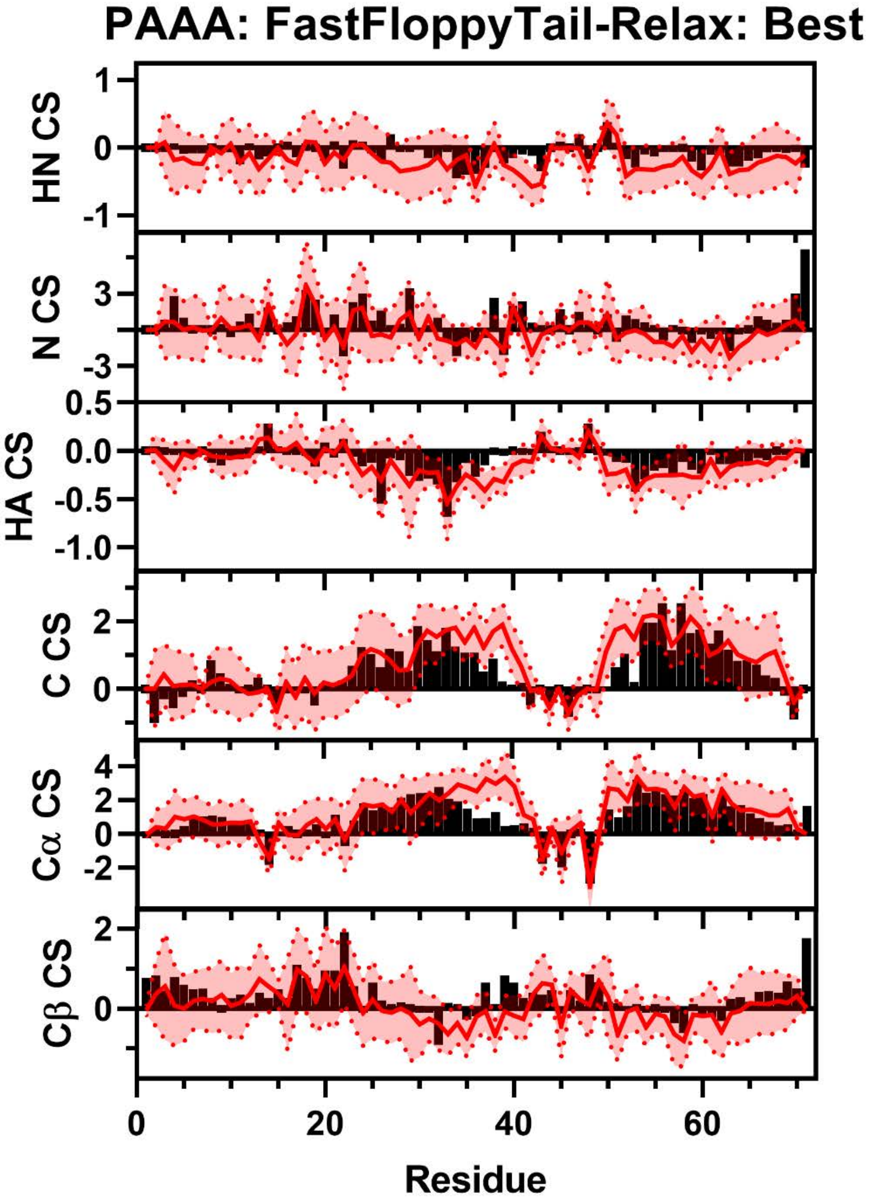


**Figure S207:** Comparison of Simulated and Experimental NMR Secondary Chemical Shift Data for NTAL. Simulated Chemical Shift values from the FastFloppyTail-Relax ensemble using best-reweighted fragment selection (red) overlayed on experimental data (black bars) of HN (Row 1), N (Row 2), HA (Row 3), C, (Row 4), Cα (Row 5), and Cβ (Row 6) chemical shifts from Streckx *et al*. [33] Neighbor corrected random coil chemical shift values from SPARTA+ have been subtracted from both simulated and experimental chemical shifts to generate secondary chemical shift data [11].

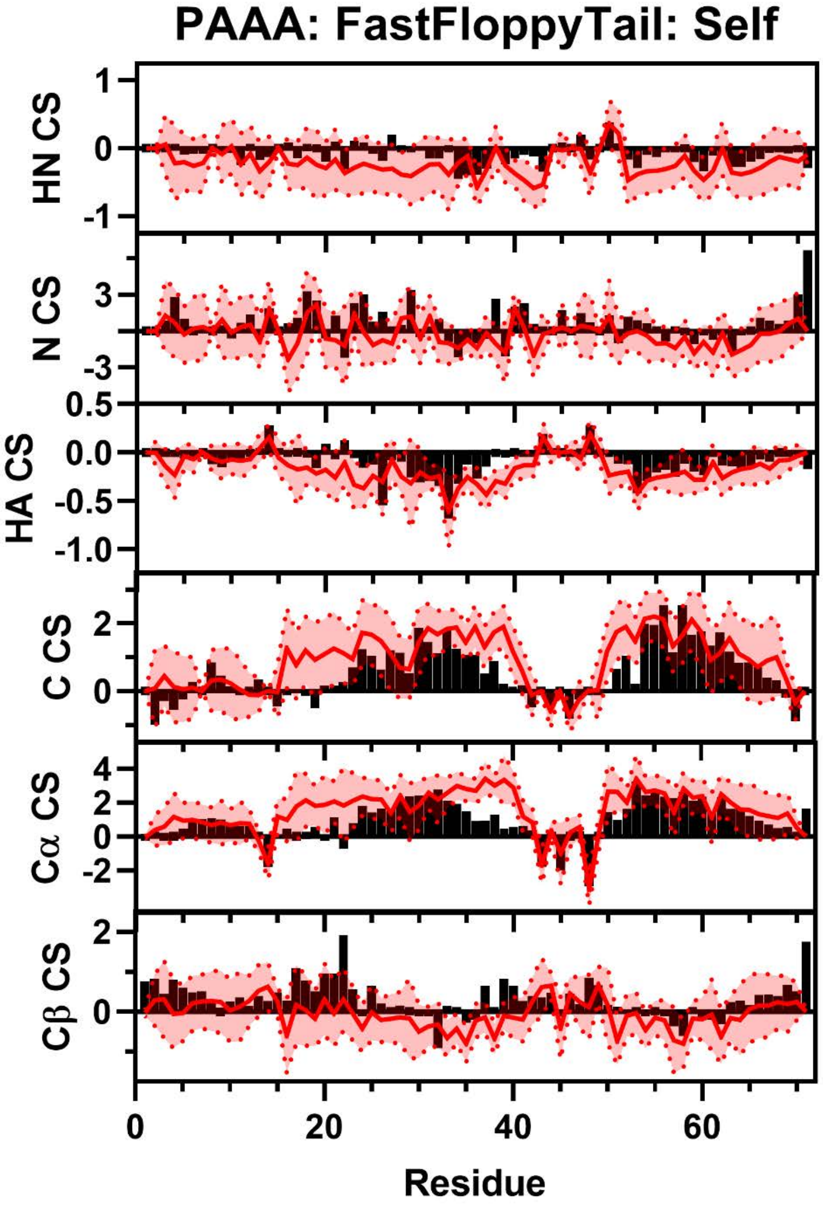


**Figure S208:** Comparison of Simulated and Experimental NMR Secondary Chemical Shift Data for NTAL. Simulated Chemical Shift values from the FastFloppyTail ensemble using self-reweighted fragment selection (red) overlayed on experimental data (black bars) of HN (Row 1), N (Row 2), HA (Row 3), C, (Row 4), Cα (Row 5), and Cβ (Row 6) chemical shifts from Streckx *et al*. [33] Neighbor corrected random coil chemical shift values from SPARTA+ have been subtracted from both simulated and experimental chemical shifts to generate secondary chemical shift data [11].

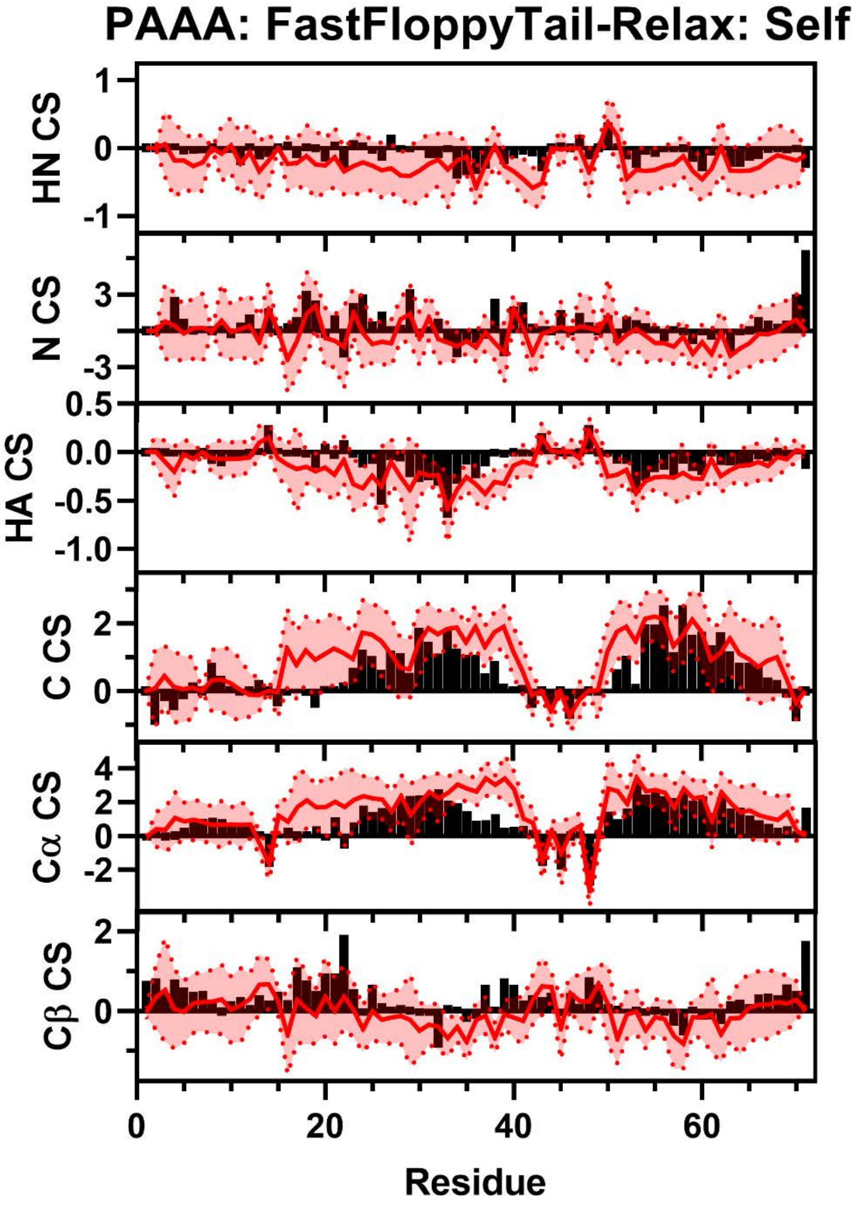


**Figure S209:** Comparison of Simulated and Experimental NMR Secondary Chemical Shift Data for NTAL. Simulated Chemical Shift values from the FastFloppyTail-Relax ensemble using self-reweighted fragment selection (red) overlayed on experimental data (black bars) of HN (Row 1), N (Row 2), HA (Row 3), C, (Row 4), Cα (Row 5), and Cβ (Row 6) chemical shifts from Streckx *et al*.[33] Neighbor corrected random coil chemical shift values from SPARTA+ have been subtracted from both simulated and experimental chemical shifts to generate secondary chemical shift data [11].

**Comparisons of SIC1 Data**

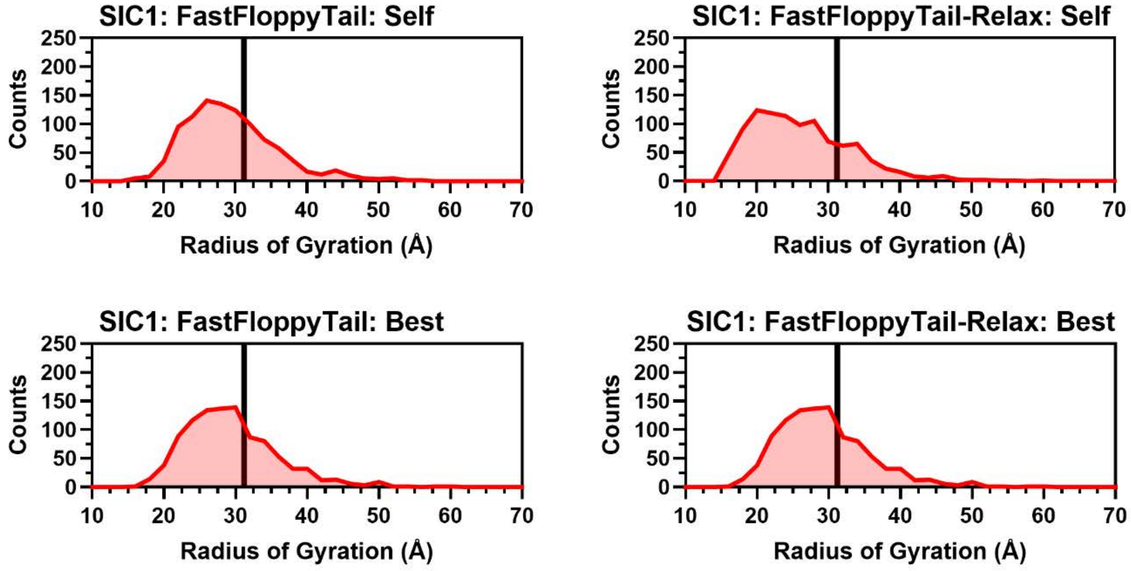


**Figure S210:** Histograms of Radii of Gyration for SIC1. Histograms of the radius of gyration from self (Top Left) and best (Top Right) reweighting in FastFloppyTail and self (Bottom Left) and best (Bottom Right) reweighting in FastFloppyTail-Relax compared to experimental values (black) reported by Mittag *et al*. [34].

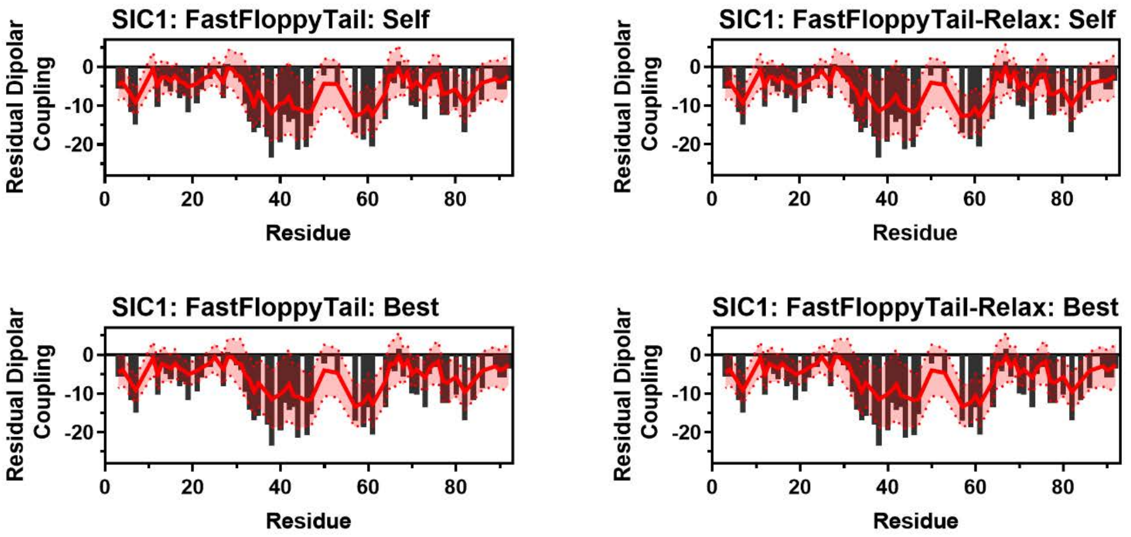


**Figure S211:** Comparison of Simulated and Experimental Residual Dipolar Coupling Data. Simulated RDC values (red line) from self (Top Left) and best (Top Right) reweighting in FastFloppyTail and self (Bottom Left) and best (Bottom Right) reweighting in FastFloppyTail-Relax overlayed on experimental data (black bars) from Mittag *et al*. [34].

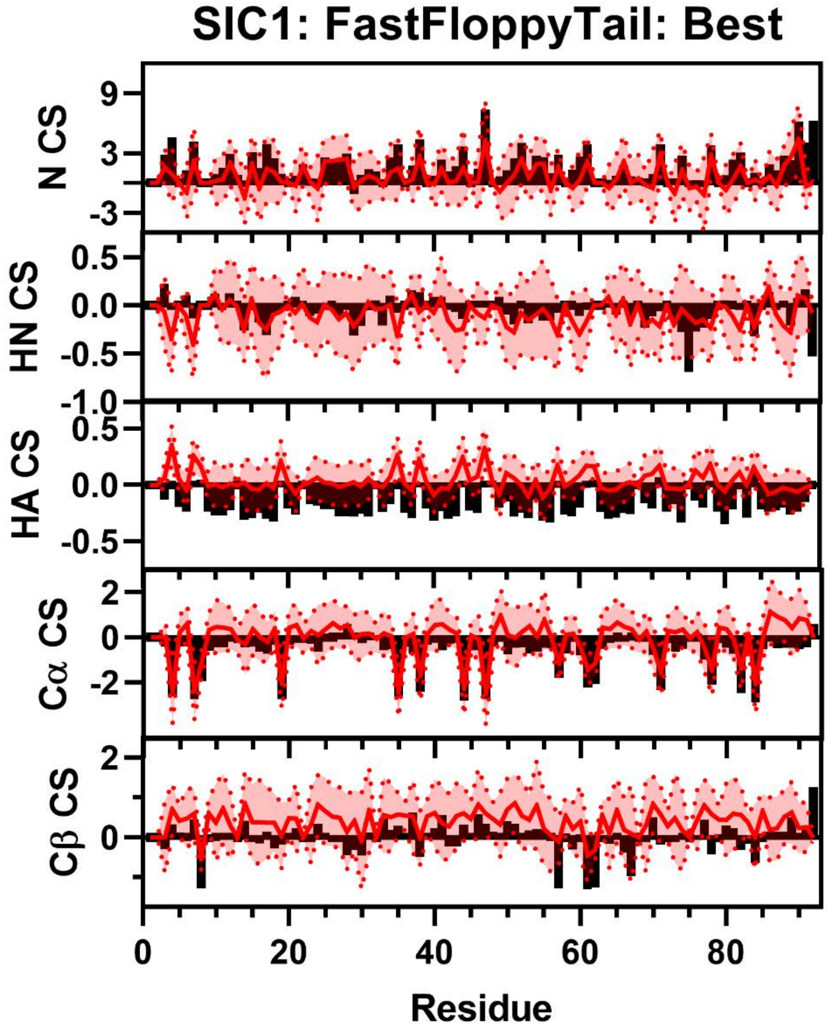


**Figure S212:** Comparison of Simulated and Experimental NMR Secondary Chemical Shift Data for SIC1. Simulated Chemical Shift values from the FastFloppyTail ensemble using best-reweighted fragment selection (red) overlayed on experimental data (black bars) of N (Row 1), HN (Row 2), HA (Row 3), Cα (Row 4), and Cβ (Row 5) chemical shifts from Mittag *et al.* [34] Neighbor corrected random coil chemical shift values from Sparta+ have been subtracted from both simulated and experimental chemical shifts to generate secondary chemical shift data [11].

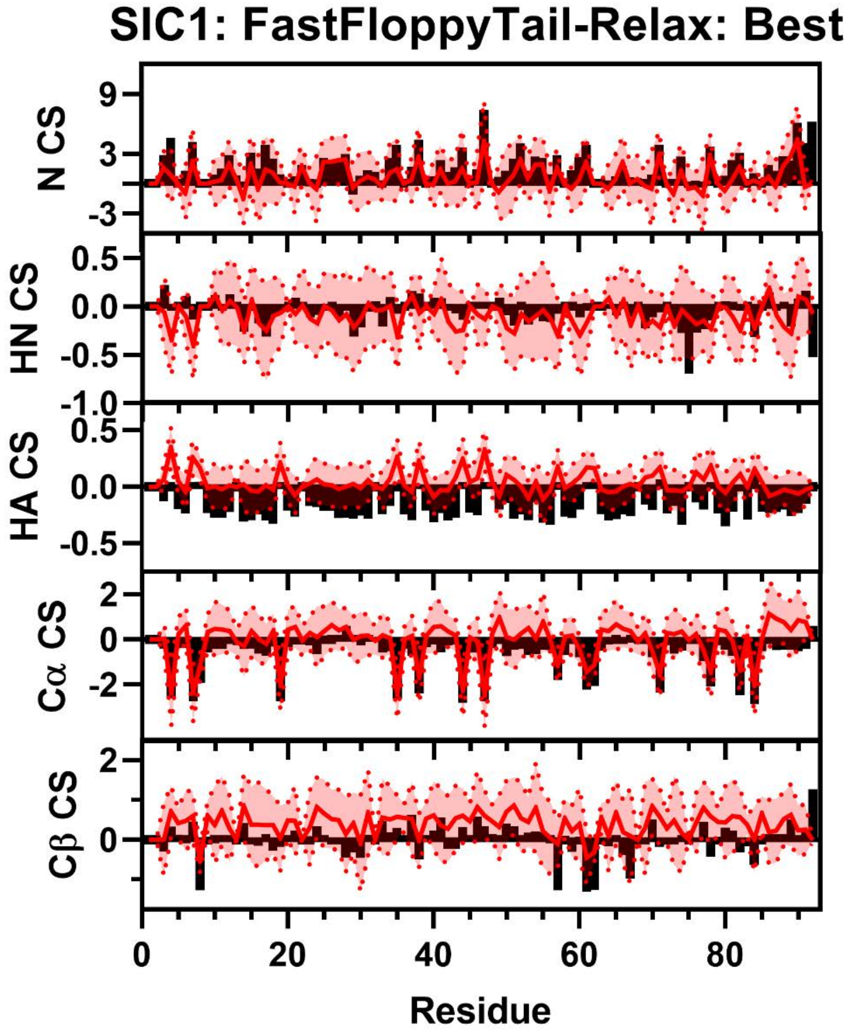


**Figure S213:** Comparison of Simulated and Experimental NMR Secondary Chemical Shift Data for SIC1. Simulated Chemical Shift values from the FastFloppyTail ensemble using best-reweighted fragment selection (red) overlayed on experimental data (black bars) of N (Row 1), HN (Row 2), HA (Row 3), Cα (Row 4), and Cβ (Row 5) chemical shifts from Mittag *et al.* [34] Neighbor corrected random coil chemical shift values from Sparta+ have been subtracted from both simulated and experimental chemical shifts to generate secondary chemical shift data [11].

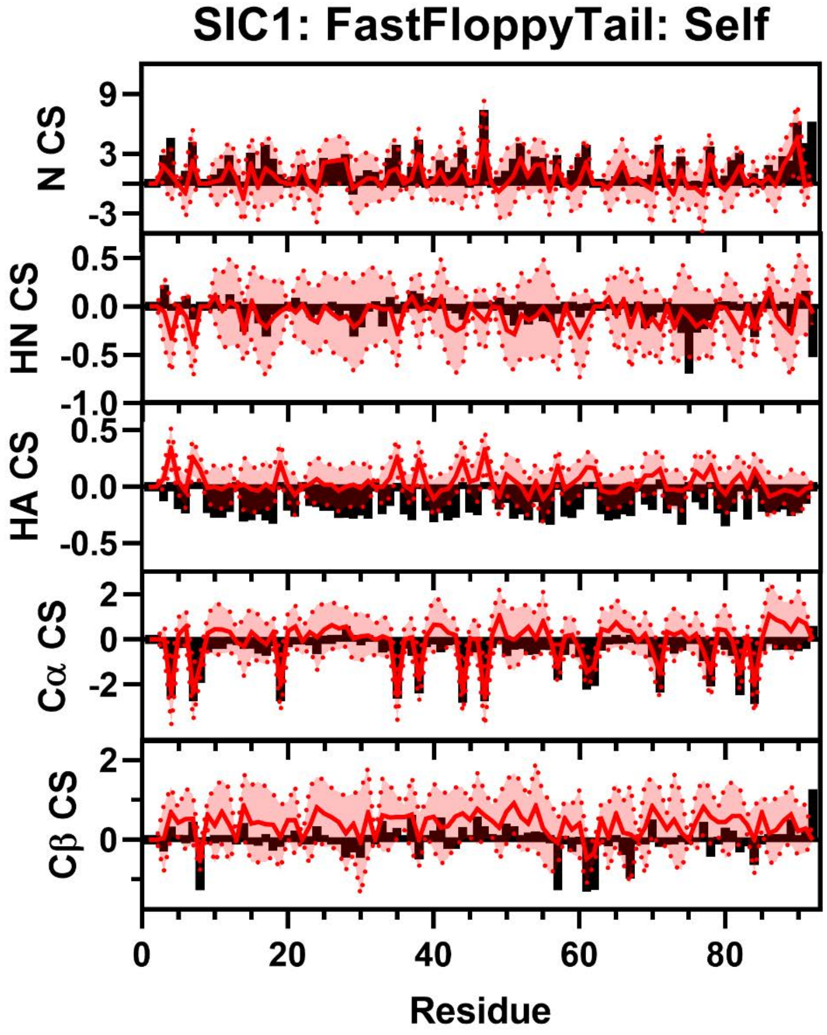


**Figure S214:** Comparison of Simulated and Experimental NMR Secondary Chemical Shift Data for SIC1. Simulated Chemical Shift values from the FastFloppyTail ensemble using best-reweighted fragment selection (red) overlayed on experimental data (black bars) of N (Row 1), HN (Row 2), HA (Row 3), Cα (Row 4), and Cβ (Row 5) chemical shifts from Mittag *et al*. [34] Neighbor corrected random coil chemical shift values from Sparta+ have been subtracted from both simulated and experimental chemical shifts to generate secondary chemical shift data [11].

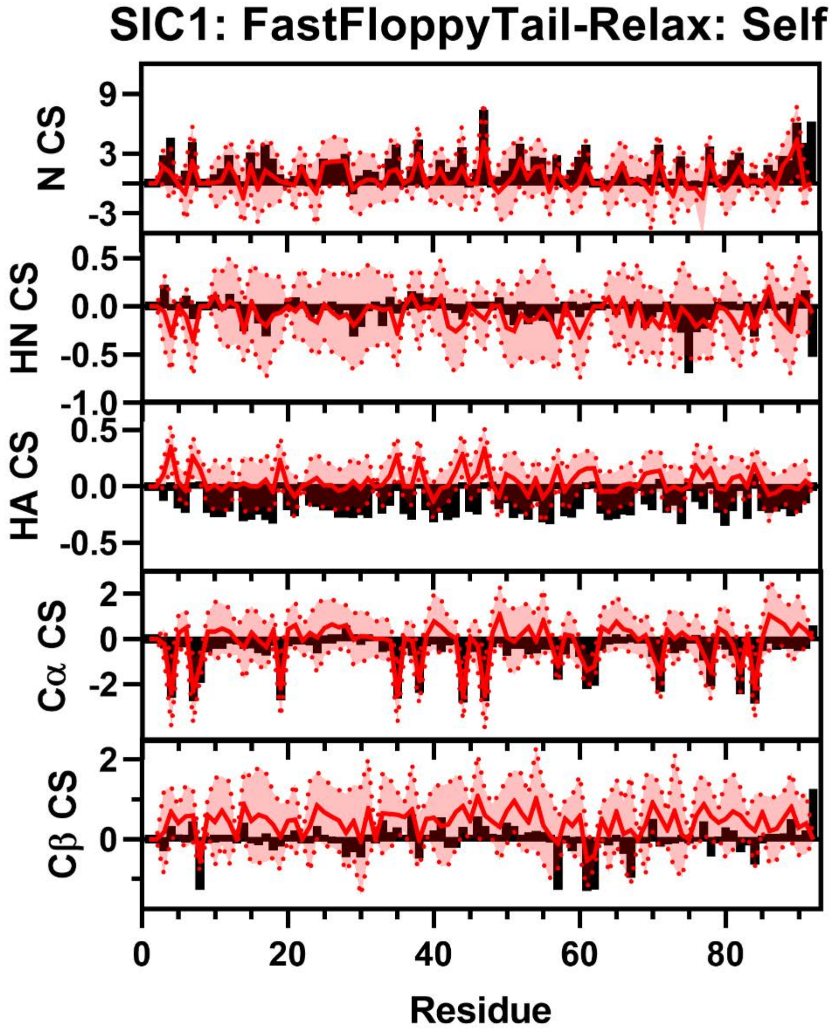


**Figure S215:** Comparison of Simulated and Experimental NMR Secondary Chemical Shift Data for SIC1. Simulated Chemical Shift values from the FastFloppyTail ensemble using best-reweighted fragment selection (red) overlayed on experimental data (black bars) of N (Row 1), HN (Row 2), HA (Row 3), Cα (Row 4), and Cβ (Row 5) chemical shifts from Mittag *et al.* [34] Neighbor corrected random coil chemical shift values from Sparta+ have been subtracted from both simulated and experimental chemical shifts to generate secondary chemical shift data [11].

### Comparisons of TAUK Data

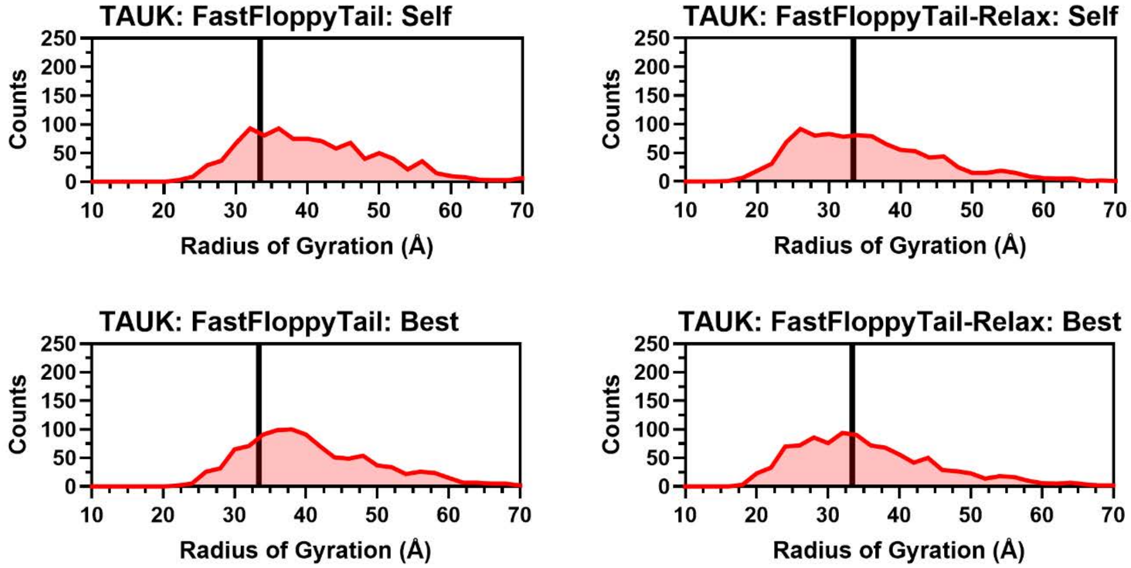


**Figure S216:** Histograms of Radii of Gyration for TAUK. Histograms of the radius of gyration from self (Top Left) and best (Top Right) reweighting in FastFloppyTail and self (Bottom Left) and best (Bottom Right) reweighting in FastFloppyTail-Relax compared to experimental values from Ozenne *et al.* (black) [35].

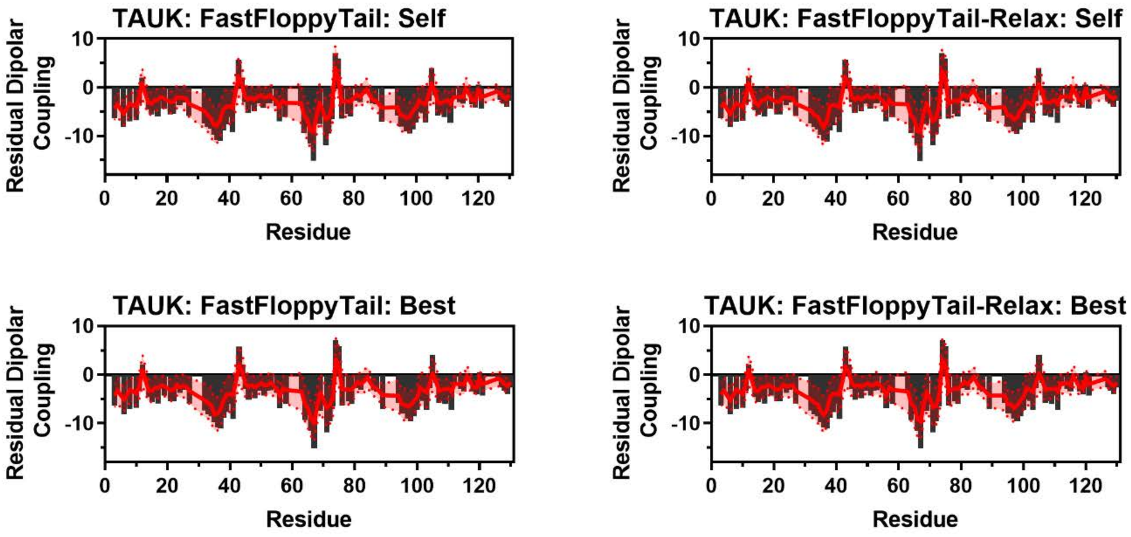


**Figure S217:** Comparison of Simulated and Experimental Residual Dipolar Coupling Data. Simulated RDC values (red line) from self (Top Left) and best (Top Right) reweighting in FastFloppyTail and self (Bottom Left) and best (Bottom Right) reweighting in FastFloppyTail-Relax overlayed on experimental data (black bars) from Ozenne *et al.* [35].

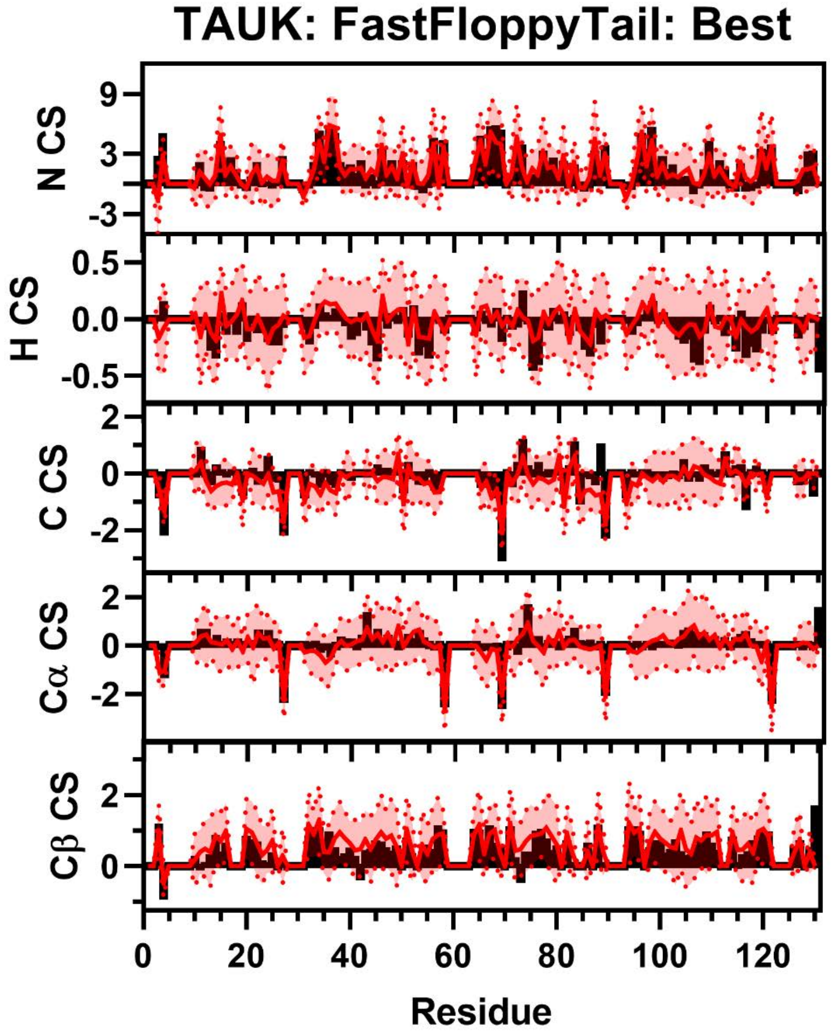


**Figure S218:** Comparison of Simulated and Experimental NMR Secondary Chemical Shift Data for TAUK. Simulated Chemical Shift values from the FastFloppyTail ensemble using best-reweighted fragment selection (red) overlayed on experimental data (black bars) of N (Row 1), H (Row 2), C (Row 3), Cα (Row 4), and Cβ (Row 5) chemical shifts from Ozenne *et al.* [35]. Neighbor corrected random coil chemical shift values from SPARTA+ have been subtracted from both simulated and experimental chemical shifts to generate secondary chemical shift data [11].

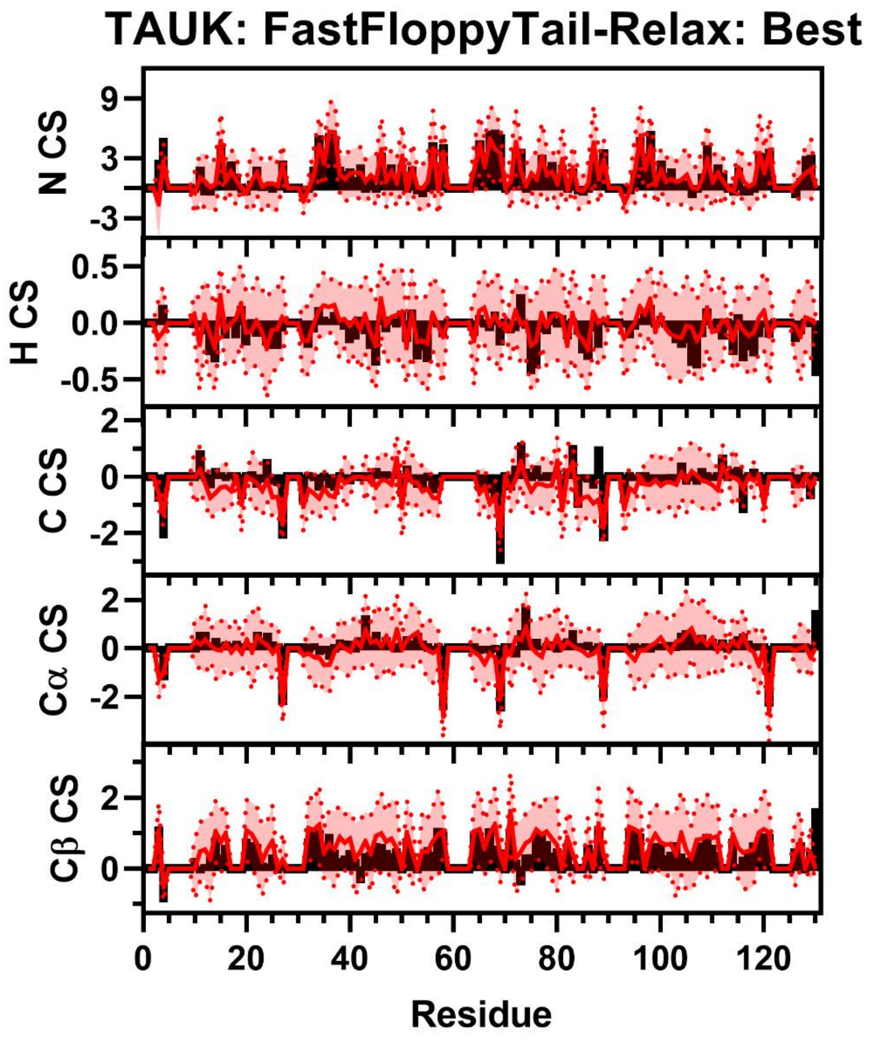


**Figure S219:** Comparison of Simulated and Experimental NMR Secondary Chemical Shift Data for TAUK. Simulated Chemical Shift values from the FastFloppyTail-Relax ensemble using best-reweighted fragment selection (red) overlayed on experimental data (black bars) of N (Row 1), H (Row 2), C (Row 3), Cα (Row 4), and Cβ (Row 5) chemical shifts from Ozenne *et al*. [35] Neighbor corrected random coil chemical shift values from SPARTA+ have been subtracted from both simulated and experimental chemical shifts to generate secondary chemical shift data [11].

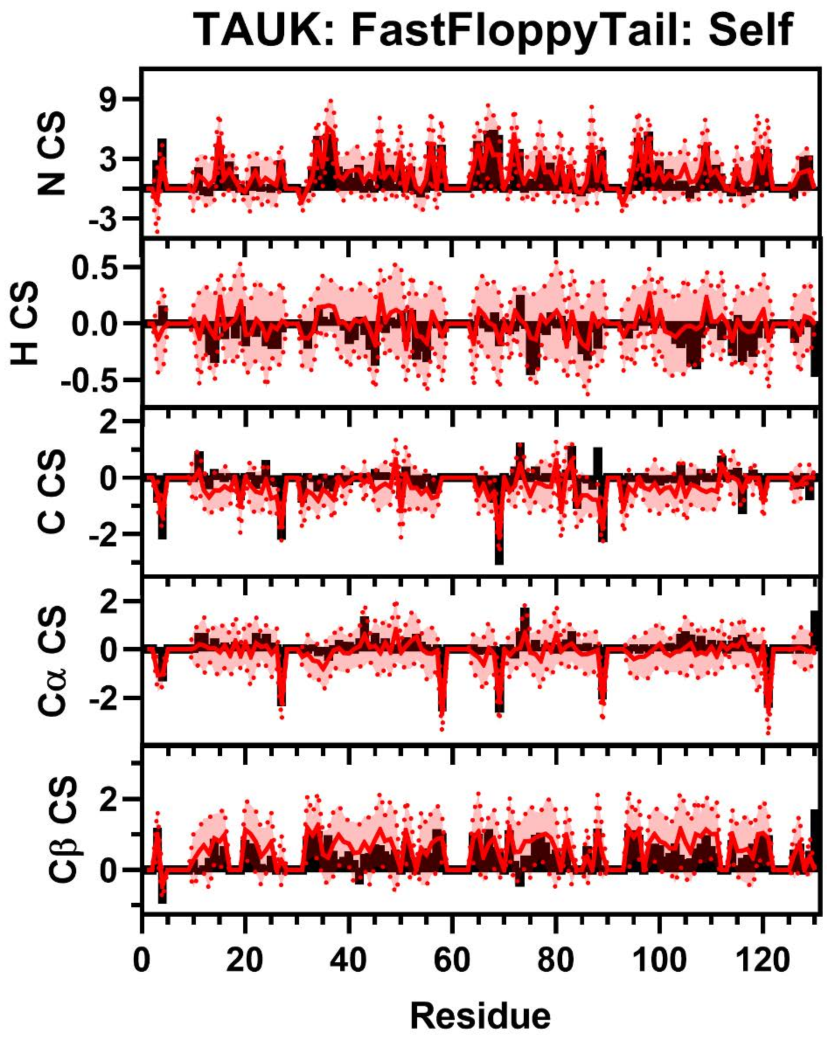


**Figure S220:** Comparison of Simulated and Experimental NMR Secondary Chemical Shift Data for TAUK. Simulated Chemical Shift values from the FastFloppyTail ensemble using self-reweighted fragment selection (red) overlayed on experimental data (black bars) of N (Row 1), H (Row 2), C (Row 3), Cα (Row 4), and Cβ (Row 5) chemical shifts from Ozenne *et al*. [35] Neighbor corrected random coil chemical shift values from SPARTA+ have been subtracted from both simulated and experimental chemical shifts to generate secondary chemical shift data [11].

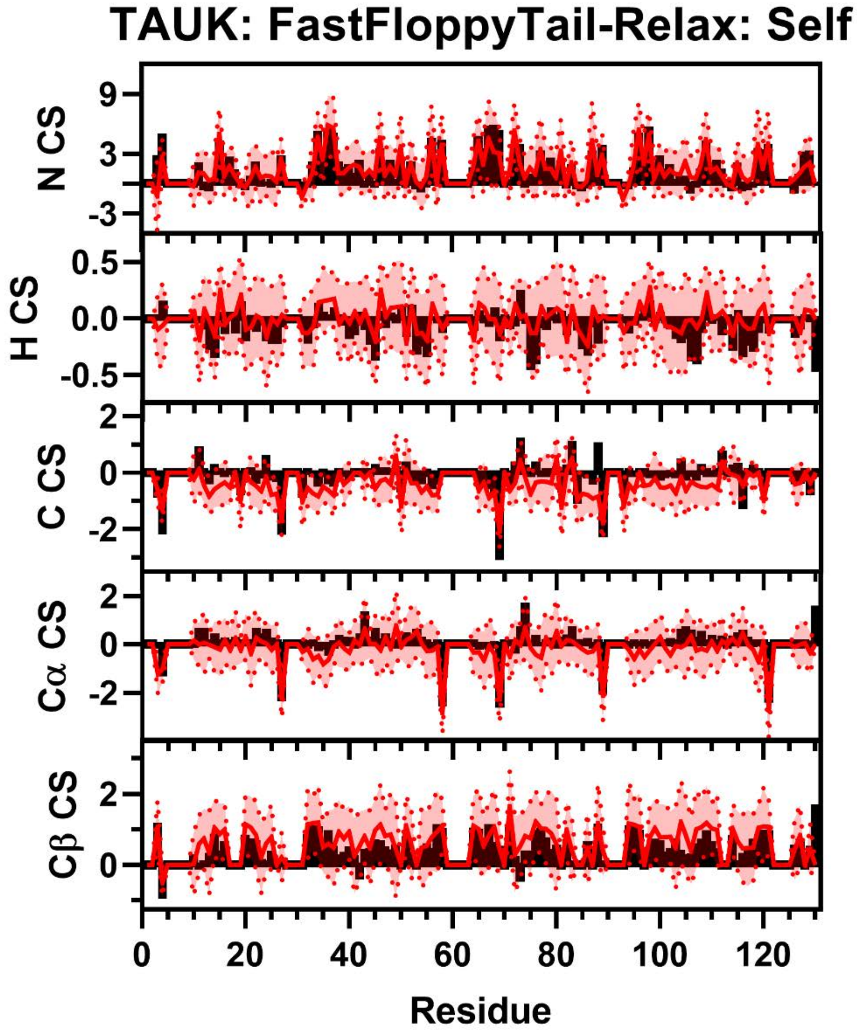


**Figure S221:** Comparison of Simulated and Experimental NMR Secondary Chemical Shift Data for TAUK. Simulated Chemical Shift values from the FastFloppyTail-Relax ensemble using self-reweighted fragment selection (red) overlayed on experimental data (black bars) of N (Row 1), H (Row 2), C (Row 3), Cα (Row 4), and Cβ (Row 5) chemical shifts from Ozenne *et al*. [35] Neighbor corrected random coil chemical shift values from SPARTA+ have been subtracted from both simulated and experimental chemical shifts to generate secondary chemical shift data [11].
